# On the utilisation and characterisation of external biotransformation systems in *in vitro* toxicology: a critical review of the scientific literature with guidance recommendations

**DOI:** 10.1101/2025.09.23.677684

**Authors:** Sebastian Lungu-Mitea, Matilda Stein Åslund, Inska Reichstein, Felipe Augusto Pinto-Vidal, Andreas Schiwy, Henner Hollert, Miriam N Jacobs, Klára Hilscherová

**Affiliations:** Masaryk University, Faculty of Science, RECETOX, Brno, Czech Republic; Uppsala University, Department of Pharmaceutical Biosciences, Toxicology and Drug Safety, Sweden; Linnaeus University, Department of Biology and Environmental Science, Kalmar, Sweden; Swedish University of Agricultural Sciences, Department of Forest Mycology and Plant Pathology, Uppsala, Sweden; Goethe University Frankfurt, Department of Evolutionary Ecology and Environmental Toxicology, Frankfurt am Main, Germany; Fraunhofer Institute for Molecular Biology and Applied Ecology (FhG-IME), Department of Environmental Media Related Ecotoxicology, Schmallenberg, Germany; UK Health Security Agency, Harwell Science and Innovation Campus, Chilton, OXON, UK

**Keywords:** biotransformation, S9, microsomes, *in vitro*, critical review, endocrine disruption, mutagenicity, genotoxicity

## Abstract

Incorporating biotransformation capabilities into *in vitro* assays represents one of the most critical challenges in toxicology, facilitating the transition from *in vivo* models to integrated *in vitro* strategies. Although emerging technologies show promise, their current limitations in scalability hinder high-throughput applications. In the short to mid-term, externally added biotransformation systems (“BTS”: S9 and microsomal liver fractions) used together with *in vitro* assays offer viable alternatives. However, despite over fifty years of use, BTS are marred by reproducibility issues, raising concerns about their reliability, and raising the question: Are BTS inherently unreliable, or has their reputation been flawed by methodological oversights?

This review critically evaluates BTS’ methodological rigour, applying a deep statistical analysis of the scientific literature. We employed Boolean operator searches across scientific literature repositories to curate a database on BTS research in conjunction with relevant *in vitro* assays, focusing on endocrine disruption, mutagenicity, and genotoxicity endpoints. Through systematic searches, screening, and eligibility criteria, we identified 229 bibliographic records. Data parameterisation and extraction were conducted across 24 domains of BTS relevance and reliability. Methodological reporting rigour was assessed via scoring (reported vs. non-reported data items) and revealed a lack of reproducible standards. Numerical measures associated with principal BTS reaction components were subjected to meta-regression analyses. No statistically significant correlations were found for BTS and related cofactor concentration-response relationships or time-related elements. Finally, descriptive statistics, multiple correspondence analysis, and *Apriori* algorithm-based relational networks identified qualitative patterns of methodological robustness and deficiencies.

In conclusion, these results emphasise shortcomings across the scientific literature in complying with appropriate methodological reporting. We offer evidence-based recommendations, in the form of a conceptual regulatory guidance framework, to enhance research practices, quality, and reproducibility of BTS applications; designed to strengthen the robustness of BTS research and its integration into regulatory-relevant hazard and risk assessment of chemicals.

**Graphical abstract:** 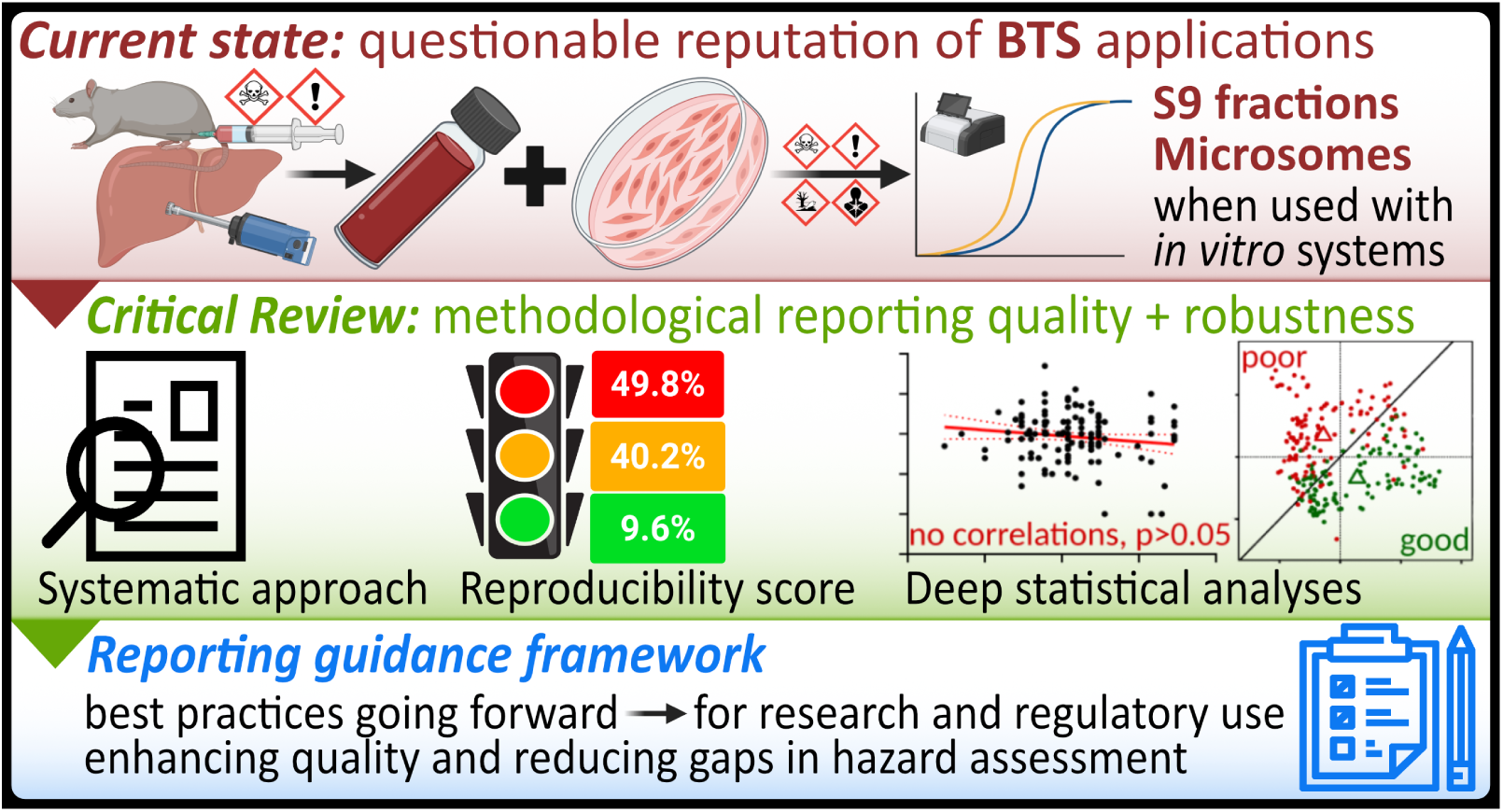

## 1. Introduction

### 1.1 Rationale

Over the past two decades, the focus of toxicological chemical hazard assessment has transitioned from relying primarily on *in vivo* animal lethality-based endpoints to embracing a mode of action and mechanistic methodology ^1–4^, whereby incorporating alternatives to *in vivo* animal testing, with the intention of ultimately replacing animal testing, has now become central to regulatory toxicology. Current European and national strategies aimed at reducing the risks posed by hazardous chemicals to health and the environment seek to minimise animal testing and implement strategies for next-generation risk assessment. In 2023, the EU Commission published a roadmap to achieve this ^5^. Whilst *in vitro* test methods can be utilised to shed light on toxicological mechanisms, the inclusion of metabolism into *in vitro* testing strategies is necessary, as the *in vitro* methods are not a one-to-one replacement for the *in vivo* animal test methods. Combinations of *in vitro* test methods are needed to address endpoints of concern adequately.

Furthermore, a common limitation of many *in vitro* systems, as used in regulatory safety assessment and reported in the literature, is their inadequate or degraded biotransformation capacity ^6–9^. To date, metabolising systems have been routinely applied to standardised *in vitro* test guidelines (TGs) in relation to genotoxicity and mutagenicity, the *in vitro* micronucleus test ^10^ and the bacterial reverse mutation test, or Ames test ^11^. As such, these standardised test methods are routinely requested for most industrial chemical, pharmaceutical, and agrochemical hazard testing. Further, OECD TGs do recommend that “exogenous metabolising systems should be used in cell systems without metabolic capacity”.

In the early 2000s, efforts to review and improve the metabolic status of *in vitro* systems were undertaken by the OECD ^7–9^, with short, medium, and long-term recommendations. Short to medium-term recommendations included the addition of non-cytotoxic rodent liver tissue fractions such as microsomal systems and supernatants after 9000g centrifugation of liver homogenate (S9), containing Phase 1 and Phase 2 metabolic enzymes, i.e., Cytochrome P450 (CYP450), Uridine-diphosphate-glucuronosyltransferase (UGT), and cytosolic enzymes (biotransformation systems, hereafter “BTS”).

The longer-term recommendations advocated for the development of a relevant cell system, such as the HepaRG CYP induction test method and the use of cryopreserved human hepatocytes. These have progressed, are discussed in, e.g. ^12,13^, and are not part of the work described here.

The International Council for Harmonisation of Technical Requirements for Pharmaceuticals for Human Use (ICH) has recently published the M12 Guideline, which provides specific and detailed guidance regarding the application of various hepatic *in vitro* systems and considerations regarding the application of Physiologically Based Pharmacokinetic (PBPK) computational modelling for drug interaction studies ^14^. The ICH guidance, however, is designed for the safety assessment of pharmaceuticals that will be entering clinical trials. Whilst for chemical testing and hazard assessment purposes, the objective is not to ensure the safety and efficacy of a pharmaceutical intended for human consumption, but to characterise the intrinsic hazards of the chemical as a protection and prevention measure in the design, application, risk assessment, and management of a chemical.

Other complementary research efforts include developing novel test systems incorporating biotransformation capabilities, such as 3D cell cultures and microfluidic chips, although they are not yet ready for regulatory applications or high-throughput screening (HTS) ^15,16^. The US EPA is exploring metabolically active tips to add to HTS cellular systems and has included metabolic assessment of cell line systems in Toxcast ^17,18^. Additional approaches include the stable or transient augmentation of microorganisms or cell lines via a genetic modification to include or enhance metabolic enzymes ^19–21^. Some of these approaches are not recent ^22^ and have not gained broad application or regulatory acceptance.

BTS, as they are derived from animals, represent an *ex vivo* refinement rather than a complete replacement of *in vivo* methods. Nonetheless, their utility is high, evidenced by well-established applications in biomedicine, pharmacology, toxicology, and other fields. This is supported by over a century of historical use and experience ^23,24^. Indeed, many protocols, including BTS, are available within an *in vitro* setup. Furthermore, S9 and microsomal BTS sourced from various vendors cover diverse species and configurations, including those obtained from chemically induced animals for enhanced BTS activity. (Historically, increased BTS-related metabolic capacity was induced into model test species by dietary exposure to mixtures of polychlorinated biphenyls (PCBs) known as “Aroclors”, particularly the Aroclor 1254 variant. However, following the phase-out of PCBs under the Stockholm Convention on Persistent Organic Pollutants ^25^, mixtures of β-naphthoflavone and phenobarbital (BNF/PB) have been utilised as substitutes).

BTS definition and variability are recognised as the major drawbacks for their integration into combinations with *in vitro* systems, including HTS applications. Common rodent-derived BTS often lack comprehensive characterisation of vital aspects affecting their biotransformation capacities, such as specifications of BTS preparation, BTS protein concentration, enzyme activity, and experimental conditions used in the assay ^6^. This information is essential for accurately categorising and utilising historical data. Prospectively, precise characterisation and categorisation of BTS are crucial to ensure the quality, reproducibility, and comparability of their *in vitro* applications, especially for chemical biotransformation. Current BTS manufacturing protocols are often based upon procedures that have been in long-term use ^26–28^, but the new ICH guidance ^14^ provides more contemporary quality criteria. Notably, many scientific literature studies have used self-manufactured BTS without explicitly detailing production protocols, and greater transparency in the production process is needed from commercial manufacturers to understand the inherent uncertainties when interpreting the resultant data. Awareness of variance and variability in BTS has been acknowledged for over forty years ^29–33^, and in practice, a reasonably substantial amount of data using BTS with the Ames and micronucleus test methods is included in, e.g., agrochemical study report submissions to regulatory bodies.

Considering the ultimate objective of enhancing the human and environmental relevance of *in vitro* methodologies by incorporating metabolic components and facilitating scalable, reproducible HTS methods, we have conducted a focused examination of the hazard assessment requirements for BTS to attain a regulatory-acceptable quality level. This entails ensuring adequate, consistent reporting characterisation of BTS to guarantee reliability, relevance, and reproducibility. As pointed out above, previous detailed reviews and communications ^6–8^ have pinpointed the deficiencies in BTS utilisation, recommending that these issues be addressed. Yet, these recommendations have been largely overlooked in applied research. This review critically evaluates the existing scientific literature with a focus on BTS methodological rigour by providing complex data and subsequent analysis.

### 1.2 Objectives

The overarching goal of this critical review is to evaluate the reporting standards of BTS within the existing scientific literature. This entails a meta-analysis of the relevant information on BTS and cofactor formulations, their concentrations, and reaction conditions (in the sense of chemical kinetics). Considerations for the development of a research and regulatory-relevant guidance framework to establish specific reporting criteria for BTS applications in conjunction with *in vitro* assays for chemical hazard assessment are provided. It is intended to complement the new ICH pharmaceutical guidelines ^14^. The purpose is to enhance the quality of BTS applications in *in vitro* regulatory contexts, including HTS.

Given BTS’ diverse origins and applications across a wide array of chemical compounds, this review is focused on the application of BTS to *in vitro* test systems for the detection of mutagens, genotoxicants, and endocrine active substances/disruptors, such as in, e.g., reporter-gene assays.

Next to the most commonly used rat-derived BTS, the review also encompasses human ^34^ and fish-derived BTS. The primary focus is on BTS’ implementation and methodological reporting. This is achieved by evaluating and grading three primary domains (“BTS characterisation”, “BTS reaction components”, and “BTS-specific experimental setup”) and 24 subdomains within the selected literature. Both quantitative and qualitative meta-analyses were conducted on the gathered data. Following analysis and synthesis, quality criteria are proposed to support researchers in improving their experimental designs, thereby enhancing the utility and applicability of their BTS experiments in assessing toxicological metabolic hazards.

In summary, this review:

A. Critically assesses the scientific rigour of BTS reporting standards in the published, peer-reviewed literature, focusing on the reproducibility, reliability, and robustness of the methodologies used via a grading scheme approach.
B. Deconstructs and interprets concentration-response and reaction conditions of BTS protein and cofactor concentrations as reported in the literature.
C. Investigates the extracted data for additional patterns, correlations, and identification of knowledge gaps.
D. Formulates a guidance framework for future BTS applications in chemical hazard assessment, targeting regulatory and Test Guideline applications.

## 2. Methods

The critical review follows a systematic approach by employing tools for framing, scoping, eligibility, literature search, and screening, as utilised in systematic reviews (SR ^35,36^), systematic evidence maps (SEM ^37,38^), and scoping reviews (ScR ^39^), with specific modifications towards the field of toxicology ^40–42^. Due to the nature of our investigation, we could not comply with canonical SR, SEM, or ScR outcomes ^43,44^. Instead, we conceptualised our structural framework to fit the specific needs (Fig. S1 in the supplementary manuscript (SM)). The SM provides further details in sections 2.4, 2.11, and 2.15. Previously, a review protocol was conceptualised, published, and shared among peers according to the PRISMA-P guidelines ^45,46^. The latter was uploaded on Figshare ^47^ (https://doi.org/10.6084/m9.figshare.21494616.v2).

This method section highlights the most critical aspects. For a more detailed insight into the applied methods and techniques, please refer to the SM, section 2, which presents the materials and methods in a PRISMA-aligned manner and provides further details for all sections elaborated below. All raw data and metadata are provided as additional supplementary information materials (SI) 1 to 11 and are available on Figshare (https://doi.org/10.6084/m9.figshare.28504826.v1).

This review did not include grey literature, nor JMPR and EFSA/ECHA summary reports on agrochemicals and industrial chemicals.

### 2.1 Study eligibility criteria

Limitations: Peer-reviewed, English, scientific literature. Only original scientific articles, no meta-analyses or review articles. No Limitations: Articles’ geographic origin or publication date. For more details, eligibility criteria reasoning, and explanations, see the SM, section 2.3. PICO/PECO(TS) (“population, intervention, exposure, comparator, outcomes, target conditions, study design”) criteria are described in sections 2.4 and 2.9 of the SM.

### 2.2 Information sources

Standard scientific literature repositories were employed to construct an information collection database using Boolean operator searches detailed below in section 2.3 and SM, section 2.6. These repositories include PubMed (https://pubmed.ncbi.nlm.nih.gov/), Web of Science (https://www.webofscience.com/), and Scopus (https://www.scopus.com/).

The search strategy was refined, and data item domains were coded using a “piloting” process. A subset of 73 highly relevant publications previously identified by the reviewers (see SI5 and the reference list “BTS3” in SM, section 5.4) was initially screened and utilised for this purpose. For more details on the piloting process, see SM, section 2.5. The coding of data items is detailed below in section 2.6.

### 2.3 Search strategy

The named repositories were searched using the following syntax and operators:

Search string 1: “biotransformation” AND “metab*” AND “S9” AND (“endocrine” OR “thyroid” OR “*estro*” OR “andro*”)

Search string 2: “biotransformation” AND “metab*” AND “S9” AND “cells” AND “cytotox*” AND (“genotox*” OR “mutagen*”)

The main article, the SM, and supporting supplementary information material files (SI) refer to the databases and resulting bibliographies from the search strings as “BTS1/endocrine” and “BTS2/mutagen”. The search parameters were specifically tailored to focus on BTS implementation in *in vitro* studies addressing mutagenic, genotoxic, and endocrine-disrupting endpoints, to align with the objectives outlined in section 1.2, and maintain a manageable scope of search results. More details on Boolean operator specifics, search strategy reasoning, and search timelines are given in the SM, section 2.6.

### 2.4 Selection of Studies

The search results from the specified literature repositories were downloaded and imported into the Sysrev GUI (https://sysrev.com/) using PubMed identifiers (PMID). All search outputs were consolidated into a single database within Sysrev. Four independent reviewers employed this platform for article selection using an article sequence randomisation approach. The reviewers screened titles and abstracts based on the pre-established eligibility criteria in section 2.1 and the SM, section 2.3. Article inclusion was based on reviewer consensus, with 75% agreement as a threshold. For more details on study selection, bibliographic file formatting, and duplicate removal, consult the SM, section 2.7.

### 2.5 Data collection

Post-selection, the revised bibliography underwent a second round of manual duplicate removal and full-text PDFs were retrieved.

Data extraction and evaluation adhered to the DEERS protocol (Data Extraction, Evaluation, and Reliability Schema; see SM, appendix 6.1 and section 2.11). Each article’s DEERS information extracts were classified, organised, and summarised in a Microsoft Excel spreadsheet using a predefined extraction form for each reviewer (details in SI6). A unique identification number was allocated to each article, ensuring comprehensive data collection across all primary and secondary data domains.

Two reviewers independently conducted detailed data extraction. Four independent reviewers then cross-verified and refined the entire extraction dataset. For more details on the used data formats, software, and data extraction, consult SM, section 2.8.

### 2.6 Data items and coding

The coding for data items was established through a preliminary evaluation of 73 studies (part of “BTS3/historical”) deemed highly relevant (“piloting”) while referencing prior publications on key BTS experimental parameters ^6,8,9,16,48^. This process involved discussions and consensus among four reviewers to determine the coding strategy. In total, 24 domains were identified as essential for a comprehensive evaluation of the studies’ methodology, reporting, and reliability standards (Tab. 1). Of these 24 domains, nine were consistently detailed in all piloting studies and were considered significant for assessing study relevance but not for evaluating reliability. These nine domains are referred to as “data items of relevance” and are not included in the reliability assessment (see also SM, Tab. S2, delineated as “non-critical”). The remaining 15 domains were deemed vital for methodological reliability assessment (outcome A; see also SM, Tab. S2, delineated as “critical”). The data items discussed in this section align with the criteria outlined in the DEERS protocol (refer to the SM, appendix 6.1). Methodological data item measures for four primary and 24 subdomains were extracted, as summarised in Tab. 1. For more information on the data item domain measures, consult the SM, section 2.9. Table S2, within the SM, also provides a hypothetical example of data extraction following the established coding parameters.

**Table 1:** Total data item domains extracted from the bibliography database.

| Primary domains | Subdomains |
| --- | --- |
| BTS characterisation | Origin, species, strain, pooling (and sex), husbandry, and (xeno-)metabolic induction |
| BTS reaction components | Total protein concentration, buffer system, dilution factor, primary cofactors, secondary cofactors, and used solvent |
| BTS experimental setup | Incubation period, incubation temperature, and BTS-related controls |
| Data items of relevance | Author, year, journal, test system, endpoint, experimental methodology, type of applied external BTS, exposure, and post-BTS procedure |

### 2.7 Outcomes, effect measures, and synthesis methods

We employed various effect measures and synthesis methods for each outcome based on the prioritisation hierarchy outlined in section 1.2, “Objectives”, and the SM, section 2.10, “Outcomes and prioritisation”. Each primary study outcome (A to D) is aligned with the study objectives listed in section 1.2. Outcomes A to C are assessed via the meta-analyses of extracted data item measures. In contrast, the synthesis of the latter directs outcome D (regulatory and research guidance framework) (see also Fig. S1 for the structural framework of the critical review).

For outcome A (“scoring”), which focused on assessing BTS methodological reporting standards and scientific rigour as per the DEERS protocol, data items were scored binary (reported vs. non-reported), resulting in a net score of one for each reported subdomain. This measure was applied to qualitative or quantitative data item domains (SM, Tab. S2, and SI6). We only scored critical data item domains for reliability (see SM, Tab. S2), with 16 scoring categories derived from 15 critical domains, including the bifurcated “BTS pooling” domain. Each reported data item received a positive score of 1, whereas a non-reported item received a neutral score of 0. Data was manually entered into the DEERS protocol and transferred to a Microsoft Excel sheet (SI6) for summarisation in a queryable table format database. Details on non-reported data items, visualisation of scoring outcomes, and statistical analysis are provided in SM, section 2.12.

For the secondary study outcome B (“meta-regression”), we aimed to meta-analyse quantitative data item domain measures to identify concentration-response or reaction condition-related patterns in BTS applications. The parameters related to BTS, including the incubation period, BTS protein concentration, and primary cofactor concentrations, exhibit hypothetical interdependence. For instance, an increase in BTS protein concentration can provide a reduction in incubation time.

Consequently, we hypothesised that the relationship between these parameters should be characterised by either a linear or exponential correlation. Data item measures were log-transformed, checked for the fulfilment of parametric test criteria (see SM, section 2.12), and analysed through simple linear regressions to derive adjusted R^2^ values for model accuracy. Pearson’s correlation was used to determine the coefficients of correlation and determination. Additionally, paired comparisons were employed where quantitative data item measures were matched up from the exact origin of the article record. We conducted computations, statistical analyses, and graphical plotting in GraphPad Prism 8. For more methodological details, consult the SM, section 2.12.

As a tertiary study outcome, outcome C (“mapping”) involved the exploratory mapping of all qualitative data item (sub-)domain measures. We simplified these domain measures for machine readability (see SM, Tab. S4, and SI9) and analysed them using descriptive and summary statistics, multiple factor correspondence analyses (MCA), and association networks (*Apriori* algorithms). For brevity, the following paragraphs summarise the utilised mapping analyses. More details are provided in SM, section 2.12.

Descriptive and summary statistics, including histograms, Euler plots, and Upset plots, were generated in R Studio ^49^ and R software version 4.1.2 (R Core Team, 2023), utilising the R packages ggplot2, eulerr, and UpSetR ^51–53^.

MCA was performed using FactoMineR ^54^ in R Studio. The analysis included all qualitative data item subdomains from the simplified coding dataset (see SM, Tab. S4, and SI9) as active variables, which were transformed into a binary information matrix (for details see SM, section 2.12, and SI9), except for “publication year” (grouped in five-year intervals), “publication group” (extraction datasets BTS1 to 3), and total scores, which were defined as qualitative supplementary variables and did not influence the analysis mathematically. Visualisation in ggplot2 ^53^ was restricted to levels with more than five relative observations, with data ellipses representing normal probability contours at a 0.9 confidence level.

Lastly, for data association and relational networks, we utilised association rule mining with the *Apriori* algorithm in the arules package ^55^ in R Studio to identify relational networks among qualitative data item subdomains in the simplified coding dataset (see SM, Tab. S4, and SI9). This process involved setting a support threshold of 10% and a confidence threshold of 80%, with visualisation aided by the arulesviz package ^56^.

Data item subdomains that exhibited robustness markers, either positively or negatively (see SM, Tab. S9), in both MCA and *Apriori* frameworks, were further analysed (confirmatory analyses ^57^) to evaluate the effect sizes in different scoring subpopulations. For groups comprising two scoring subpopulations, non-parametric, two-sided Mann-Whitney-U tests were employed (alpha level = 0.05). In multiple comparisons, the Kruskal-Wallis test was utilised as the primary analytical tool, followed by Dunn’s post-hoc test (alpha level = 0.05). All mentioned tests were performed with GraphPad Prism 8.

## 3. Results and Discussion

Our comprehensive analysis, detailed in the subsequent sections and further elaborated in the supplementary manuscript (SM), revealed that the literature generally fell short of reaching adequate methodological reporting quality that would assure reproducibility.

In the following sections, we outline the screening and selection processes employed to retrieve records, which were first evaluated for methodological soundness via the DEERS framework (objective to outcome A). Additionally, extracted data item measures were subjected to meta-analyses. Quantitative data item measures were further explored in meta-regression (objective to outcome B). Following a process of coding simplification (SM, Table S4 and SI9), qualitative data item measures were scrutinised iteratively via multivariate statistics (Multiple Correspondence Analysis, MCA) and machine-learning, relational networks (*Apriori* algorithms) to identify patterns of methodological robustness and shortcomings (objective to outcome C).

Discussions pertinent to the findings are presented immediately following their introduction. Section 4 integrates these results and discussions, synthesising the insights obtained into a coherent body of knowledge. This synthesis is then directed towards the development of a research and regulatory guidance framework (objective to outcome D), detailed in section 5.

### 3.1 Search, screening, and selection processes

We retrieved 229 publications from the comprehensive Boolean operator search and screening processes (Fig. 1). For initial piloting, including data item domain coding and refining search strings, we utilised 73 highly relevant publications previously known to the authors. After cross-referencing, this database also contributed 101 publications to the BTS3 database (“historical”). The Boolean searches yielded 39 publications in BTS1 (“endocrine”) and 89 in BTS2 (“mutagen”). This process was repeated twice, initially after three weeks and again after ten months. The first repetition yielded no new outcomes, while the second added two eligible publications ^58,59^, which were not included retrospectively in our assessment.

**Fig. 1:**
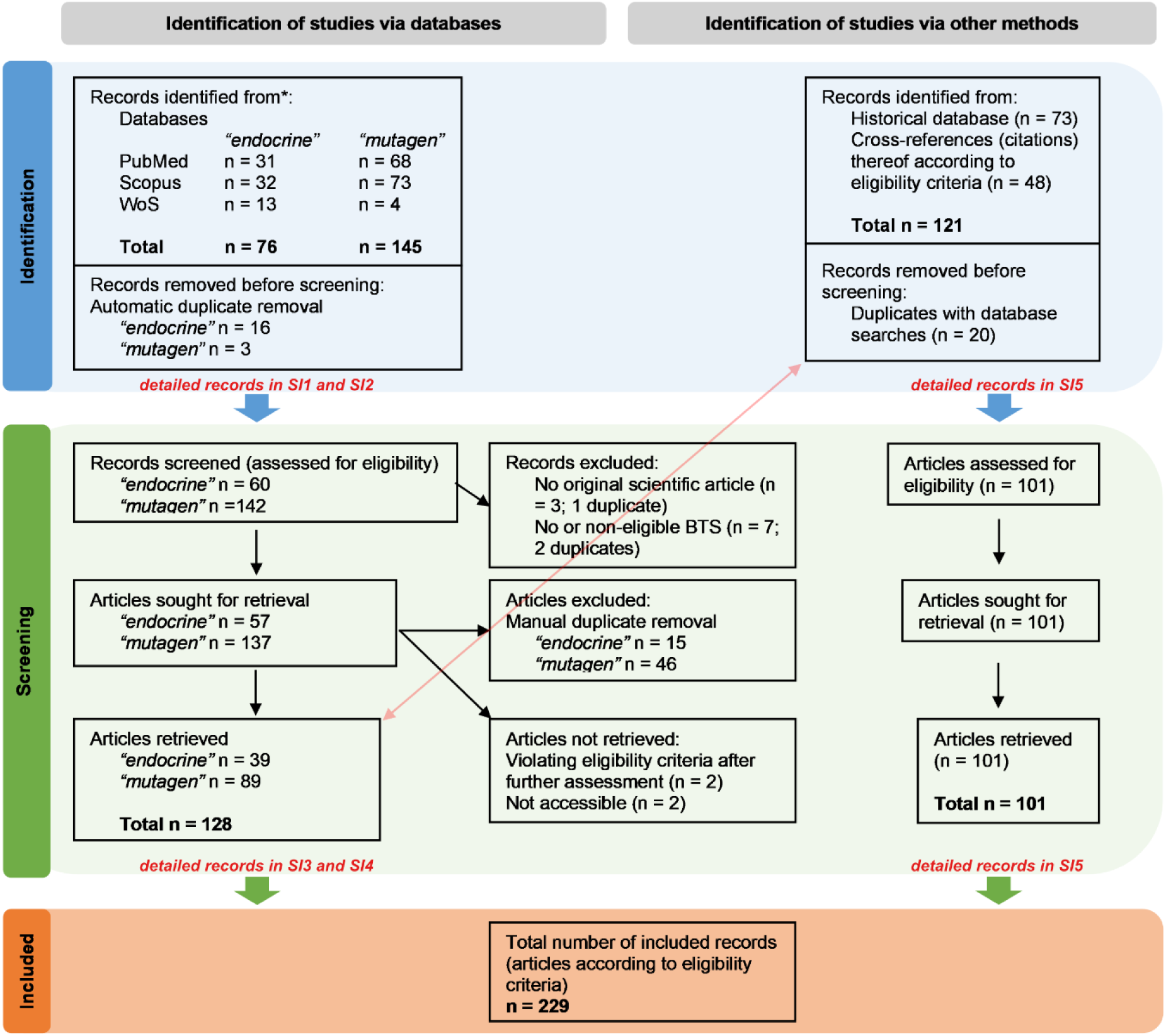
Flow diagram of search and selection processes. The chart was designed according to PRISMA guidance ^45,46^. Detailed records are given in the supplementary information material files (SI3 to SI6). Section 3.1 of the supplementary manuscript (SM) outlines reasons for record exclusion according to the eligibility criteria.

The SM, section 3.1, provides a more thorough discussion of the screening and selection processes, including the naming and reasoning of record exclusion according to the study eligibility criteria, as depicted in Fig. 1. Study characteristics and citations for each bibliographical database (BTS1 to 3) are provided in SI3 to SI6, and separate bibliographical database reference lists are available in the SM, section 5.

### 3.2 Methodological reliability assessment and scoring (DEERS, outcome A)

Within the DEERS protocol (detailed in the SM, appendix 6.1 and section 2.11), we coded data item domains guided by specific reviews ^6,7,9,16^. We transferred these to a queryable wide-range format table in Microsoft Excel (SI6), serving as the primary database. Data item measures were incorporated into the extraction form (SI6), and a reliability assessment of methodological rigour was performed, as outlined in section 2.7, utilising the DEERS scoring approach. Fig. 2A displays the hierarchical absolute scores for each article, ordered from lowest to highest. Articles are identified by their assigned ID numbers, with more detailed identification available in SI6. Bibliographic identifiers were excluded from the main article to maintain professional courtesy and acknowledge the context of historical scientific development.

**Fig. 2:**
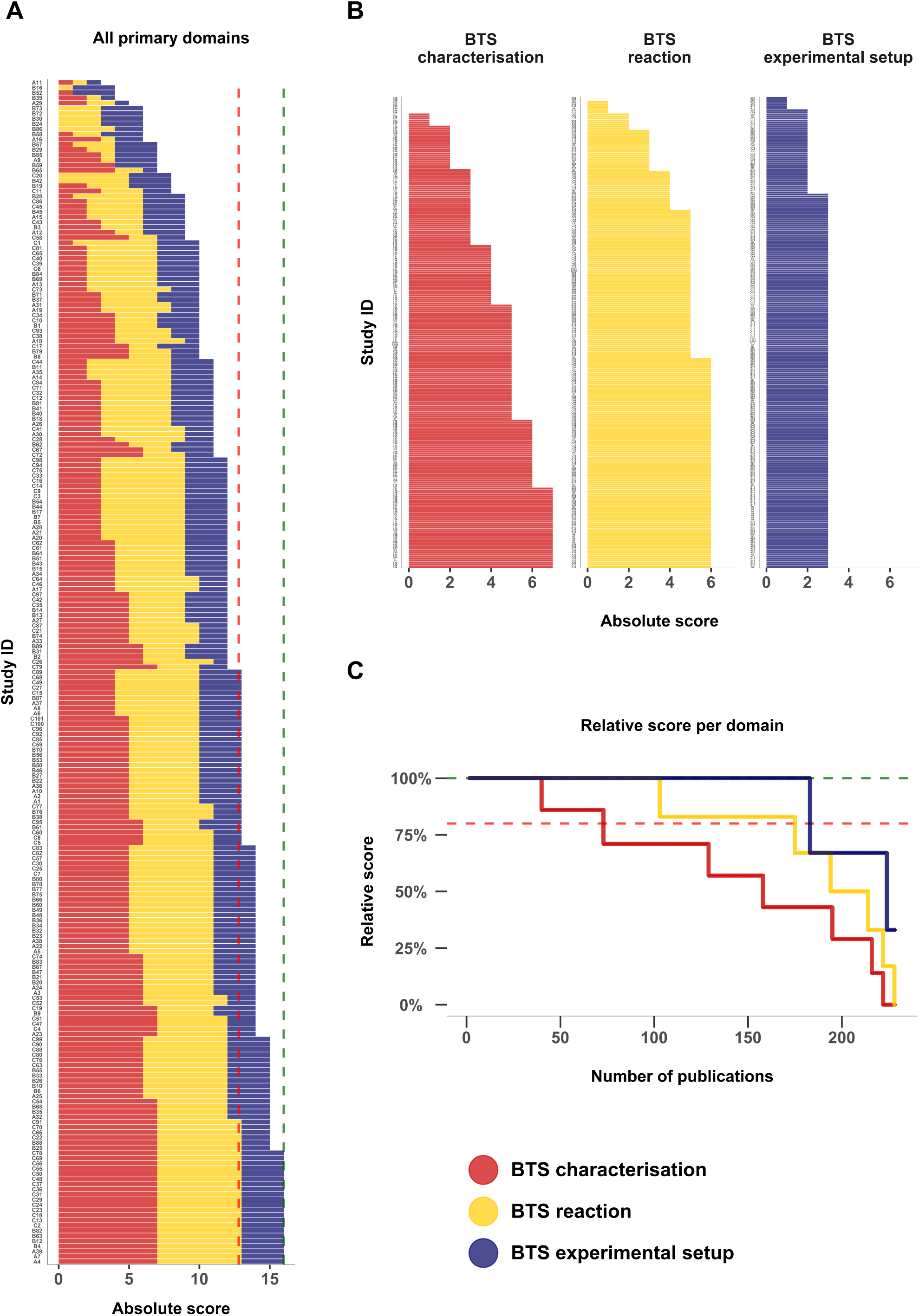
Scoring values for all n = 229 assessed studies via DEERS. (A) Total hierarchical absolute scores for all set studies by primary data item domains; “BTS characterisation” (red, max. score = 7), “BTS reaction” (yellow, max. score = 6), and “BTS experimental setup” (blue, max. score = 3). The red dotted line indicates a threshold of tolerable quality acceptance of 80%. Only a 100% score is considered fully reproducible and robust; see the green dotted line. (B) Separate absolute scores for the primary data item domains “BTS characterisation”, “BTS reaction”, and “BTS experimental setup”. (C) Relative scores for all primary data item domains for comparison. For more details, consult SI7. Refer to Tab. 1 for respective subdomains.

Our non-weighted scoring for data item (sub-)domains showed that only 22 publications (9.6% of total records) fully met reproducibility standards. The scoring criteria were conceptualised so that only studies reaching 100% reporting status were considered fully reproducible, whereas an overall score of approximately 80% (translating to 13 absolute points) was considered a lower boundary. This 80% threshold was established on the rationale that studies scoring within this range could likely attain full reproducibility with minor modifications in their methodological descriptions, as detailed in sections 4 and 5. Among the evaluated studies, a total of 115 articles attained scores of 13 points or higher. However, 114 articles (49.8% of the total evaluated studies) fell below this threshold, demonstrating a concerning level of insufficient BTS reporting quality. This indicates that half of the analysed studies were considered not to be improvable and easily rectifiable in terms of their reproducibility.

Particularly, the “BTS characterisation” primary domain exhibited the lowest scores, with only 39 studies achieving full marks and 72 surpassing the threshold (Fig. 2B, C). For “BTS reaction”, 102 studies reached a full score, and 174 exceeded the threshold. The “BTS experimental setup” domain exhibited the highest reporting standards, with 182 studies attaining a full score and an equal number surpassing the threshold (Fig. 2B, C). Within the primary domain “BTS characterisation”, especially the secondary data item subdomains “strain” (69% relative score), “pooling – sex” (57%), “pooling – number of individuals” (22%), and “husbandry details” (33%) were poorly reported (see SM, Tab. S5). Additionally, within the primary domain of “BTS reaction”, the subdomain “BTS protein concentration” also scored poorly (57%). Tabs. S5 and S6 in the SM highlight areas where reporting standards should be namely improved. In contrast, subdomains within the “BTS experimental setup” primary domain were comparatively well-documented, with 87 to 95% relative scoring correspondence observed (see SM, Tab. S7). Readers are referred to the SM (section 3.2 and Tabs. S5 to S7) for in-depth analysis and specific scoring of subdomains. Potential quality improvement measures are addressed below in Section 4.

Boxplots were generated for each primary data item domain to enhance the visualisation and comparison of scoring distributions across all assessed articles. These boxplots illustrate the relative scoring, facilitating comparisons between search strategies and databases (see Fig. 3 with summary statistics detailed in the SM, Tab. S8). The overall dataset’s sub-threshold mean scoring value (76%, Fig. 3D and Table S8 in SM) further underscores concerns about methodology reporting. The means (74 and 73%) and medians (both 75%) of Boolean search-derived datasets (BTS1 and 2) scored below the threshold (Figs. 3A, 3B). The overall median reached 81% due to higher methodological robustness within BTS3 (Fig. 3C and Fig. S2 in SM for statistics). Likewise, we encountered positive methodological robustness markers associated with the BTS3 dataset within the iterative mapping analyses (outcome C, section 3.4 below). It can be argued that BTS3 encounters some selection bias, given that it is originally based on a set of 73 publications known to the authors and deemed highly relevant. Accordingly, the methodological robustness might be inherently higher, given their prominence. Details on potential biases are elaborated in the SM, section 4.1.

**Fig. 3:**
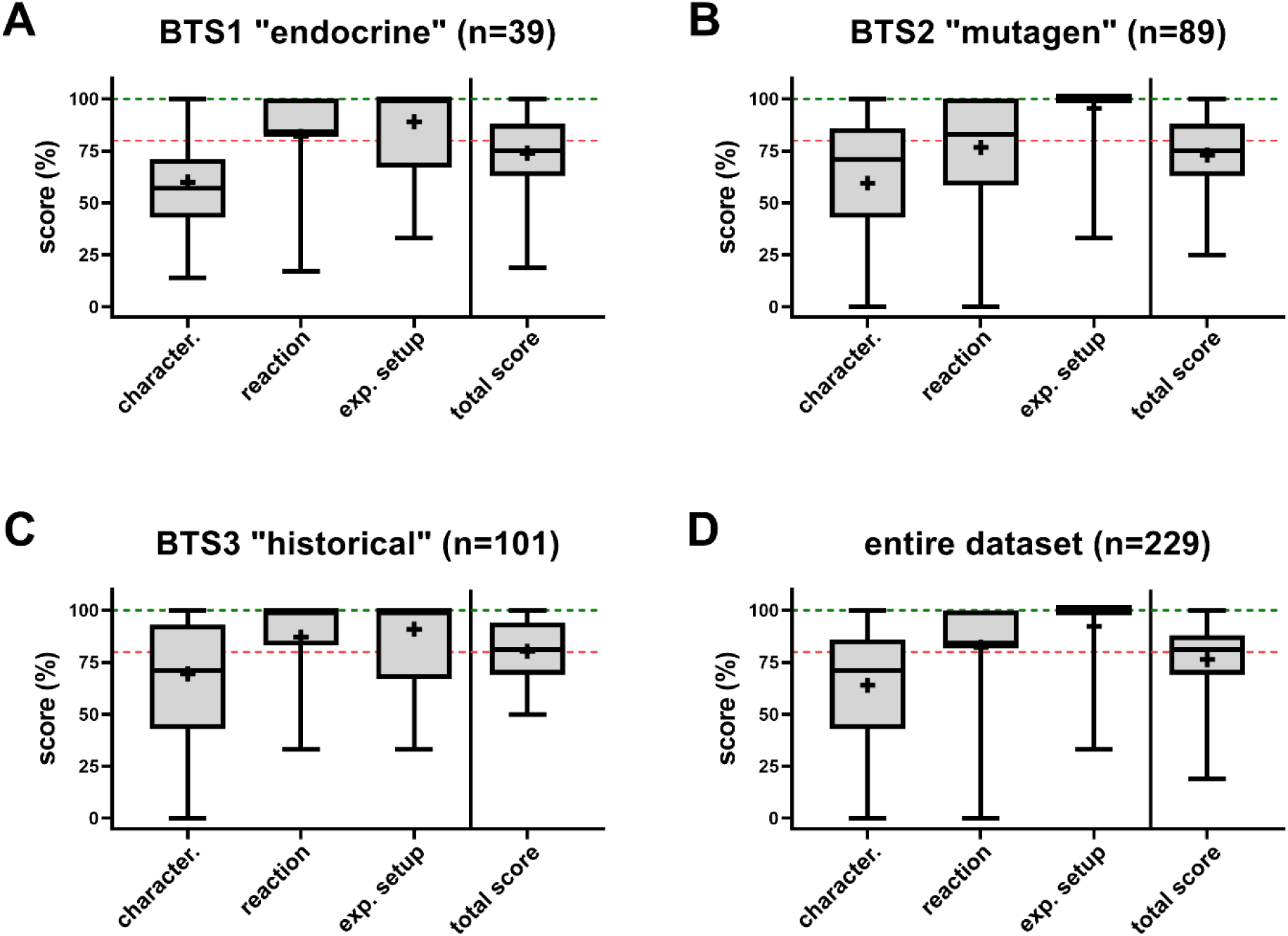
Boxplots depicting relative scoring populations of reviewed and assessed articles for all primary data item domains. Scoring distribution is given for every dataset as defined by the search strategies (panels A, B, and C) and all datasets together (D). Whiskers indicate the upper (max.) to lower (min.) boundaries. Boxes indicate the 75^th^ and 25^th^ percentiles, and the in-between line represents the median. Crosses represent mean scores. The upper green dotted lines indicate full data reproducibility and robustness, and the lower red dotted lines represent 80% reporting quality thresholds. The number of included articles within every dataset is given in the titles.

In summary, to date, the overall body of scientific literature has poor methodological robustness. In most cases, it is not possible to reproduce or adapt for follow-up applications (e.g., *in silico* biotransformation models ^60,61^).

### 3.3 Meta-regression of quantitative BTS reaction components (outcome B)

In the meta-regression analyses, we focused on the numerical data item measures relating to BTS protein concentrations, primary cofactor concentrations, and BTS incubation periods. Given the hypothesised linear or exponential relationships between these data items, we investigated the potential statistical correlations between BTS protein concentration and incubation period (Fig. 4A), primary cofactor concentration and incubation period (Fig. 4B), and BTS protein concentration and primary cofactor concentration (Fig. 4C).

**Fig. 4:**
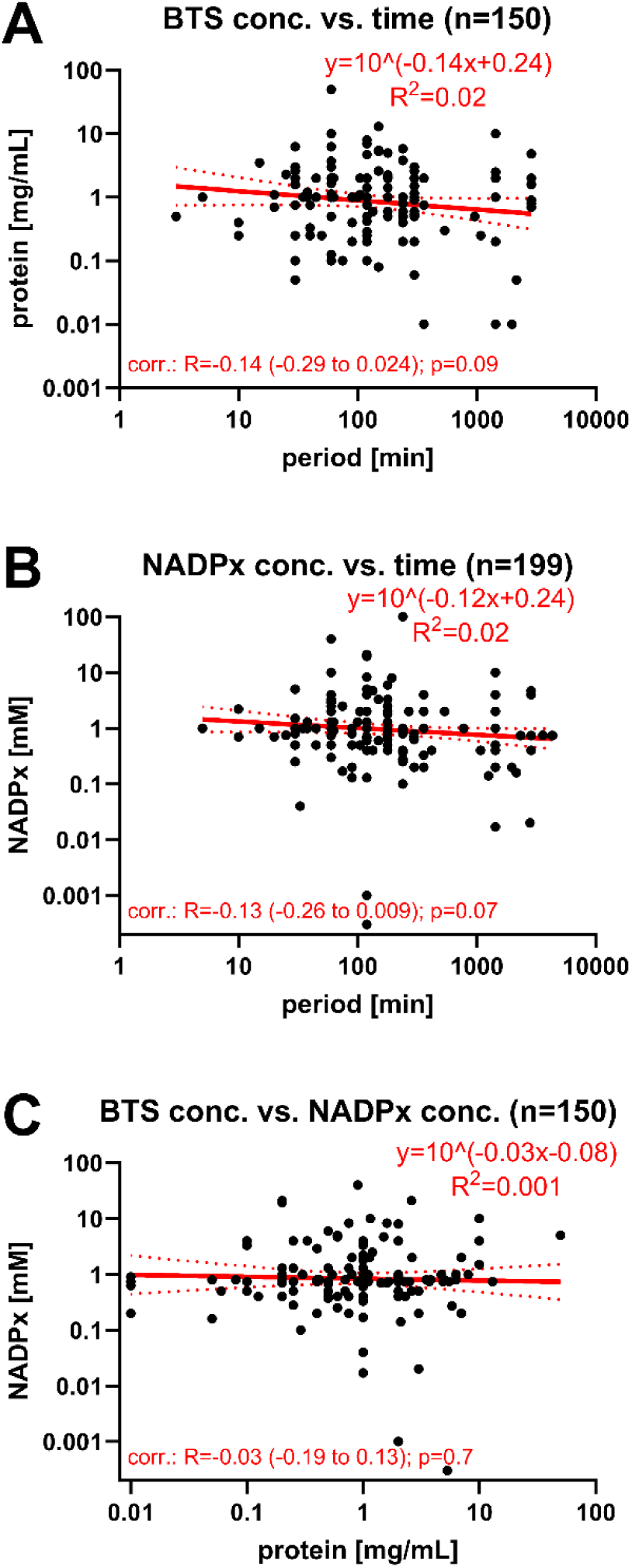
Simple linear regression fits of numerical data item measures hypothesised to show a particular mathematical relationship. Data were extracted from the overall database (SI6) and manually curated as outlined in section 2.7, with detailed seminal populations presented in SI8. Numerical data were log-log transformed before fitting a simple linear regression. Equations and fits (R^2^ values) are given within the respective graphs. Single measures are illustrated as dots. The seminal populations are provided within the titles. The regression fits are displayed as red lines, with 95% CIs as dotted red lines. Pearson’s correlation was computed between variables, the respective coefficients of correlation (R), coefficients of determination (R^2^, same as for linear fit), and 95% CIs are given within the graphs. P-values determine the statistical significance of correlation. The following pairs were tested: (A) BTS protein concentration vs. BTS incubation period; (B) primary cofactor concentration vs. BTS incubation period; (C) BTS protein concentration vs. primary cofactor concentration. Additional paired or selected comparisons (without regeneration systems) are given in the SM, section 3.3, Fig. S3.

Additionally, to explore the potential kinetic and dynamic impacts of cofactor regeneration systems, regression analyses were explicitly conducted for paired analyses (measures taken from the same publication record) and BTS setups that employed only NADPH, without a cofactor regeneration system (see SM, Fig. S3 and section 3.3). Within the graphs, “NADPx” refers to the various forms of nicotinamide adenine dinucleotide phosphate used as reducing agents in BTS reactions ^62,63^. Thereby, NADPx is either applied directly in its reduced state (NADPH) or its oxidised form (NADP+), including a redox-cycling regeneration system. While adding NADPH directly is the more straightforward approach, integrating NADP+ with a regeneration system is both more economical and ensures stable reducing agent concentration throughout prolonged incubation times.

Our analyses revealed no significant statistical relationships among the investigated quantitative measures – neither for overall (Fig. 4), paired (Fig. S3A and C), nor selected (without a cofactor regeneration system, Fig. S3B and C) comparisons. The goodness-of-fit (R^2^) values were consistently low, ranging from 0.001 to 0.2, and the regression lines were predominantly horizontal. The correlation coefficients (R) hovered around zero (−0.14 to 0.13), with 95% confidence intervals (CIs) spanning both positive and negative values, indicating a lack of directional correlation. The coefficients of determination (R^2^) varied between 0.1 to 2%, reflecting minimal variance in one variable being explained by the variance in another. Furthermore, the p-values ranged from 0.07 to 0.7, indicating insufficient evidence to reject the null hypothesis of no correlation between the variables.

According to the analyses, it is uncertain if proper assessments of concentration-response and reaction condition components of BTS-induced biotransformation processes have ever been thoroughly conducted throughout the extensive literature. Instead, component selection seems influenced by historical protocols ^26–28^ and has been continuously pursued, or the selection happened in a rather *ad hoc* manner, as demonstrated by numerical BTS parameters ranging over several orders of magnitude without correlative trajectories (Fig. 4 and Fig. S3, SM). The inclusion of all reaction parameters (substrate concentration, BTS protein concentration, and cofactor concentration at different levels) is unrealistic to test and assess, if not the prime target of the investigation, due to time and cost limits. Only one study ^64^ comprehensively set these parameters using a “Doehlert” experimental design matrix.

In conclusion, the meta-regression analyses suggest a lack of significant correlations between the key quantitative components of the BTS reaction. This indicates the need for more detailed investigations into BTS reaction dynamics to set appropriate standards.

### 3.4 Iterative mapping analyses of qualitative data items (outcome C)

After further coding simplification (detailed in SM, Tab. S4, and SI9), qualitative data item measures were made machine-readable and forwarded to data mapping analyses. As such, the data was subjected to descriptive statistics (represented as histograms, Euler plots, and Upset plots), multiple correspondence analysis (MCA), and *Apriori* algorithms (for data association and relational networks). The results are provided and discussed in the following sections.

#### 3.4.1 Descriptive statistics of qualitative data item subdomains

Descriptive statistics of qualitative data item subdomains (more details and illustrations in the SM, section 3.4) revealed evolving dynamic trends over time, including shifts in publication focus (see SM, Fig. S4), diversification of study endpoints (SM, Fig. S6), and changes in the species used for BTS (SM, Figs. S9 and S10). A notable coincidence was observed between external sourcing of BTS (SM, Fig. S8) and a decline in reporting accuracy for certain subdomains (“strain”, “BTS pooling”, and “BTS induction”, SM, Figs. S11, S12, and S14).

Another pattern identified within the descriptive statistics analyses relates to the primary cofactor (“NADPx”) regeneration systems ^62,63^. Noteworthy, oxidised NADP+ is utilised alongside glucose-6-phosphate (“G6P”) or isocitrate (“iso.”) for redox-cycling regeneration systems. Our analysis primarily focused on identifying whether these regeneration systems were mentioned. Detailed components of these systems, particularly dehydrogenases (“dh”), were not exhaustively scored to avoid overcomplicating the assessment. However, it is critical to note that for a reproducible system, both the presence and concentrations of applied dehydrogenases should be reported. Of 177 studies using cofactor regeneration systems, only 35 studies provided comprehensive details on the essential dehydrogenase components (see SM, Figs. S16 to S18).

We also encountered inconsistencies in the literature regarding the use of cofactors for phase 1 or 2 biotransformation studies. Several studies allegedly examining phase 2 reactions predominantly utilised phase 1 inducing systems (NADPH or NADP+ with regeneration systems). Conversely, some research focusing on phase 1 metabolism incorporated phase 2 specific cofactors (e.g., GSH) into the BTS mixture. Furthermore, there were instances where studies claimed to explore specific phase 2 reactions that are impractical to assess in mitochondrial fractions (microsomes). With respect to the intricacies of cofactor utilisation, the reader is directed to relevant literature, e.g., ^14,65,66^. In conclusion, there is a pressing need for substantial enhancements in reporting primary cofactors and their regeneration systems to meet adequate quality for BTS applications.

#### 3.4.2 Multiple correspondence analyses (MCA) of qualitative data item subdomains

Qualitative data measures underwent MCA across various subdomains to discern patterns among categorical interdependent (active) and independent (supplementary) variables. MCA facilitates the reduction of data dimensionality analogous to Principal Component Analysis (PCA), yet it is distinctively suited for qualitative, categorical variables. Notably, the variables “publication year”, “publication group” (datasets BTS1 to BTS3), and “total scores” were treated as supplementary variables, thus exerting no influence on the outcome of the analysis. The two principal dimensions (Dim.1, Dim.2) identified by MCA explain most of the variance (21.7% total, 13.3% and 8.4%, respectively, e.g., Fig. 6) within categorical experimental parameters among the entire dataset, with publication records similar in experimental parameter selection clustering together.

Notably, it has been described that MCA underestimates the apprehended variance, as the underlying computation introduces additional inertia ^67,68^. As a rule of thumb, the explained variance computed via MCA can be multiplied by a factor of two for comparison with familiar PCA values. Thus, the explained variance computed here is reasonable, especially for a multi-dimensional dataset. Moreover, the variance is of secondary relevance; the emerging patterns were considered to be of greater interest.

Superimposing the MCA computations with the supplementary variable “total score” revealed that publications comparable in methodological parameter selection are also similar in scored robustness (Fig. 5A). Furthermore, we observed that the line of unity roughly divides the dataset into segments of acceptable and non-acceptable methodological rigour, when the 80% threshold criterion (see section 3.2) is applied (equivalent to 12.8 or 13 absolute points, Fig. 5B). Therefore, variable patterns with centroids to the right of the line of unity, not or barely intersected by their 90% confidence interval (CI) ellipses, are considered methodologically robust, and, *vice versa*, variable patterns with identical centroid and CI ellipses characteristics to the left of the line of unity as methodologically unsound. Especially patterns clustering in the lower right quadrant can be in majority identified as methodologically robust, whereas patterns in the upper left quadrant tend to be methodologically unsound. This delineation via the line of unity was further utilised to identify experimental parameters linked to robustness patterns among the qualitative data item subdomains. Here, we focus on subdomains with distinct relational patterns, with additional data in section 3.5 of the SM. To enhance readability and graphical identification, not all CI ellipses were plotted; only ellipses of the discussed categories were plotted. For complete illustrations, please consult the SM, section 3.5, Figs. S21 to S24.

**Fig. 5:**
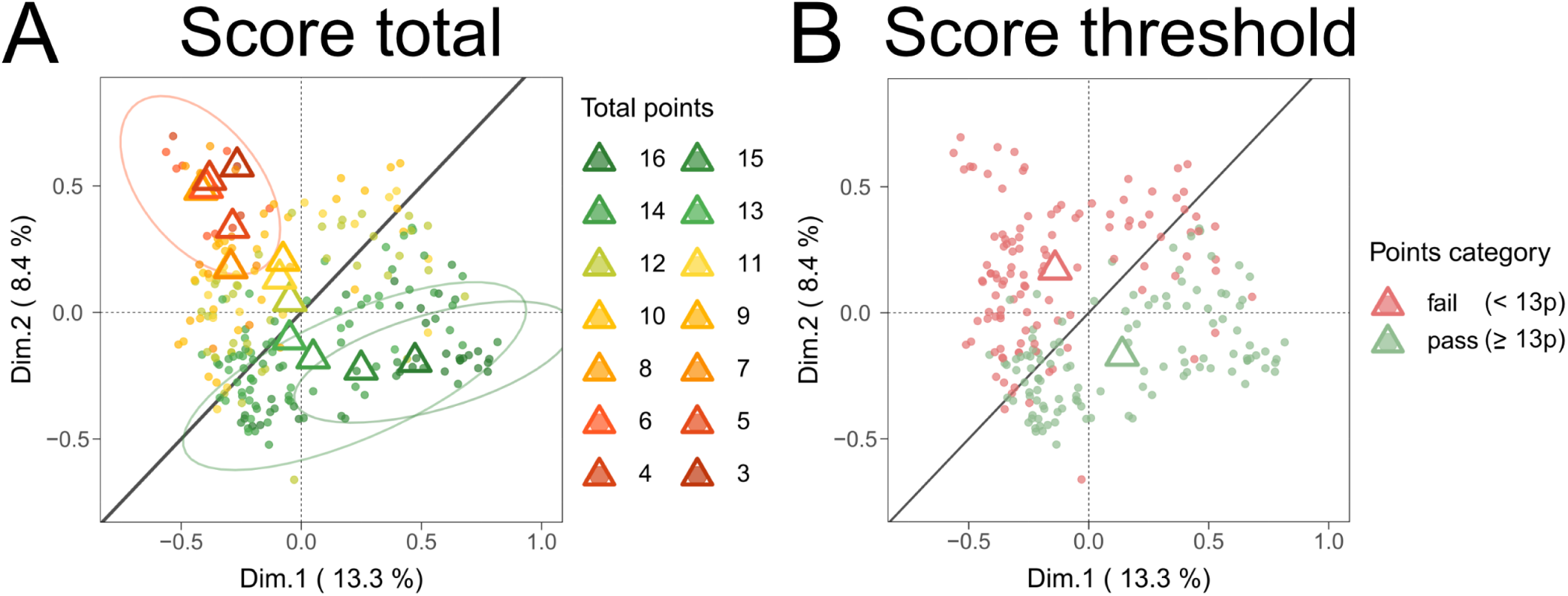
Multiple Correspondence Analysis (MCA) plots showing the clustering of n = 229 publications regarding the total attributed score (A) or divided by the 80% threshold (B). Centroids (respectively coloured triangles) mark the mean individual dimensional coordinates per variable. Ellipses (coloured) represent normal probability contours at a 0.9 confidence level. The line of unity is given in dark grey.

MCA identified patterns of methodological robustness and unsoundness within qualitative subdomain measures, hinting at the shortcomings in the methodological reporting of external BTS applications, while also indicating possible solutions.

Robustness markers were identified in the “BTS characterisation” primary domain, when examining the subdomains “BTS origin” (producer and system origin), “species”, “pooling” (sex), “husbandry”, and “BTS induction” (Fig. 6). For “BTS origin”, internally produced BTS showed more robust trends compared to externally purchased ones (Fig. 6B), and *in vitro*-derived BTS (e.g., derived from hepatocyte cultures) exhibited higher methodological soundness (Fig. 6A). In the “species” subdomain, fish and human-derived BTS systems showed greater robustness than rat-derived and undefined (“nd”) systems (Fig. 6C). Regarding “pooling”, combinations of “female and male”, or “female only” systems were more robust than “male only” or undefined (“nd”) systems (Fig. 6D).

**Fig. 6:**
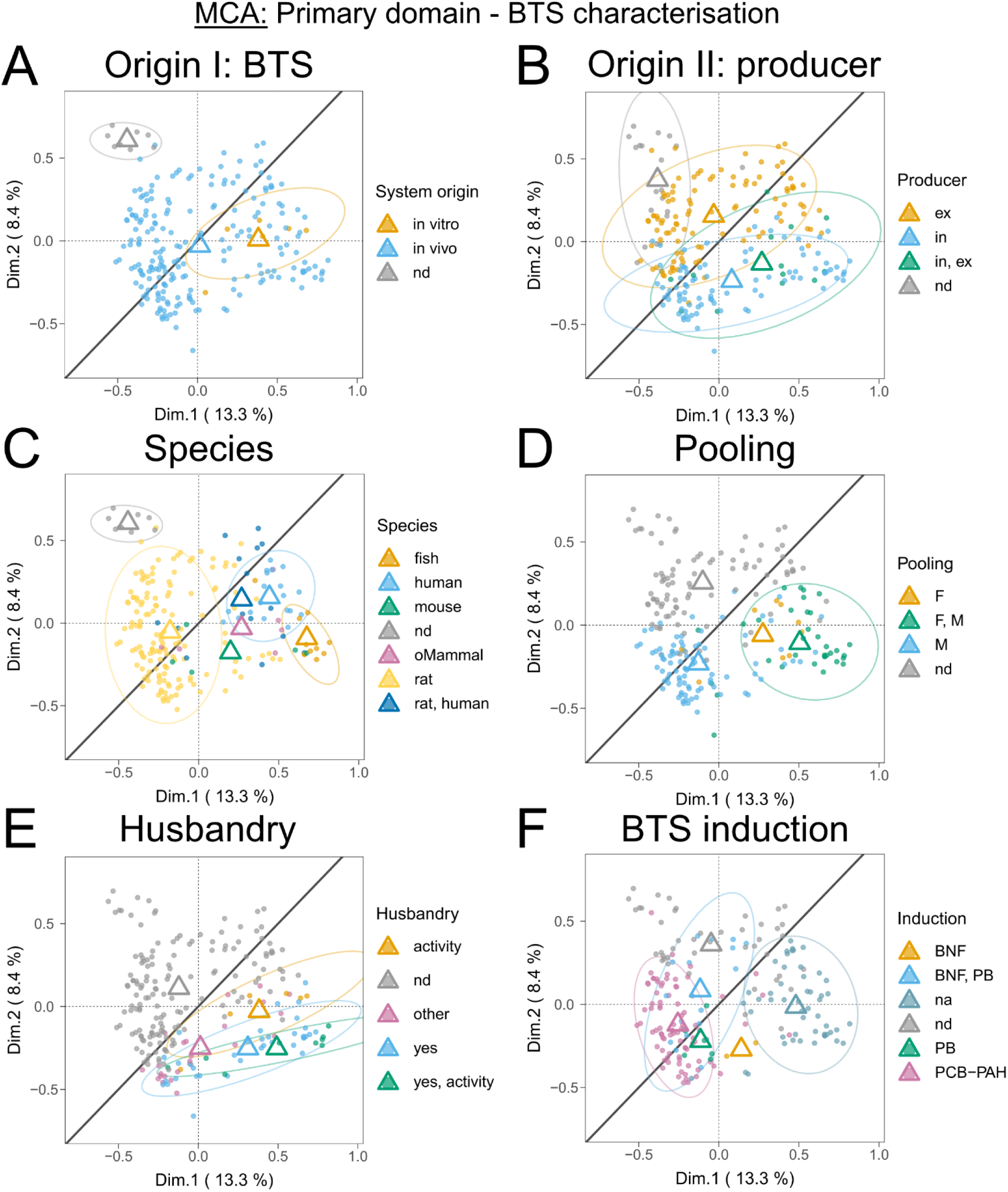
Multiple Correspondence Analysis (MCA) plots showing the clustering of n = 229 publications by the primary data item domain “BTS characterisation”. Depicted are clusters of the inquired subdomains “BTS origin” (BTS system origin, panel A; producer, panel B), “species” (C), “pooling” (D), “husbandry” (E), and “BTS induction” (F). Centroids (respectively coloured triangles) mark the mean individual dimensional coordinates per category. Ellipses (coloured) represent normal probability contours at a 0.9 confidence level. The line of unity is given in dark grey. Abbreviations: nd – not defined; na – not applicable; ex – external; in – internal: oMammal – other Mammals; F – female; M – male; BNF - beta-Naphthoflavone; PB – phenobarbital; PCB-PAH - polychlorinated biphenyls and polycyclic aromatic hydrocarbons (note that Aroclors, other PCBs, and methylcholanthrene (PAH) were summarised in this category for MCA analysis to meet computation criteria, more details are given in the SM, section 2.12, Tab. S4, and SI9).

Records reporting husbandry and enzymatic activity related to biotransformation displayed higher robustness (Fig. 6E). Finally, non-induced BTS, likely correlating with human and fish-derived systems, had higher robustness than induced systems (Fig. 6F). Especially when compared to classical BNF/PB and Aroclor-induced BTS (“PCB-PAH”; note that Aroclors, other PCBs, and methylcholanthrene (PAH) were summarised in this category for MCA analysis to meet computation criteria, more details are given in the SM, section 2.12, Tab. S4, and SI9).

We also encountered robustness markers in the primary reliability domains “BTS reaction components”, subdomains “solvent” (Fig. 7A), with studies employing organic solvents, and “BTS experimental setup”, subdomain “BTS-related control” (Fig. 7B), for records featuring the categories “w/o cofactors” and “inactivated BTS”. In general terms, for studies utilising two BTS-related controls, all centroids were located right to the line of unity, whereas only employing a single control was located to the left. The subdomain “cofactor class” depicted a robust pattern for relative observations investigating only phase 2 metabolism (Fig. S22B). However, an ellipse could not be computed due to the low number of relative observations (n = 3).

**Fig. 7:**
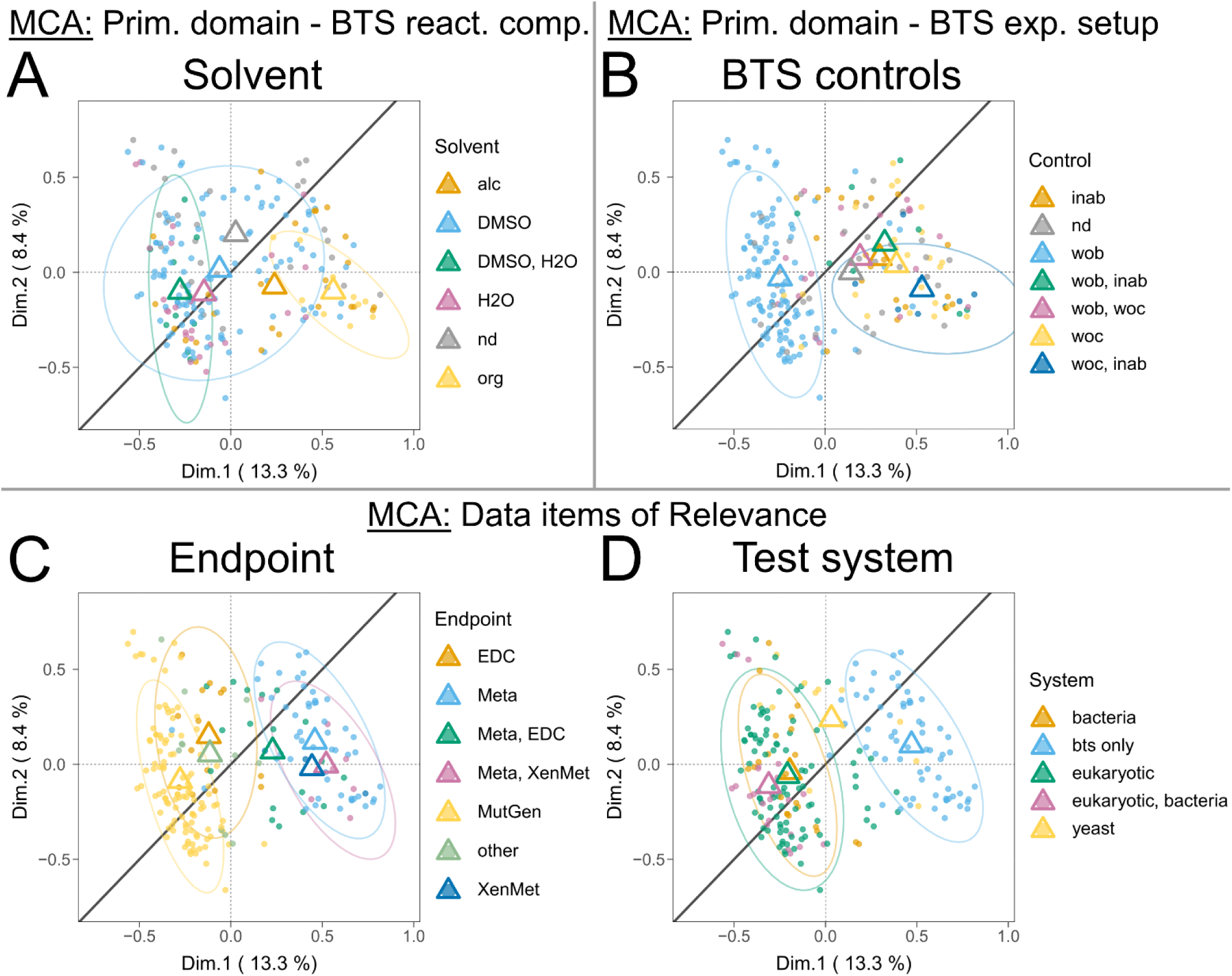
Multiple Correspondence Analysis (MCA) plots showing the clustering of n = 229 publications by the primary data item domains “BTS reaction components”, “BTS experimental setup”, and data item domains of relevance. Depicted are clusters of the inquired subdomains “solvent” (A), “BTS-related controls” (B), “endpoint” (C), and “test system” (D). Centroids (respectively coloured triangles) mark the mean individual dimensional coordinates per category. Ellipses (coloured) represent normal probability contours at a 0.9 confidence level. The line of unity is given in dark grey. Abbreviations: nd – not defined; alc – alcohol-based solvent; DMSO - dimethyl sulfoxide-based solvent; H20 – water-based solvent; org – organic solvent; inab – inactivated BTS; wob – without BTS; woc – without cofactors; EDC – endocrine disruption; Meta – metabolites; MutGen – mutagenicity and genotoxicity; XenMet – xenobiotic metabolism functionality.

Robustness markers were also observed in the “endpoint” and “test system” relevance domains (Fig. 7C and D). Studies focusing on “metabolites” (identification) and “xenobiotic metabolism” (enzymatic activity) endpoints depicted higher robustness (Fig. 7C). BTS-only test systems generally exhibited higher methodological robustness (Fig. 7D), particularly when compared to test systems that employed eukaryotes and bacteria.

However, markers of methodological unsoundness were less evident, with mainly non-defined (“nd”) data item measures per subdomain (e.g., Fig. 6A to C) clustering in the upper left quadrant. Besides that, negative trends are identified for categories with centroids left of the line of unity but crossing ellipses. Specifically, these include rat-derived BTS (Fig. 6C), BNF/PB and PCB or PAH-induced BTS (Fig. 6F), DMSO + H_2_O solvent-utilising test systems (Fig. 7A), setups using only the “w/o BTS” control (Fig. 7B), and studies focusing on mutagenicity, genotoxicity, and endocrine disruption endpoints (Fig. 7C), all depicting negative robustness tendencies.

For other qualitative data item subdomains, the overall methodological quality and reporting patterns have remained consistent across various scientific fields over time, as no or only minor discernible robustness pattern direction could be established. Other patterns also seemed less critical for our analyses. Additionally, the subdomains “strain” and “buffer system” are further discussed in the SM, section 3.5.

The MCA’s emerging patterns are instrumental in pinpointing specific areas of methodological soundness and shortcomings and provide a platform for developing procedures to address and potentially rectify these issues (as further discussed in Section 4). In summary, the subdomains “BTS origin”, “BTS induction”, “BTS controls”, “pooling”, “husbandry”, “solvent”, and “species” showed significant divergence throughout positive robustness markers (Figs. 6 and 7). Although less evident, we also identified positive robustness trajectories for the subdomains “test system”, “endpoint”, “cofactor (class)”, and “strain”. Internally produced, *in vitro*, fish, or human-derived, and non-chemically induced BTS yielded stronger robustness patterns.

#### 3.4.3 Data association rule mining and relational networks via *Apriori* algorithms

Association data rule mining, utilising *Apriori* algorithms, was implemented to discern relational networks among qualitative data item subdomains. These algorithms identify sets of qualitative data item subdomain measures that frequently coincide and establish associative connections (rules) between them. The “support” value quantifies their co-occurrence, reflecting the frequency of interactions between measures. Another key metric, “lift”, indicates the strength of item associations. Lift values above 1 signify a positive association, indicating co-occurrence beyond random expectation. Absolute scores “pass” (≥13 points) and “fail” (<13 points), as derived from the 80% relative score criterion threshold, were included in the analyses as parameters to force network anchoring and facilitate interpretation, as otherwise, the networks would be hardly interpretable to the human inspector.

Fig. 8 showcases a relational data network constrained to the 50 most prominent rules. Anchored by scoring robustness, the network bifurcates into two distinct clusters, one representing high methodological robustness (right-hand side) and the other poor quality (left-hand side), with both converging at the central association of “*in vivo*-derived BTS system origin”.

**Fig. 8:**
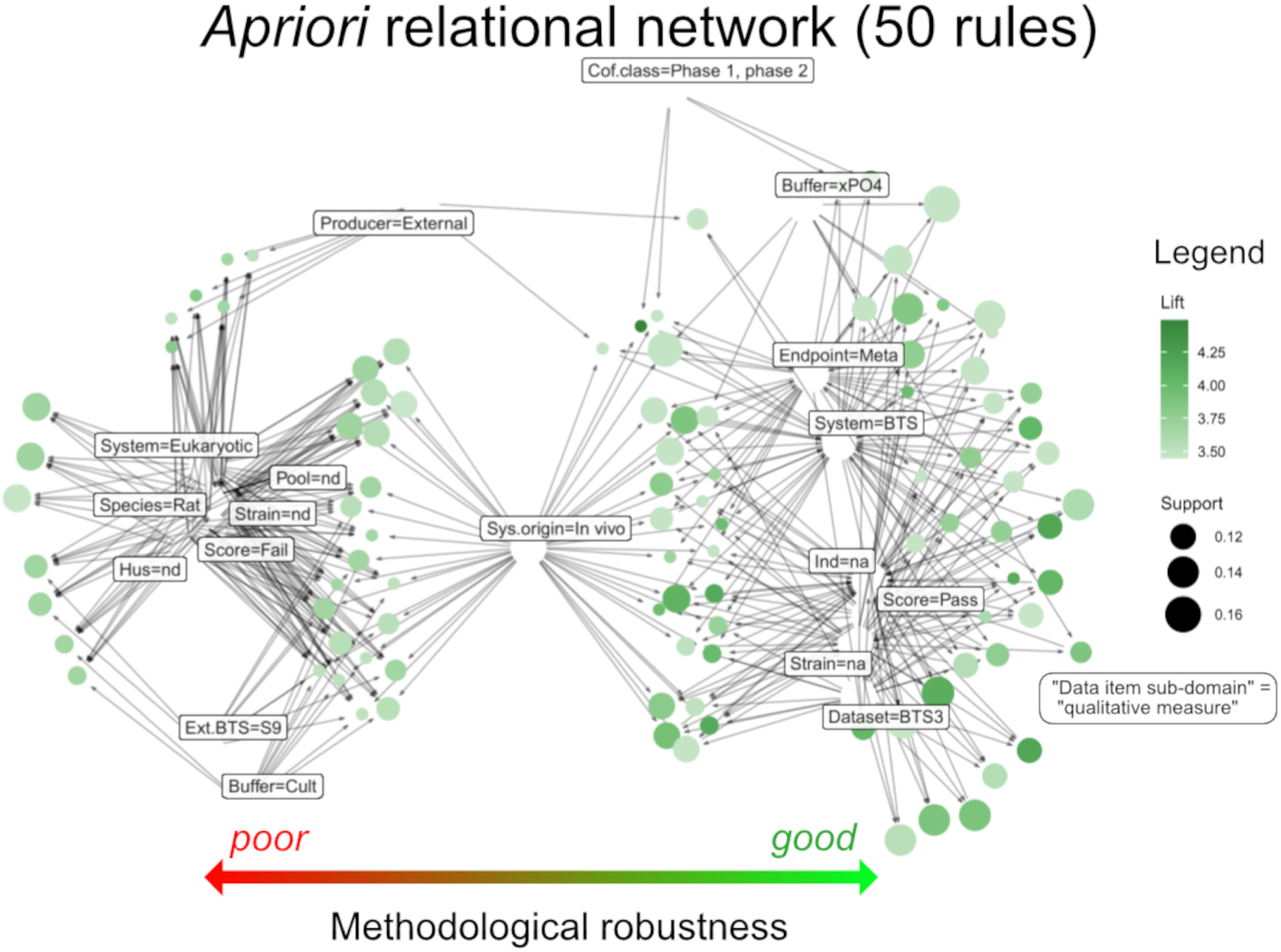
Relational data network derived from all qualitative subdomains included in the simplified, machine-readable coding book (SM, Tab. S4, and SI9). Patterns were emphasised by anchoring; relational networks to the left are related to poor methodological reliability, while clusters to the right are robust. The network comprises records from n = 229 publications. 50 rules (association connections) are depicted. Rules are shown as dots, with the size illustrating the support and the colour representing the lift of the association. Arrows indicate the direction of the rules, as associated with frequent data item measures (squares). The data item measures are located within the voids but have been moved to increase readability. Rule mining has been conducted at a support threshold of 10% and a confidence threshold of 80%. Interactive plots for 100 to 5000 rules are given in SI10. Abbreviations: nd – not defined; Cult – cultivation medium; Cof.class – cofactor class; xPO4 – generic phosphate buffer; Meta – metabolites, na – not applied; BTS3 – dataset BTS3/historical.

In the robust pattern, methodological strength correlates with the exclusive use of “BTS (-only)” systems (excluding other *in vitro* systems), assessing “metabolites” as endpoints, absence of chemical induction, and non-applicability of rodent strains – suggesting non-rodent BTS origins. This pattern is also loosely associated with using phosphate buffers and examining both phase 1 and 2 metabolism. Notably, the “BTS3/historical” database links to higher robustness, a fact that is pursued in the bias discussion (SM, section 4.1).

Conversely, the poor-quality pattern aligns with undefined measures (“nd”). Interestingly, other defined measures cluster in direct proximity, such as rat-derived BTS and test systems incorporating “eukaryotic” cells in conjunction with the BTS. Further associations include “S9 BTS”, externally sourced BTS, and systems using cellular culture medium as a buffer.

The supplementary material folder SI10 includes interactive networks based on 100 to 5000 rules. In the 100-rule network, robustness patterns mirror those in the 50-rule network (Fig. 8). The 200-rule network reveals additional negative subdomain to data item measure associations, such as “BTS-related control” to “w/o BTS” (without BTS) and “cofactor class” to “phase 1”, near lower scores. At 500 rules, new clusters emerge (“dataset” to “BTS2” (BTS2/mutagen), “endpoint” to “MutGen” (mutagenicity and genotoxicity), “solvent” to “DMSO”, and “field” to “Tox” (classic toxicology)), albeit closer to lower methodological robustness. Beyond 500 rules, score anchoring loses relevance, and the complexity renders the patterns uninterpretable by human analysis.

In summary, robustness correlated with the exclusive use of “BTS-only” test systems, assessing “metabolites” as endpoints, absence of chemical induction, and non-rodent BTS origins. Poor-quality patterns were associated with undefined measures (“nd”), the use of rat-derived BTS, studies focusing on the endpoints mutagenicity and genotoxicity, records employing culture medium as a buffer system, and studies utilising eukaryotic *in vitro* systems in combination with BTS.

#### 3.4.4 Follow-up, confirmatory analyses

With the help of the explorative mapping analyses, we discerned methodological robustness patterns and markers, which defined directions of scientific rigour and soundness (positive/negative). Table S9 in the SM, section 3.7, summarises robustness markers for identified qualitative data item subdomains and their respective measures. For subdomains that illustrated distinctive positive or negative robustness patterns in MCA and *Apriori* assessment frameworks simultaneously, we followed up with confirmatory analyses ^57^ and effect size measures of the respective seminal scoring populations to consolidate the mapping results.

In eight of nine subdomains, significant scoring population variances emerged upon examining specific measure combinations (Fig. 9). For the “BTS system” subdomain, studies employing “only BTS” in a buffer system demonstrated greater methodological robustness than those using “eukaryotic” cells in conjunction or other test systems (Fig. 9A). For studies investigating the endpoints “xenobiotic metabolism” and “metabolites”, robustness was notably higher compared to “mutagenicity and genotoxicity” and “endocrine disruption” studies (Fig. 9B). “Internally” produced BTS outperformed “externally” acquired BTS (Fig. 9C). Fish and human-derived BTS were superior in robustness to those from other organisms, particularly rats (Fig. 9D), with fish-derived BTS scoring highest in mean and median absolute scores. *In vitro*-derived BTS (e.g., from hepatocyte culture) ranked higher than classically *in vivo*-derived BTS systems (Fig. 9E). Non-induced BTS showed a generally higher absolute score than chemically induced BTS (Fig. 9F). Records utilising phosphate buffer outperformed those with culture media (Fig. 9G). Lastly, employing two or more BTS-related control types (w/o BTS, w/o cofactors, or inactivated BTS) was more advantageous than using a single BTS-related control type (Fig. 9H). No significant difference emerged in studies focusing solely on phase 2 metabolism compared to phase 1 metabolism (SM, section 3.7, Fig. S25). However, the lack of statistical significance in the latter comparison is likely due to the low power of the non-parametric test and the small sample size in one of the comparator groups (phase 2 metabolism), despite the effect measures demonstrating a clear divergence.

**Fig. 9:**
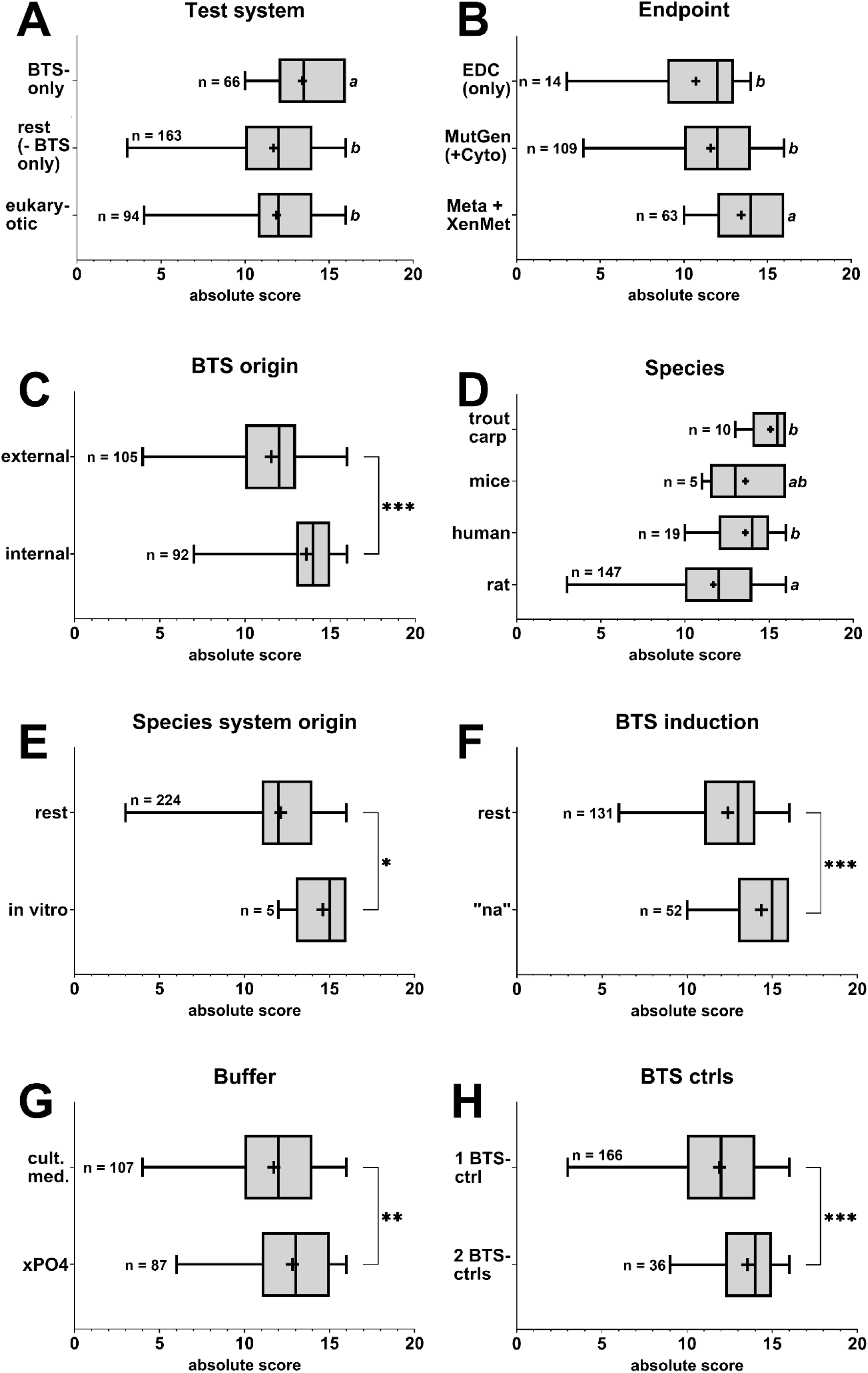
Boxplots depicting absolute scoring populations of reviewed and assessed articles for respectively curated sub-datasets per qualitative data item subdomain. Whiskers indicate the populations’ upper (max.) to lower (min.) boundaries. Boxes indicate the 75^th^ and 25^th^ percentiles, and the in-between line represents the median. Crosses represent mean population values. Statistical analyses of variance between datasets were conducted via two-sided Mann-Whitney U tests (pairwise comparison, alpha level = 0.05) or Kruskal-Wallis tests, followed by Dunn’s posthoc test (multiple comparisons, alpha level = 0.05). Asterisks indicate statistically differing significance between means of respective datasets for pairwise comparisons (*p < 0.05, **p < 0.01, ***p < 0.001). For multiple comparisons, varying letters indicate statistical significance (p < 0.05). The number of included records per seminal population analysis is given in the graphs. More details are provided in SI11. Abbreviations: EDC – endocrine disruption; MutGen – mutagenicity and genotoxicity; Cyto – cytotoxicity; Meta – metabolites; XenMet – xenobiotic metabolism functionality; “na” – not applied; xPO4 – generic phosphate buffer; cult. med. – culture medium; ctrl – control.

A previous methodological harmonisation for *in vitro* hepatic clearance studies also identified the variables “concentration”, “species”, and “culture medium” as most influential in a random forest regression analysis ^69^. Overall, the results from MCA, *Apriori*, and confirmatory analyses corroborate each other.

In conclusion, follow-up analyses on scoring population effect sizes confirmed the iterative mapping observations, showing statistically significant differences across subdomains, which reinforces the methodological robustness markers identified in the earlier analyses. Altogether, these findings are synthesised in the following sections to build a foundation for the subsequent guidance framework.

## 4. Synthesis of results and discussion

From the DEERS scoring assessment (outcome A), we deduced that the overall body of literature has a poor methodological reporting status regarding the application of BTS. Additionally, meta-regression analyses (outcome B) revealed a significant lack of correlations between key components of the BTS reaction. The iterative mapping analyses identified clear patterns of methodological robustness (outcome C) and unsoundness, which we substantiated in subsequent confirmatory analyses of effect sizes. In the following sections, we synthesise these results and propose practical ways to enhance BTS reporting. The ensuing suggestions are structured around primary data item domains, offering targeted recommendations for improving reporting practices.

### 4.1 “BTS experimental setup” is overall adequate

The primary domain, “BTS experimental setup”, was noted for its high level of methodological rigour (Figs. 2 and 3). Within this primary domain, the subdomain “BTS-related controls” emerged as a key area for potential improvement, scoring a mean relative score of 87% compared to 95% for the other two subdomains (SM, Tab. S7). Positive robustness patterns were associated with using at least two BTS-related controls (Figs. 7B and 9H). However, this did not lead to different scoring, as solely one BTS control was considered sufficient for positive evaluation and, hence, had no influence on the absolute score per record.

Within the analysed body of literature, we identified several types of BTS-related controls which are being employed to differentiate from fully active BTS reaction mixtures (BTS plus respectively used cofactors). Table S11 in the SM provides detailed descriptions of various BTS-related control setups that are employed alongside prokaryotic or eukaryotic test systems. For data parametrisation and coding procedures, we classified BTS-related controls into three categories, as follows: “w/o BTS” (per se the plain *in vitro* assay – without BTS and without cofactors), “w/o cofactors” (with BTS but lacking cofactors), and “inactivated BTS” (heat or chemically treated; cofactors are optional in relation to the test setup, see Tab. S11). Although this categorisation does not capture the full complexity of BTS-related controls, which is influenced by the parallel *in vitro* systems used, post-BTS incubation procedures, and the research questions, it was adequate for our analysis and interpretation, serving as a basis for further recommendations.

Each control type plays a crucial role in evaluating different aspects that might influence the BTS-driven biotransformation process. We address the critical aspects of BTS controls for setups that incorporate externally added BTS into pro-or eukaryotic test systems, which are also applicable to assays that use cell lysates or extracts in conjunction with the BTS. Notably, standalone BTS applications are not a primary subject of this review and may require specific control strategies, which are beyond the scope of this article. The “w/o BTS” control is essential for distinguishing between systems with and without induced biotransformation ^70–72^. As such, it is used to determine whether the substance undergoes biotransformation and whether this process alters its bioactivity (within the practicable margins of the test setup). The “w/o cofactor” control accounts for the impact of self-sustained biotransformation potential within the microsomes or S9 fractions themselves, due to the low inherent quantities of cofactors remaining as residues from homogenisation ^73^. This might be specifically important for studies utilising short BTS incubation times (e.g., 10 min) or focusing on specific phase 1 or 2 enzymatic activities. Especially for S9 fractions, other biotransformation reactions may occur simultaneously, in addition to those specifically investigated by adding respective phase 1 or 2 cofactors. Lastly, the “inactivated BTS” control is crucial for assessing the impact of BTS addition on the bioavailability of chemicals within the test system. BTS addition can significantly alter the protein content and, consequently, the distribution of hydrophobic/lipophilic compounds ^74,75^. The additional protein may act as a sink compartment for hydrophobic/lipophilic compounds and impact the actual free concentration of the chemical under study ^76^. This is especially crucial in studies using complex culture media as buffer systems, such as those employing BTS alongside eukaryotic *in vitro* systems, for example, reporter gene assays. Additional BTS control considerations are especially required when developing and troubleshooting novel methods, which are further addressed in the SM, Tab. S11. Overall, it ultimately rests in the experimenter’s judgement to carefully select the optimal setup of necessary controls, which needs to be clearly described in the methodology.

Given the results of our exploratory analyses and the preceding discussion, we advocate for the inclusion of at least two BTS-related controls. Experimenters should consistently include both “w/o BTS” (per se, the plain *in vitro* assay) and “inactivated BTS” (without cofactors) controls, especially in studies employing BTS alongside pro-or eukaryotic test systems, to account for total biotransformation impact and bioavailability issues.

### 4.2 Minor improvements can have a significant impact, “BTS reaction”

In our analyses of the “BTS reaction” primary domain, it was observed that all subdomains, except for “BTS protein concentration”, were somewhat satisfactorily described, with relative scores ranging from 81 to 97% (SM, Tab. S6). However, the “BTS protein concentration” subdomain significantly lagged, achieving only a 57% relative score. Accurately specifying BTS protein concentration in mg/mL is vital for several reasons. Firstly, it enables the normalisation of results across various study endpoints, enhancing the comparability of different studies ^16,77,78^. Secondly, elevated concentrations of BTS can induce cytotoxic effects, which are particularly problematic in studies involving prokaryotic or eukaryotic test systems ^79,80^. As well, an improved definition of cofactor-related measures is needed overall, for which detailed insights have already been provided above in section 3.4.1.

A common shortfall in many studies is reporting BTS concentration as a percentage of the final reaction mixture (% v/v) without specifying the initial concentration. This approach is likely rooted in historical protocols by Ames *et al.* ^26–28^, which introduced the Ames mutagenicity test, a significant catalyst of BTS utilisation within toxicology. Ames *et al.*’s initial protocols mentioned BTS protein concentrations of “approximately” 40 mg/mL. However, they also acknowledged batch-to-batch variations and differences arising from using various species or rat strains. Over time, not only did the % v/v concentrations in the Ames protocols vary (ranging from 4 to 30%, see SM, Tab. S10), but discrepancies also appeared between these protocols and the foundation publications they were based on ^81,82^. This inconsistency underlines the methodological variations within this body of literature. Lastly, even the approximated 40 mg/mL BTS concentration cannot be used as a rough standard since we have encountered literature that employs the Ames protocols for BTS production but states divergent BTS protein concentrations (e.g., ^83–85^). While the historical Ames protocols focused on activity normalisation using the number of revertant colonies in response to a reference control, such as aflatoxin B1, this method is not universally applicable to other BTS-relevant endpoints. Therefore, many studies referencing Ames *et al.* for BTS-related methodologies fail to specify critical parameters, rendering this approach inadequate for precise BTS characterisation due to the inherent parameter variability and the evolution of these protocols over time.

Expert panel, best practice reports on the methodological implementation of the *in vitro* Comet Assay ^71^ and the *in vitro* micronucleus assay ^72^ maintain the use of %v/v metrics, while the guidance for the chromosomal aberration assay ^70^ recommends presenting BTS protein concentrations in mg/mL. Similarly, respective and more recent *in vitro* mutagenicity and genotoxicity-related OECD test guidelines (TGs no. 471, 473, 476, and 487; ^11,86–88^) also do not specify reporting BTS concentrations in mg/mL. While all TGs demand defining S9 BTS type and composition within the TG test report, including the final S9 concentration in culture medium, they abstain from defining concentration metrics. Contextually, all of the named TGs address S9 concentrations in % v/v.

Regulatory authority guidance relating to standardised *in vitro* genotoxicity tests (TGs), such as by the UK government C*ommittee on the mutagenicity of chemicals in food, consumer products and the environment* (COM), mentions the utilisation of S9 BTS but does not give details on practical utilisation or reporting standards ^89,90^. Finally, a recent volume of *Methods in Molecular Biology* published a series of protocols related to *in vitro* genotoxicity and mutagenicity ^91^. While many protocols define BTS mixtures in detail regarding employed cofactors, regeneration systems, and buffers, all rely on the % v/v for S9 BTS. Adherence to Ames’ historical protocols, and thereby the % v/v metric, may influence the methodological robustness in studies of mutagenicity and genotoxicity.

Another area for improvement is the “BTS dilution” subdomain. Although most records scored well (89% relative score, see SM, Tab. S6), some publications either did not define (not defined – “nd”) or were unclear (not clear - “nc”) in describing the dilution steps from BTS and cofactor stocks to final reaction concentrations. This lack of clarity means that the experimental setup is not reproducible; however, this issue is easily rectified.

To conclude, it is essential to always define BTS protein concentrations in mg/mL and explicitly state the concentrations of BTS and cofactors at the stock, mix, and final reaction stages to ensure methodological robustness and reproducibility.

### 4.3 “BTS characterisation” - the primary culprit

In section 3.2, we highlighted the primary data item domain “BTS characterisation” as exhibiting the lowest methodological soundness in our study, particularly in the subdomains “strain”, “pooling” (sex and number of individuals), “husbandry”, and “BTS induction” (see SM, Tab. S5). The iterative mapping analyses revealed a troubling trend: Negative robustness patterns associated with externally purchased BTS, especially those derived from rats and used in conjunction with eukaryotic test systems and cell culture mediums (Fig. 8). A notable increase in the use of externally sourced BTS over the years has been accompanied by a decline in the quality of BTS characterisation reporting (see SM, Figs. S8C, S11C, S12C, and S14C). In contrast, BTS produced in-house consistently showed higher standards of reporting (Fig. 9C). Per se, this is self-evident, given that experimenters who hand-craft the BTS are likelier to report on the specific parameter of BTS characterisation. On the other hand, purchasing BTS from an external supplier was associated with non-compliance with reporting standards and a lack of certification/documentation from the suppliers. Dependence upon the mention of a BTS supplier or producer is insufficient, as it withholds specific details necessary for interpretation.

The rising trend in sourcing BTS externally is concerning, considering the critical role of BTS in high-throughput applications, such as eukaryotic reporter systems. These applications often involve studying various toxicity pathways’ modes of action, where robustness in BTS characterisation is crucial. To address this, we compiled a non-exhaustive list of BTS characterisation items from different (anonymised) producers encountered within the investigated literature (see SM, Tab. S12). Our findings indicate inconsistent BTS characterisation among producers, particularly with respect to the assessment of phase 1 and 2 capacities. Producers offer a variety of BTS, differing in type (S9 or microsomes), species of origin, pooling, and chemical induction. The lack of standardisation in BTS production protocols further complicates this issue, particularly where production has been discontinued, making it impossible to request or obtain any documentation. A standardised BTS production protocol is needed to guide production and ensure consistent reporting. Producers are encouraged to consider offering well-characterised BTS to meet these quality requirements, and the end-user can promote this process by requesting such information from their supplier.

In practical applications, mainly when BTS is used in conjunction with pro- and eukaryotic test systems, the BTS protein concentrations typically range between 0.01 – 0.1 mg/mL in the final reaction mix (i.e., in the well of a microtiter plate). Various studies have identified concentrations within this range as optimal ^77,92–96^. Higher concentrations have been linked to cytotoxic effects or the introduction of BTS-derived cellular debris that could interfere with later-stage assay readouts, such as luminescence, fluorescence, or absorbance readings ^97^ (Lungu-Mitea *et al.*, in preparation). Thus, a small initial aliquot of well-characterised batches of BTS (5 - 10 mL) could suffice for an entire project. Ideally, producers could conduct characterisation in a high-throughput manner during production, ensuring economic efficiency. This approach could align with the research and regulatory guidance framework discussed in section 5.

### 4.4 Possible solutions and a way forward

In contrast to the challenges highlighted in the previous sections, our analyses identified good robustness patterns where the literature studies focused on metabolites and xenobiotic metabolism functionality as endpoints (Figs. 7C, 8, 9B). These studies predominantly utilised BTS in generic phosphate buffers (Figs. 8, 9G) and employed non-induced BTS (Figs. 6F, 8, 9F), usually of non-rodent origin, such as human or fish. This trend suggests that studies specifically targeting xenobiotic metabolism functionality are more likely to accurately address all BTS components, owing to the precise focus of their objectives. Similarly, studies involving metabolite characterisation post-BTS incubation, e.g., via mass spectrometry, often employing simpler BTS systems, also tend to report all relevant BTS constituents more consistently.

An intriguing robustness pattern emerged with the use of *in vitro*-derived BTS (Figs. 6A, 9E), where BTS is extracted from immortalised hepatocyte cell cultures rather than animal-derived liver homogenates (e.g., ^98–100^). This approach is particularly noteworthy as it aligns with the principles of reducing, refining, and replacing animal experiments. However, only a limited number of publications (n = 5) have adopted this method, suggesting that it is an emerging aspect but also highlighting the potential for significant growth and improvement in this area.

Moreover, the source species of BTS substantially influenced the quality of reporting (Figs. 6C, 8, 9D). Human, mouse, and fish-derived BTS generally received higher scores than rat-derived BTS. Human- and fish-derived BTS outperformed rat-derived BTS in overall scoring. Also, the comparison of fish-derived BTS to the rest of the dataset indicated greater data quality (see SM, Fig. S26).

Human-derived BTS is often more thoroughly reported, likely due to its status as a valuable pharmaceutical resource obtained from donor organs, coupled with the inherent variability in human phase 1 and 2 enzyme activity ^14,101^. However, the ethics in using human tissue for commercially related activities and the high variability reduce acceptability when used for chemical regulatory purposes at the global level. Whilst in pharma, it is useful to have this variability, for chemical hazard assessment test methods, reproducibility is essential for regulatory acceptance. Indeed, the issues discussed around human serum are equally applicable to human BTS ^102^.

As mentioned earlier, the contemporary ICH M12 Guideline recommends using hepatic *in vitro* systems and PBPK *in silico* modelling for drug interaction studies ^14^. It specifies using human liver tissue fractions, such as microsomal systems from at least ten donors. Microsomal protein concentrations should be minimised, and standardised assay conditions, including buffer strength and pH, must be applied. BTS should be characterised with selective *in vitro* probe substrates for phenotyping experiments to confirm enzyme activity. It also lists probe substrates with marker reactions and advises including inhibitors or inducers as positive controls. Our analysis aligns closely with and complements the ICH M12 guidance, offering potential synergies for the scientific and regulatory communities.

Fish-derived BTS, typically from species like trout and carp, are rarely sourced externally and are usually prepared following established protocols ^103^, which have been adopted into OECD TG 319B^104^. This guideline details essential steps and reporting recommendations, thereby inherently promoting methodological rigour in the research that applies them.

In contrast, “classical” BTS (male rat, Sprague-Dawley strain-derived, Aroclor or BNF/PB-induced, co-utilised with bacterial or eukaryotic test systems, buffered in culture medium, with exposure chemicals solved in DMSO or aqueous solutions, employing solely “w/o BTS” as a control, and investigating the endpoints mutagenicity, genotoxicity, and endocrine disruption) were associated with methodological limitations in at least one analysis conducted in outcome C. Given these findings, we propose the development of a research and regulatory guidance framework applicable to all BTS types. Such a framework could enhance the standardisation of BTS characterisation and reporting, potentially integrating it into broader *in vitro* toxicology guidance, such as the GIVIMP ^105^ and the Guidance Document for Describing Non-Guideline In Vitro Test Methods ^106^.

## 5. Guidance framework to support regulatory-relevant and research applications (outcome D)

Building upon the insights developed from the critical analyses of our results, synthesis, and discussion, we propose an evidence-based reporting guidance framework for external biotransformation systems (“BTS”, S9 or microsomal liver fractions), which is particularly relevant for experimenters employing BTS in conjunction with pro- or eukaryotic test systems, such that their work can achieve greater utility. This framework aims to enhance methodological rigour and augment the value of datasets for subsequent analyses. The framework centres around key recording categories or data item (sub-)domains that scientific publications and reports should routinely include. By following these recommendations, researchers can contribute to a more robust and better-utilised body of scientific literature. The guidance is intended to be self-reliant, allowing the reader to use it without familiarising themselves with the preceding article. However, in some instances, referencing other literature or sections of the main manuscript is necessary to link to additional details, whilst ensuring coherence and conciseness.

Our guidance takes inspiration from existing protocols for fish-derived BTS, which our study identified as one of the most methodologically sound systems ^103,104^, combined with additional criteria derived from our analyses, together with the M12 guideline from the International Council for Harmonisation of Technical Requirements for Pharmaceuticals for Human Use (ICH) ^14^. This approach also considers the challenges and issues previously discussed in the literature regarding the use of external BTS in toxicological *in vitro* systems ^6,7,9,16^. We provide a roadmap to help researchers navigate these challenges more effectively.

The proposed reporting categories are supplemented with brief discussions elucidating the scientific rationale behind their necessity. A separate reporting checklist is available as a specific, single supplement provided with the main article, which can be used to develop suitable study designs and facilitate report preparation for publication. The reporting categories essentially mirror the data item domains extracted in our study, focusing on the reliability aspects. If these data requirements are routinely included in scientific publications and regulatory study reports using BTS, a concomitant improvement in the quality of BTS-related research is expected.

## I. BTS characterisation

### I. Determine BTS origin (internally produced or externally purchased)

If produced in-house, define the production protocol in detail, including husbandry, organism source, organism supplier, housing conditions, organism weight, the weight of extracted liver tissue, gonadosomatic index, etc. Detailed BTS protocolling and reporting instructions for in-house production can be found in TG 319B and associated publications pertaining to trout S9 ^103,104^. All the following points (I.A to I.E) should be addressed when reporting on in-house produced and applied BTS. If externally purchased, name the producer and give as many details as possible (aligned with points I.A to I.E). Ideally, request this information from the supplier prior to purchase. Until a common standardisation of BTS production has been agreed upon and also implemented for BTS suppliers, we strongly recommend that investigators document all details of BTS characterisation provided by the producer/supplier.

#### I.A Define the type of BTS, e.g., microsomal or S9

Defining the type and source of BTS is crucial, as microsomes and S9 fractions have different biotransformation capacities. While S9 BTS incorporate both mitochondrial and cytosolic fractions of the hepatic homogenate and are capable of both phase 1 and phase 2 biotransformation processes if supplied with the specific cofactors (see more details in points II.D and II.E), microsomes contain only mitochondrial fractions. Thus, they are limited to phase 1 metabolism and UDP-glucuronosyltransferase (UGT, phase 2) activities. Notably, even when investigating identical phase 1 processes, differences have been encountered between microsomal and S9 BTS ^107^.

Given the scenario in which BTS are *in vitro*-derived (e.g., ^99,100^), it is essential to specify the cell culture origin and its biotransformation capacities, as some metabolism pathways may be limited in permanent or recombinant cell lines or by specific induction patterns sought and applied ^98,108^.

#### I.B Define the species of BTS origin

Species of origin plays a pivotal role in the functionality of BTS due to varying biotransformation capacities and pathways, as extensively reported in the literature ^109–112^. Thus, specifying the species from which the BTS is derived is essential.

#### I.C Define the strain of the above species, if applicable

Strain-specific differences in rodent biotransformation capacities have been well-documented ^113–117^.

#### I.D Provide details on BTS pooling procedures, especially the number of individuals and sex

In humans, high variability within CYP isoforms has led to pooling across several individuals to reduce variability ^101,118^. Sex differences in rodent BTS production have been reported to impact biotransformation capacities on several occasions ^119–121^. Moreover, the biopsy location within the liver of the same individual used for BTS production can lead to variations in biotransformation capacity, highlighting the need for detailed specification ^114^ and, eventually, pooling. The ICH M12 guideline ^14^ specifies that a pool of at least 10 donors is preferred for human liver microsomes.

#### I.E Define the biotransformation-inducing chemical agents, if any, that were used. Explicitly state if no chemical induction was performed to avoid confusion

Studies have shown that different chemical inducers in source animals can lead to varying BTS capacities ^122–124^. With the Stockholm Convention ^25^, for OECD TGs, PCB-induced rat livers (e.g., via Aroclor) have been phased out in favour of phenobarbital plus beta-naphthoflavone mixtures (PB/BNF). PB/BNF induction has been described as qualitatively similar to Aroclor but with quantitative differences ^110,125,126^. Furthermore, “Aroclor” does not refer to a uniform mixture. Instead, it can encompass diverse profiles of PCB, PCDD, and PCDF congeners ^127^.

## II. BTS reaction components

### II. Clearly define all reaction components and their respective concentrations within the final BTS reaction mixture

The final BTS reaction mixture constitutes S9 or microsomal fractions, cofactors, buffers, solvents, and exposure chemicals (see points II.A to II.D below) after their addition to the employed *in vitro* assay and/or chemical exposure within the reaction vessel (microtiter well, cuvette, etc.) at the time point when all reaction components are simultaneously present, and the biotransformation reaction starts (often also referred to as “incubation”, see also point III.A). Also, defining components’ concentrations within stocks and master mixes (dependent on the utilised test setup) is advised. For recommended concentration units, refer to points II.C and II.D. Finally, clearly marking the dilution/titration steps between stocks, master mixes, and the final BTS reaction mixture facilitates the reproducibility and clarity of the experimental protocol and is therefore recommended.

#### II.A Define the BTS buffer system in which the final BTS reaction takes place

The buffer system in which the final BTS reaction occurs is critical, especially for cell culture medium-buffered systems. Different components in media, such as amino acids, vitamins, and cofactors, as well as pH, can impact BTS activity ^128,129^.

#### II.B Specify the utilised solvent(s) for chemical exposure, especially the solvent concentrations within the final BTS reaction mixture

Several publications indicated that various organic solvents could impact phase 1 and phase 2 BTS enzymatic activity if utilised within the same reaction mixture ^65,78,130^.

#### II.C Provide the S9 or microsomal fraction protein concentration in mg/mL, not percentages

Most importantly, provide the S9 or microsomal fraction protein concentration in the final BTS reaction mixture. S9 or microsomal fraction protein concentration in mg/mL can be utilised to normalise the results of downstream readouts related to BTS biotransformation processes. Reporting BTS protein concentrations in mg/mL has been independently suggested by different authors (e.g., ^16,77,78^). Further, defining the BTS protein concentration can give the reader additional insight into the potential cytotoxic effects of BTS ^79,80^.

#### II.D Name the utilised cofactors appropriately and define their molar concentrations at least within the final BTS reaction mixture

Familiarise yourself with the cofactors needed for your research question. If a NADPH regeneration system is utilised, define all components fully. Optimal use of cofactor combinations, regarding BTS utilisation in a specific experimental setup, and NADPH regeneration systems are discussed in previous studies (e.g., ^62,63,65,66^). Characterising the utilised regeneration systems is vital, as different systems can lead to varying results ^63,131^. It is also noted that cofactors and regeneration systems can introduce cytotoxicity themselves in specific scenarios ^132^.

#### II.E Define, report, or measure BTS enzymatic activity

If BTS were purchased externally, request this information from your supplier. If necessary or for in-house produced BTS, conduct the measurement yourself (e.g., CYP1A, CYP3A, CYP2B6, CYP2C9, GST). Lack of standardisation in assessing BTS-induced enzymatic biotransformation activity necessitates obtaining the producer’s activity information as part of the Certificate of Analysis or determining it independently if not forthcoming from the supplier. For now, when purchasing external BTS, we recommend indicating all information given by the producers and requesting missing key information. Suggestions on enzymatic activity endpoints and protocols can be found in the literature ^14,78,133,134^. For studies investigating toxicological mechanisms of action, e.g., via eukaryotic test systems complemented with BTS, assessing all phase 1 and phase 2 reactions for which specific cofactors have been added is recommended. A standard approach would assess CYPs, GST, UGT, and SULT activity, e.g. via HPLC MS-MS ^12,135^ or metabolic turnover of fluorometric substrates ^136^. Notably, a live-cell, fluorometric, enzymatic activity assessment battery protocol for these endpoints is underway (Lungu-Mitea et al., in preparation).

## III. BTS experimental setup

### III.A Define BTS incubation time and temperature

Incubation time must be considered in relation to the employed BTS protein and cofactor concentrations so that biotransformation reactions can realistically proceed ^16^. Studies employing prolonged incubation must assess potential cytotoxicity impacts ^137^. If non-mammalian BTS are applied, the incubation temperature should be adjusted accordingly ^107^. Optimally, experimenters are advised to also report on the circumstances of incubation, including the use of shaking devices and the type of reaction vessel (glass, plastic, etc.).

### III.B Define BTS-related controls

Carefully select a set of BTS-related controls according to the study design and objectives; their proper implementation and documentation are necessary for any BTS applications. Appropriate BTS controls are vital for interpreting the impact of the chemical biotransformation on its bioactivity, distinguishing between systems with induced biotransformation ^70–72^, assessing self-sustained biotransformation potential ^73^, and evaluating the impact of BTS on the bioavailability of chemicals ^74,75^. For studies that employ the BTS alongside pro- or eukaryotic test systems, we recommend using at least the “w/o BTS” (without BTS and without cofactors, per se the plain *in vitro* assay) and heat or chemically “inactivated BTS” (excluding cofactors) controls to evaluate biotransformation impact and address bioavailability-related issues, respectively. More detailed reasoning on the necessity of these BTS-related controls is given in section 4.1 of the main article and Table S11 in the SM. Additional BTS control scenarios might be necessary or informative when developing, characterising, and troubleshooting novel methods, as well as for specific BTS applications, which is also addressed in Table S11.

## Supporting Information

This article is accompanied by a Supplementary Manuscript (SM), eleven Supplementary Information files (SI1–SI11), and a BTS reporting checklist. The SM and reporting checklist are provided with the article, while the SI files are accessible via FigShare (https://doi.org/10.6084/m9.figshare.28504826.v1).

A summary of the supplementary materials is as follows:

- SM (PDF): detailed conducted methodologies, additional results and discussion, references of all bibliographic records, R-code, DEERS data extraction form, and an abbreviation list.
- BTS reporting checklist (PDF): a concise, user-friendly checklist for BTS methodological reporting.
- SI1 (XLSX): complete dataset for endocrine disruption endpoint searches (“BTS1” dataset).
- SI2 (XLSX): complete dataset for mutagenicity and genotoxicity endpoint searches (“BTS2” dataset).
- SI3 (XLSX): dataset of endocrine disruption endpoint screening selections.
- SI4 (XLSX): dataset of mutagenicity and genotoxicity endpoint screening selections.
- SI5 (XLSX): bibliographic records previously identified as relevant by reviewers and associated cross-references (“BTS3” dataset), including their respective screening selections.
- SI6 (XLSX): comprehensive dataset of all extracted quantitative and qualitative data item measures across (sub)domains, forming the foundation for all subsequent meta-analyses.
- SI7 (XLSX): DEERS assessment-derived total scores.
- SI8 (XLSX): raw and meta-level data from meta-regression analyses.
- SI9 (XLSX): coding simplifications of data item measures (from SI6) for machine readability, including a binary information matrix for Multiple Correspondence Analysis.
- SI10 (folder): additional association networks (100 to 5000 rules) generated using Apriori algorithms.
- SI11 (XLSX): raw and meta-level data from follow-up, confirmatory analyses.

## Data availability statement

All data generated or analysed throughout this study are included in this article (and its supplementary files; as mentioned above, the SM, SI1-11, and the reporting checklist). The SM provides in-depth documentation of the utilised systematic tools, illustrates additional data, and discusses the conceptual framework of the review. Additional supplementary information comprising raw data, metadata, and analyses have been uploaded to Figshare and are available under the following link: (https://doi.org/10.6084/m9.figshare.28504826.v1).

## Supporting information

Supplementary manuscript

BTS reporting checklist

## Acknowledgements

The authors are grateful to Dr. Klára Komprdová, Pharmacokinetic Modelling, MUNI-RECETOX, Czech Republic, for statistical consultation. Further, we thank Mgr. Libuše Janská, Department of Oncology-Pathology, Karolinska Institute, Sweden, for reviewing and proofreading the manuscript.

## Author contributions: CRediT

Conceptualisation: SLM, MJ, AS, and KH; Data curation: SLM and MSA; Formal analysis: SLM and MSA; Funding acquisition and resources: KH, SLM, AS, and HH; Investigation: SLM and MSA; Methodology: SLM, FPV, and IR; Project administration: SLM and KH; Writing, original draft: SLM and MJ; Writing, review and editing: all authors; Supervision: KH, MJ, and HH; Guarantors: SLM and KH

## References

(1) WHO, 2007. IPCS mode of action framework, Chemical Safety and Health Unit. WHO Press. URL: https://www.who.int/publications/i/item/9789241563499

(2) NRC. Toxicity Testing in the 21st Century; National Academies Press: Washington, D.C., 2007. 10.17226/11970.

(3) Collins, F. S.; Gray, G. M.; Bucher, J. R. Transforming Environmental Health Protection. Science (1979) 2008, 319 (5865), 906–907. 10.1126/science.1154619.

(4) Tice, R. R.; Austin, C. P.; Kavlock, R. J.; Bucher, J. R. Improving the Human Hazard Characterization of Chemicals: A Tox21 Update. Environ Health Perspect 2013, 121 (7), 756–765. 10.1289/ehp.1205784.

(5) European Commission. COMMUNICATION FROM THE COMMISSION on the European Citizens’ Initiative (ECI) ‘Save Cruelty-Free Cosmetics – Commit to a Europe without Animal Testing’; 2023; Vol. Official J. https://eur-lex.europa.eu/legal-content/EN/TXT/?uri=OJ:JOC_2023_290_R_0001.

(6) Coecke, S.; Ahr, H.; Blaauboer, B. J.; Bremer, S.; Casati, S.; Castell, J.; Combes, R.; Corvi, R.; Crespi, C. L.; Cunningham, M. L.; Elaut, G.; Eletti, B.; Freidig, A.; Gennari, A.; Ghersi-Egea, J.-F.; Guillouzo, A.; Hartung, T.; Hoet, P.; Ingelman-Sundberg, M.; Munn, S.; Janssens, W.; Ladstetter, B.; Leahy, D.; Long, A.; Meneguz, A.; Monshouwer, M.; Morath, S.; Nagelkerke, F.; Pelkonen, O.; Ponti, J.; Prieto, P.; Richert, L.; Sabbioni, E.; Schaack, B.; Steiling, W.; Testai, E.; Vericat, J.-A.; Worth, A. Metabolism: A Bottleneck in In Vitro Toxicological Test Development. Alternatives to Laboratory Animals 2006, 34 (1), 49–84. 10.1177/026119290603400113.

(7) Jacobs, M.; Janssens, W.; Bernauer, U.; Brandon, E.; Coecke, S.; Combes, R.; Edwards, P.; Freidig, A.; Freyberger, A.; Kolanczyk, R.; Mc Ardle, C.; Mekenyan, O.; Schmieder, P.; Schrader, T.; Takeyoshi, M.; Burg, B. The Use of Metabolising Systems for In Vitro Testing of Endocrine Disruptors. Curr Drug Metab 2008, 9 (8), 796–826. 10.2174/138920008786049294.

(8) OECD. Detailed Review Paper on the State of the Science on Novel In Vitro and In Vivo Screening and Testing Methods and Endpoints for Evaluating Endocrine Disruptors; OECD Series on Testing and Assessment; OECD, 2008. 10.1787/9789264221352-en.

(9) Jacobs, M. In Vitro Metabolism and Bioavailability Tests for Endocrine Active Substances: What Is Needed next for Regulatory Purposes? ALTEX 2013, 30 (3), 331–351. 10.14573/altex.2013.3.331.

(10) OECD. Test No. 487: In Vitro Mammalian Cell Micronucleus Test; OECD Publishing, 2014. 10.1787/9789264224438-en.

(11) OECD. Test No. 471: Bacterial Reverse Mutation Test; OECD Guidelines for the Testing of Chemicals, Section 4; OECD, 2020. 10.1787/9789264071247-en.

(12) Bernasconi, C.; Pelkonen, O.; Andersson, T. B.; Strickland, J.; Wilk-Zasadna, I.; Asturiol, D.; Cole, T.; Liska, R.; Worth, A.; Müller-Vieira, U.; Richert, L.; Chesne, C.; Coecke, S. Validation of in Vitro Methods for Human Cytochrome P450 Enzyme Induction: Outcome of a Multi-Laboratory Study. Toxicology in Vitro 2019, 60 (March), 212–228. 10.1016/j.tiv.2019.05.019.

(13) Jacobs, M. N.; Kubickova, B.; Boshoff, E. Candidate Proficiency Test Chemicals to Address Industrial Chemical Applicability Domains for in Vitro Human Cytochrome P450 Enzyme Induction. Frontiers in Toxicology 2022, 4 (June), 1–25. 10.3389/ftox.2022.880818.

(14) ICH M12. ICH HARMONISED GUIDELINE - DRUG INTERACTION STUDIES M12; 2024. URL: https://database.ich.org/sites/default/files/ICH_M12_Step4_Guideline_2024_0521_0.pdf.

(15) Funk, C.; Roth, A. Current Limitations and Future Opportunities for Prediction of DILI from in Vitro. Arch Toxicol 2017, 91 (1), 131–142. 10.1007/s00204-016-1874-9.

(16) Gouliarmou, V.; Lostia, A. M.; Coecke, S.; Bernasconi, C.; Bessems, J.; Dorne, J. Lou; Ferguson, S.; Testai, E.; Remy, U. G.; Brian Houston, J.; Monshouwer, M.; Nong, A.; Pelkonen, O.; Morath, S.; Wetmore, B. A.; Worth, A.; Zanelli, U.; Zorzoli, M. C.; Whelan, M. Establishing a Systematic Framework to Characterise in Vitro Methods for Human Hepatic Metabolic Clearance. Toxicology in Vitro 2018, 53 (July), 233–244. 10.1016/j.tiv.2018.08.004.

(17) Qu, W.; Crizer, D. M.; DeVito, M. J.; Waidyanatha, S.; Xia, M.; Houck, K.; Ferguson, S. S. Exploration of Xenobiotic Metabolism within Cell Lines Used for Tox21 Chemical Screening. Toxicology in Vitro 2021, 73 (October 2020), 105109. 10.1016/j.tiv.2021.105109.

(18) Franzosa, J. A.; Bonzo, J. A.; Jack, J.; Baker, N. C.; Kothiya, P.; Witek, R. P.; Hurban, P.; Siferd, S.; Hester, S.; Shah, I.; Ferguson, S. S.; Houck, K. A.; Wambaugh, J. F. High-Throughput Toxicogenomic Screening of Chemicals in the Environment Using Metabolically Competent Hepatic Cell Cultures. NPJ Syst Biol Appl 2021, 7 (1), 7. 10.1038/s41540-020-00166-2.

(19) DeGroot, D. E.; Swank, A.; Thomas, R. S.; Strynar, M.; Lee, M.-Y.; Carmichael, P. L.; Simmons, S. O. MRNA Transfection Retrofits Cell-Based Assays with Xenobiotic Metabolism. J Pharmacol Toxicol Methods 2018, 92 (January), 77–94. 10.1016/j.vascn.2018.03.002.

(20) Yoshitomi, S.; Ikemoto, K.; Takahashi, J.; Miki, H.; Namba, M.; Asahi, S. Establishment of the Transformants Expressing Human Cytochrome P450 Subtypes in HepG2, and Their Applications on Drug Metabolism and Toxicology. Toxicology in Vitro 2001, 15 (3), 245–256. 10.1016/S0887-2333(01)00011-X.

(21) Li, X.; He, X.; Le, Y.; Guo, X.; Bryant, M. S.; Atrakchi, A. H.; McGovern, T. J.; Davis-Bruno, K. L.; Keire, D. A.; Heflich, R. H.; Mei, N. Genotoxicity Evaluation of Nitrosamine Impurities Using Human TK6 Cells Transduced with Cytochrome P450s. Arch Toxicol 2022, 96 (11), 3077–3089. 10.1007/s00204-022-03347-6.

(22) Hagiwara, Y.; Watanabe, M.; Oda, Y.; Sofuni, T.; Nohmi, T. Specificity and Sensitivity of Salmonella Typhimurium YG1041 and YG1042 Strains Possesing Elevated Levels of Both Nitroreductase and Acetyltransferase Activity. Mutation Research/Environmental Mutagenesis and Related Subjects 1993, 291 (3), 171–180. 10.1016/0165-1161(93)90157-U.

(23) Walton, A. J. THE EFFECT OF VARIOUS TISSUE EXTRACTS UPON THE GROWTH OF ADULT MAMMALIAN CELLS IN VITRO. Journal of Experimental Medicine 1914, 20 (6), 554–572. 10.1084/jem.20.6.554.

(24) Mueller, G. C.; Miller, J. A. THE METABOLISM OF 4-DIMETHYLAMINOAZOBENZENE BY RAT LIVER HOMOGENATES. Journal of Biological Chemistry 1948, 176 (2), 535–544. 10.1016/S0021-9258(19)52671-0.

(25) UN. Stockholm Convention on Persistent Organic Pollutants. In United Nations Treaty Collection; 2004. URL: https://treaties.un.org/pages/ViewDetails.aspx?src=TREATY&mtdsg_no=XXVII-15&chapter=27&clang=_en

(26) Ames, B. N.; Durston, W. E.; Yamasaki, E.; Lee, F. D. Carcinogens Are Mutagens: A Simple Test System Combining Liver Homogenates for Activation and Bacteria for Detection. Proceedings of the National Academy of Sciences 1973, 70 (8), 2281–2285. 10.1073/pnas.70.8.2281.

(27) Ames, B. N.; McCann, J.; Yamasaki, E. Methods for Detecting Carcinogens and Mutagens with the Salmonella/Mammalian-Microsome Mutagenicity Test. Mutation Research/Environmental Mutagenesis and Related Subjects 1975, 31 (6), 347–363. 10.1016/0165-1161(75)90046-1.

(28) Maron, D. M.; Ames, B. N. Revised Methods for the Salmonella Mutagenicity Test. Mutation Research/Environmental Mutagenesis and Related Subjects 1983, 113 (3–4), 173–215. 10.1016/0165-1161(83)90010-9.

(29) Dunkel, V. C.; Zeiger, E.; Brusick, D.; McCoy, E.; McGregor, D.; Mortelmans, K.; Rosenkranz, H. S.; Simmon, V. F. Reproducibility of Microbial Mutagenicity Assays: II. Testing of Carcinogens and Noncarcinogens In Salmonella Typhimurium And Escherichia Coli. Environ Mutagen 1985, 7 (S5), 1–19. 10.1002/em.2860070902.

(30) Dunkel, V. C.; Zeiger, E.; Brusick, D.; McCoy, E.; McGregor, D.; Mortelmans, K.; Rosenkranz, H. S.; Simmon, V. F. Reproducibility of Microbial Mutagenicity Assays: I. Tests With Salmonella Typhimurium And Escherichia Coli Using a Standardized Protocol. Environ Mutagen 1984, 6 (S2), 201–251. 10.1002/em.2860060706.

(31) Hubbard, S. A.; Brooks, T. M.; Gonzalez, L. P.; Bridges, J. W. Preparation and Characterisation of S9 Fractions. In Comparative Genetic Toxicology; Palgrave Macmillan UK: London, 1985; pp 413–438. 10.1007/978-1-349-07901-8_50.

(32) Poiley, J. A.; Raineri, R. Species Specificity Affects the Choice of S9 Preparations for Use in the Hamster Embryo Cell Transformation System. In Vitro 1984, 20 (8), 602–606. 10.1007/BF02619608.

(33) Shahin, M. M. Studies on the Mutagenicity of Aniline in Association with Norharman in the Salmonella /Mammalian Microscome Assay. Int J Cosmet Sci 1989, 11 (3), 129–140. 10.1111/j.1467-2494.1989.tb00502.x.

(34) JRC; Coecke, S.; Bartnicka, J.; Langezaal, I. Towards Animal-Free in Vitro Methods in the Thyroid Validation Study: Final Report; Publications Office, 2021. doi10.2760/544332

(35) Page, M. J.; Moher, D.; Bossuyt, P. M.; Boutron, I.; Hoffmann, T. C.; Mulrow, C. D.; Shamseer, L.; Tetzlaff, J. M.; Akl, E. A.; Brennan, S. E.; Chou, R.; Glanville, J.; Grimshaw, J. M.; Hróbjartsson, A.; Lalu, M. M.; Li, T.; Loder, E. W.; Mayo-Wilson, E.; McDonald, S.; McGuinness, L. A.; Stewart, L. A.; Thomas, J.; Tricco, A. C.; Welch, V. A.; Whiting, P.; McKenzie, J. E. PRISMA 2020 Explanation and Elaboration: Updated Guidance and Exemplars for Reporting Systematic Reviews. BMJ 2021, 372, n160. 10.1136/bmj.n160.

(36) Page, M. J.; McKenzie, J. E.; Bossuyt, P. M.; Boutron, I.; Hoffmann, T. C.; Mulrow, C. D.; Shamseer, L.; Tetzlaff, J. M.; Akl, E. A.; Brennan, S. E.; Chou, R.; Glanville, J.; Grimshaw, J. M.; Hróbjartsson, A.; Lalu, M. M.; Li, T.; Loder, E. W.; Mayo-Wilson, E.; McDonald, S.; McGuinness, L. A.; Stewart, L. A.; Thomas, J.; Tricco, A. C.; Welch, V. A.; Whiting, P.; Moher, D. The PRISMA 2020 Statement: An Updated Guideline for Reporting Systematic Reviews. BMJ 2021, 372, n71. 10.1136/bmj.n71.

(37) James, K. L.; Randall, N. P.; Haddaway, N. R. A Methodology for Systematic Mapping in Environmental Sciences. Environ Evid 2016, 5 (1), 7. 10.1186/s13750-016-0059-6.

(38) Wolffe, T. A. M.; Whaley, P.; Halsall, C.; Rooney, A. A.; Walker, V. R. Systematic Evidence Maps as a Novel Tool to Support Evidence-Based Decision-Making in Chemicals Policy and Risk Management. Environ Int 2019, 130 (June), 104871. 10.1016/j.envint.2019.05.065.

(39) Tricco, A. C.; Lillie, E.; Zarin, W.; O’Brien, K. K.; Colquhoun, H.; Levac, D.; Moher, D.; Peters, M. D. J.; Horsley, T.; Weeks, L.; Hempel, S.; Akl, E. A.; Chang, C.; McGowan, J.; Stewart, L.; Hartling, L.; Aldcroft, A.; Wilson, M. G.; Garritty, C.; Lewin, S.; Godfrey, C. M.; Macdonald, M. T.; Langlois, E. V.; Soares-Weiser, K.; Moriarty, J.; Clifford, T.; Tunçalp, Ö.; Straus, S. E. PRISMA Extension for Scoping Reviews (PRISMA-ScR): Checklist and Explanation. Ann Intern Med 2018, 169 (7), 467–473. 10.7326/M18-0850.

(40) Food, E.; Authority, S. Application of Systematic Review Methodology to Food and Feed Safety Assessments to Support Decision Making. EFSA Journal 2010, 8 (6). 10.2903/j.efsa.2010.1637.

(41) NTP-OHAT. Handbook for Conducting a Literature-Based Health Assessment Using OHAT Approach for Systematic Review and Evidence Integration; 2019. https://ntp.niehs.nih.gov/go/ohathandbook; URL: https://ntp.niehs.nih.gov/sites/default/files/ntp/ohat/pubs/handbookmarch2019_508.pdf

(42) US EPA. Application of Systematic Review in Tsca Risk Evaluations; 2018. https://www.epa.gov/assessing-and-managing-chemicals-under-tsca/draft-protocol-systematic-review-tsca-risk-evaluations. URL: https://www.epa.gov/sites/default/files/2018-06/documents/final_application_of_sr_in_tsca_05-31-18.pdf

(43) Munn, Z.; Peters, M. D. J.; Stern, C.; Tufanaru, C.; McArthur, A.; Aromataris, E. Systematic Review or Scoping Review? Guidance for Authors When Choosing between a Systematic or Scoping Review Approach. BMC Med Res Methodol 2018, 18 (1), 143. 10.1186/s12874-018-0611-x.

(44) Khalil, H.; Tricco, A. C. Differentiating between Mapping Reviews and Scoping Reviews in the Evidence Synthesis Ecosystem. J Clin Epidemiol 2022, 149, 175–182. 10.1016/j.jclinepi.2022.05.012.

(45) Moher, D.; Shamseer, L.; Clarke, M.; Ghersi, D.; Liberati, A.; Petticrew, M.; Shekelle, P.; Stewart, L. A. Preferred Reporting Items for Systematic Review and Meta-Analysis Protocols (PRISMA-P) 2015 Statement. Syst Rev 2015, 4 (1), 1. 10.1186/2046-4053-4-1.

(46) Shamseer, L.; Moher, D.; Clarke, M.; Ghersi, D.; Liberati, A.; Petticrew, M.; Shekelle, P.; Stewart, L. A. Preferred Reporting Items for Systematic Review and Meta-Analysis Protocols (PRISMA-P) 2015: Elaboration and Explanation. BMJ 2015, 349 (jan02 1), g7647–g7647. 10.1136/bmj.g7647.

(47) Lungu-Mitea, S.; Åslund, M. S.; Reichstein, I.; Vidal, F.; Schiwy, A.; Hollert, H.; Jacobs, M. N.; Hilscherová, K. On the Utilisation and Characterisation of External Biotransformation Systems in in Vitro Toxicology: Protocol for a Critical Review. August 17, 2023. 10.6084/m9.figshare.21494616.v2.

(48) OECD. Revised Guidance Document 150 on Standardised Test Guidelines for Evaluating Chemicals for Endocrine Disruption; OECD Series on Testing and Assessment; OECD, 2018. 10.1787/9789264304741-en.

(49) Posit team. RStudio: Integrated Development Environment for R. Posit Software, PBC: Boston, MA 2023. http://www.posit.co/.

(50) Team, R. C. R: A Language and Environment for Statistical Computing. R Foundation for Statistical Computing, Vienna, Austria. 2023. https://www.r-project.org/.

(51) Larsson, J. Eulerr: Area-Proportional Euler and Venn Diagrams with Ellipses. 2022. https://cran.r-project.org/package=eulerr.

(52) Gehlenborg, N. UpSetR: A More Scalable Alternative to Venn and Euler Diagrams for Visualizing Intersecting Sets. 2019. https://cran.r-project.org/package=UpSetR.

(53) Wickham, H. Ggplot2: Elegant Graphics for Data Analysis; Springer New York: New York, 2016. 10.1007/978-3-319-24277-4

(54) Lê, S.; Josse, J.; Husson, F. FactoMineR : An R Package for Multivariate Analysis. J Stat Softw 2008, 25 (1), 253–258. 10.18637/jss.v025.i01.

(55) Hahsler, M.; Buchta, C.; Gruen, B.; Hornik, K. Arules: Mining Association Rules and Frequent Itemsets. 2023. https://cran.r-project.org/package=arules.

(56) Hashler, M. ArulesViz: Visualizing Association Rules and Frequent Itemsets. 2023. https://cran.r-project.org/package=arulesViz.

(57) Hair, J. F.; Black, W. C.; Babin, B. J.; Anderson, R. E. Multivariate Data Analysis; Cengage, 2019. ISBN: 9781473756557

(58) Chelcea, I.; Örn, S.; Hamers, T.; Koekkoek, J.; Legradi, J.; Vogs, C.; Andersson, P. L. Physiologically Based Toxicokinetic Modeling of Bisphenols in Zebrafish (Danio Rerio) Accounting for Variations in Metabolic Rates, Brain Distribution, and Liver Accumulation. Environ Sci Technol 2022, 56 (14), 10216–10228. 10.1021/acs.est.2c01292.

(59) Harding, C.; Viljanto, M.; Habershon-Butcher, J.; Taylor, P.; Scarth, J. Equine Metabolism of the Selective Androgen Receptor Modulator YK-11 in Urine and Plasma Following Oral Administration. Drug Test Anal 2023, 15 (4), 388–407. 10.1002/dta.3425.

(60) Zheng, Z.; Peters, G. M.; Arp, H. P. H.; Andersson, P. L. Combining in Silico Tools with Multicriteria Analysis for Alternatives Assessment of Hazardous Chemicals: A Case Study of Decabromodiphenyl Ether Alternatives. Environ Sci Technol 2019, 53 (11), 6341–6351. 10.1021/acs.est.8b07163.

(61) Zheng, Z.; Arp, H. P. H.; Peters, G.; Andersson, P. L. Combining in Silico Tools with Multicriteria Analysis for Alternatives Assessment of Hazardous Chemicals: Accounting for the Transformation Products of DecaBDE and Its Alternatives. Environ Sci Technol 2021, 55 (2), 1088–1098. 10.1021/acs.est.0c02593.

(62) Miller, G. E.; Brabec, M. J.; Kulkarni, A. P. Mutagen Activation of 1,2-dibromo-3-chloropropane by Cytosolic Glutathione S-transferases and Microsomal Enzymes. J Toxicol Environ Health 1986, 19 (4), 503–518. 10.1080/15287398609530948.

(63) Otto, M.; Hansen, S. H.; Dalgaard, L.; Dubois, J.; Badolo, L. Development of an in Vitro Assay for the Investigation of Metabolism-Induced Drug Hepatotoxicity. Cell Biol Toxicol 2008, 24 (1), 87–99. 10.1007/s10565-007-9018-x.

(64) Kropf, C.; Begnaud, F.; Gimeno, S.; Berthaud, F.; Debonneville, C.; Segner, H. In Vitro Biotransformation Assays Using Liver S9 Fractions and Hepatocytes from Rainbow Trout (Oncorhynchus Mykiss): Overcoming Challenges with Difficult to Test Fragrance Chemicals. Environ Toxicol Chem 2020, 39 (12), 2396–2408. 10.1002/etc.4872.

(65) Ku, W. W.; Bigger, A.; Brambilla, G.; Glatt, H.; Gocke, E.; Guzzie, P. J.; Hakura, A.; Honma, M.; Martus, H.-J.; Obach, R. S.; Roberts, S. Strategy for Genotoxicity Testing—Metabolic Considerations. Mutation Research/Genetic Toxicology and Environmental Mutagenesis 2007, 627 (1), 59–77. 10.1016/j.mrgentox.2006.10.004.

(66) Richardson, S.; Bai, A.; A. Kulkarni, A.; F. Moghaddam, M. Efficiency in Drug Discovery: Liver S9 Fraction Assay As a Screen for Metabolic Stability. Drug Metab Lett 2016, 10 (2), 83–90. 10.2174/1872312810666160223121836.

(67) Abdi, H.; Valentin, D. Multiple Correspondence Analysis. Encyclopedia of measurement and statistics 2007, 2 (4), 651–657. URL: https://personal.utdallas.edu/~herve/Abdi-MCA2007-pretty.pdf

(68) Lebart, L. Validation Techniques in Multiple Correspondence Analysis; 2006; pp 179–195. 10.1201/9781420011319.ch7.

(69) Louisse, J.; Alewijn, M.; Peijnenburg, A. A. C. M.; Cnubben, N. H. P.; Heringa, M. B.; Coecke, S.; Punt, A. Towards Harmonization of Test Methods for in Vitro Hepatic Clearance Studies. Toxicology in Vitro 2020, 63 (August 2019), 104722. 10.1016/j.tiv.2019.104722.

(70) Galloway, S. M.; Aardema, M. J.; Ishidate, M.; Ivett, J. L.; Kirkland, D. J.; Morita, T.; Mosesso, P.; Sofuni, T. Report from Working Group on in Vitro Tests for Chromosomal Aberrations. Mutat Res 1994. DOI: 10.1016/0165-1161(94)00012-3

(71) Tice, R. R.; Agurell, E.; Anderson, D.; Burlinson, B.; Hartmann, A; Kobayashi, H.; Miyamae, Y.; Rojas, E.; Ryu, J. C.; Sasaki, Y. F. Single Cell Gel/Comet Assay: Guidelines for in Vitro and in Vivo Genetic Toxicology Testing. Environ Mol Mutagen 2000, 35 (3), 206–221. DOI: 10.1002/(sici)1098-2280(2000)35:3<206::aid-em8>3.0.co;2-j

(72) Kirsch-Volders, M.; Sofuni, T.; Aardema, M.; Albertini, S.; Eastmond, D.; Fenech, M.; Ishidate, M.; Kirchner, S.; Lorge, E.; Morita, T.; Norppa, H.; Surrallés, J.; Vanhauwaert, A.; Wakata, A. Report from the in Vitro Micronucleus Assay Working Group. Mutation Research/Genetic Toxicology and Environmental Mutagenesis 2003, 540 (2), 153–163. 10.1016/j.mrgentox.2003.07.005.

(73) Lebsanft, J.; McMahon, J. B.; Steinmann, G. G.; Shoemaker, R. H. A Rapid in Vitro Method for the Evaluation of Potential Antitumor Drugs Requiring Metabolic Activation by Hepatic S9 Enzymes. Biochem Pharmacol 1989. DOI: 10.1016/0006-2952(89)90659-x

(74) Escher, B. I.; Cowan-Ellsberry, C. E.; Dyer, S.; Embry, M. R.; Erhardt, S.; Halder, M.; Kwon, J.-H.; Johanning, K.; Oosterwijk, M. T. T.; Rutishauser, S.; Segner, H.; Nichols, J. Protein and Lipid Binding Parameters in Rainbow Trout (Oncorhynchus Mykiss) Blood and Liver Fractions to Extrapolate from an in Vitro Metabolic Degradation Assay to in Vivo Bioaccumulation Potential of Hydrophobic Organic Chemicals. Chem Res Toxicol 2011, 24 (7), 1134–1143. 10.1021/tx200114y.

(75) Kwon, J.-H.; Lee, H.-J.; Escher, B. I. Bioavailability of Hydrophobic Organic Chemicals on an in Vitro Metabolic Transformation Using Rat Liver S9 Fraction. Toxicology in Vitro 2020, 66 (November 2019), 104835. 10.1016/j.tiv.2020.104835.

(76) Proença, S.; Escher, B. I.; Fischer, F. C.; Fisher, C.; Grégoire, S.; Hewitt, N. J.; Nicol, B.; Paini, A.; Kramer, N. I. Effective Exposure of Chemicals in in Vitro Cell Systems: A Review of Chemical Distribution Models. Toxicology in Vitro 2021, 73 (February), 105133. 10.1016/j.tiv.2021.105133.

(77) Shao, Y.; Schiwy, A.; Glauch, L.; Henneberger, L.; König, M.; Mühlenbrink, M.; Xiao, H.; Thalmann, B.; Schlichting, R.; Hollert, H.; Escher, B. I. Optimization of a Pre-Metabolization Procedure Using Rat Liver S9 and Cell-Extracted S9 in the Ames Fluctuation Test. Science of The Total Environment 2020, 749, 141468. 10.1016/j.scitotenv.2020.141468.

(78) Jia, L.; Liu, X. The Conduct of Drug Metabolism Studies Considered Good Practice (II): In Vitro Experiments. Curr Drug Metab 2007, 8 (8), 822–829. 10.2174/138920007782798207.

(79) Langsch, A.; Nau, H. In Vitro Metabolism : Applications in Pharmacology and Toxicology Metabolic Activation for In Vitro Systems. ALTEX 2005, No. 2, 354–358.

(80) Myhr, B. C.; Mayo, J. K. Mutagenicity of Rat-Liver S9 to L5178Y Mouse Lymphoma Cells. Mutation Research/Genetic Toxicology 1987, 189 (1), 27–37. 10.1016/0165-1218(87)90030-9.

(81) Kamath, S. A.; Kummerow, F. A.; Narayan, K. A. A Simple Procedure for the Isolation of Rat Liver Microsomes. FEBS Lett 1971, 17 (1), 90–92. 10.1016/0014-5793(71)80571-9.

(82) Garner, R. C.; Miller, E. C.; Miller, J. A. Liver Microsomal Metabolism of Aflatoxin B 1 to a Reactive Derivative Toxic to Salmonella Typhimurium TA 1530. Cancer Res 1972, 32 (10), 2058–2066. PMID: 4404160

(83) Liewen, M. B.; Marth, E. H. Evaluation of 1,3-Pentadiene for Mutagenicity by the Salmonella/Mammalian Microsome Assay. Mutation Research/Genetic Toxicology 1985, 157 (1), 49–52. 10.1016/0165-1218(85)90048-5.

(84) Lynch, B.; Lau, A.; Baldwin, N.; Hofman-Hüther, H.; Bauter, M. R.; Marone, P. A. Genotoxicity of Dried Hoodia Parviflora Aerial Parts. Food and Chemical Toxicology 2013, 55, 272–278. 10.1016/j.fct.2013.01.014.

(85) Sarrif, A. M.; Arce, G. T.; Krahn, D. F.; O’Neil, R. M.; Reynolds, V. L. Evaluation of Carbendazim for Gene Mutations in the Salmonella/Ames Plate-Incorporation Assay: The Role of Aminophenazine Impurities. Mutation Research/Genetic Toxicology 1994, 321 (1–2), 43–56. 10.1016/0165-1218(94)90119-8.

(86) OECD. Test No. 473: In Vitro Mammalian Chromosomal Aberration Test; OECD Guidelines for the Testing of Chemicals, Section 4; OECD, 2016. 10.1787/9789264264649-en.

(87) OECD. Test No. 476: In Vitro Mammalian Cell Gene Mutation Tests Using the Hprt and Xprt Genes; OECD Guidelines for the Testing of Chemicals, Section 4; OECD, 2016. 10.1787/9789264264809-en.

(88) OECD. Test No. 487: In Vitro Mammalian Cell Micronucleus Test; OECD Publishing, 2014. 10.1787/9789264224438-en.

(89) COM UK GOV. Guidance on a Strategy for Genotoxicity Testing of Chemicals; 2021. https://www.gov.uk/government/publications/a-strategy-for-testing-of-chemicals-for-genotoxicity. URL: https://assets.publishing.service.gov.uk/media/61c316cdd3bf7f1f71aa7ca7/strategy-for-genotoxicity-testing-of-chemicals-guidance.pdf

(90) COM UK GOV. Guidance on a Strategy for Genotoxicity Testing of Chemical Substances - in Vitro Genotoxicity Testing; 2021. https://www.gov.uk/government/publications/a-strategy-for-testing-of-chemicals-for-genotoxicity. URL: https://assets.publishing.service.gov.uk/media/61c3176fe90e071971e255fa/stage-1-in-vitro-genotoxicity-testing.pdf

(91) Dhawan, A.; Bajpayee, M. Genotoxicity Assessment; Dhawan, A., Bajpayee, M., Eds.; Methods in Molecular Biology; Springer New York: New York, NY, 2019; Vol. 2031. 10.1007/978-1-4939-9646-9.

(92) Mollergues, J.; Van Vugt-Lussenburg, B.; Kirchnawy, C.; Bandi, R. A.; Van Der Lee, R. B.; Marin-Kuan, M.; Schilter, B.; Fussell, K. C. Incorporation of a Metabolizing System in Biodetection Assays for Endocrine Active Substances. ALTEX 2017, 34 (3), 389–398. 10.14573/altex.1611021.

(93) van Vugt-Lussenburg, B. M. A.; van der Lee, R. B.; Man, H.-Y.; Middelhof, I.; Brouwer, A.; Besselink, H.; van der Burg, B. Incorporation of Metabolic Enzymes to Improve Predictivity of Reporter Gene Assay Results for Estrogenic and Anti-Androgenic Activity. Reproductive Toxicology 2018, 75, 40–48. 10.1016/j.reprotox.2017.11.005.

(94) Brendt, J.; Crawford, S. E.; Velki, M.; Xiao, H.; Thalmann, B.; Hollert, H.; Schiwy, A. Is a Liver Comparable to a Liver? A Comparison of Different Rat-Derived S9-Fractions with a Biotechnological Animal-Free Alternative in the Ames Fluctuation Assay. Science of The Total Environment 2020, No. xxxx, 143522. 10.1016/j.scitotenv.2020.143522.

(95) Reichstein, I. S.; Becker, A. H.; Johann, S.; Braunbeck, T.; Schiwy, S.; Hollert, H.; Schiwy, A. Biotechnological Metabolization System Has the Potential to Improve the Predictive Ability of the Fish Embryo Acute Toxicity (FET) Test with the Zebrafish (Danio Rerio). Environ Sci Eur 2024, 36 (1), 91. 10.1186/s12302-024-00913-w.

(96) Reichstein, I. S.; König, M.; Wojtysiak, N.; Escher, B. I.; Henneberger, L.; Behnisch, P.; Besselink, H.; Thalmann, B.; Colas, J.; Hörchner, S.; Hollert, H.; Schiwy, A. Replacing Animal-Derived Components in in Vitro Test Guidelines OECD 455 and 487. Science of The Total Environment 2023, 868 (January), 161454. 10.1016/j.scitotenv.2023.161454.

(97) Gonzalez, R. J.; Tarloff, J. B. Evaluation of Hepatic Subcellular Fractions for Alamar Blue and MTT Reductase Activity. Toxicology in Vitro 2001, 15 (3), 257–259. 10.1016/S0887-2333(01)00014-5.

(98) Brendt, J.; Lackmann, C.; Heger, S.; Velki, M.; Crawford, S. E.; Xiao, H.; Thalmann, B.; Schiwy, A.; Hollert, H. Using a High-Throughput Method in the Micronucleus Assay to Compare Animal-Free with Rat-Derived S9. Science of The Total Environment 2021, 751, 142269. 10.1016/j.scitotenv.2020.142269.

(99) Vignati, L.; Turlizzi, E.; Monaci, S.; Grossi, P.; Kanter, R. De; Monshouwer, M. An in Vitro Approach to Detect Metabolite Toxicity Due to CYP3A4-Dependent Bioactivation of Xenobiotics. Toxicology 2005, 216 (2–3), 154–167. 10.1016/j.tox.2005.08.003.

(100) Géniès, C.; Jacques-Jamin, C.; Duplan, H.; Rothe, H.; Ellison, C.; Cubberley, R.; Schepky, A.; Lange, D.; Klaric, M.; Hewitt, N. J.; Grégoire, S.; Arbey, E.; Fabre, A.; Eilstein, J. Comparison of the Metabolism of 10 Cosmetics-relevant Chemicals in EpiSkinTM S9 Subcellular Fractions and in Vitro Human Skin Explants. Journal of Applied Toxicology 2020, 40 (2), 313–326. 10.1002/jat.3905.

(101) Hakura, A.; Suzuki, S.; Sawada, S.; Sugihara, T.; Hori, Y.; Uchida, K.; Kerns, W. D.; Sagami, F.; Motooka, S.; Satoh, T. Use of Human Liver S9 in the Ames Test: Assay of Three Procarcinogens Using Human S9 Derived from Multiple Donors. Regulatory Toxicology and Pharmacology 2003, 37 (1), 20– 27. 10.1016/S0273-2300(02)00024-7.

(102) Jacobs, M. N.; Bult, J. M.; Cavanagh, K.; Chesne, C.; Delrue, N.; Fu, J.; Grange, E.; Langezaal, I.; Misztela, D.; Murray, J.; Paparella, M.; Stoddart, G.; Tonn, T.; Treasure, C.; Tsukano, M.; Versteegen, R. OECD Workshop Consensus Report: Ethical Considerations with Respect to Human Derived Products, Specifically Human Serum, in OECD Test Guidelines. Frontiers in Toxicology 2023, 5 (February), 1–10. 10.3389/ftox.2023.1140698.

(103) Johanning, K.; Hancock, G.; Escher, B.; Adekola, A.; Bernhard, M. J.; Cowan-Ellsberry, C.; Domoradzki, J.; Dyer, S.; Eickhoff, C.; Embry, M.; Erhardt, S.; Fitzsimmons, P.; Halder, M.; Hill, J.; Holden, D.; Johnson, R.; Rutishauser, S.; Segner, H.; Schultz, I.; Nichols, J. Assessment of Metabolic Stability Using the Rainbow Trout (Oncorhynchus Mykiss) Liver S9 Fraction. Curr Protoc Toxicol 2012, 53 (1), 1–28. 10.1002/0471140856.tx1410s53.

(104) OECD. Test No. 319B: Determination of in Vitro Intrinsic Clearance Using Rainbow Trout Liver S9 Sub-Cellular Fraction (RT-S9); OECD Guidelines for the Testing of Chemicals, Section 3; OECD, 2018; Vol. Section 3,. 10.1787/9789264303232-en.

(105) OECD. Guidance Document on Good In Vitro Method Practices (GIVIMP); OECD Series on Testing and Assessment; OECD, 2018. 10.1787/9789264304796-en.

(106) OECD. Guidance Document for Describing Non-Guideline In Vitro Test Methods; OECD Series on Testing and Assessment; OECD, 2014. 10.1787/9789264274730-en.

(107) Ladd, M. A.; Fitzsimmons, P. N.; Nichols, J. W. Optimization of a UDP-Glucuronosyltransferase Assay for Trout Liver S9 Fractions: Activity Enhancement by Alamethicin, a Pore-Forming Peptide. Xenobiotica 2016, 46 (12), 1066–1075. 10.3109/00498254.2016.1149634.

(108) Brendt, J.; Crawford, S. E.; Velki, M.; Xiao, H.; Thalmann, B.; Hollert, H.; Schiwy, A. Is a Liver Comparable to a Liver? A Comparison of Different Rat-Derived S9-Fractions with a Biotechnological Animal-Free Alternative in the Ames Fluctuation Assay. Science of The Total Environment 2021, 759 (xxxx), 143522. 10.1016/j.scitotenv.2020.143522.

(109) Takehisa, S.; Kanaya, N.; Rieger, R. Promutagen Activation by Vicia Faba: An Assay Based on the Induction of Sister-Chromatid Exchanges in Chinese Hamster Ovary Cells. Mutation Research/Fundamental and Molecular Mechanisms of Mutagenesis 1988, 197 (2), 195–205. 10.1016/0027-5107(88)90093-0.

(110) Johnson, T. E.; Umbenhauer, D. R.; Galloway, S. M. Human Liver S-9 Metabolic Activation: Proficiency in Cytogenetic Assays and Comparison with Phenobarbital/β-Naphthoflavone or Aroclor 1254 Induced RatS-9. Environ Mol Mutagen 1996, 28 (1), 51–59. 10.1002/(SICI)1098-2280(1996)28:1<51::AID-EM8>3.0.CO;2-H.

(111) Pelkonen, O. Comparison of Metabolic Stability and Metabolite Identification of 55 ECVAM/ICCVAM Validation Compounds between Human and Rat Liver Homogenates and Microsomes - a Preliminary Analysis. ALTEX 2009, 26 (3), 214–222. 10.14573/altex.2009.3.214.

(112) Cox, J. A.; Fellows, M. D.; Hashizume, T.; White, P. A. The Utility of Metabolic Activation Mixtures Containing Human Hepatic Post-Mitochondrial Supernatant (S9) for in Vitro Genetic Toxicity Assessment. Mutagenesis 2016, 31 (2), 117–130. 10.1093/mutage/gev082.

(113) Kishida, T.; Muto, S.; Hayashi, M.; Tsutsui, M.; Tanaka, S.; Murakami, M.; Kuroda, J. Strain Differences in Hepatic Cytochrome P450 1A and 3A Expression between Sprague-Dawley and Wistar Rats. J Toxicol Sci 2008, 33 (4), 447–457. 10.2131/jts.33.447.

(114) Rudeck, J.; Bert, B.; Marx-Stoelting, P.; Schönfelder, G.; Vogl, S. Liver Lobe and Strain Differences in the Activity of Murine Cytochrome P450 Enzymes. Toxicology 2018, 404–405 (February), 76–85. 10.1016/j.tox.2018.06.001.

(115) Eggleston, L. V.; Krebs, H. A. Strain Differences in the Activities of Rat Liver Enzymes. Biochemical Journal 1969, 114 (4), 877–879. 10.1042/bj1140877.

(116) Matsui, M.; Nagai, F.; Aoyagi, S. Strain Differences in Rat Liver (UDP-Glucuronyltransferase Activity towards Androsterone. Biochemical Journal 1979, 179 (3), 483–487. 10.1042/bj1790483.

(117) DeNucci, S. M.; Tong, M.; Longato, L.; Lawton, M.; Setshedi, M.; Carlson, R. I.; Wands, J. R.; de la Monte, S. M. Rat Strain Differences in Susceptibility to Alcohol-Induced Chronic Liver Injury and Hepatic Insulin Resistance. Gastroenterol Res Pract 2010, 2010, 1–16. 10.1155/2010/312790.

(118) Sumida, K.; Ooe, N.; Nagahori, H.; Saito, K.; Isobe, N.; Kaneko, H.; Nakatsuka, I. An in Vitro Reporter Gene Assay Method Incorporating Metabolic Activation with Human and Rat S9 or Liver Microsomes. Biochem Biophys Res Commun 2001, 280 (1), 85–91. 10.1006/bbrc.2000.4071.

(119) Schenkman, J. B.; Frey, I.; Remmer, H.; Estabrook, R. W. Sex Differences in Drug Metabolism by Rat Liver Microsomes. Mol Pharmacol 1967, 3 (6), 516–525. PMID: 6059868

(120) Hales, B. F.; Jain, R. Effects of Phenobarbital and β-Naphthoflavone on the Activation of Cyclophosphamide to Mutagenic Metabolites in Vitro by Liver and Kidney from Male and Female Rats. Biochem Pharmacol 1980, 29 (14), 2031–2037. 10.1016/0006-2952(80)90488-8.

(121) Ou, Y.; Lin, J. Biotransformation of Butachlor through Mercapturic Acid Pathway in Rat Tissue Homogenates. J Toxicol Environ Health 1992, 35 (1), 19–28. 10.1080/15287399209531590.

(122) Winckler, K.; Obe, G.; Madle, S.; Nau, H. Mutagenic Activities of Cyclophosphamide (NSC-26271) and Its Main Metabolites in Salmonella Typhimurium, Human Peripheral Lymphocytes and Chinese Hamster Ovary Cells. Mutation Research/Fundamental and Molecular Mechanisms of Mutagenesis 1984, 129 (1), 47–55. 10.1016/0027-5107(84)90122-2.

(123) Kitamura, S.; Sanoh, S.; Kohta, R.; Suzuki, T.; Sugihara, K.; Fujimoto, N.; Ohta, S. Metabolic Activation of Proestrogenic Diphenyl and Related Compounds by Rat Liver Microsomes. Journal of Health Science 2003, 49 (4), 298–310. 10.1248/jhs.49.298.

(124) Kitamura, S.; Ohmegi, M.; Sanoh, S.; Sugihara, K.; Yoshihara, S.; Fujimoto, N.; Ohta, S. Estrogenic Activity of Styrene Oligomers after Metabolic Activation by Rat Liver Microsomes. Environ Health Perspect 2003, 111 (3), 329–334. 10.1289/ehp.5723.

(125) Callander, R. D.; Mackay, J. M.; Clay, P.; Elcombe, C. R.; Elliott, B. M. Evaluation of Phenobarbital/β-Naphthoflavone as an Alternative S9-Induction Regime to Aroclor 1254 in the Rat for Use in in Vitro Genotoxicity Assays. Mutagenesis 1995, 10 (6), 517–522. 10.1093/mutage/10.6.517.

(126) Franco, S. G. Alternatives in the Induction and Preparation of Phenobarbital/Naphthoflavone-Induced S9 and Their Activation Profiles. Mutagenesis 1999, 14 (3), 323–326. 10.1093/mutage/14.3.323.

(127) Johnson, G. W.; Hansen, L. G.; Hamilton, M. C.; Fowler, B.; Hermanson, M. H. PCB, PCDD and PCDF Congener Profiles in Two Types of Aroclor 1254. Environ Toxicol Pharmacol 2008, 25 (2), 156–163. 10.1016/j.etap.2007.10.011.

(128) Doostdar, H.; Duthie, S. J.; Burke, M. D.; Melvin, W. T.; Grant, M. H. The Influence of Culture Medium Composition on Drug Metabolising Enzyme Activities of the Human Liver Derived Hep G2 Cell Line. FEBS Lett 1988, 241 (1–2), 15–18. 10.1016/0014-5793(88)81021-4.

(129) Hewitt, N. J.; Hewitt, P. Phase I and II Enzyme Characterization of Two Sources of HepG2 Cell Lines. Xenobiotica 2004, 34 (3), 243–256. 10.1080/00498250310001657568.

(130) Easterbrook, J.; Lu, C.; Sakai, Y.; Li, A. P. Effects of Organic Solvents on the Activities of Cytochrome P450 Isoforms, UDP-Dependent Glucuronyl Transferase, and Phenol Sulfotransferase in Human Hepatocytes. Drug Metab Dispos 2001, 29 (2), 141–144.

(131) Lindblad, W. J.; Jackim, E. Mechanism for the Differential Induction of Mutation by S9 Activated Benzo[a]Pyrene Employing Either a Glucose-6-Phosphate-Dependent NADPH-Regenerating System or an Isocitrate-Dependent System. Mutation Research/Fundamental and Molecular Mechanisms of Mutagenesis 1982, 96 (1), 109–118. 10.1016/0027-5107(82)90021-5.

(132) Bimboes Detlev; Greim Helmut. Human Lymphocytes as Target Cells in a Metabolizing Test System in Vitro for Detecting Potential Mutagens. Mutation Research/Fundamental and Molecular Mechanisms of Mutagenesis 1976, 35 (1), 155–159. 10.1016/0027-5107(76)90177-9.

(133) OECD. Guidance Document On The Determination Of In Vitro Intrinsic Clearance Using Cryopreserved Hepatocytes (RT-HEP) Or Liver S9 Sub-Cellular Fractions (RT-S9) From Rainbow Trout And Extrapolation To In Vivo Intrinsic Clearance Series On Testing And Assessmen. 2018, No. 280, 47. URL: https://one.oecd.org/document/ENV/JM/MONO(2018)12/en/pdf

(134) Galluzzi, L.; Aaronson, S. A.; Abrams, J.; Alnemri, E. S.; Andrews, D. W.; Baehrecke, E. H.; Bazan, N. G.; Blagosklonny, M. V.; Blomgren, K.; Borner, C.; Bredesen, D. E.; Brenner, C.; Castedo, M.; Cidlowski, J. A.; Ciechanover, A.; Cohen, G. M.; De Laurenzi, V.; De Maria, R.; Deshmukh, M.; Dynlacht, B. D.; El-Deiry, W. S.; Flavell, R. A.; Fulda, S.; Garrido, C.; Golstein, P.; Gougeon, M.-L.; Green, D. R.; Gronemeyer, H.; Hajnóczky, G.; Hardwick, J. M.; Hengartner, M. O.; Ichijo, H.; Jäättelä, M.; Kepp, O.; Kimchi, A.; Klionsky, D. J.; Knight, R. A.; Kornbluth, S.; Kumar, S.; Levine, B.; Lipton, S. A.; Lugli, E.; Madeo, F.; Malorni, W.; Marine, J.-C.; Martin, S. J.; Medema, J. P.; Mehlen, P.; Melino, G.; Moll, U. M.; Morselli, E.; Nagata, S.; Nicholson, D. W.; Nicotera, P.; Nuñez, G.; Oren, M.; Penninger, J.; Pervaiz, S.; Peter, M. E.; Piacentini, M.; Prehn, J. H. M.; Puthalakath, H.; Rabinovich, G. A.; Rizzuto, R.; Rodrigues, C. M. P.; Rubinsztein, D. C.; Rudel, T.; Scorrano, L.; Simon, H.-U.; Steller, H.; Tschopp, J.; Tsujimoto, Y.; Vandenabeele, P.; Vitale, I.; Vousden, K. H.; Youle, R. J.; Yuan, J.; Zhivotovsky, B.; Kroemer, G. Guidelines for the Use and Interpretation of Assays for Monitoring Cell Death in Higher Eukaryotes. Cell Death Differ 2009, 16 (8), 1093–1107. 10.1038/cdd.2009.44.

(135) Liu, Y.; Jiao, J.; Zhang, C.; Lou, J. A Simplified Method to Determine Five Cytochrome P450 Probe Drugs by HPLC in a Single Run. Biol Pharm Bull 2009, 32 (4), 717–720. 10.1248/bpb.32.717.

(136) Donato, M. T.; Gómez-Lechón, M. J. Fluorescence-Based Screening of Cytochrome P450 Activities in Intact Cells. In Methods in Molecular Biology; 2013; Vol. 987, pp 135–148. 10.1007/978-1-62703-321-3_12.

(137) Sheu, C. J.; Lee, J. K.; Rodriguez, I.; Randolph, S. C. The Use of Uninduced Rat Liver S-9 to Supplement BALB/3T3 Cells in the in Vitro Transformation Assay. Drug Chem Toxicol 1991. DOI: 10.3109/01480549109017871

