## Supplementary manuscript for "On the utilisation and characterisation of external biotransformation systems in *in vitro* toxicology: a critical review of the scientific literature with guidance recommendations"

Special inquiries

 (regulatory inquiries)

 (technical inquiries)

ORCID IDs

SLM: https://orcid.org/0000-0001-8192-9134

MSÅ: https://orcid.org/0009-0005-0061-2722

IR: https://orcid.org/0000-0002-2968-4659

FPV: https://orcid.org/0000-0002-7159-9781

AS: https://orcid.org/0000-0002-9142-3345

HH: https://orcid.org/0000-0001-5776-5619

MJ: https://orcid.org/0000-0002-4858-0118

KH: https://orcid.org/0000-0001-6320-8093

1. List of additional supplementary information material (SI) besides the supplementary manuscript (SM). SIs have been uploaded to Figshare and are available under the link: (https://doi.org/10.6084/m9.figshare.28504826.v1).

**Table S1**: Description of all provided supplementary elements.

| Name | Content |
|---|---|
| SM | Supplementary manuscript (this file) |
| SI1 | Raw search outputs from search string 1 (“endocrine”) |
| SI2 | Raw search outputs from search string 2 (“mutagen”) |
| SI3 | Details of screening and selection processes conducted for “BTS1/endocrine” |
| SI4 | Details of screening and selection processes conducted for “BTS2/mutagen” |
| SI5 | Composition of the database “BTS3/historical” |
| SI6 | Queryable, wide-format, total database including extracted data item measures per subdomain for every publication record. |
| SI7 | Summarised scores from DEERS assessment. |
| SI8 | Raw data on meta-regression analyses. |
| SI9 | Machine-readable simplifications of coded data items. |
| SI10 | Additional interactive relational network graphs from *Apriori* analyses. |
| SI11 | Raw data on follow-up, confirmatory analyses. |

### 2. Supplementary material & methods

As described in the main article, the critical review of BTS reporting standards and methodology follows a systematic approach, as utilised in systematic reviews (SR), systematic evidence maps (SEM), and scoping reviews (ScR). To improve brevity and readability, the main article only highlights the most important methodological details. Here, within the supplementary manuscript, all methods conducted are exemplified in a PRISMA-aligned manner (Page et al., 2021a, 2021b). Additionally, the SM contains all conducted analyses and results, as the main article's narrative only highlights the most important aspects.

To meet the requirements of this workstream, the structural framework of the critical review was adapted from canonical SR, SEM, or ScR outcomes (Khalil and Tricco, 2022; Munn et al., 2018). Alternations to canonical systematic review frameworks are further discussed in section 2.15.

#### 2.1 Protocol and registration

Following the PRISMA guidelines, a protocol was conceptualised, published, and shared amongst peers. The critical review protocol has been prepared according to the PRISMA-P guidelines (preferred reporting items for systemic review and meta-analysis protocols (Moher et al., 2015; Shamseer et al., 2015)) and the European Food Safety Authority's (EFSA) guidelines for systematic reviews (EFSA, 2010). The protocol was uploaded to Figshare (https://doi.org/10.6084/m9.figshare.21494616.v2) on November 3rd, 2022, published on November 21st, 2022, and updated on August 17th, 2023.

#### 2.2 Description of the project team and areas of expertise related to the conducted review

The authors have accumulated expertise in *in vitro* toxicology, xenobiotic metabolism, regulatory approaches, and ample BTS experience. Short individual summaries are given alphabetically, as follows.

AS – *in vitro* toxicology, development of biotechnological metabolisation systems ewoS9R, genotoxicology

FPV – *in vitro* toxicology

HH – evolutionary ecology and environmental toxicology, with a focus on bioanalytical toxicology, mechanistic-specific toxicity, mixture toxicity, alternative animal testing methods, effect-directed analysis, and omics – beyond others

IR – *in vitro* toxicology, xenobiotic metabolism, endocrine disruption, application of (biotechnological) BTS

KH – *in vitro* toxicology, endocrine disruption, bioassays development, and implementation for hazard and risk assessment

MJ – regulatory toxicology, (environmental) hazard and risk assessment, endocrine disruption, reproductive & developmental toxicology, metabolism and metabolic disruption, *in silico* tool development QSAR, Integrated Approaches for Testing and Assessment (IATA), validation and test guideline development

MSA - Multivariate statistics in R

SLM – *in vitro* toxicology and method development; lead in the practical implementation of external BTS into *in vitro* testing batteries, as conducted within the Horizon 2020 ERGO project (European Commission, ID: 825753)

### 2.3 Eligibility criteria

The studies chosen were limited to those in English and which had undergone peer review, given that this process should guarantee methodological rigour. Officially recognised international guidance and communication is referenced and not categorised as “grey literature”. Grey literature types, such as unreviewed internal reports, were excluded from consideration. Both meta-analyses and review articles were removed from the dataset. There were no limitations based on the studies' geographic origin or publication date. In principle, every peer-reviewed publication employing an exogenous BTS in an *in vitro* setup was eligible for our analysis within the limits of the defined search strategy (see section 2.6, in SM). The following BTS parameters were defined as of utmost importance via "piloting" (EFSA, 2010): protein concentration, origin, manufacturing, and spatiotemporal application, as they were shown to impact the results of the biotransformation procedure. It was postulated that eligible studies should contain information about the latter to ensure scientific and methodological rigour.

Primarily, we focused on rat-derived BTS, given their overall abundance, representation, and commercial availability in the field, but not exclusively. Therefore, we added BTS derived from humans, mice, fish, and others for comparison. Both S9 (incorporating both cytosolic and microsomal fractions) and microsome-only type BTS were included in the study, even though the search string prioritises S9 BTS due to search size delimitations. Besides applied BTS concentrations and details on origin, we focused on the employed cofactors during the BTS reaction. Depending on the employed cofactors, the investigator can interrogate phase 1 or 2 biotransformation reactions as appropriate.

### 2.4 Statement on PICO/PECO(TS) criteria

As our investigation is not centred around an intervention/exposure-to-outcome framework but on scientific and methodological rigour in BTS application and their resulting exploratory interconnections, we will abstain from assigning PICO(S) ("population, intervention, comparator, outcomes, study setup") or PECOTS ("population, exposure, comparator, outcomes, target condition, study design" – described in the Conduct of Systematic Reviews in Toxicology and Environmental Health Research (COSTER, (Whaley et al., 2020)) criteria. In practice (Table S3), we can assign a PO statement ("population + outcome" (James et al., 2016)) to our closed-framed, confirmatory hypotheses (aims A and B, within the main article). However, this framework does not rely on the actual defined data items but our interpretation thereof, with the population being coded and extracted data item measures and the outcome final scoring assessment via the DEERS framework (see appendix 6.1) or meta-analysis results of quantitative data item measures (see section 2.10 below).

PRISMA guidelines have been devised for clinical studies, which are not necessarily appropriate for toxicological evaluations. In theory, data item domains defined in the coding book (see section 2.9) can be assigned to PICO/PECOTS, and we conducted this as an exercise in Table S3. For relevance (Table S3), PICOS/PECOTS criteria were recorded for every screened and credited publication (EFSA, 2010). However, they are not at the core of the meta-analysis or discussion. Our approach to handling PICOS/PECOTS criteria is discussed further below in sections 2.9 ("data items and coding") and 2.12 ("effect measures and synthesis methods"). In conclusion, we deem this assignment an artificial construct and prefer to use our framework DEERS (Data Extraction, Evaluation, and Reliability Schema; see appendix 6.1 and section 2.11).

2.5 Information sources

Standard scientific literature repositories, Pubmed, Web of Science, and Scopus, were used to construct a database for information collection via Boolean operator searches (see section 2.6).

Additionally, a subset of highly relevant publications previously known to the reviewers (n = 73, see SI5 and reference list "BTS3" in section 5.4) was initially screened for piloting. The subset was used to refine the search strategy. Four reviewers (MJ, SLM, AS, KH) practically refined the search strategy and conducted exploratory searches. Further, the subset was checked for cited cross-references according to the data selection processes described below. Cross-referenced publications not incorporated by the below-described automated collection processes were manually added to the literature database and forwarded to analysis and synthesis (total manually added a subset of n = 121 publications; see results section 3.1 in the main article). This database is referred to as "historical/BTS3" in the following.

2.6 Search strategy

The Boolean search operators are named in the main article, section 2.3.

Noteworthy, the Web of Science repository does not support prefixed operators, such as "*estro*". Therefore, in this specific case, the search was conducted once with the operator "estro*" and once with "oestro*" to account for different notations. However, no differences in output were obtained. The databases and derived bibliographies from the search strings are referred to as "endocrine/BTS1" or "mutagen/BTS2" within the main article, the SM, and all the supplementary information material files (SI). SLM, MJ, KH, and AS discussed, conceptualised, and agreed upon the search strategies. No limits or filters were applied to the search strategy beyond the eligibility criteria (see section 2.3 above). The search operators were optimised towards mutagenic, genotoxic, and endocrine-disruption endpoints recorded in *in vitro* systems jointly applied with BTS to concur with the objectives (see section 1.2, main article) and keep search outputs within a manageable margin. Eukaryotic, continuous cellular *in vitro* systems were favoured by adding the operator "cytotox*" and S9 BTS by adding the operator "S9" to the respective search terms.

SLM conducted searches on March 7, 2022, and reiterated them on March 29, 2022, with identical results. During the main article's preparation process, SLM repeated the searches on January 24, 2023 (see results section 3.1, main article). The raw search outputs of BTS1 and BTS2 are given in SI1 and SI2, respectively.

2.7 Study selection process

The inquiry outputs from the above-stated literature repositories were extracted as readable bibliography formats (RIS), subjected to automatic duplicate removal in Mendeley Desktop (version 1.19.8, Glyph & Cog, LLC), and imported into the Sysrev GUI (https://sysrev.com/) via PubMed identifiers (PMID). All inquiry outputs were collected into a single database within the application. The Sysrev tool was utilised for article selection by four reviewers (AS, FPV, IR, SLM) in a randomised manner. Titles and abstracts were screened and selected according to the stated eligibility criteria (see section 2.3 above). All four reviewers worked independently. The Sysrev tool randomises the sequence of articles for every reviewer. Article inclusion was handled as follows. All records reaching a 100% or 75% inclusion rate from all reviewers were instantly accepted, and all references reaching 25% or 0% were immediately excluded. For articles matching a 50% inclusion rate, titles and abstracts were scrutinised more carefully, the full text was retrieved and inspected, and a consensus was reached via final discussion (see SI4 and SI5 for more details).

### 2.8 Data collection process

After selection, the adjusted bibliography was exported as EndNote XML files and imported into Mendeley Desktop. Duplicates were checked and removed manually. Afterwards, full-text PDFs of every article were retrieved. The data collection was hand-curated in Mendeley Desktop.

Data curation, extraction, and evaluation were conducted via the devised DEERS protocol (Data Extraction, Evaluation, and Reliability Schema; see appendix 6.1 and section 2.11 below). DEERS output of every article was categorised, sorted, and summarised in a Microsoft Excel spreadsheet as prepared as an extraction form for every reviewer (SI6). As pointed out in the DEERS protocol, a unique ID number was assigned to every article, and specific data items were collected for all primary data item domains and subdomains.

For every article, two reviewers independently conducted data extraction in detail (SLM, approximately 70% of the final bibliography; IR, about 30%). The entire extracted dataset for every data item was then double-checked and curated independently by four reviewers (AS, FPV, IR, SLM). Disagreement was resolved via discussion in case of discrepancies within reviewers' collection processes. Section 2.9 ("data items and coding") and section 2.6 of the main article present methodological information extracted for three primary and 24 subdomains.

### 2.9 Data items and coding

Coding for data item domains is described in the main article, section 2.6, and summarised in Table 1 of the main article. Four reviewers (MJ, KH, AS, SLM) discussed, conceptualised, and agreed upon the coding approach. We coded for 24 subdomains allocated to three primary domains (BTS characterisation, reaction components, and experimental setup). The subdomains were divided into “data items of relevance” (“non-critical” in Tab. S2) and “data items of reliability” (“critical” in Tab. S2), with the second category feeding into the methodological reliability assessment (outcome A) via DEERS (appendix 6.1). The reasoning for data item domain categorisation is outlined in the main article, section 2.6. A hypothetical example of extracted data measures in accordance with the coding book is given in Table S2.

The measures of the extracted data items are qualitative (descriptive, categorical, dichotomous) or quantitative (numerical, continuous). Total BTS protein concentration is illustrated as mass per volume. Primary and secondary cofactors are depicted as molar concentrations. Further, the incubation period is recorded in minutes or hours, and the temperature is degrees Celsius. All other data items were documented qualitatively (e.g., BTS induction via Aroclor or BNF/PB-spiked diet). Table S2 gives examples of extracted data item measures for every domain. Critical data item domains were assessed for reliability, whereas non-critical data items were only evaluated for relevance. As for some articles, multiple measures of data items were retrieved, specific annotations were made in DEERS, and explanations were indicated within the extraction forms. For a more detailed resolution and precise data handling of item domains and their respective measures, please consult the "explanations" sheet in SI6.

**Table S2**: Examples of data item measures. Non-critical domains are only assessed in terms of relevance, and critical domains are evaluated in terms of reliability.

| **(Sub-)Domain** | **Measure** | **Assessment** | **Example** |
|---|---|---|---|
| Author | Qualitative | Non-critical | *Allaben et al.* |
| Publication year | Numerical | Non-critical | *1979* |
| Journal | Qualitative | Non-critical | *Cancer Letters* |
| Test system | Qualitative | Non-critical | *CHO cells* |
| Endpoint | Qualitative | Non-critical | *Genotoxicity* |

| | | | |
|---|---|---|---|
| Methodology | Qualitative | Non-critical | *ER and AR Calux assays* |
| External BTS type | Qualitative | Non-critical | *S9* |
| BTS origin | Qualitative | Critical | *Self-made* |
| Species | Qualitative | Critical | *Rat* |
| Strain | Qualitative | Critical | *Sprague-Dawley* |
| BTS pooling | Qualitative | Critical | *female, male, nd* |
| Husbandry | Qualitative | Critical | *Details given* |
| BTS induction | Qualitative | Critical | *Aroclor* |
| BTS protein concentration | Numerical | Critical | *1 mg/mL* |
| Buffer system | Qualitative | Critical | *Culture medium* |
| BTS dilution | Percental, fractal | Critical | *final* |
| Primary cofactors | Numerical | Critical | *0.74 mM NADPH* |
| Other cofactors | Numerical | Critical | *5 mM G6P* |
| Exposure | Numerical, qualitative | Non-critical | *5 mM benzo[a]pyrene* |
| Solvent | Qualitative | Critical | *DMSO* |
| BTS incubation period | Numerical | Critical | *40 min* |
| BTS incubation temperature | Numerical | Critical | *37°C* |
| Post BTS procedure | Qualitative | Non-critical | *quenched by ACN, extraction, CA* |
| BTS-related controls | Qualitative | Critical | *w/o BTS* |

As the critical review assesses reporting bias and methodological reporting rigour, data items could not be retrieved for every domain from any article. In the scenario that specific data items were not available, they were categorised as "nd" ("not defined") in the DEERS protocol and data extraction sheet (SI6). Further, if the true nature of a data item was concealed by unclear formulation or lacking definition, e.g., non-determined final concentration due to omitted dilution factor indication, data items were defined as "nc" ("not clear"). Finally, if data items were evidentially not applied or redundant in a specific methodological scenario, e.g., BTS induction regimes in human-derived microsomes, they were defined as "na" ("not applicable/assessable"). Section 2.12 iterates how missing data items are dealt with in the reliability assessment.

As mentioned previously, PICOS/PECOTS criteria could not be the central basis of the synthesis, but these criteria for the data collection facilitated a stringent characterisation of relevance. The subdomains can be assigned to PICOS/PECOTS as shown in Table S3. PICOS/PECOTS criteria are named and defined here for stringency reasons to adhere to the original PRISMA and COSTER guidelines. However, in our specific case, we did not regard PICOS/PECOTS as directly applicable to the reviewed topic and focused on the DEERS evaluation systems (see appendix 6.1 and section 2.11 below).

**Table S3:** Data item domains according to PICOS/PECOTS criteria.

| **PICOS/PECOTS criterium** | **Respective subdomain** |
|---|---|
| Population | Species, strain, pooling (and sex) |
| Intervention/Exposure | Exposure |
| Comparator | BTS-related (and study-specific) controls |

| | |
|---|---|
| Outcomes | Not assessed, heterogeneous |
| Target conditions | Endpoint |
| Study design | BTS experimental setup (incubation period, incubation temperature, and BTS-related controls). |

2.10 Outcomes and prioritisation

Primary and secondary study outcomes are defined as closed-framed hypotheses, formulated as objectives A and B (main article, section 1.2). Further, outcomes were determined for all data item domains named in Table 1, main article, given that they were logically feasible, and enough data could be extracted for overall analysis. First, the primary outcome is an assessment of BTS's methodological reporting standards and scientific rigour. We employed a grading scheme (DEERS, see appendix 6.1 and section 2.11 below) to depict the quality status of methodological reporting within the articles of the derived bibliography (aim to outcome A). This assessment was conducted for the entire bibliographic database by definition of the eligibility criteria. Second, quantitative data items (see section 2.9 above, "data items and coding") were plotted as time-response (concentration) or concentration-concentration-related functions of BTS incubation time, BTS protein concentration, and BTS-related cofactor concentration (aim to outcome B). We hypothesise that the quantitative data should relate to linear, exponential, or asymptotic behaviour. For the tertiary outcome (aim to outcome C), extracted data item measures of every domain were simplified in a further coding step to facilitate downstream analysis of categorical data and potential for machine reading (Table S4 and SI9). For the tertiary outcome, exploratory analyses were conducted in a data mapping manner, facilitating the development of a subsequent regulatory guidance framework (aim to outcome D). Finally, using descriptive statistics, all qualitative data items were illustrated as tertiary outcomes.

Secondary and tertiary outcomes were only retrieved where possible and feasible. Due to an expected rate of incomplete reports, it was assumed that recovering data for all data items would not be possible. The final report states the seminal population of every specific outcome. For more details on the respective outcomes, consult section 2.12, "synthesis methods," and the results sections of the main article and this document.

2.11 Risk of bias & DEERS

Compared to standard systematic reviews, we are not investigating experimental, clinical, or trial outcomes, but the reporting standards of methodological data items. Per se, the critical review focuses on reporting bias as an outcome. Hence, the investigated outcomes do not have the character of measurement, including an error, but are fixed dichotomous characteristics (reported vs. not reported). Further, we are not analysing and synthesising the studies' observations and conclusions. The reported concentrations, temperature, or time points were used for further meta-analyses of numerical and continuous outcomes, but these data points have no intrinsic error. An overall meta-analysis of all extracted data item measures was impossible, given that the data items are incompletely reported. Instead, a separate population/dataset was curated for every evaluated data item/outcome.

Further, an assessment of publication and selective non-reporting bias was omitted. I.e., methodological reporting rigour might only be positively affected by publication bias (e.g., an article heavily below the methodological quality standards of the reviewed literature would never be published; its potential overall low score would further negatively impact the aggregated quality of the database). Thus, we deemed assessing the risk of bias unfit in this case. A discussion of the potentially biased assumptions and frameworks of this critical review at hand is given in section 4.1 below. Instead, we orientated towards the SWiM guideline ("Synthesis without meta-analysis in systemic reviews" (Campbell et al., 2020)). We employed methods to examine heterogeneity and assess the certainty of evidence among selected studies. Hence, a reliability assessment in the form of a critical appraisal was conducted instead of a risk of bias analysis. Therefore, we devised a data extraction, evaluation, and reliability schema (DEERS, see appendix 6.1) in line with the reliability schemata ToxRTool (Schneider et al., 2009), CRED ("Criteria for reporting and evaluating ecotoxicity data" (Moermond et al., 2016)), and the Klimisch method (Klimisch et al., 1997).

Contrary to the above methods (Klimisch et al., 1997), we abstained from categorising the studies regarding reliability quality. We want to depict the reporting issues retrospectively but not judge individual publications. OECD and ISO guideline methodology relevant to metabolising systems were utilised where appropriate.

The weighing of criteria was neglected in this review, given that using weighted scoring might be deemed scientifically controversial (EFSA, 2010). Instead, all scores were summed up equipotently to the final quality assessment score. Additionally, we outline how current BTS approaches can improve the characterisation of BTS (see the discussion part in the main article).

2.12 Effect measures and synthesis methods

The following section provides effect measures and synthesis methods for every assessed outcome regarding their prioritisation hierarchy, defined in section 2.10, "outcomes and prioritisation,", and within the main article.

**Effect measures and synthesis methods for outcome A (“scoring”):** The primary outcome is an assessment of BTS methodological reporting standards and scientific rigour (outcome to aim A), as elaborated via the DEERS protocol. Extracted measures for every data item were scored binary, reported vs. non-reported, resulting in a net score of one per reported data item subdomain. The measure of the data item domain was of a qualitative or quantitative character, as defined in Table S2 and more detailed in SI6. As described above, only critical data item domains were scored for the reliability assessment. Out of the 15 critical data item domains, 16 scoring categories were derived, given that the "BTS pooling" domain was divided into two separate scoring domains (pooling sex and pooling number of individuals).

A reported data item received a positive binary score of 1, whereas a non-reported item received a neutral score of 0. Non-reported or concealed data items were marked as "nd" (“not defined”) or "nc" (“not clear”) within the data extraction form. Data items evidentially not applied or redundant in a specific BTS scenario were marked as "na" (“not applicable/assessable”) and received a positive score. For more details on missing data handling, see also section 2.9 above. Scores were registered manually in the DEERS protocol and transferred to the data extraction sheet (SI6) in Microsoft Excel. All extracted data was summarised in a wide, queryable table format database (SI6). The scores were summed according to their primary domains (Table 1 in the main article) and forwarded for descriptive analyses in R (R CoreTeam, 2023). Total and relative scores for each domain were visualised using the ggplot2 package (Wickham, 2016). Overall scores were plotted as hierarchically stacked bar plots of primary data item domains or as scoring population boxplots per primary data item domain. Per se, only a 100% score was determined to be fully reproducible and assessable in methodological terms. However, an arbitrary 80% overall achieved score threshold was defined as a lower boundary of potential rectification.

For the scoring results derived from outcome A, statistical interference of inquired sub-populations was conducted via non-parametric Kruskal-Wallis tests. In the scenario where more than two subgroups were analysed, the latter was followed by a two-sided Dunn's test for multiple comparisons (alpha level = 0.05). Statistical analyses were conducted in GraphPad Prism 8. Statistical interference of scoring sub-populations was conducted for sub-datasets, as patterns emerged from the exploratory analysis (aim C and tertiary study outcomes).

**Effect measures and synthesis methods for outcome B ("meta-regression"):** Meta-analyses of quantitative, numerical data item domain measures were conducted as a secondary outcome (outcome to aim B). We intended to identify or deconstruct concentration-response or temporal patterns of BTS applications. Hence, it was hypothesised that specific BTS parameters should constitute approximate linear or exponential relationships, such as for the BTS incubation period, BTS protein concentration, and BTS primary cofactor concentrations.

Incubation period vs. BTS protein concentrations, BTS incubation period vs. primary cofactor concentration, and BTS protein concentration vs. primary cofactor concentrations were plotted and analysed regarding their mathematical and statistical relation (simple linear regression and Pearson's correlation analyses). Respective quantitative data item measures, BTS concentrations in mg/mL, cofactor concentrations in mM, and time in minutes were log-transformed to align with parametric test assumptions. Compliance with parametric assumptions was tested and inspected: (semi-) normality of residuals was confirmed via the Shapiro-Wilk test and visually via Normal Q-Q plot of residuals; homoscedasticity of residuals was assessed via Levene's test and visually via Actual vs. Fitted residual plots. Simple linear regressions were fitted to the subgroups, and adjusted $R^2$ values were derived as goodness-of-fit (model accuracy) measures. Further, Pearson's parametric correlation was utilised to determine the coefficient of correlation and the coefficient of determination of the above-stated variables. Additionally, paired comparisons were conducted for measures originating from identical publication records. Finally, to account for potential toxicokinetic effects, we fitted experimental setups which do not apply BTS regeneration systems.

During data collection, all data item measures were converted as follows:

All time-related measures were expressed in minutes (min), and all concentration-related measures were defined in molarity (mM). If a dilution factor was applied, the final BTS reaction components and concentrations were calculated and used for the analyses. A subgroup was hand-curated for every meta-analysis due to the reports' incompleteness of data item measures within selected studies. Also, for some articles, multiple measures were extracted for single data item subdomains, given that some studies applied different setups or ranges of periods and BTS reagent concentrations. If different methodological setups were involved, all measures were extracted separately, meaning multiple measure parameters for a single domain. In the scenario in which a range was applied, the mean value was calculated and noted as the final measure (see SI8 for details). The seminal populations of these subgroups are reported within the results section (this manuscript and main article). Computation, statistical analyses, and graphical plotting were conducted in GraphPad Prism 8 (GraphPad Software, La Jolla, USA).

**Effect measures and synthesis methods for outcome C ("mapping"):** As a tertiary study outcome (outcome to aim C), all qualitative data item domains were subjected to exploratory mapping analyses. All qualitative data item measures were simplified in a second coding approach (Table S4), making them machine-readable and accessible for further analysis. We used descriptive and summary statistics to depict all extracted data (histograms, Euler diagrams, Upset plots). Coded data items were subjected to multiple factor analysis (multiple correspondence analysis, “MCA”) to identify associative patterns within the data. *Apriori* algorithms were utilised for data mining approaches, which helped identify qualitative data association rules and build relational networks.

**Descriptive and summary statistics**: All data preparation and visualisation for descriptive and summary statistics were conducted using R Studio (Posit team, 2023)and R software version 4.1.2 (R Core Team, 2021). Bar plots, stacked bar plots, and stepped line graphs to visualise absolute and relative scores per primary domain were produced using ggplot2 (Wickham, 2016). Euler diagrams for individual subdomains were created using the eulerr package (Larsson, 2022). Euler diagrams illustrate relation frequencies (simultaneous occurrence (intersection) of qualitative data item measures) with circle size corresponding to the categorical sub-population. Upset plots were generated for each subdomain with the UpSetR package (Gehlenborg, 2019), visualising the main bar plot, the set size bar plot, and the matrix plot. Upset plots depict interaction frequencies, i.e., an intersection matrix with the rows corresponding to sets of categories (absolute observations) and the columns corresponding to the intersections between these sets of categories (relative observations). Data underwent a numerical transformation for histograms, with each observation allocated proportional scores for applicable domains. Specifically, an observation linked to three qualitative data item measures was evenly distributed, each receiving a proportion of 0.333. Stacked histograms for each subdomain were produced with the bin size set to five years using ggplot2 (Wickham, 2016). The histograms illustrate distribution frequencies over the investigated time period (years 1973 to 2022).

**Multiple Correspondence Analysis (MCA)** was employed to visualise the associations among categorical interdependent and supplementary variables using the FactoMineR package (Lê et al., 2008) in R Studio (Posit team, 2023) and R software version 4.1.2 (R Core Team, 2021), according to recommendations described in (Husson et al., 2017).

MCA, an extension of Correspondence Analysis (CA), analyses relationship patterns within categorical variables (Abdi and Valentin, 2007). Like Principal Component Analysis (PCA), the first dimension in an MCA captures the data’s most significant source of variability. It represents the primary pattern or relationship between categories that explain the most significant proportion of the total inertia. The second dimension is orthogonal (uncorrelated) to the first dimension and captures the next most significant source of variability. Each subsequent dimension continues to capture independent patterns in decreasing order of significance.

We conducted the following data processing approach to balance preserving meaningful information, reducing the dimensionality of the dataset, and preventing rare categories from exerting a too strong influence on the MCA: for every variable (data item subdomain), a maximum of five qualitative categories (data item measures) was allowed. Therefore, very rare categories ($n < 5$ absolute observations) were either reallocated to other categories (if feasible) or deleted. Please consult the specific sheets in SI9 and Tab. S4, where detailed procedures are depicted per variable. The qualitative dataset was transformed into a binary information matrix for equal spacing of multi-categorical observations. I.e., for a hypothetical variable X, with the categories A, B, and C, an observation (publication record) containing B and C would be coded as 0-1-1. The final qualitative dataset and the binary matrix are given in SI9, first sheet. Computations were conducted based on the binary information matrix. For the MCA, the active variables (interdependent) included “research field” (previous subdomain “journal”, redefined in simplification step, see Tab. S4), “test system”, “endpoint”, “external BTS” (type), “producer” (BTS origin), “species”, “species system origin” (sub-category, redefined in simplification step, see Tab. S4), “strain”, “BTS pooling sex”, “husbandry details", "BTS induction", "buffer system", "cofactor class" (primary cofactor, redefined in simplification step, see Tab. S4), "solvent", and "BTS-related controls". The supplementary variables (independent) "publication year" (grouped in five-year intervals), "publication group" (extraction datasets BTS1 to BTS3), and "total scores" were defined as qualitative supplementary variables and did not influence the analysis mathematically but were only superimposed on the graphical illustrations.

Plotting in ggplot2 (Wickham, 2016) was restricted to levels with more than five relative observations, with a few logical exceptions. Note that single observations (every recorded literature article) are often multi-categorical per variable, e.g., a publication record can be defined by both categories "Tox" and "Env" for the variable "research field". For the MCA computation, multi-categorical dependencies were dissolved by transforming the dataset into a binary information matrix (SI9, first sheet). Thus, in binary code, absolute observations can be summarised per category and variable by the additional dimensionality of the matrix. However, when plotting the MCA, the categories (qualitative data item measures) are superimposed onto the binary calculations per variable. Hence, an observation defined by both "Tox" and "Env" cannot be divided but must remain a single entity in graphical terms ("Tox" + "Env"). These multi-categorical levels are defined as relative observations.

MCA plots were generated for all active and supplementary variables. As the qualitative measures are superimposed on the binary computation, all active variables are illustrated interdependently, whereas all supplementary variables are independent. Data ellipses were added to represent normal probability contours at a 0.9 confidence level, and centroids were plotted at the mean individual coordinates for dimensions 1 and 2. Data ellipses could only be computed for relative observations with $n > 5$.

**Data association and relational networks**: Association rule mining was employed to uncover relational networks among qualitative data item domains in the simplified coding dataset (Table S4 and SI9), using the *Apriori* algorithm in the arules package (Hahsler et al., 2023) in R Studio (Posit team, 2023) and R software version 4.1.2 (R Core Team, 2021). This process identified frequent data item measure sets and generated association rules under specific parameters: a support threshold of 10%, which indicates the minimum frequency of a data item set in connections for consideration, and a confidence threshold of 80%, representing the minimum confidence level for an association rule to be significant.

The "support" parameter indicates the frequency or the proportion of relations in a dataset that contain a specific set of data item measures. It is calculated as the number of relations containing the itemset divided by the total number of relations in the dataset. The support of an itemset indicates how frequently it appears in the dataset. Higher support values suggest that the itemset is more commonly occurring.

The "lift" parameter indicates the strength of association between two data item measures within a rule (node association direction). It is calculated as the ratio of the observed support of the itemset containing both data item measures to the expected support of the itemset if the data item measures were independent. Lift values greater than 1 suggest a positive association, meaning that the measures tend to occur together more often than expected by chance. Lift values less than 1 indicate a negative association, implying that the measures occur together less often than expected by chance. A lift value of 1 suggests independence.

The maximum number of item association rules was separately set to 50, 100, 200, 500, 1000, 2000, and 5000. Publication score was categorised as pass (score $\geq$ 13) or fail (score < 13). The visualisation of these associations was facilitated using the arulesviz package (Hashler, 2023) with htmlwidget (https://www.htmlwidgets.org/) as the engine to generate interactive plots.

**Confirmatory analyses**: For qualitative subdomain measures associated with methodological robustness markers within the MCA and *Apriori* analyses (see Tab. S9), subsequent confirmatory investigations (Hair et al., 2019) were conducted to evaluate the effect sizes in different scoring subpopulations. Specifically, subdomains that exhibited robustness markers, either positively or negatively, in both MCA and *Apriori* frameworks were earmarked for in-depth analysis. The trajectory of these analyses, whether positive or negative, was determined based on the outcomes of MCA and *Apriori* assessments. Detailed information on the curated scoring subpopulations related to the examined data item subdomain and data item measure pairs is provided in SI11.

Non-parametric, two-sided Mann-Whitney-U tests were employed to compare groups comprising two scoring subpopulations (alpha level = 0.05). In multiple comparisons, the Kruskal-Wallis test was utilised as the primary analytical tool, followed by Dunn's post-hoc test (alpha level = 0.05). All the tests mentioned were performed with GraphPad Prism 8.

**Table S4**: Simplification of coding book for machine-readable measures; for more details, see SI9, simplified code given in parenthesis.

| Data item domain | Qualitative outcome scoring categories |
|---|---|
| Year | Binned into 5-year intervals |
| Journal | Redefined to "field": biomedicine ("Biomed"), toxicology ("Tox"), environmental sciences ("Env"), nutritional sciences ("Nut"), general biosciences ("BioSci"), analytical chemistry ("AnChem"), veterinary sciences ("Vet")<br><br>For MCA computation, the categories "BioMed", "BioSci", and "Vet" were pooled to "oBioSci" (other bio sciences) |
| Test system | Simplified to: Prokaryotic ("bacteria"), eukaryotic (w/o) fungi ("eukaryotic"), fungi ("yeast"), BTS only ("bts only"), and other ("other")<br><br>For MCA computation, "other" (n=4) has been removed |
| Endpoint | Simplified to: Mutagenicity & Genotoxicity ("MutGen"), endocrine disruption ("EDC"), metabolites identification & characterisation ("Meta"), cytotoxicity ("Cyto"), xenobiotic metabolism ("XenMet"), neuronal & developmental toxicity ("NeuDev"), and other ("other")<br><br>For MCA computation, "NeuDev" was pooled with "other"; "Cyto" appeared overwhelmingly as cytotoxic control (n=82), e.g., MutGen + Cyto, EDC + Cyto, etc. -> in this case, "Cyto" was deleted; otherwise (IDs B5, B6, B33, B42, C15; n=5), "Cyto" was defined as "other" |

| Methodology | Omitted due to complexity |
|---|---|
| External BTS | As is: “S9”, “microsomes”, “cellular”<br><br>For MCA computation, “cellular” (n = 2) was deleted |
| BTS origin | Simplified to: internal producer (“in”), external producer (“ex”), not defined (“nd”)<br><br>For MCA illustration plotted as “Origin II: producer” |
| Species | Further simplifications to: “mammal”, “bird”, “fish”, “invertebrate”, and not defined (“nd”);<br><br>Further characterisation to “species system origin”: “in vitro” or “in vivo”<br><br>For MCA computation, hamster, pig, dog, bovine, etc., were pooled to “oMammal” (other mammals; all fish species were pooled; non-vertebrates (n = 2) were removed<br><br>For MCA illustration plotted as “Origin I: BTS” |
| Strain | Only evaluated for rat-derived BTS;<br><br>Simplified to: Sprague-Dawley, BALB/c (“SD”), Wistar (“Wi”), Fisher (“Fi”), Long-Evans (“LE”), and not defined (“nd”)<br><br>For MCA computation, "LE" and "Fi" were pooled with "other" |
| BTS pooling sex | Simplified to: female (“F”), male (“M”), and not defined (“nd”)<br><br>For MCA computation, “na” (n = 3) was removed |
| Husbandry details | Simplified to: reported details (“yes”), further details on biotransformation capacity of BTS system (“activity”), details given elsewhere (“other”), and not defined (“nd”) |
| BTS induction | Simplified to: Aroclor and other PCBs (“Aro”), beta-naphthoflavone (“BNF”), phenobarbital (“PB”), Methylcholanthrene (“MCA”), applicable/assessable (“na”), and not defined (“nd”)<br><br>For MCA computation, “Aro” and “MCA” were pooled as “PCB-PAH”; one "other" (ID C79) was redefined as "PCB-PAH"; "other" (n =4) was removed |
| Buffer system | Simplified to: generic phosphate buffer (“xPO4”), cell culture medium (“cult”), Tris-HCl |

| | |
|---|---|
| | (“Tris”), phosphate-buffered saline (“PBS”), “Hepes”, and not defined (“nd”)<br><br>For MCA computation, "PBS" (n=3) was integrated into "xPO4" |
| Primary cofactors | Simplified to “cofactor class”: Phase 1 system (“ph1”), Phase 2 system (“ph2”), not defined (“nd”) |
| Secondary cofactors | Redefined to ”cofactor regeneration system”: glucose-6-phosphate (“g6p”), isocitrate (“iso”), applicable/assessable (“na”), and not defined (“nd”)<br><br>Additional categorisation “defined dehydrogenase system”: “yes”, “no”, previously not applicable/assessable in a nested hierarchy (“na”), previously not defined in a nested hierarchy (“nd”) |
| Solvent | Simplified to: alcoholic solvents (”alc”), other organic solvents (“org”), “DMSO”, water-based solvents (“H2O”), and not defined (“nd”) |
| BTS incubation period | As is, not assessed qualitatively |
| BTS incubation temperature | As is, not assessed qualitatively |
| BTS-related controls | Simplified to: without BTS (“wob”), without cofactors (“woc”), inactivated BTS (“inab”) |
| Exposure | Omitted due to complexity and focus on BTS methodology |

#### 2.13 Meta-biases

Common meta-bias analysis frameworks, such as the Grading of Recommendations Assessment, Development, and Evaluation (GRADE), the Navigation Guides of the National Toxicology Program's (NTP) Office of Health Assessment and Translation (OHAT) and the Office of the Report on Carcinogens (ORoC), and the Integrated Risk Information System of the U.S. Environmental Protection Agency (EPA-IRIS), are primarily centred on PICOS/PECOTS criteria (Rooney et al., 2016) and are assessing if the design and conduct of studies compromise the credibility link between intervention/exposure and an (adverse) outcome. As such, these frameworks are not directly suited for our investigation, as we investigate the overall scientific and methodological rigour of externally added BTS applications (see section 2.9 above). Thus, we abstain from meta-bias analyses, as is the case for SEM (Wolffe et al., 2020, 2019) and ScR (Tricco et al., 2018) procedures.

#### 2.14 Confidence in cumulative evidence

Certainty of evidence assessment relies on frameworks such as GRADE (Morgan et al., 2019), grading and summarising risk of bias, inconsistency, indirectness, imprecision, publication bias, the magnitude of effect, and others, according to the used framework. As we can neither determine these categories due to the structure of our investigation nor integrate them within our DEERS reliability assessment scheme, the certainty of evidence assessment is omitted. Instead, extracted data are assessed and evaluated via DEERS in a critical appraisal format (see appendix 6.1).

### 2.15 Structural framework of the critical review

We avoid defining the critical review as an SR, SEM, or ScR. Due to its nature, the critical review cannot deliver canonical SR, SEM, or ScR outcomes (Munn et al., 2018; Khalil and Tricco, 2022). Unlike canonical SR, which focuses its synthesis on a measured adverse outcome or physiological condition, scientific and methodological rigour within the documented literature is central to our critical review. Therefore, an SR-type risk of bias and certainty of evidence assessment is not possible, as, e.g., discussed in (Rooney et al., 2016; Morgan et al., 2019). We designed a reliability assessment schema for critical appraisal to alleviate this shortcoming ("DEERS", see sections 2.11 and the appendix 6.1). The reliability assessment schema is conceptualised to guide us through the systematic critical evaluation of methodological rigour within a BTS context and provide data item measures for meta-analyses.

Likewise, we cannot wholly adhere to the exploratory frameworks of SEM (James et al., 2016; Wolffe et al., 2019) and ScR (Tricco et al., 2018), as we aim to conclude the study with guidance communication for efficient and accurate BTS reporting and substantiate our claims via the meta-analysis of methodology-related data item measures. Further, our investigation does not need to examine emerging evidence and clarify concepts in an ScR-manner (Armstrong et al., 2011; Munn et al., 2018). The issues with BTS reporting have been acknowledged for almost two decades (Coecke et al., 2006; Gouliarmou et al., 2018; Jacobs, 2013; Jacobs et al., 2008; OECD, 2018, 2008). Therefore, we pursue both closed-framed, confirmatory hypotheses (for aims A and B, see section 1.2 of the main article) and openly-framed, exploratory hypotheses (aim C) and, thereby, extend the utility of the synthesis within a scientific and regulatory context (aim D) (see Fig. S1 for framework).

In theory, SRs in a BTS context would be possible once all experimental BTS parameters were appropriately defined and enough studies fit a homogenous outcome. As long as BTS parameters remain a black box, it is impossible to derive interference for a specific outcome via meta-analysis (e.g., comparator = well-defined but varying BTS parameters, outcome = adverse outcome for a particular mode of action or biological level of complexity). Here, we intend to set the groundwork for such future studies. Therefore, we need to gather the evidence and statistically interfere with the results – from a methodological and experimental standpoint, not an outcome perspective.

The protocol and the critical review follow the PRISMA structure (Page et al., 2021b, 2021a), with additional implementations from COSTER (Whaley et al., 2020). For the exploratory parts, SEM guidance is applied (James et al., 2016; Wolffe et al., 2019, 2020). Further, additional alterations are needed when translating guidelines derived from a clinical trial background to an *in vitro* toxicology context (EFSA, 2010; NTP-OHAT, 2019; US EPA, 2018). Specific alterations are highlighted in the sections above.

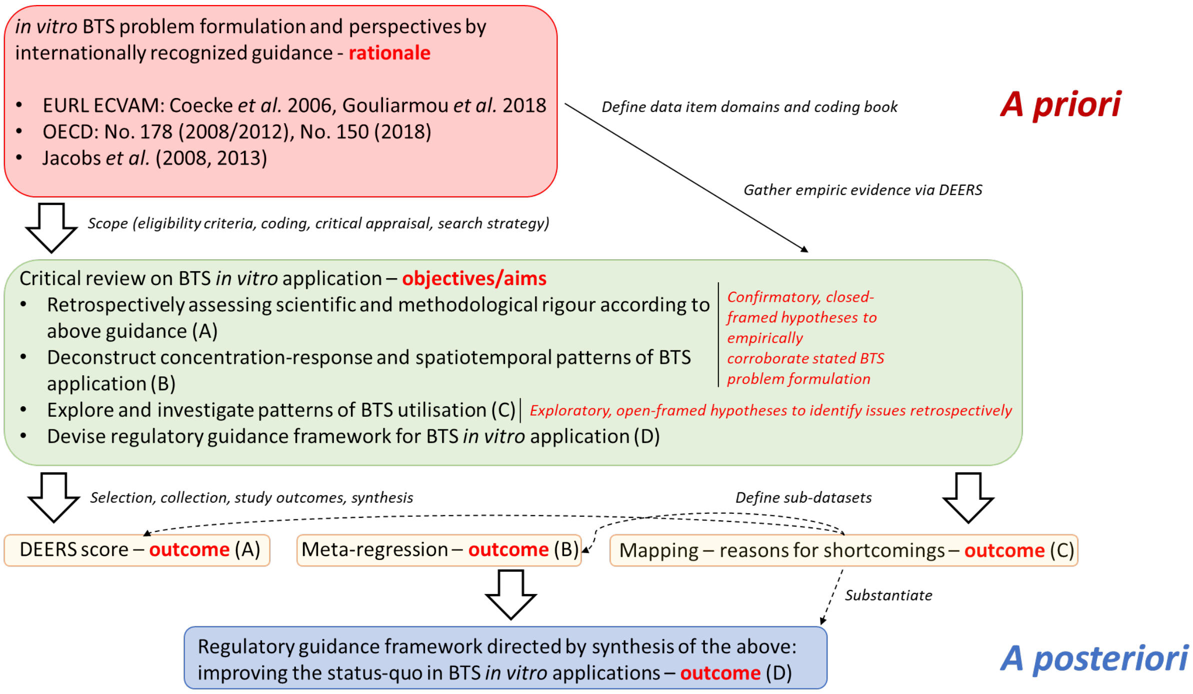

**Fig. S1:** Structural framework of the critical review regarding rationale, objectives, and study outcomes.

### 3. Supplementary results

#### 3.1 Selection of sources

The comprehensive search using defined Boolean operators yielded n = 221 records, with n = 76 from the "endocrine" query (“BTS1”) and n = 145 from the "mutagen" query (“BTS2”), as illustrated in Fig. 1 (main article) and detailed SI1 and SI2. These records were retrieved from PubMed, Scopus, and Web of Science. The initial search was conducted on March 7, 2022, and repeated on March 29, 2022, and January 24, 2023. The repeat searches confirmed the initial results, with a minor exception noted during the last search. Specifically, the "endocrine" search in the Web of Science database revealed two additional publications (Chelcea et al., 2022; Harding et al., 2023), as detailed in SI1. Despite meeting the eligibility criteria, these publications were not retrospectively added to the comprehensive bibliography. These reiterations underscore the manageable pace of publication in this context, suggesting our findings remain valid for publication and several years thereafter.

Concurrently, n = 121 records were sourced from the "historical" database and its cross-references (“BTS3”, see SI5). An initial automated duplicate removal was attempted using Mendeley Desktop but proved inefficient due to format discrepancies among different repositories. Consequently, manual duplicate removal was employed throughout the screening process (SI3 to SI5).

The retrieved records were processed through the Sysrev tool for title and abstract selection based on eligibility criteria outlined in section 2.3 here and section 2.1 of the main article (detailed in SI3, SI4, and SI5). Exclusions were made for various reasons, including non-originality (such as reviews by (Combes, 2012) or reports by (Galloway et al., 1994)), *in vivo* biotransformation studies (Liu et al., 2011), or absence of external BTS (Chang et al., 1988).

Post manual duplicate removal, n = 132 articles were earmarked for retrieval. Four were subsequently excluded for not meeting eligibility criteria: one article in Chinese (Zhang et al., 2016), a book chapter (Felton et al., 1984), and two articles lacking available full-text resources (Ampy and Asseffa, 1988; Siegers et al., 1987), leaving n = 128 articles for data extraction. From the n = 121 records identified in the "historical" database, 20 duplicates already present in the "endocrine" or "mutagen" searches and selection processes were removed, resulting in n = 101 articles from this source. In total, n = 229 records were advanced to data extraction, meta-analyses, and reliability assessment procedures.

Detailed study characteristics and citations for each bibliographical database are provided in SI3 to SI6, and a separate bibliographical database reference list is available in the SM, section 5.

#### 3.2 Reliability assessment via DEERS protocol and scoring of methodological rigour (outcome A) – additional results

Reporting deficiencies can be further investigated when looking at the relative scores of every data item subdomain (Tabs. S5 to S7). For the primary domain "BTS characterisation", four out of seven subdomains ("strain”, “pooling” (sex and number of individuals), and “husbandry details") showed below threshold reporting standards within the overall dataset, and all sub-datasets (Tab. S5). Especially "pooling” (number of individuals) and "husbandry details" scored low, with 22% and 33% within the overall dataset. For the sub-dataset "BTS1/endocrine", the subdomain "BTS induction" scored below the threshold (67%), and the overall dataset barely reached the standard (80%). The subdomains "species" and "BTS origin" scored relatively high numbers, with 96% and 92% respectively.

For the primary data item domain "BTS reaction" (Tab. S6), only one out of six categories, "BTS protein concentration", scored below the threshold within the total dataset (57%) and every sub-dataset (56, 43, and 70%, respectively). For the "BTS2/mutagen" dataset, the subdomains "primary" and "other cofactors" scored below the threshold (both 76%). Overall, the categories "buffer system" (97%), "BTS dilution factor" (89%), and "solvent" (87%) were rather well-described, whereas "primary" and "other cofactors" barely reached the threshold (81 and 83%, respectively).

Finally, the primary data item domain "BTS experimental setup" was best described, with total overall scores well above the threshold for all subdomains (Tab. S7). Especially "BTS incubation period" and "BTS incubation temperature" reached high reporting standards (both overall 95%). "BTS-related controls" were acceptably described but showed room for improvement (87% overall).

**Table S5:** Relative scores (in %) of every assessed subdomain of the primary domain "BTS characterisation". Scores below 80% are highlighted in red. For details, see also SI7.

| | **Species** | **BTS origin** | **Strain** | **Pooling – sex** | **Pooling – n** | **Husbandry details** | **BTS induction** |
|---|---|---|---|---|---|---|---|
| **BTS1 "endocrine"** | 100 | 95 | **69** | **46** | **21** | **23** | **67** |
| **BTS2 "mutagen"** | 90 | 85 | **63** | **56** | **12** | **30** | 80 |
| **BTS3 "historical"** | 99 | 96 | **74** | **61** | **31** | **40** | 85 |
| **Total** | 96 | 92 | **69** | **57** | **22** | **33** | 80 |

**Table S6:** Relative scores (in %) of every assessed subdomain of the primary domain "BTS reaction". Scores below 80% are highlighted in red. For details, see also SI7.

| | **BTS protein conc.** | **Buffer system** | **BTS dilution** | **Primary cofactors** | **Other cofactors** | **Solvent** |
|---|---|---|---|---|---|---|
| **BTS1 "endocrine"** | **56** | 92 | 87 | 82 | 85 | 90 |
| **BTS2 "mutagen"** | **43** | 96 | 83 | **76** | **76** | 87 |
| **BTS3 "historical"** | **70** | 99 | 94 | 85 | 87 | 87 |
| **Total** | **57** | 97 | 89 | 81 | 83 | 87 |

**Table S7:** Relative scores (in %) of every assessed subdomain of the primary domain "BTS experimental setup". For details, see also SI7.

| | **BTS incubation period** | **BTS incubation temperature** | **BTS related controls** |
|---|---|---|---|
| **BTS1 "endocrine"** | 87 | 95 | 85 |
| **BTS2 "mutagen"** | 96 | 97 | 94 |
| **BTS3 "historical"** | 97 | 94 | 81 |
| **Total** | 95 | 95 | 87 |

**Table S8:** Descriptive statistics of relative score populations (in %) of primary data item domains for the described datasets. Mean and median values below the 80% threshold are highlighted below in red.

| **BTS1** | **BTS char.** | **BTS rea.** | **BTS exp.** | **Total** | **BTS2** | **BTS char.** | **BTS rea.** | **BTS exp.** | **Total** |
|---|---|---|---|---|---|---|---|---|---|
| **N** | 39 | 39 | 39 | 39 | | 89 | 89 | 89 | 89 |
| **Minimum** | 14 | 17 | 33 | 19 | | 0 | 0 | 33 | 25 |
| **25% percentile** | 43 | 83 | 67 | 63 | | 43 | 58.5 | 100 | 63 |
| **Median** | **57** | 83 | 100 | **75** | | **71** | 83 | 100 | **75** |
| **75% percentile** | 71 | 100 | 100 | 88 | | 86 | 100 | 100 | 88 |
| **Maximum** | 100 | 100 | 100 | 100 | | 100 | 100 | 100 | 100 |
| **Mean** | **60** | 82 | 89 | **74** | | **59** | **77** | 96 | **73** |
| **BTS3** | **BTS char.** | **BTS rea.** | **BTS exp.** | **Total** | **BTS entire dataset** | **BTS char.** | **BTS rea.** | **BTS exp.** | **Total** |
| **N** | 101 | 101 | 101 | 101 | | 229 | 229 | 229 | 229 |
| **Minimum** | 0 | 33 | 33 | 50 | | 0 | 0 | 33 | 19 |
| **25% percentile** | 43 | 83 | 67 | 69 | | 43 | 83 | 100 | 69 |
| **Median** | **71** | 100 | 100 | 81 | | **71** | 83 | 100 | 81 |
| **75% percentile** | 93 | 100 | 100 | 94 | | 86 | 100 | 100 | 88 |
| **Maximum** | 100 | 100 | 100 | 100 | | 100 | 100 | 100 | 100 |
| **Mean** | **70** | 87 | 91 | 80 | | **64** | 82 | 92 | **76** |

Variation between datasets' primary domains was compared statistically to analyse coherence (Fig. S2). Absolute scoring populations were analysed via Kruskal-Wallis tests accompanied by Dunn's post hoc test for multiple comparisons. Statistically significant variance ($p < 0.05$) between primary domains "BTS characterisation", "BTS reaction", and the overall total score was computed for the "BTS2/mutagen" vs "BTS3/historical" datasets (Fig. S2A, B, and D). Taken together with Fig. 3 (main article), this implicates some study selection bias, which is further elaborated within the discussion (section 4.1, below).

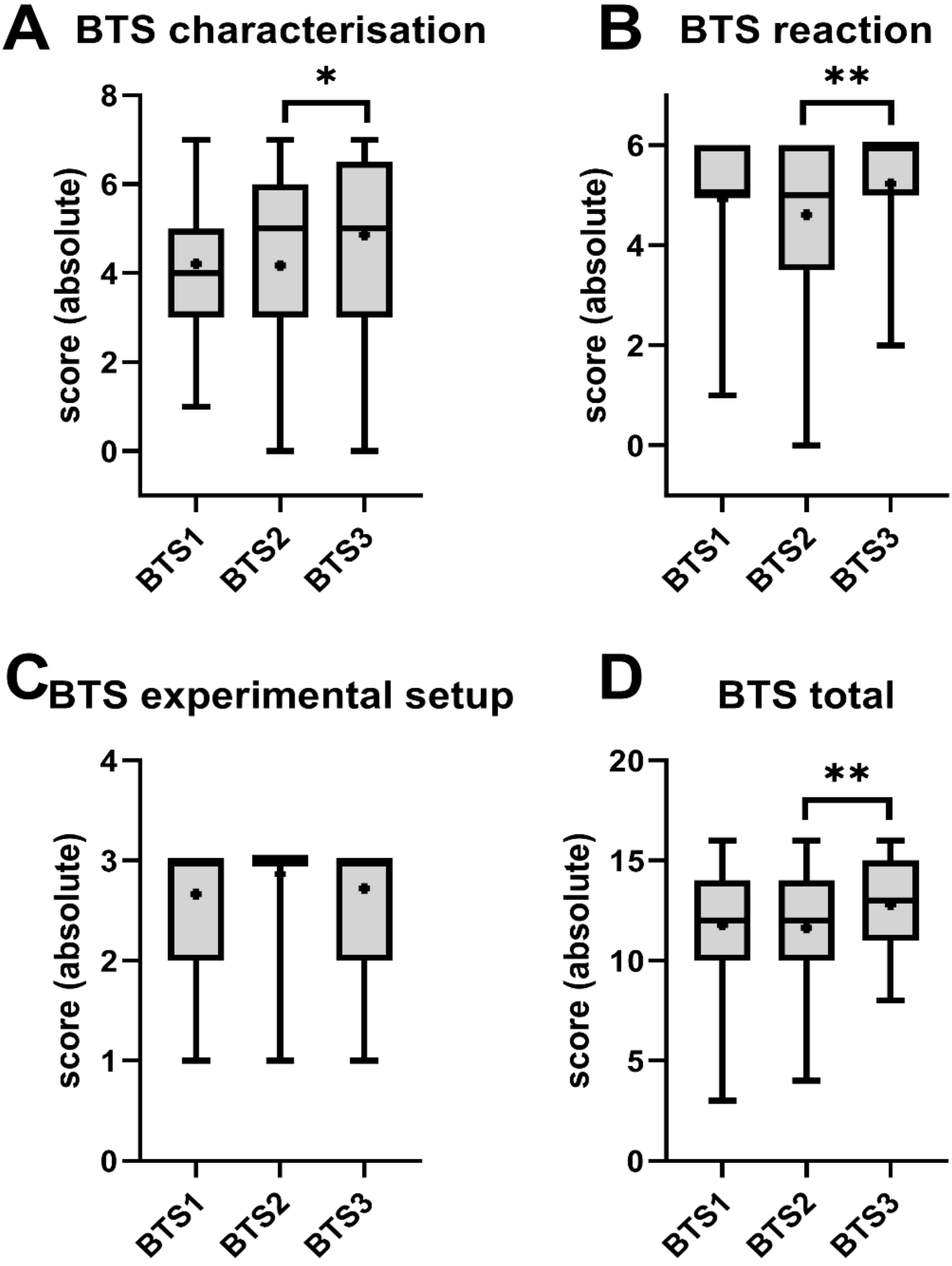

**Fig. S2**: Boxplots depicting absolute scoring populations of reviewed and assessed articles for all datasets, ordered by primary data item domains for comparison (panels A to D). Whiskers indicate the populations' upper (max.) to lower (min.) boundaries. Boxes indicate the 75th and 25th percentile, and the in-between line represents the median. Dots represent mean population values. Statistical analyses of variance between datasets were conducted via Kruskal-Wallis tests accompanied by Dunn's post hoc test for multiple comparisons. Asterisks indicate statistically differing significance between means of respective datasets (*$p < 0.05$, **$p < 0.01$). The number of included articles within every dataset is given, e.g., in Tab. S8.

3.3 Meta-regression of quantitative BTS reaction components (outcome B) – additional results

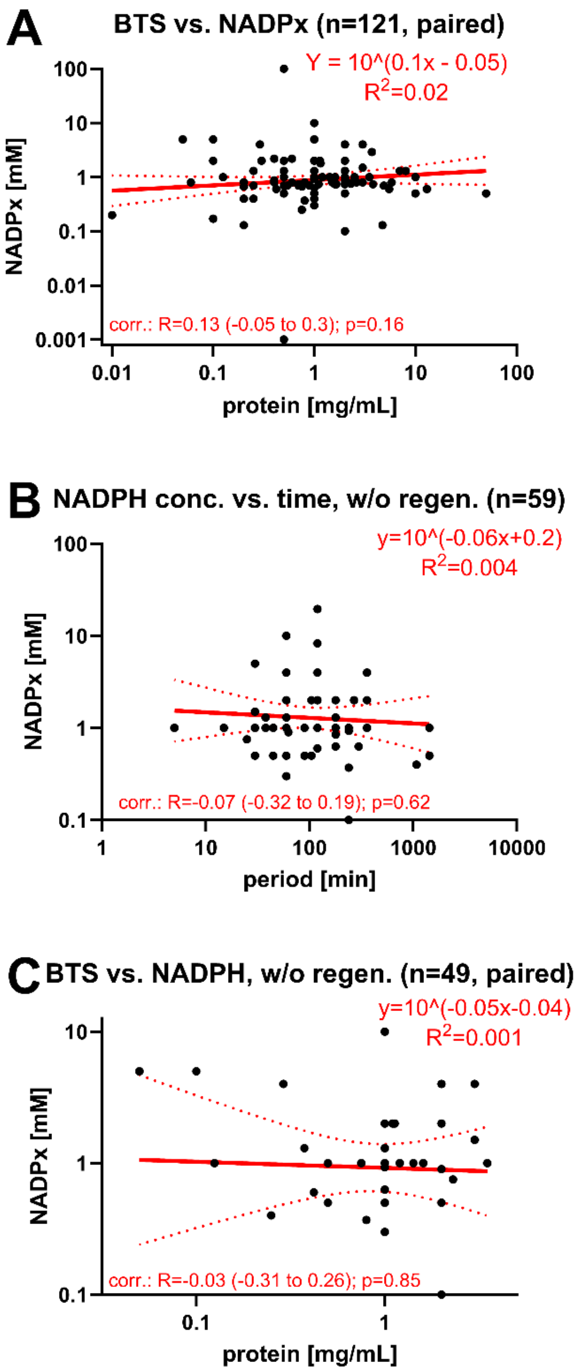

**Fig. S3:** Simple linear regression fits of numerical data item measures hypothesised to show a particular mathematical relationship. Data were extracted from the overall database (SI6) and manually curated as outlined in section 2.7 (main article), with detailed seminal populations presented in SI8. Numerical data were log-log transformed before fitting a simple linear regression. Equations and fits ($R^2$ values) are given within the respective graphs. Single measures are illustrated as dots. The seminal populations are provided within the titles. The regression fits are displayed as red lines, with 95% CIs as dotted red lines. Pearson's correlation was computed between variables, the respective coefficients of correlation (R), coefficients of determination ($R^2$, same as for linear fit), and 95% CIs are given within the graphs. P-values determine the statistical significance of correlation. The following pairs were tested: (A) paired BTS protein concentration vs. primary cofactor concentration; (B) primary cofactor concentration (w/o regeneration system) vs. BTS incubation period; (C) paired BTS protein concentration vs. primary cofactor concentration (w/o regeneration system).

#### 3.4 Descriptive statistics of qualitative data item subdomains (outcome C) – results in detail by subdomain

The following sections of the SM (3.4.1 to 3.4.14) present more detailed descriptive statistics of qualitative data item subdomains, where every analysed subdomain is presented and shortly discussed independently. This preceding section summarises the most notable patterns observed.

In terms of chronological patterns, there has been a substantial increase in the diversification of the "field/journal" subdomain of BTS-related publications post-millennium. This shift has been from classical toxicology to a broader range that includes environmental toxicology and analytical chemistry, among other fields, as elaborated in Fig. S4C. Similarly, the "study endpoints" subdomain has evolved, showing increasing diversification. The initial focus on mutagenicity and genotoxicity studies has expanded towards encompassing the assessment of endocrine disruptors, metabolites, and xenobiotic metabolism functionality, as illustrated in Fig. S6C. Furthermore, the "species" derivation subdomain within BTS has also diversified. The trend has moved from predominantly rat-based systems to a more varied range, including systems derived from humans, mice, fish, and other species, as detailed in Fig. S9C. The last decade has witnessed an increase in the utilisation of externally purchased BTS, as seen in Fig. S8C. This increase in external sourcing correlates with a decrease in reporting accuracy in subdomains such as “strain” (Fig. S11C), “pooling” (Fig. S12C), and “induction” (Fig. S14C).

Another noteworthy pattern in the descriptive statistics analyses relates to the primary cofactor (NADPx) regeneration systems. Out of the n = 177 studies employing cofactor regeneration systems, only n = 35 provide detailed reporting on the essential dehydrogenase components. This gap in reporting is evident in Figs. S16 to S18.

##### 3.4.1 Journal/Field

Toxicology (n = 147) and environmental sciences (n = 69) are the most prevalent fields within the overall records (Fig. S4). The relation frequency (Fig. S4B) of other areas, except for nutritional sciences, places them in context with (environmental) toxicology, analytical chemistry, and biomedicine. The diversification of the field of publication has increased since the onset of the new millennium (Fig. S4C). Since the 2000s, the diversification in publication fields has expanded (Fig. S4C), with toxicology historically leading but now sharing prominence with environmental toxicology, analytical chemistry (n = 19), and veterinary sciences (n = 6).

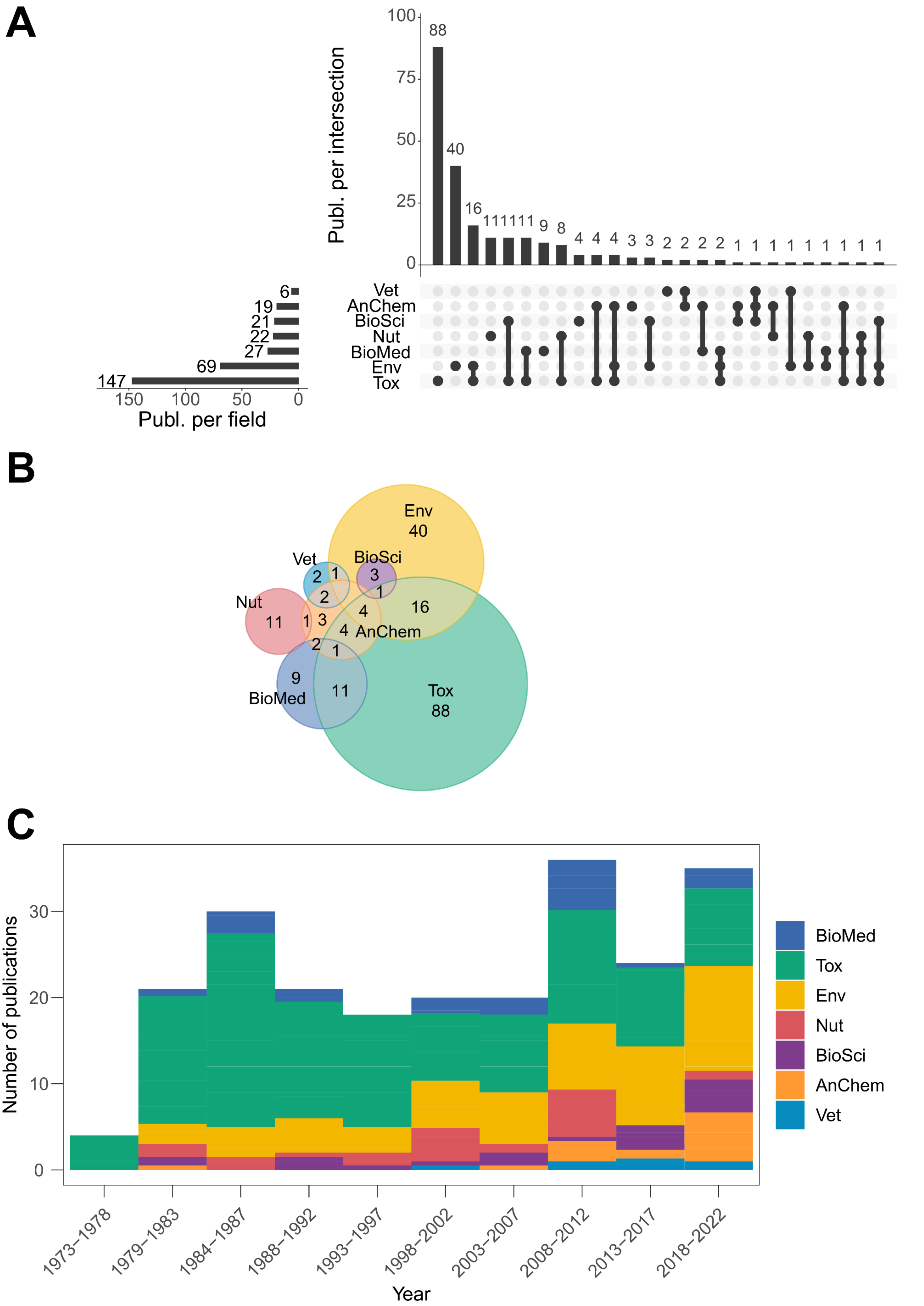

**Fig. S4**: Descriptive statistics of the data item subdomain "journal" (simplified to "field"), illustrated as interaction frequencies (upset plot, panel A), relation frequencies (Euler diagram, panel B), and distribution frequencies over the years (histogram, panel C). Abbreviations: BioMed – biomedicine; Tox – toxicology; Env – environmental sciences; Nut – nutritional sciences; BioSci – general biosciences; AnChem – analytical chemistry; Vet – veterinary sciences.

#### 3.4.2 Test system

Studies employing eukaryotic systems along BTS are most frequent (n = 124), followed by BTS-only test systems (n = 71) and test systems utilising bacteria (n = 52) (Fig. S5A and B). The combination of eukaryotic and prokaryotic systems (n = 24) primarily originates from early mutagenicity and genotoxicity studies. Historically, eukaryotic *in vitro* test systems, together with BTS systems, were more common but have recently been supplanted by BTS-only systems (Fig. S5C).

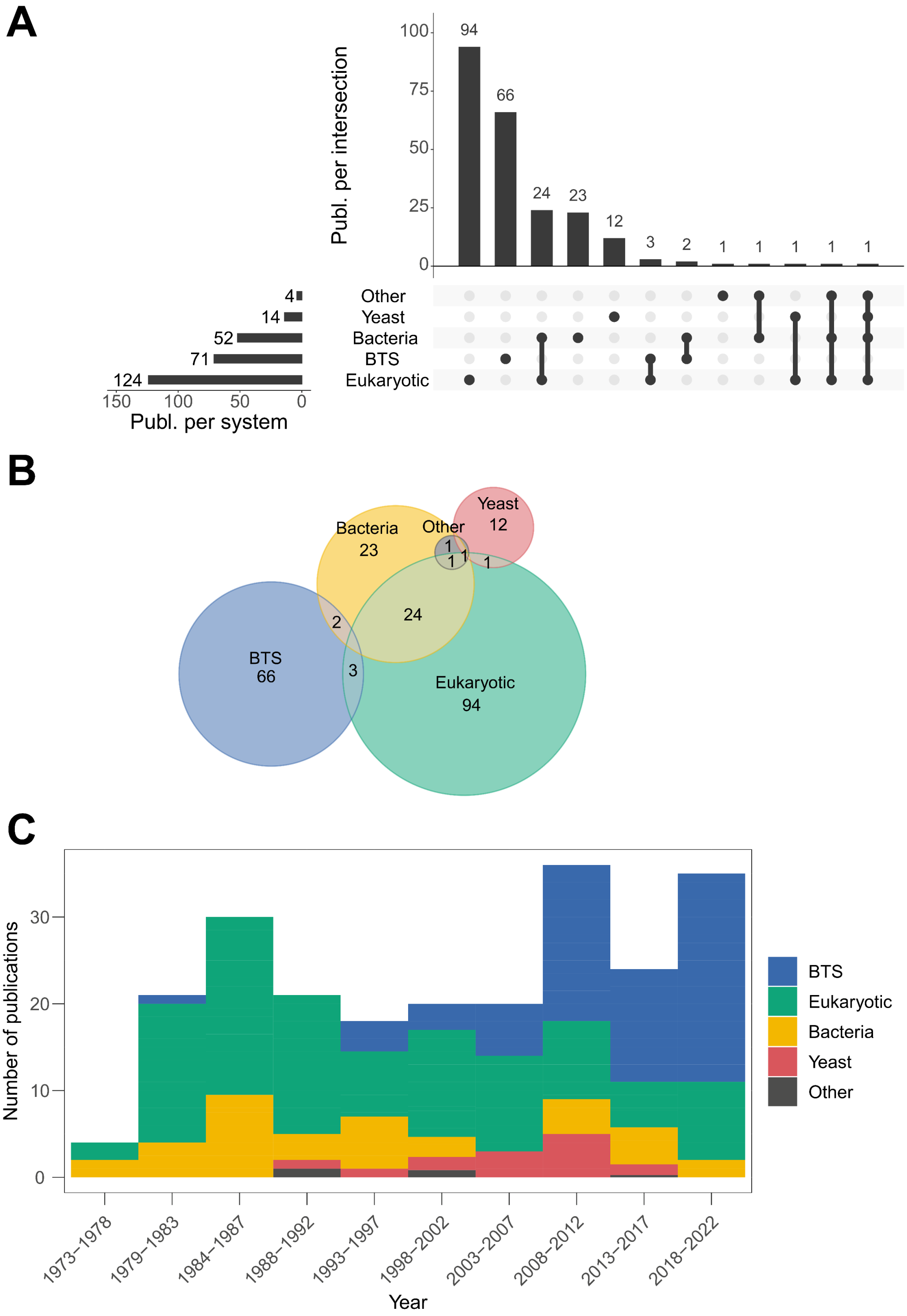

**Fig. S5**: Descriptive statistics of the data item subdomain “test system”, illustrated as interaction frequencies (upset plot, panel A), relation frequencies (Euler diagram, panel B), and distribution frequencies over the years (histogram, panel C).

3.4.3 Endpoint

Studies frequently cover mutagenicity and genotoxicity (n = 122), cytotoxicity (n = 87), metabolites (n = 87), endocrine disruptors (n = 34), and xenobiotic metabolism activity (n = 17) endpoints (Fig. S6A). Mutagenicity and genotoxicity studies often coincide frequency-wise with assessing cytotoxicity (n = 72) (Fig. S6A and B), whereas studies investigating biotransformation metabolites coincide with recording endocrine disruption (n = 17) and assessing xenobiotic metabolism mechanisms (n = 7). Study endpoints have become more diverse over the years (Fig. S6C). While studies on mutagenicity and genotoxicity were dominant until the dawn of the new millennium, a shift in study endpoints towards endocrine disruptors and biotransformation metabolites has been observed post-millennium, with a recent rise in neurological and developmental endpoint studies (Fig. S6C).

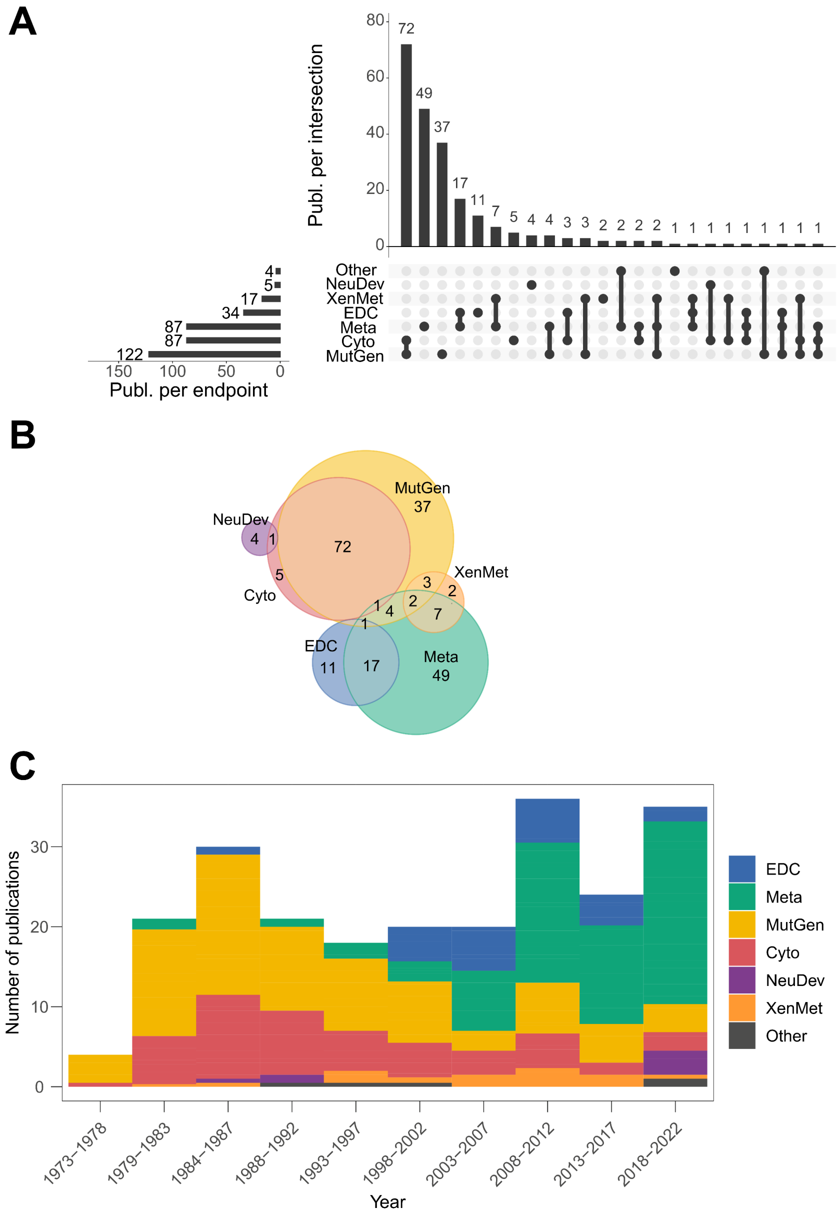

**Fig. S6**: Descriptive statistics of the data item subdomain investigated "endpoint", illustrated as interaction frequencies (upset plot, panel A), relation frequencies (Euler diagram, panel B), and distribution frequencies over the years (histogram, panel C). Abbreviations: EDC – endocrine disruption; Meta – metabolites; MutGen – mutagenicity and genotoxicity; Cyto – cytotoxicity; NeuDev – neurosciences and developmental biology; XenMet – xenobiotic metabolism functionality.

#### 3.4.4 BTS type

S9 (n = 200) and microsomes (n = 63) are the most prevalent BTS types (Fig. S7) frequency-wise. Utilisation of both types within a specific study is relatively frequent (n = 34, Fig. S7A and B). Only very few studies (n = 2) compared S9 directly with primary hepatocytes.

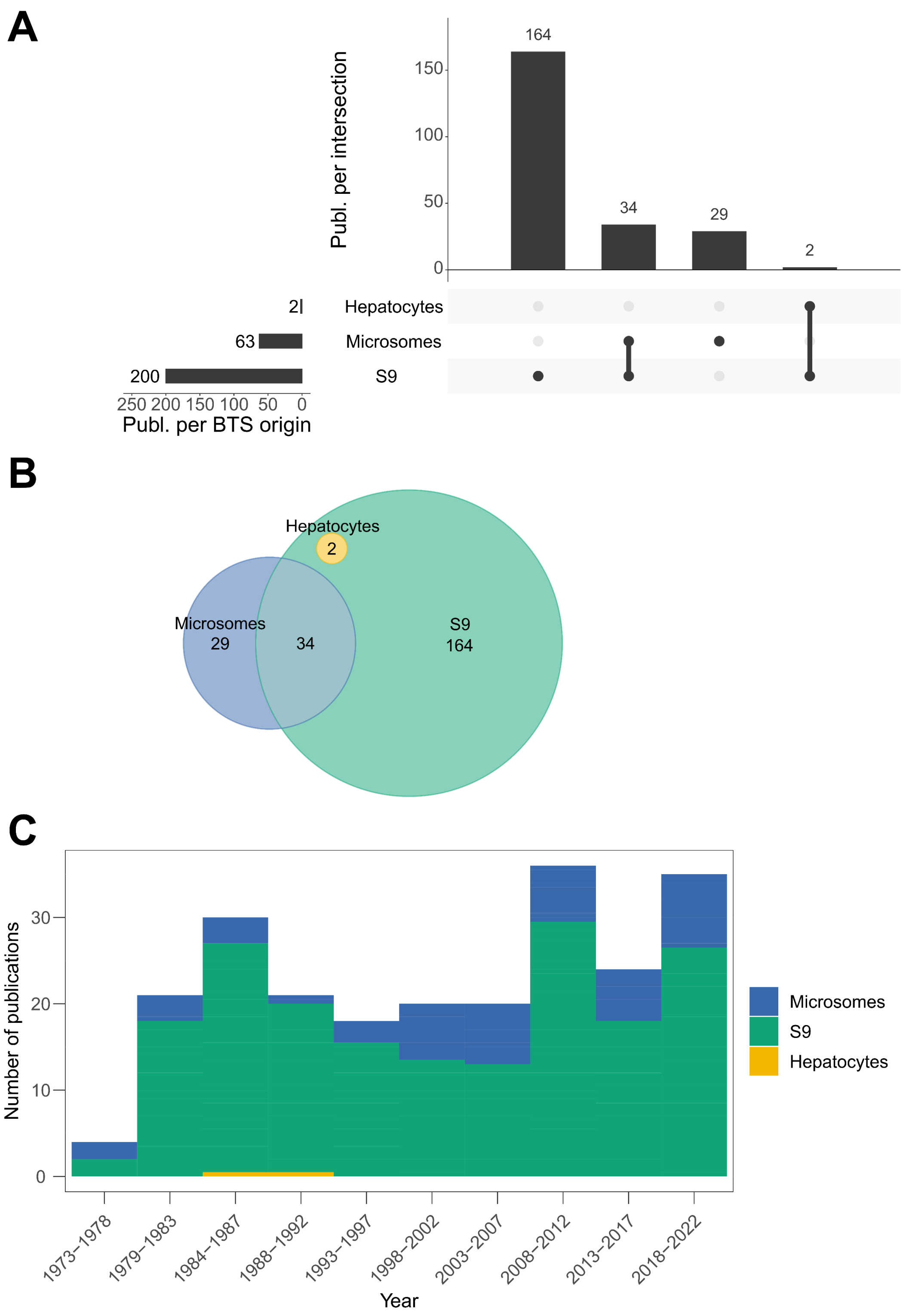

**Fig. S7**: Descriptive statistics of the data item subdomain "BTS type", illustrated as interaction frequencies (upset plot, panel A), relation frequencies (Euler diagram, panel B), and distribution frequencies over the years (histogram, panel C).

3.4.5 BTS origin

Externally sourced BTS (n = 118) have gained prominence over time (Fig. S8C). The number of studies utilising both internally produced and externally purchased BTS is rather low (n = 13, Fig. S8A and B). A noteworthy number of studies (n = 19) do not disclose BTS origin.

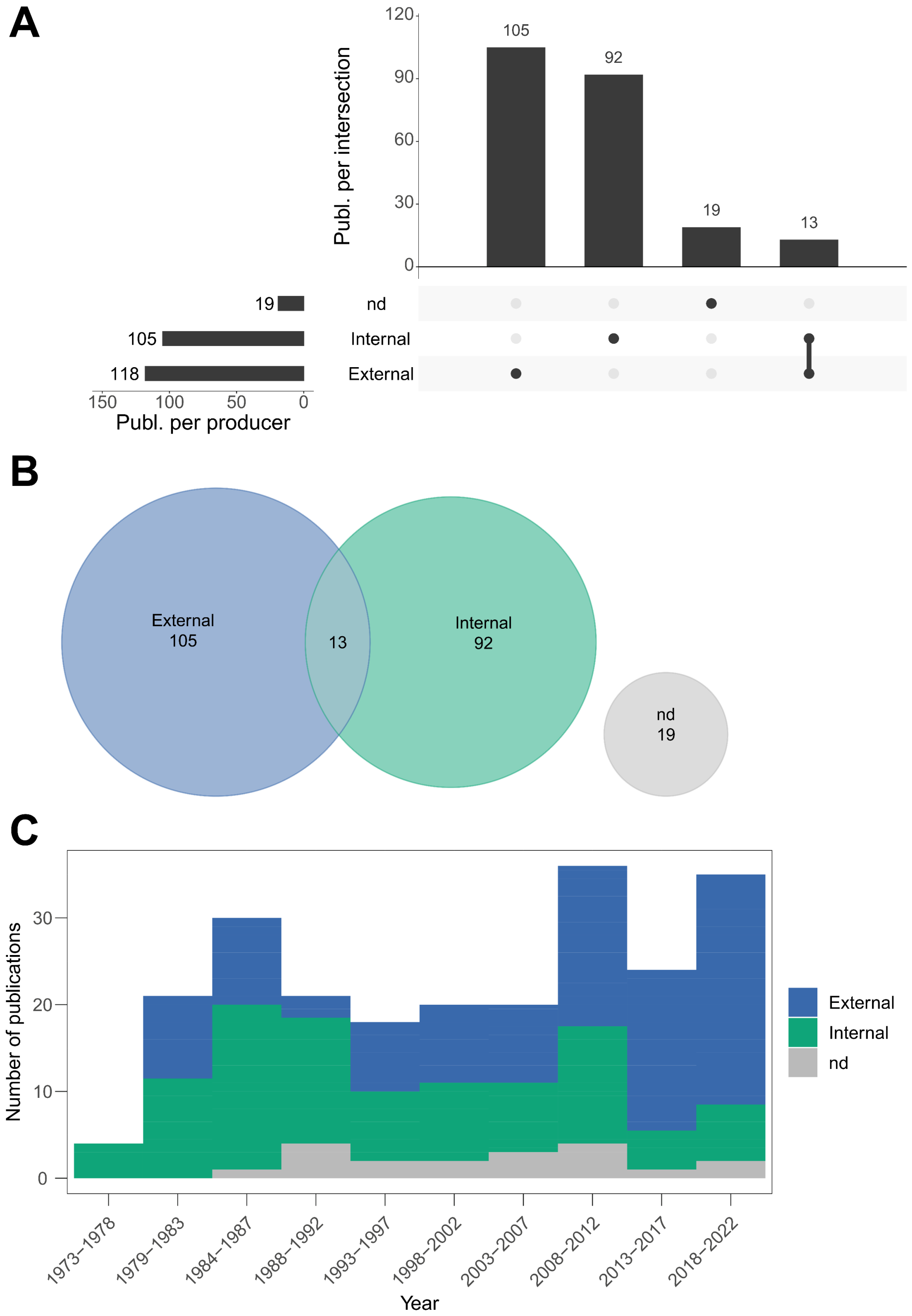

**Fig. S8**: Descriptive statistics of the data item subdomain "BTS origin", illustrated as interaction frequencies (upset plot, panel A), relation frequencies (Euler diagram, panel B), and distribution frequencies over the years (histogram, panel C).

3.4.6 Species

Species diversification in BTS sources has significantly increased over the last 20 years (Fig. S9C). However, mammalian systems remain dominant (n = 208, Fig. S10B). Rat-derived BTS was most common until the 1990s. It is rather uncommon to utilise more than one species of BTS origin (n = 34) (Fig. S9A). Besides rats, other mentionable BTS sources are humans (n = 42), mice (n = 12), and fish (n = 17).

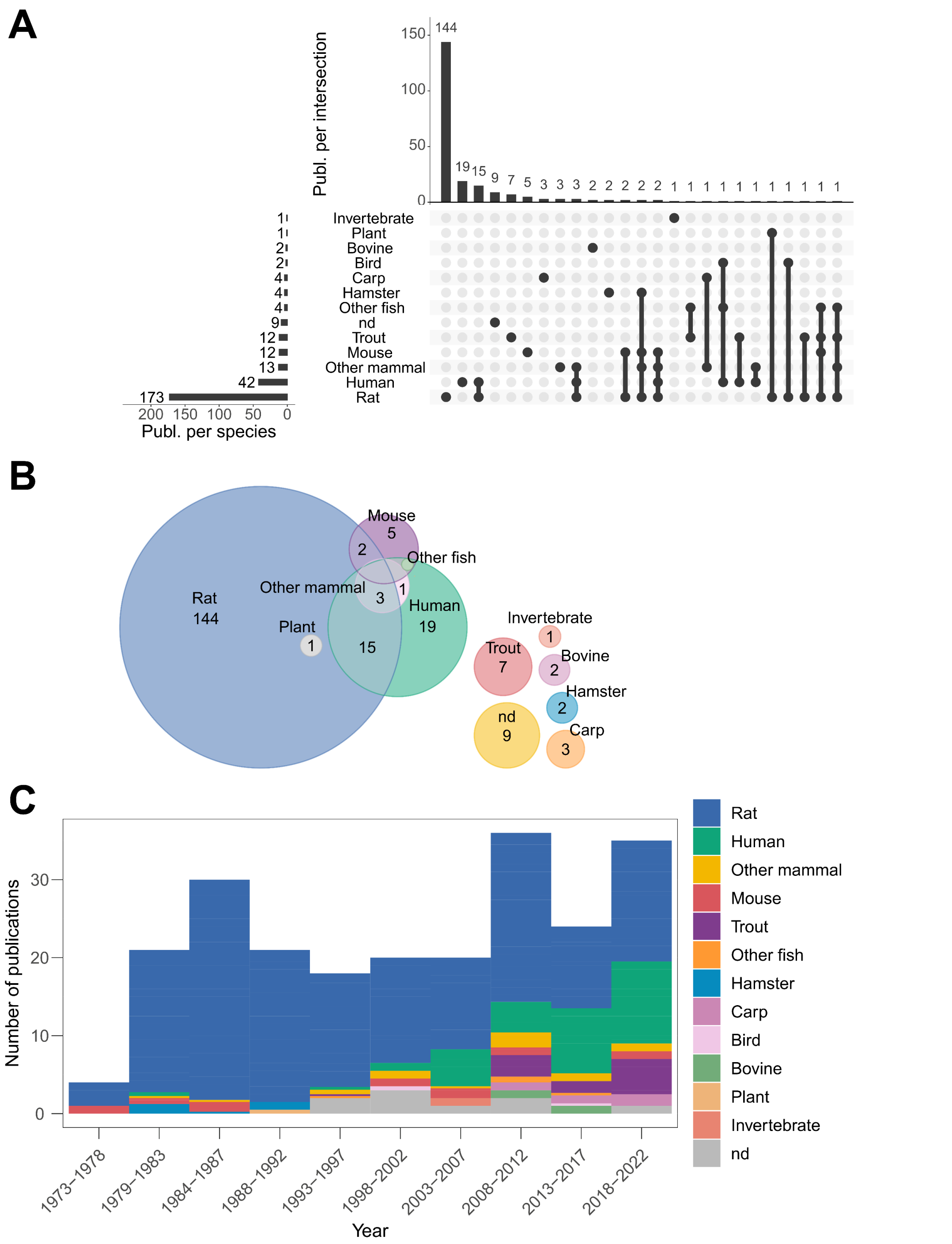

**Fig. S9**: Descriptive statistics of the data item subdomain "species" of BTS derivation, illustrated as interaction frequencies (upset plot, panel A), relation frequencies (Euler diagram, panel B), and distribution frequencies over the years (histogram, panel C).

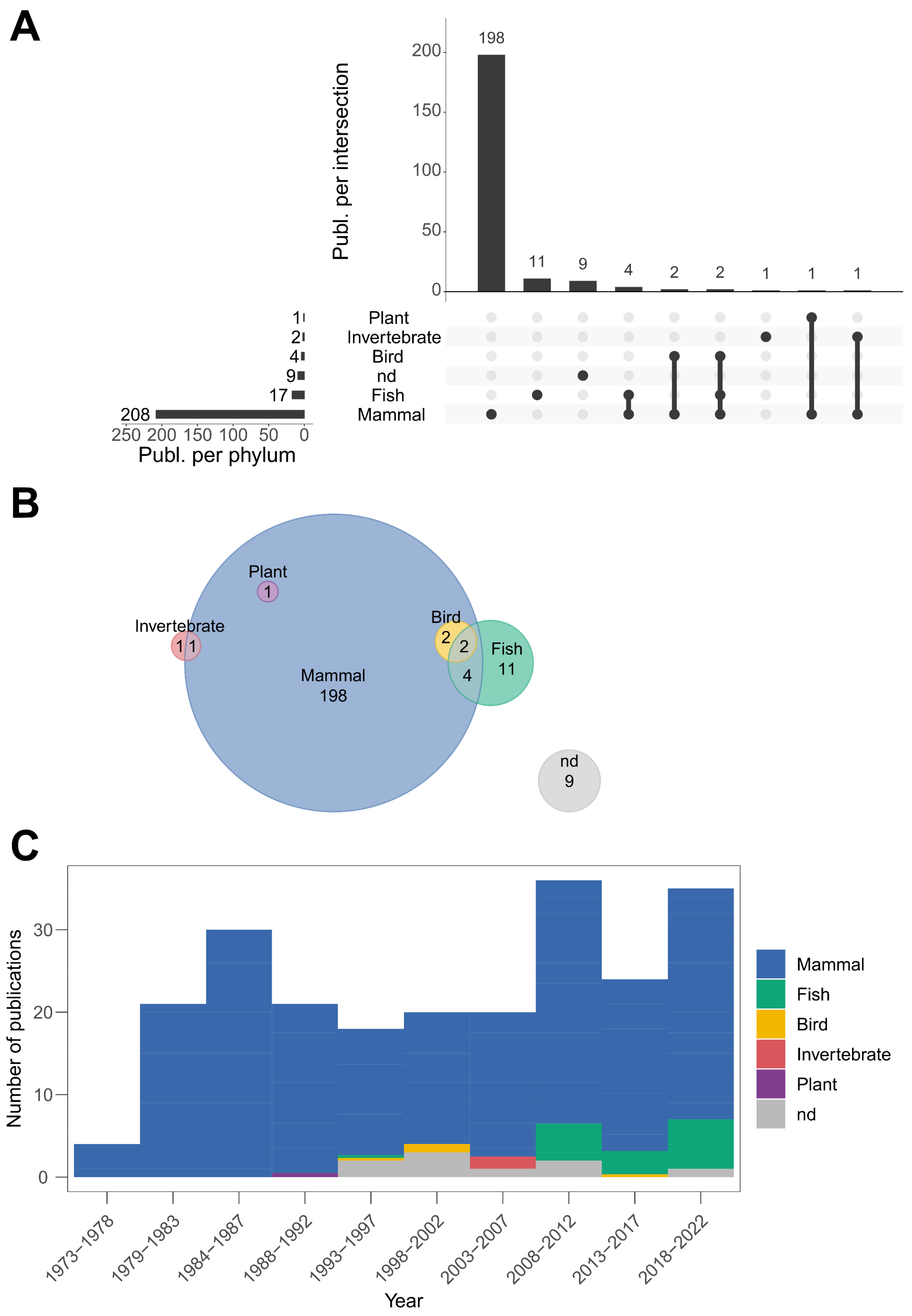

682

683 **Fig. S10**: Descriptive statistics of the data item subdomain "species" of BTS derivation (summarised to clade), illustrated as
684 interaction frequencies (upset plot, panel A), relation frequencies (Euler diagram, panel B), and distribution frequencies
685 over the years (histogram, panel C).

#### 3.4.7 Strain

As described above, an increase in non-rodent BTS frequency is evident throughout the years (here reflected as not applicable – “na”, n = 56). The Sprague-Dawley strain is the most common (n = 84), followed by Wistar rats (n = 22) (Fig. S11A and B). Only n = 5 studies employ multiple rat strains. The pattern of rat strain utilisation remained stable throughout the years (Fig. S11C). Many studies (n = 60) do not report the strain of BTS derivation.

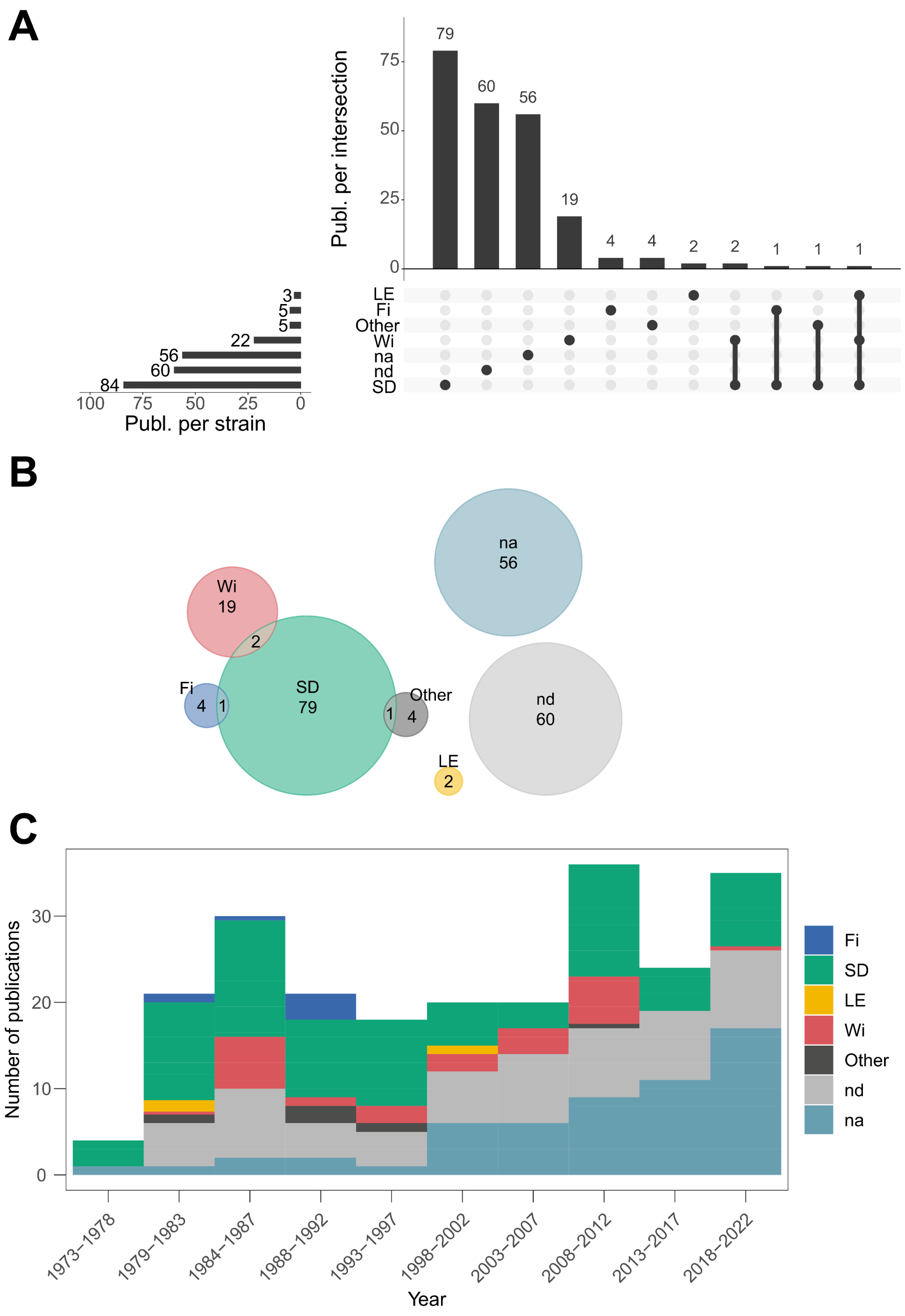

**Fig. S11**: Descriptive statistics of the data item subdomain utilised rodent “strain”, illustrated as interaction frequencies (upset plot, panel A), relation frequencies (Euler diagram, panel B), and distribution frequencies over the years (histogram, panel C). Abbreviations: Fi – Fischer; SD - Sprague Dawley; LE - Long Evans; Wi – Wistar.

3.4.8 BTS pooling

Male-only derived BTS is the most frequent (n = 88) type, followed by mixed derivation systems (n = 34) (Fig. S12). Female-only systems are rarely applied (n = 9) The prevalence of “not defined” (“nd”) has increased over the years (n = 95), most likely associated with external BTS sourcing.

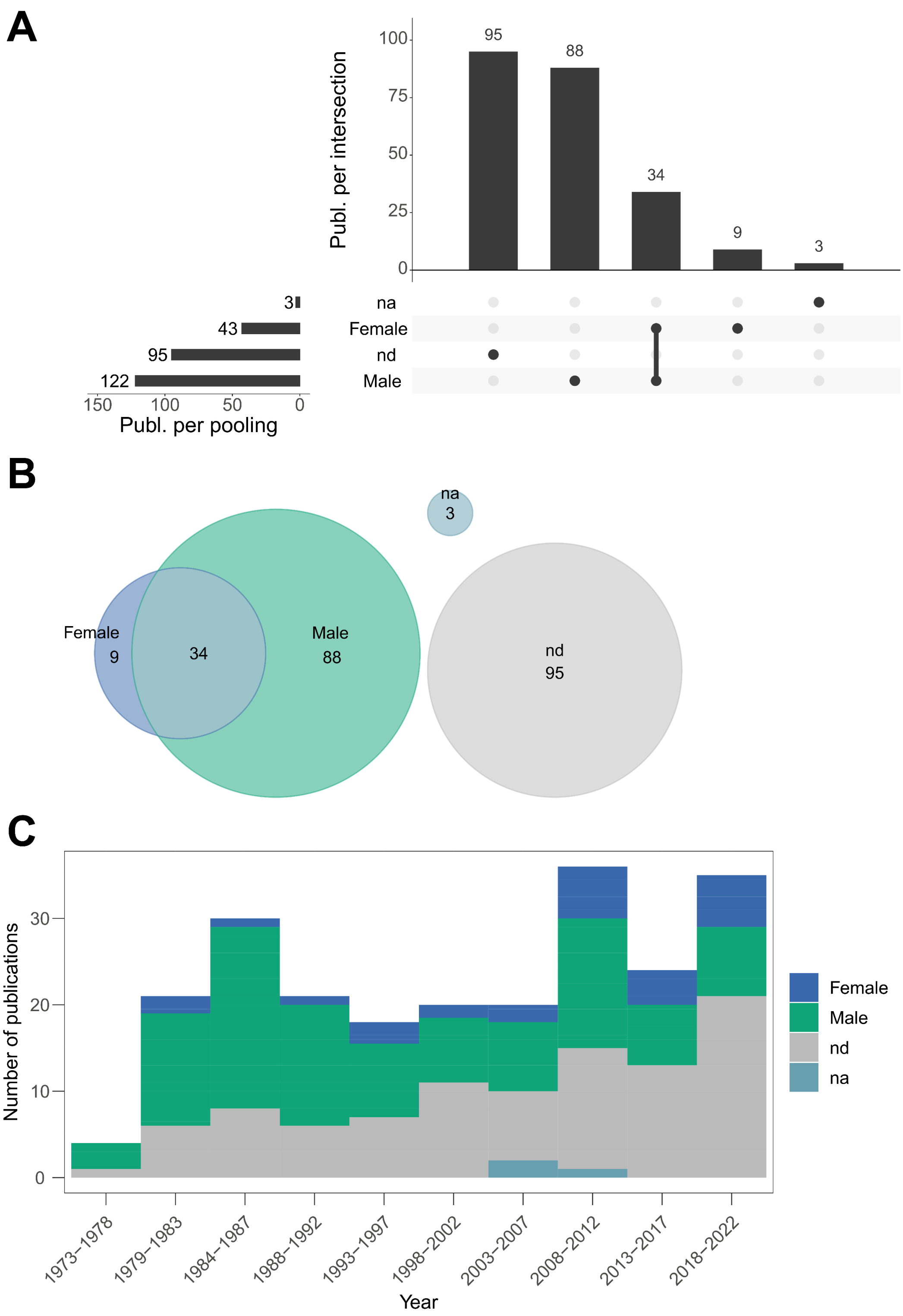

**Fig. S12**: Descriptive statistics of the data item subdomain “BTS pooling”, illustrated as interaction frequencies (upset plot, panel A), relation frequencies (Euler diagram, panel B), and distribution frequencies over years (histogram, panel C).

#### 3.4.9 Husbandry

Husbandry details are generally poorly described (Fig. S13B), with n = 154 studies defined as “nd”. Few studies detailing husbandry (n = 45) also report BTS enzymatic activity (n = 9). Studies mentioning only activity are either *in vitro* or human-derived systems (n = 6, Fig. S13A). A decline in husbandry reporting is likely due to increased external BTS sourcing (Fig. S13C). The abbreviation “other” (n = 24) defines studies where the husbandry details could be found in a direct citation or protocol.

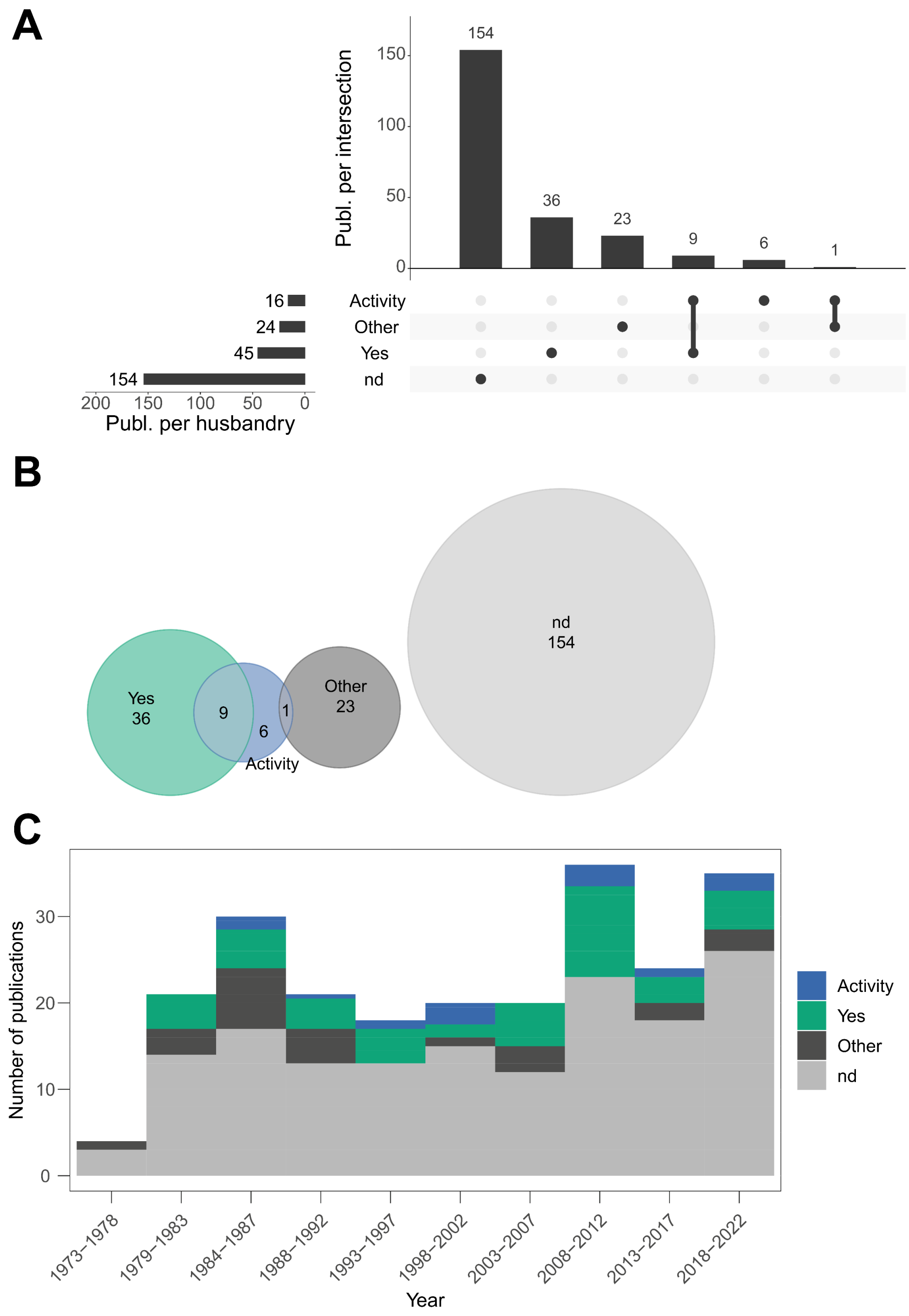

**Fig. S13:** Descriptive statistics of the data item subdomain “husbandry”, illustrated as interaction frequencies (upset plot, panel A), relation frequencies (Euler diagram, panel B), and distribution frequencies over years (histogram, panel C).

3.4.10 BTS induction

PCB-induced ("Aro") BTS are most frequent throughout the recorded and analysed literature (n = 99) (Fig. S14), followed by non-induced systems (n = 69) and BNF/PB mixtures (n = 16). Only a few studies investigate alternative induction regimes (MCA, n = 5; other, n = 5). The prevalence of using solely non-induced BTS testing systems (n = 51) has increased over the years, as has the number of "nd" studies (n = 46). The phaseout of Aroclor-induced BTS is evident in the data (Fig. S14C).

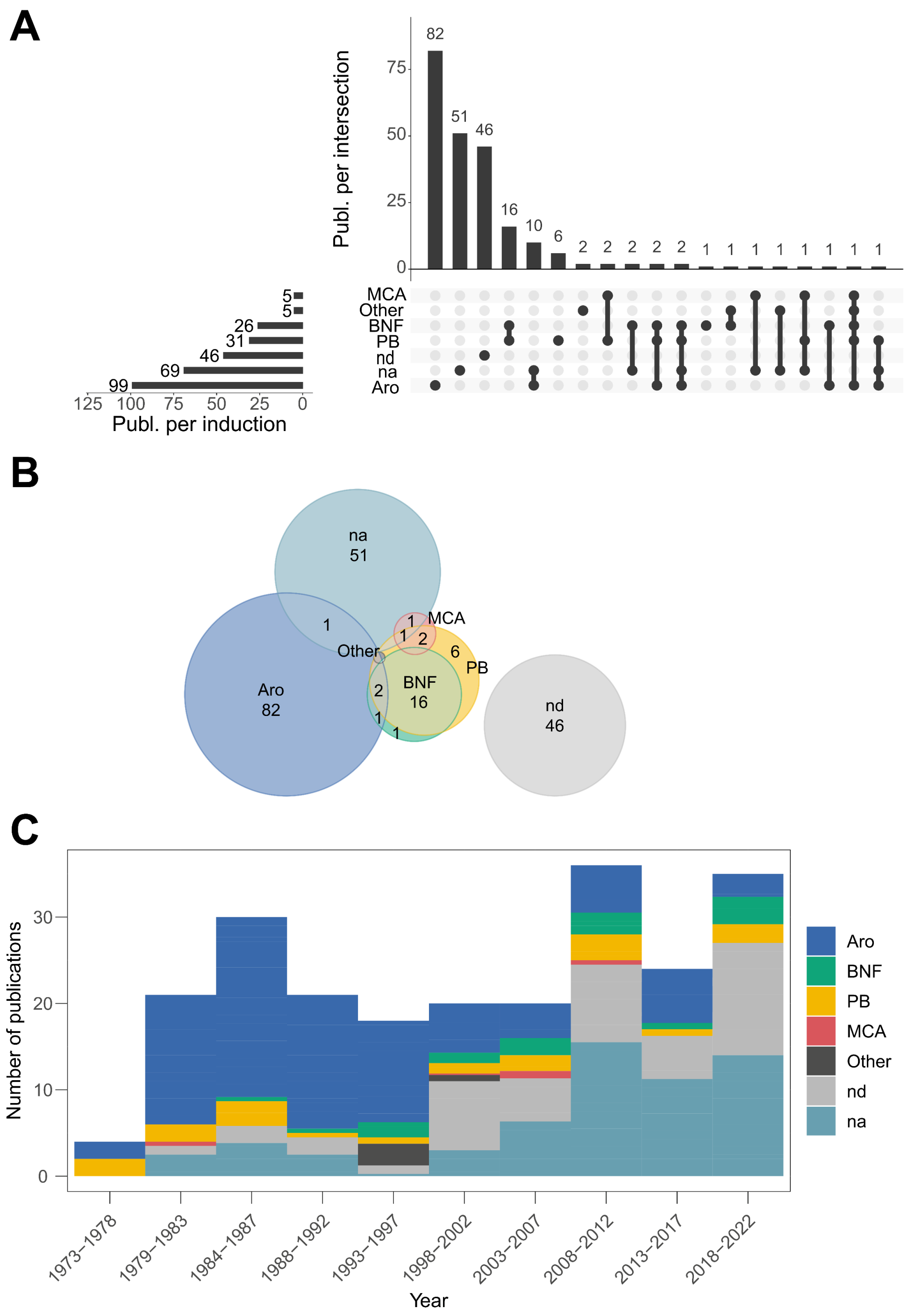

**Fig. S14**: Descriptive statistics of the data item subdomain "BTS induction", illustrated as interaction frequencies (upset plot, panel A), relation frequencies (Euler diagram, panel B), and distribution frequencies over the years (histogram, panel C). Abbreviations: Aro – Aroclor and other PCBs; BNF - beta-Naphthoflavone; PB – phenobarbital; MCA - methylcholanthrene.

3.4.11 Buffer system

Cell culture medium is the most frequent buffer system (n = 122), followed by phosphate buffers (n = 100) and Tris buffers (n = 14), the latter used primarily in analytical chemistry studies (Fig. S15). Only a few studies (n = 16) simultaneously employ various buffer systems, typically investigating endpoint or test systems (bioanalytics and chemical analytics). Phosphate buffer use has risen alongside BTS-only test systems (Fig. S15C).

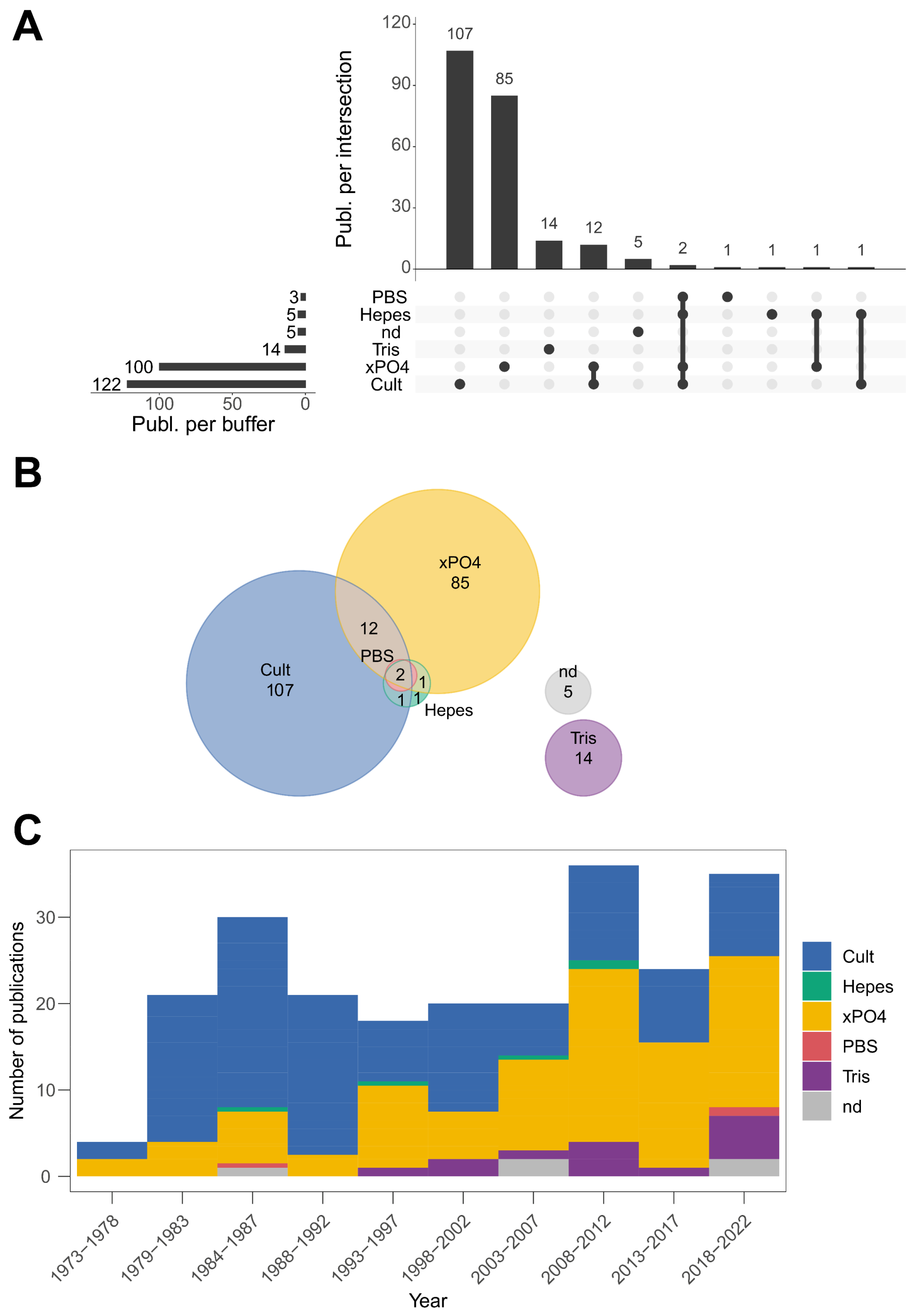

**Fig. S15**: Descriptive statistics of the data item subdomain "buffer system", illustrated as interaction frequencies (upset plot, panel A), relation frequencies (Euler diagram, panel B), and distribution frequencies over the years (histogram, panel C). Abbreviations: Cult – culture medium; xPO4 – various forms of phosphate buffers.

#### 3.4.12 Cofactors

Studies focusing on phase 2 biotransformation (n = 3) are rare (Fig. S16). Most studies focus solely on phase 1 metabolism (n = 156), with some investigating both phases (n = 39). The number of studies employing NADPx-regeneration systems (n = 178) has decreased over the years (Fig. 17C). Very few of the studies employing regeneration systems record necessary details (n = 35, dehydrogenase cofactors, see Fig. 18B). Unfortunately, failure to report technical details about the dehydrogenase system is very common (n = 109 + 33).

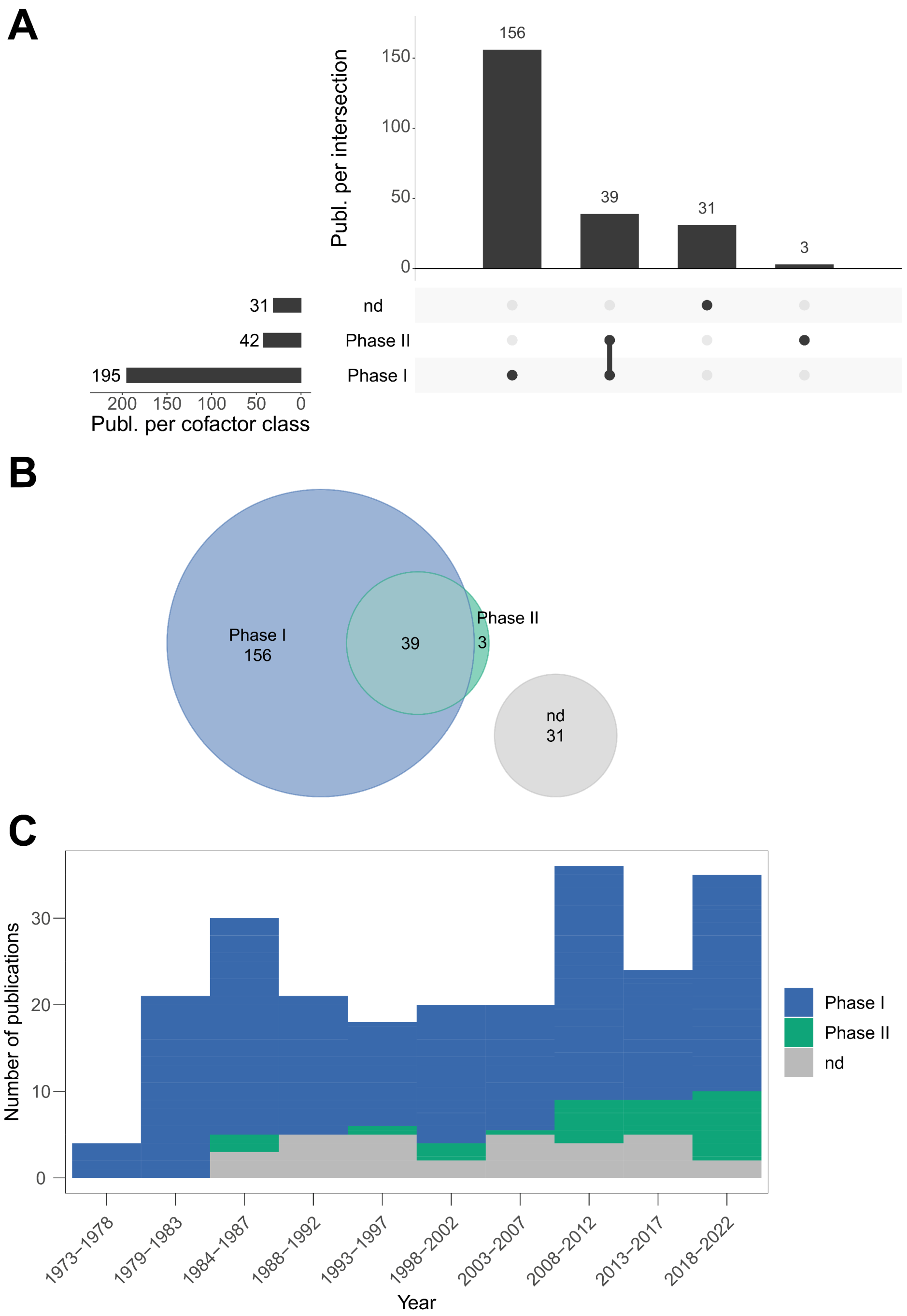

**Fig. S16**: Descriptive statistics of the data item subdomain "cofactor" (as phase classes 1 or 2), illustrated as interaction frequencies (upset plot, panel A), relation frequencies (Euler diagram, panel B), and distribution frequencies over the years (histogram, panel C).

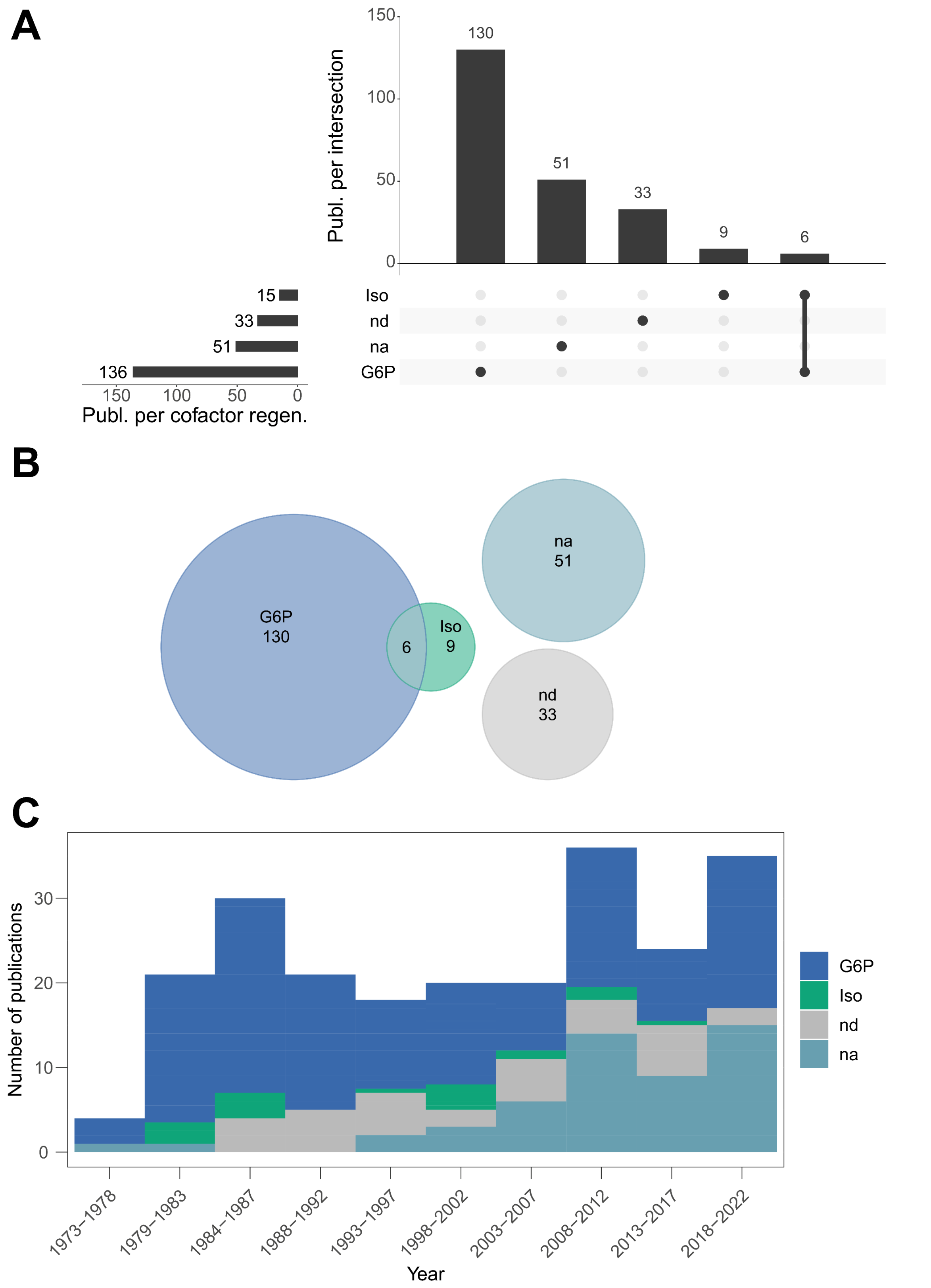

**Fig. S17**: Descriptive statistics of the data item subdomain "cofactor" (as dehydrogenase system), illustrated as interaction frequencies (upset plot, panel A), relation frequencies (Euler diagram, panel B), and distribution frequencies over the years (histogram, panel C). Abbreviations: G6P – glucose-6-phosphate; Iso – isocitrate; nd - dehydrogenase system not defined although necessary in this experimental setup; na – dehydrogenase system not applicable (uses NADPH).

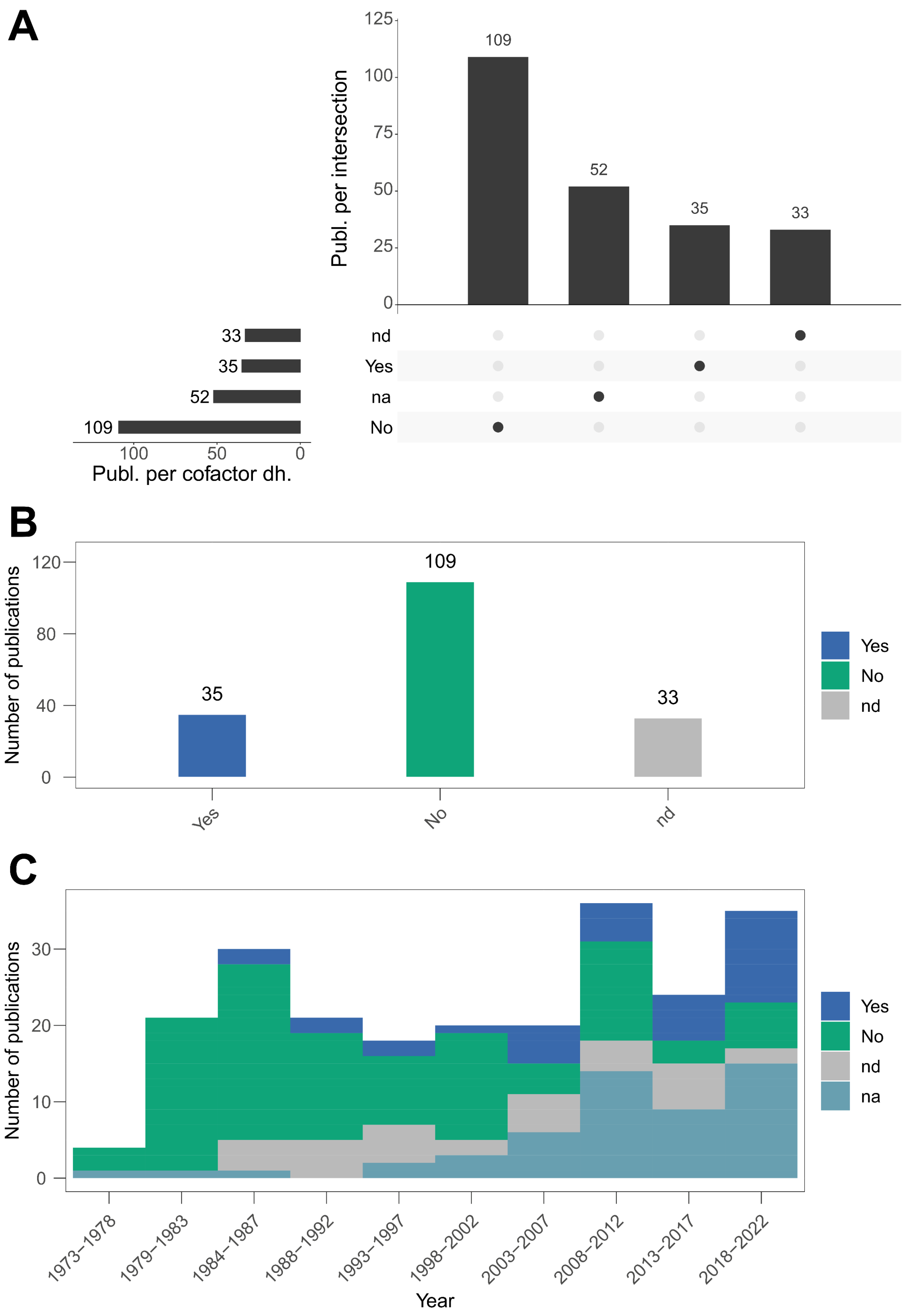

**Fig. S18**: Descriptive statistics of the data item subdomain “cofactor” (as dehydrogenase system fully defined), illustrated as interaction frequencies (upset plot, panel A) and distribution frequencies over the years (histograms, panels B and C). Abbreviation: dh – dehydrogenase; Yes – fully defined dh system; No – dh system definition is missing crucial information; nd - dehydrogenase system not defined although necessary in this experimental setup; na – dehydrogenase system not applicable (uses NADPH).

#### 3.4.13 Solvents

DMSO (n = 132) is the most frequently utilised solvent, followed by aqueous solutions (H20, n = 45), alcoholic solvents (Alc, n = 45), and various other organic solvents (n = 19) (Fig. S19). Solvent application frequency has remained relatively unchanged over time (Fig. S19C).

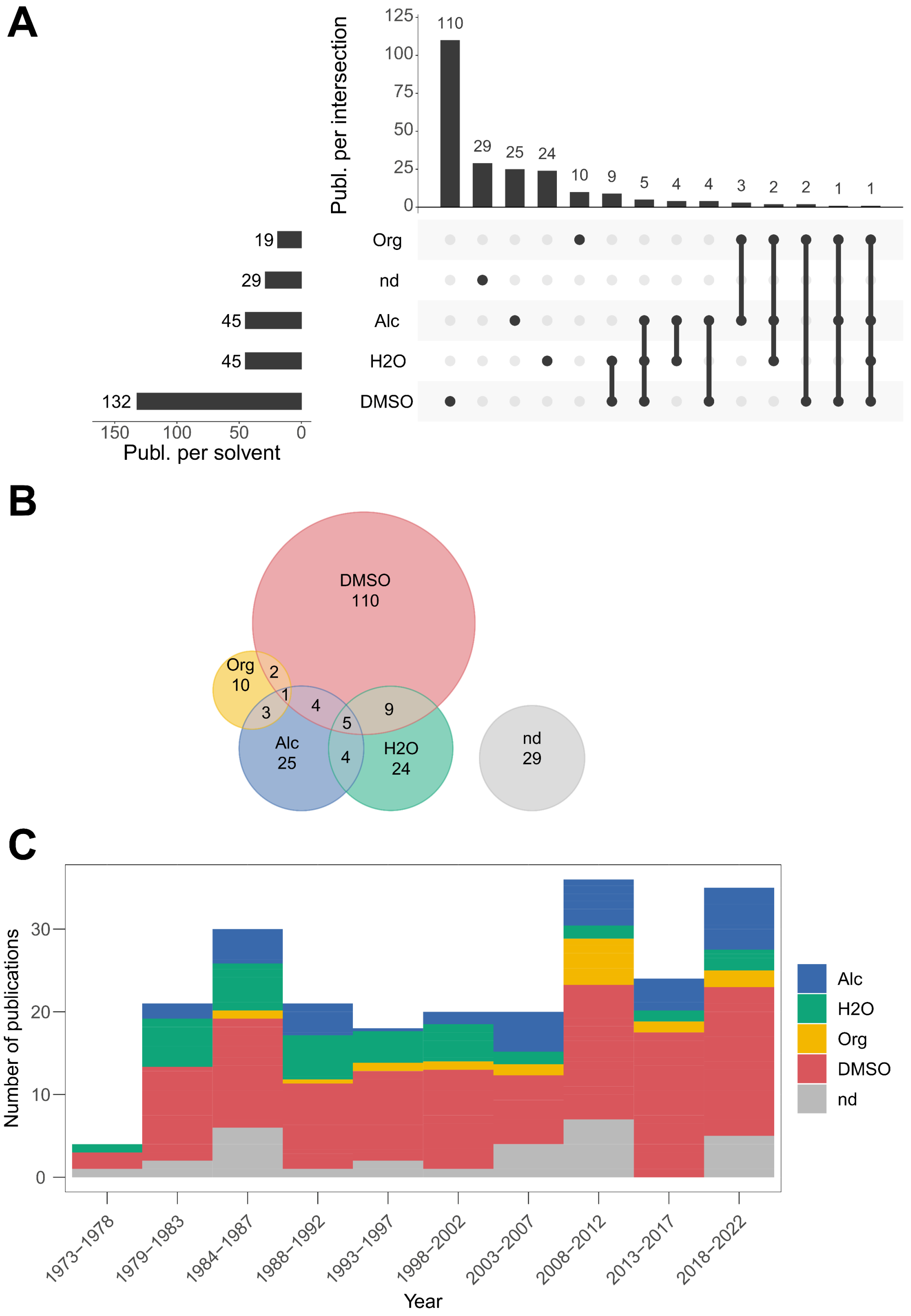

**Fig. S19**: Descriptive statistics of the data item subdomain “solvents”, illustrated as interaction frequencies (upset plot, panel A), relation frequencies (Euler diagram, panel B), and distribution frequencies over the years (histogram, panel C). Abbreviations: Alc – alcoholic solvents; H20 – water-based, aqueous solvents; Org – other organic solvents.

#### 3.4.14 BTS-related controls

Initially, solely w/o BTS was used in most cases as a BTS-related control. The number of additional controls (inactivated BTS, w/o cofactors) has increased over time (Fig. S20).

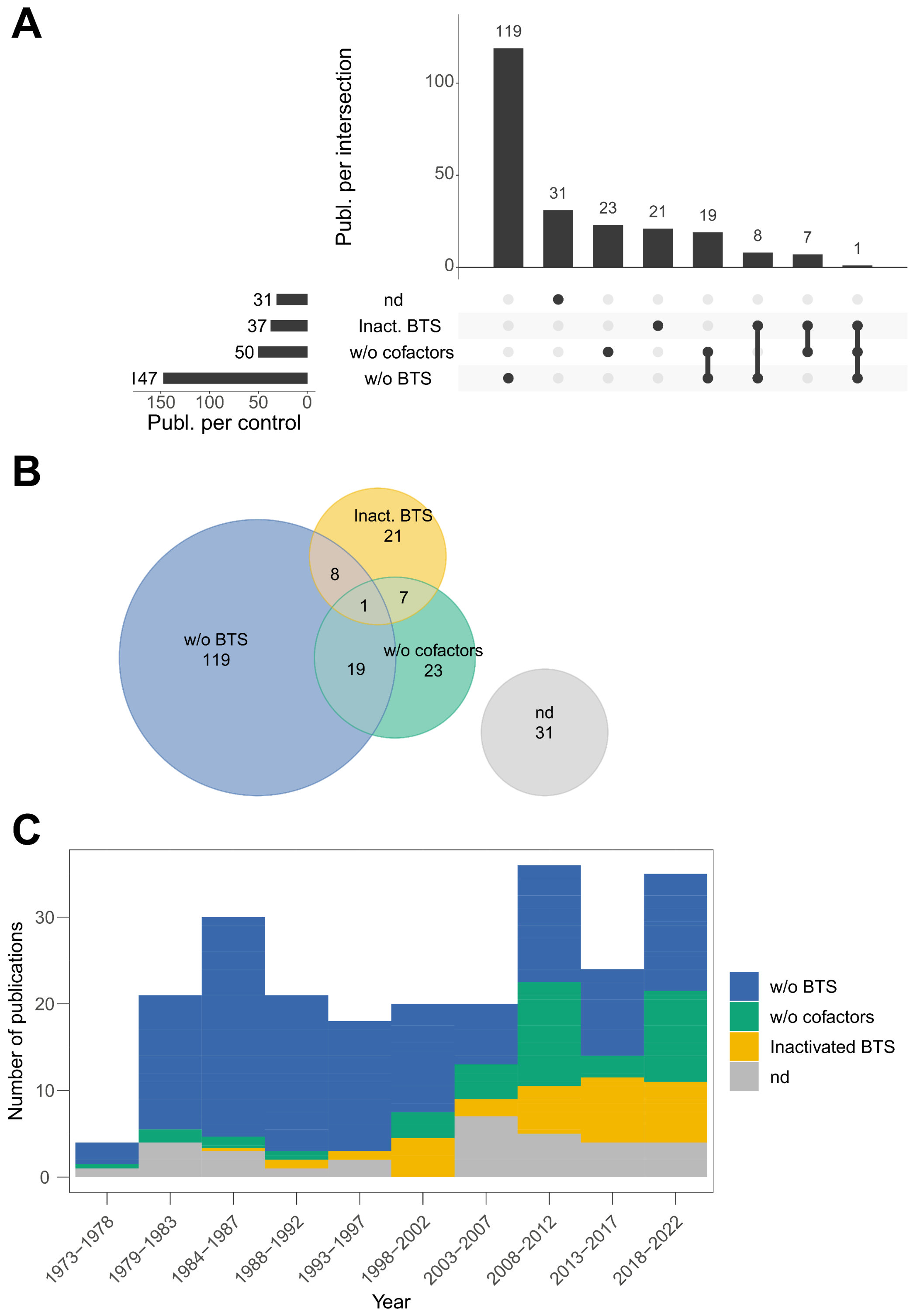

**Fig. S20:** Descriptive statistics of the data item subdomain "BTS-related controls", illustrated as interaction frequencies (upset plot, panel A), relation frequencies (Euler diagram, panel B), and distribution frequencies over the years (histogram, panel C).

3.5 Multiple correspondence analyses (MCA) of qualitative data item subdomains (outcome C) – additional results

The subdomain “strain” depicted a positive robustness tendency for studies utilising Wister-strain rats to produce BTS (Fig. S21D).

There are no clear emerging patterns for the subdomain “buffer system.” However, we can discern between Tris-buffered systems (slight positive tendency) and culture medium-buffered systems (slight negative tendency) (Fig. S22A).

No evident robustness markers could be derived from the subdomains “year of publication” (Fig. S23A), “field” (former: “journal”, Fig. S23B), and “type of BTS” (Fig. 23D). However, for “field” we can distinguish between studies allocated to “analytical chemistry” and “other biosciences”, which are rather positively connotated, and studies in the field comprising both nutritional science and toxicology, which are rather negatively connotated.

The supplementary variable “dataset” (BTS1 to BTS3) depicted no clear patterns. However, it is noticeable that the centroids of BTS1 and BTS3 align rather well, whereas BTS2/mutagen has a rather negative connotation (Fig. S24C).

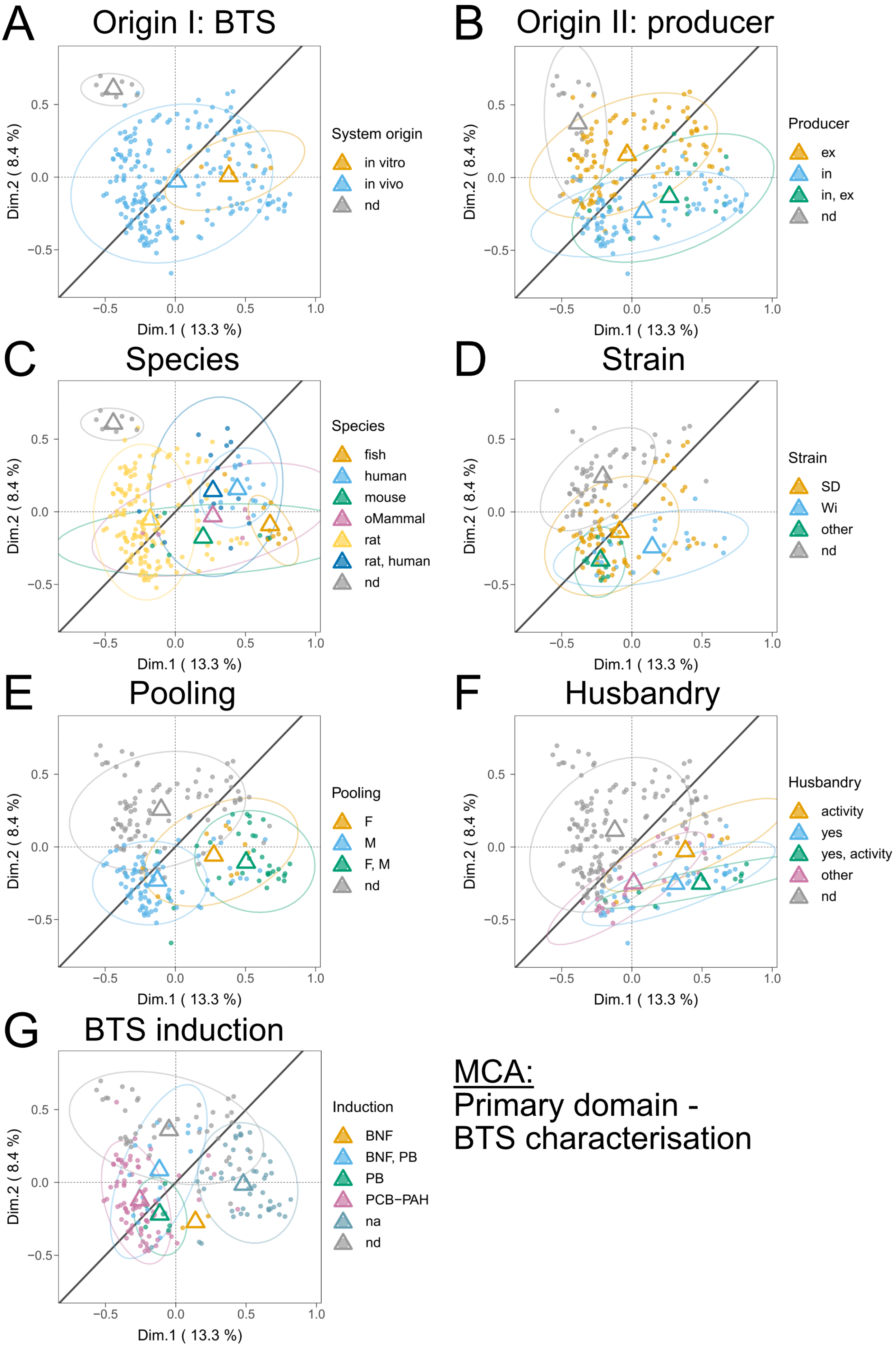
A
Origin I: BTS
System origin
in vitro
in vivo
nd
B
Origin II: producer
Producer
ex
in
in, ex
nd
C
Species
Species
fish
human
mouse
oMammal
rat
rat, human
nd
D
Strain
Strain
SD
Wi
other
nd
E
Pooling
Pooling
F
M
F, M
nd
F
Husbandry
Husbandry
activity
yes
yes, activity
other
nd
G
BTS induction
Induction
BNF
BNF, PB
PB
PCB−PAH
na
nd
Dim.1 ( 13.3 %)
Dim.2 ( 8.4 %)
−0.5
0.0
0.5
1.0
MCA:
Primary domain -
BTS characterisation

**Fig. S21**: Multiple Correspondence Analysis (MCA) plots showing the clustering of n = 229 publications. Depicted are clusters of the inquired primary data item domain BTS characterisation, subdomains “origin I: BTS” (A), “origin II: producer” (B), “species” (C), “strain” (rodent) (D), “pooling” (E), “husbandry” (F), and “BTS induction” (G). Centroids (respectively coloured triangles) mark the mean individual dimensional coordinates per category. Ellipses (coloured) represent normal probability contours at a 0.9 confidence level. The line of unity is given in dark grey. Abbreviations: nd – not defined; na – not applicable; ex – external; in – internal: oMammal – other Mammals; SD – Sprague-Dawley rat strain; Wi – Wistar rat strain; F – female; M – male; BNF - beta-Naphthoflavone; PB – phenobarbital; PCB-PAH - polychlorinated biphenyls and polycyclic aromatic hydrocarbons (note that Aroclors, other PCBs, and methylcholanthrene (PAH) were summarised in this category for MCA analysis to meet computation criteria, more details are given in the SM, section 2.12, Tab. S4, and SI9).

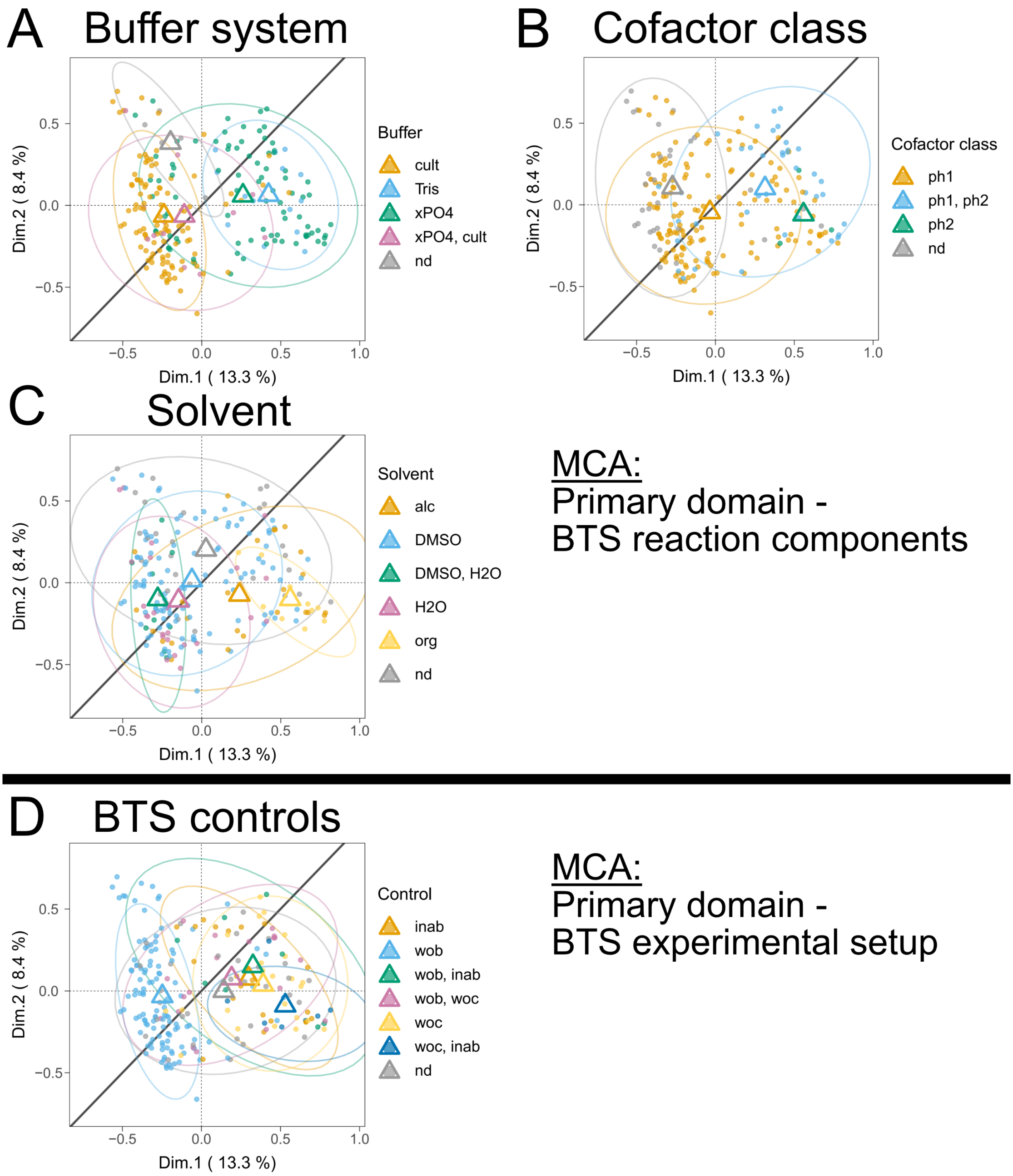

**Fig. S22**: Multiple Correspondence Analysis (MCA) plots showing the clustering of n = 229 publications. Depicted are clusters of the inquired primary data item domains of BTS reaction components and BTS experimental setup, subdomains “buffer system” (A), “cofactor class” (B), “solvent” (C), and “BTS controls” (D). Centroids (respectively coloured triangles)

mark the mean individual dimensional coordinates per category. Ellipses (coloured) represent normal probability contours at a 0.9 confidence level. The line of unity is given in dark grey. Abbreviations: nd – not defined; Cult – culture medium; xPO4 – various forms of phosphate buffers; ph1 – phase 1-related cofactors; ph2 – phase 2-related cofactors; Alc – alcoholic solvents; DMSO – dimethyl sulfoxide; H20 – water-based, aqueous solvents; Org – other organic solvents; inab – inactivated BTS; wob – without BTS; woc – without cofactors.

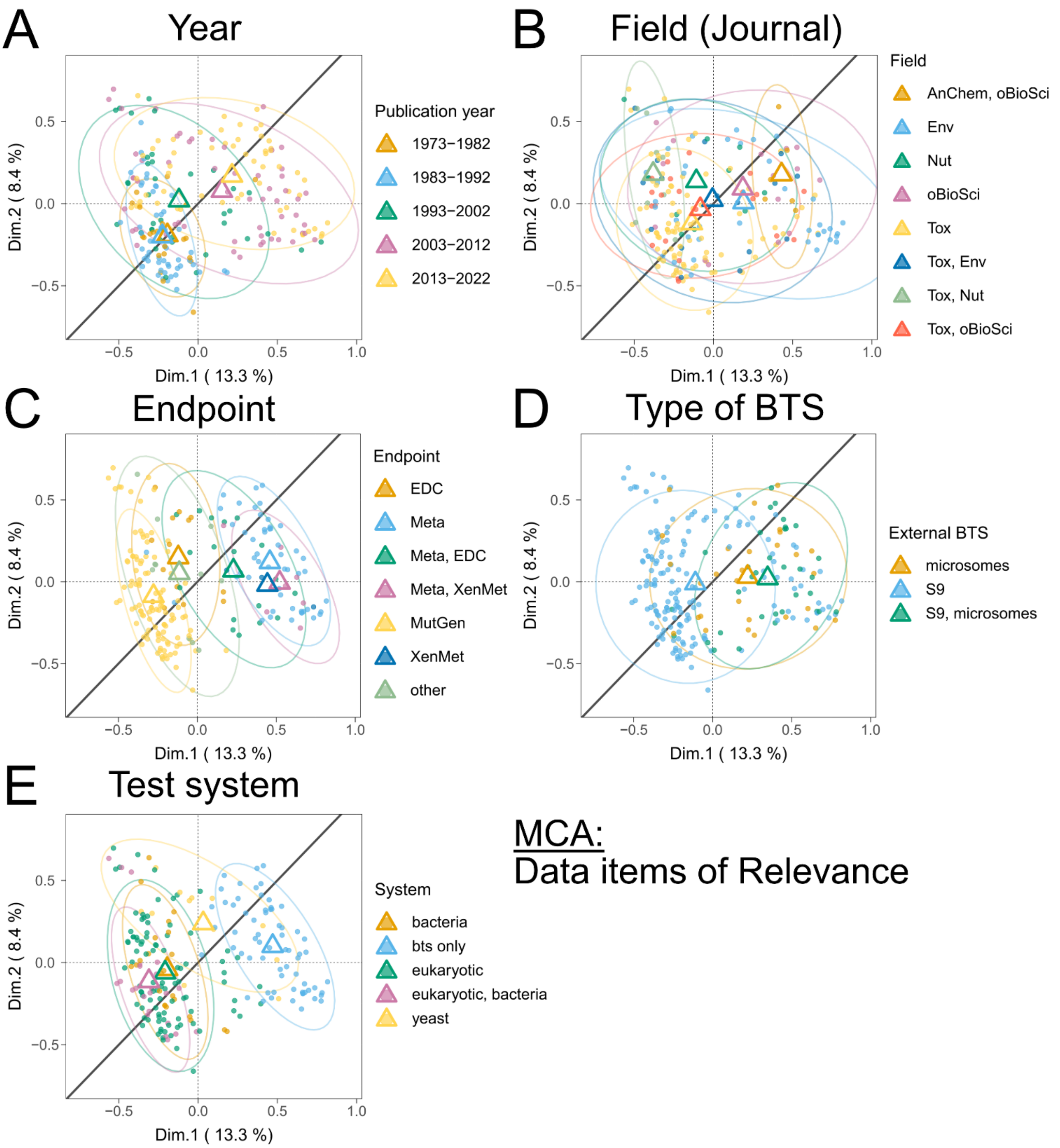

**Fig. S23**: Multiple Correspondence Analysis (MCA) plots showing the clustering of n = 229 publications. Depicted are clusters of the inquired data item domains of relevance “year” (A), “field” (journal) (B), “endpoint” (C), “type of BTS” (D), and “test system” (E). Centroids (respectively coloured triangles) mark the mean individual dimensional coordinates per category. Ellipses (coloured) represent normal probability contours at a 0.9 confidence level. The line of unity is given in dark grey. Abbreviations: AnChem – analytical chemistry; oBioSci – other biosciences; Env – environmental sciences and environmental toxicology; Nut – nutritional sciences; Tox – toxicology (classic, human); ; EDC – endocrine disruption; Meta – metabolites; MutGen – mutagenicity and genotoxicity; XenMet – xenobiotic metabolism functionality.

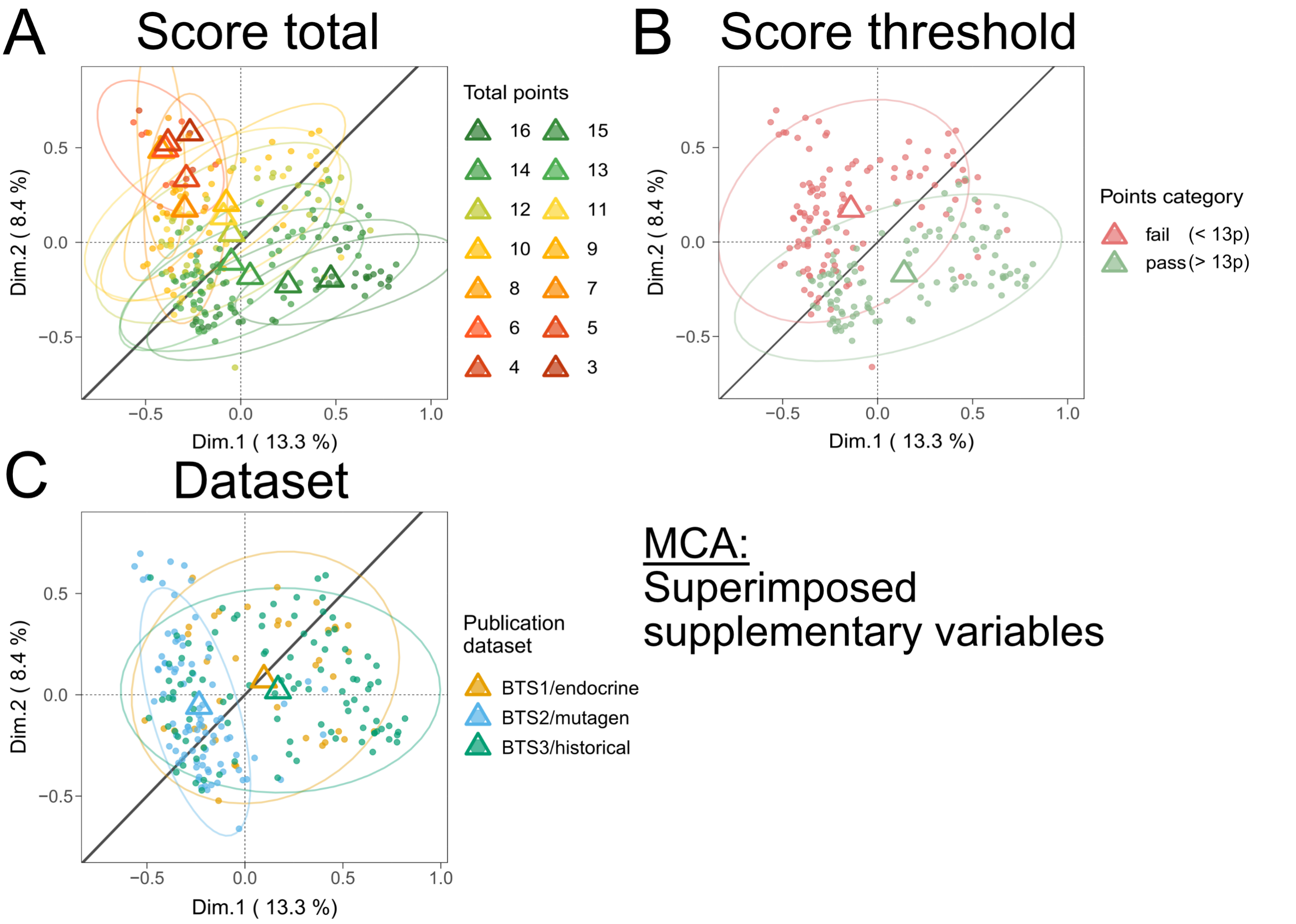

**Fig. S24**: Multiple Correspondence Analysis (MCA) plots showing the clustering of n = 229 publications. Depicted are clusters of the superimposed supplementary variables “total score” (A), “score threshold” (B), and “dataset” (C). Centroids (respectively coloured triangles) mark the mean individual dimensional coordinates per category. Ellipses (coloured) represent normal probability contours at a 0.9 confidence level. The line of unity is given in dark grey.

### 3.6 Data association rule mining and relational networks via *Apriori* algorithms (outcome C) – additional data

The supplementary information material folder SI10 provides additional interactive plots for 100, 200, 500, 1000, 2000, and 5000 association rules.

### 3.7 Follow-up, confirmatory analyses – additional results and data

**Table S9**: Summary of methodological robustness markers, as derived from MCA and *Apriori* analyses, and data item subdomain to data item measure listing.

| Qualitative data item subdomain | Qualitative data item measure | Analysis | Methodological robustness marker |
|---|---|---|---|
| Jornal/Field | “Tox” | Apriori | Negative |
| Jornal/Field | “Tox” + “Nut” | MCA | Negative (tendency) |
| Test system | “BTS only” | Apriori | Positive |
| Test system | “BTS only” | MCA | Positive (tendency) |
| Test system | “eukaryotic” | Apriori | Negative |
| Test system | “eukaryotic” | MCA | Negative (tendency) |

| | | | |
|---|---|---|---|
| Endpoint | "Meta" | MCA | Positive |
| Endpoint | "XenMet" | MCA | Positive |
| Endpoint | "Meta" | Apriori | Positive |
| Endpoint | "MutGen" | Apriori | Negative |
| Endpoint | "MutGen" | MCA | Negative (tendency) |
| External BTS (type) | "S9" | Apriori | Negative |
| BTS origin (producer) | "internal" | MCA | Positive |
| BTS origin (producer) | "external" | Apriori | Negative |
| Species | "human" | MCA | Positive |
| Species | "fish" | MCA | Positive |
| Species | "rat" | Apriori | Negative |
| Species system origin (redefined in simplification) | "in vitro" | MCA | Positive |
| Strain | "na" | Apriori | Positive |
| Strain | "Wistar" | MCA | Positive (tendency) |
| BTS pooling sex | "male + female" | MCA | Positive |
| BTS pooling sex | "na" | MCA | Positive |
| BTS induction | "na" | MCA | Positive |
| BTS induction | "na" | Apriori | Positive |
| BTS induction | "BNF/PB", "PCB-PAH" | MCA | Negative (tendency) |
| Buffer system | "xPO4" | Apriori | Positive |
| Buffer system | "Tris" | MCA | Positive (tendency) |
| Buffer system | "culture media" | Apriori | Negative |
| Buffer system | "culture media" | MCA | Negative (tendency) |
| Cofactor class (redefined in simplification) | "Phase 2" | MCA | Positive |
| Cofactor class (redefined in simplification) | "Phase 1 + Phase 2" | Apriori | Positive |
| Cofactor class (redefined in simplification) | "Phase 1" | Apriori | Negative |
| Solvent | "DMSO" | Apriori | Negative |
| Solvent | "DMSO" + "H2O" | MCA | Negative (tendency) |
| Solvent | "org" | MCA | Positive |
| BTS-related control | "w/o BTS" | Apriori | Negative |
| BTS-related control | "w/o BTS" | MCA | Negative (tendency) |
| BTS-related control | "w/o cofactor", "inactivated BTS" | MCA | Positive |
| Dataset (supplementary variable) | "BTS3" | Apriori | Positive |
| Dataset (supplementary variable) | "BTS2" | Apriori | Negative |

822

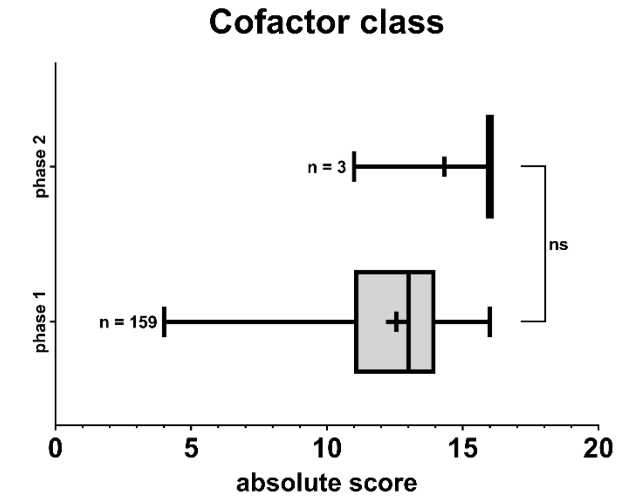

**Fig. S25**: Boxplots depicting absolute scoring populations of reviewed and assessed articles for respectively curated sub-datasets per qualitative data item subdomain. Whiskers indicate the populations' upper (max.) to lower (min.) boundaries. Boxes indicate the 75$^{th}$ and 25$^{th}$ percentile and the in-between line represents the median. Crosses represent mean population values. Statistical analysis of variance between datasets was conducted via two-sided Mann-Whitney U tests (pairwise comparison, alpha level = 0.05). Asterisks indicate statistically differing significance between means of respective datasets for pairwise comparisons (ns=non-significant). The number of included records per seminal population analysis is given in the graphs. More details are given in SI11.

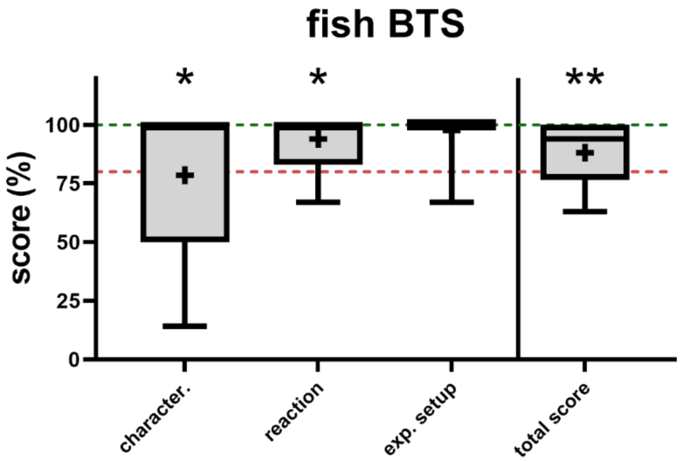

**Fig. S26**: Boxplots depicting relative scoring populations of reviewed and assessed articles utilising a fish-derived BTS (partly besides others in multi-species context, n = 17). Whiskers indicate the populations' upper (max.) to lower (min.) boundaries. Boxes indicate the 75th and 25th percentiles and the in-between line represents the median. Crosses represent mean population scores. The green dotted lines indicate full data reproducibility and robustness, and the red dotted lines represent a threshold of minimal quality acceptance (80%). Statistical significance towards the rest of the data set was tested in pairs by utilising two-sided Mann-Whitney-U tests (alpha level = 0.05). Asterisks indicate statistical significance (* $p < 0.05$, ** $p < 0.01$).

4. Supplementary discussion – additional material

**Table S10**: Summarised details from historical Ames *et al.* publications. Abbreviations: SD – Sprague-Dawley; m. – male; f. – female; hus. – husbandry; Cof. – cofactors; G6p – glucose-6-phosphate; dh – dehydrogenase; PB – phenobarbital; Aro – Aroclor.

| Ref. | Strain | Hus. | induction | Protein conc. | Cof. | Other | Mix | Ratio | Incubation |
|---|---|---|---|---|---|---|---|---|---|
| Ames et al. 1973 | SD | m., n=3, hus. in Garner et al. | PB | 30 % (v/v) | 4 mM NADP+ | 5 mM G6P, 8 mM MgCl, 33 mM KCl, 100 mM Na2PO4 | 2 mL agar, 0.1 mL bacteria, 0.1 mL compound, 0.5 mL S9 mix | 2.2 to 0.5 mL, 1 to 4.4 | 2 days, 37C |
| Ames et al. 1975 | SD | m., n=3, hus. in Garner et al. | Aro. | 4 – 10% (v/v),<br><br>Approx. 40 mg/mL | As above | As above | As above | As above | 2 days, 37C or 20 min preincubation with S9 for liquids |
| Maron & Ames 1983 | SD | m., n=3, hus. in Garner et al. | Aro. | 4 and 10% (v/v)<br><br>Approx. 40 mg/mL | As above | As above | As above | As above | 2 days, 37C Optional 20 min preincubation with S9 |
| Garner et al. 1972 | CD rats, CD1 mice, guinea pigs, hamster | f. + m.,n=3, hus. | PB | 5 – 40 mg "liver equivalents" in 3 mL | 0.5 mM NADP+ | 6.6 mM G6P, 0.3 U/mL G6P-dh, 8.3 mM MgCl2 33 mM KCl, 100 mM Na2PO4 | final | final | 20 min, 37C |

**Table S11**: BTS-related controls in the assessed literature for test setups using BTS in conjunction with eukaryotic or prokaryotic test systems

| Control category | Setup | Intention | Use |
|---|---|---|---|
| "w/o BTS" | Option A)<br>- Without BTS<br>- Without cofactors | Most common<br>- To distinguish between a biotransformation capable and not capable setup<br>- To ascertain if biotransformation is relevant for the investigated chemical within the practicable margins of the test setup<br>- To assess if biotransformation affects the bioactivity (within parallel *in vitro* assays) | - All *in vitro* assays<br>- Basically, the plain *in vitro* assay without any BTS |
| "w/o BTS" | Option B)<br>- Without BTS<br>- With cofactors | - To test assay performance when new methods are established<br>- To determine if the biotransformation reactions are predominantly driven by the added BTS, or if the addition of cofactors can stimulate any biotransformation activity in the used cell lines | - Development, characterisation, and optimisation of novel *in vitro* test system used in conjunction with BTS; including new methods or BTS adaptation to already established methods |

| | | | |
|---|---|---|---|
| | | - To detect the background signal potentially induced by added cofactors within technical readouts (relates to a blank containing culture medium or buffer without the test system) | |
| "w/o cofactors" | - With BTS<br>- Without cofactors | - Self-sustained biotransformation potential within BTS<br>- Microsomal and S9 fractions have low inherent quantities of cofactors present as residues from homogenisation<br>- To test assay performance when new methods are established<br>- To detect the background signal potentially induced by BTS within technical readouts (relates to a blank containing culture medium or buffer without the test system) | - Development, characterisation, and optimisation of novel *in vitro* test system used in conjunction with BTS; including new methods or BTS adaptation to already established methods<br>- Safeguard against non-intended phase 1 or 2 reactions |
| "inactivated BTS" | Option A)<br>- Heat or chemically inactivated BTS<br>- Without cofactors | - To account for bioavailability issues; additional BTS protein might act as a structural sink for hydrophobic/lipophilic chemicals | - All *in vitro* assays |
| "inactivated BTS" | Option B)<br>- Heat or chemically inactivated BTS<br>- With cofactors | - To determine if the biotransformation reactions are predominantly driven by the added BTS, or if the addition of cofactors can stimulate any biotransformation activity in the used cell lines | - Development, characterisation, and optimisation of novel *in vitro* test system used in conjunction with BTS; including new methods or BTS adaptation to already established methods.<br>- Only to be applied when option A is also included |

**Table S12:** List of BTS characterisation-related information gathered from BTS producers mentioned in the investigated literature. For rat S9, in specific.

| **Prod.** | **Species/strain/husbandry** | **Sterility** | **Induction** | **Protein conc.** | **Buffer** | **Activity** |
|---|---|---|---|---|---|---|
| A | Wistar; male; nd | nd | PB/BNF | 30.7 mg/mL | 0.05 M Tris, pH 7.4 | Mutagen. activity Aminoanthracene and Benzo(a)pyrene |
| B | Sprague Dawley; male; details on husbandry, protocol according to Maron & Ames 1983 | random sterility sampling of batches | PB/BNF | 38.4 mg/mL | 0.15 M KCl | EROD, PROD, MROD, BROD activity (fold-change); Mutagen. activity with ethidium bromide, cyclophosphamide, aminoanthracene, and benzo(a)pyrene |
| C | Sprague Dawley; male; pooled, nd | nd | na | 20 mg/mL | nd | sulfotransferase, CYP3A, and CYP2B activity (units/mg protein) |
| D | Sprague Dawley; ~24 -74 individuals pooled | nd | na | 20 mg/mL | 150 mM KCl, 50 mM Tris, 2 mM | sulfotransferase, CYP3A, CYP2B activity (pmol/mg min) |

| | | | | | | |
|---|---|---|---|---|---|---|
| | | | | | EDTA, pH 7.5 | |
| E | Sprague Dawley; 50 individuals pooled | nd | na | 20 mg/mL | nd | P450 content (nmol/mg protein) |
| F | Sprague Dawley; male; 50 individuals pooled; details on husbandry | nd | PB/BNF | 20 mg/mL | 50 mM Tris·HCl, 150 mM KCl, 2 mM EDTA | Cytochrome P450 content (nmol/mg protein), Cytochrome b5 content (nmol/mg protein) |
| G | Sprague Dawley; male; 6 individuals | nd | various | 21.3 mg/mL | 100mM Tris-HCl, 1mM EDTA, 250mM Sucrose, pH 7.4 | Cytochrome P450 content (nmol/mg protein), various phase 1 and 2 enzyme activity |
| H | costumer-related adjustments on strain/sex/pooling | nd | PB/BNF | ~20 mg/mL | nd | Cytochrome P450 content (nmol/mg protein), ECOD activity |
| I | Sprague Dawley; male; details on husbandry, protocol according to Maron & Ames 1983 | nd | Aroclor | nc | Maron & Ames 1983 | nc |
| J | no S9 or microsomes found on webpage | | | | | |
| K | Nothing specific found on webpage, probably only the supplier | | | | | |
| L | No S9 or microsomes found on webpage, only cofactors, S9 probably discontinued | | | | | |
| L | Information only in Japanese | | | | | |
| M | Could only find OECD test with S9 as a service provider | | | | | |
| N | could not find anything on webpage | | | | | |
| O | Discontinued, unclear | | | | | |
| P | Discontinued, unclear | | | | | |
| Q | defunct in 2003 | | | | | |
| R | defunct | | | | | |
| S | defunct | | | | | |

### 4.1 Study biases

Regarding study biases, we need to point out two aspects we identified throughout the assessment and analysis processes, which might potentially impact the interpretation of study outcomes.

First, one notable bias arises from the positive non-weighted scoring of non-applicable (“na”) data item measures. This issue emerged when it was not feasible to assign specific measures, such as the inability to induce human-derived BTS chemically. In such cases, the measure was designated as “na” and received a positive score. While this approach yields valid information for qualitative analyses, it inadvertently skews the scoring assessment favourably towards the respective record. This skewness introduces a potential positive bias in the scoring assessment, leading to an over-optimistic evaluation of methodological robustness. Despite its limitations, this scoring method was adopted due to the scientific controversy surrounding weighted scoring criteria (EFSA, 2010). Given that the overall body of literature did not meet the quality threshold, this approach was considered acceptable. A more refined scoring design would only further decrease the overall scores of all records, thereby not altering the fundamental interpretation of the data.

The second bias relates to the inclusion of the historical dataset BTS3. Analysis revealed that BTS3 consistently exhibited higher scores and more robust patterns in iterative analyses (Fig. 8 in the main article, Fig. S2, and Fig. S24C). Statistical differences were observed between BTS3 and BTS2 but not between BTS1 and BTS3 (Fig. S2). This discrepancy could be attributed to the prior familiarity of the authors with articles in BTS3, potentially reflecting their inherent quality and recognition in the field. However, the absence of statistical differences between BTS1 and BTS3, along with consistent associative patterns between these datasets, supports the feasibility of incorporating BTS3 in our analysis.

### 5. References

#### 5.1 References of the supplementary manuscript

Abdi, H., Valentin, D., 2007. Multiple correspondence analysis. Encyclopedia of measurement and statistics 2, 651–657.

Ampy, F.R., Asseffa, A., 1988. Regulatory effects of testosterone and 17 beta-oestradiol on the metabolism of dimethylnitrosamine by renal and hepatic microsomal enzymes from BALB/c mice. Cytobios 55, 87–94.

Armstrong, R., Hall, B.J., Doyle, J., Waters, E., 2011. “Scoping the scope” of a cochrane review. J Public Health (Bangkok) 33, 147–150. https://doi.org/10.1093/pubmed/fdr015

Campbell, M., McKenzie, J.E., Sowden, A., Katikireddi, S.V., Brennan, S.E., Ellis, S., Hartmann-Boyce, J., Ryan, R., Shepperd, S., Thomas, J., Welch, V., Thomson, H., 2020. Synthesis without meta-analysis (SWiM) in systematic reviews: Reporting guideline. The BMJ 368, 1–6. https://doi.org/10.1136/bmj.l6890

Chang, G., Jacobson-Kram, D., Williams, J.R., 1988. Use of an established human hepatoma cell line with endogenous bioactivation for gene mutation studies. Cell Biol Toxicol.

Chelcea, I., Örn, S., Hamers, T., Koekkoek, J., Legradi, J., Vogs, C., Andersson, P.L., 2022. Physiologically Based Toxicokinetic Modeling of Bisphenols in Zebrafish ( Danio rerio ) Accounting for Variations in Metabolic Rates, Brain Distribution, and Liver Accumulation. Environ Sci Technol 56, 10216–10228. https://doi.org/10.1021/acs.est.2c01292

Coecke, S., Ahr, H., Blaauboer, B.J., Bremer, S., Casati, S., Castell, J., Combes, R., Corvi, R., Crespi, C.L., Cunningham, M.L., Elaut, G., Eletti, B., Freidig, A., Gennari, A., Ghersi-Egea, J.-F., Guillouzo, A., Hartung, T., Hoet, P., Ingelman-Sundberg, M., Munn, S., Janssens, W., Ladstetter, B., Leahy, D., Long, A., Meneguz, A., Monshouwer, M., Morath, S., Nagelkerke, F., Pelkonen, O., Ponti, J., Prieto, P., Richert, L., Sabbioni, E., Schaack, B., Steiling, W., Testai, E., Vericat, J.-A., Worth, A., 2006. Metabolism: A Bottleneck in In Vitro Toxicological Test Development. Alternatives to Laboratory Animals 34, 49–84. https://doi.org/10.1177/026119290603400113

Combes, R.D., 2012. Cell Transformation Assays: Are we Barking up the Wrong Tree? Alternatives to Laboratory Animals 40, 115–130. https://doi.org/10.1177/026119291204000211

Felton, J.S., Bjeldanes, L.F., Hatch, F.T., 1984. Mutagens in cooked foods--metabolism and genetic toxicity. Adv Exp Med Biol.

Food, E., Authority, S., 2010. Application of systematic review methodology to food and feed safety assessments to support decision making. EFSA Journal 8. https://doi.org/10.2903/j.efsa.2010.1637

Galloway, S.M., Aardema, M.J., Ishidate, M., Ivett, J.L., Kirkland, D.J., Morita, T., Mosesso, P., Sofuni, T., 1994. Report from working group on in vitro tests for chromosomal aberrations. Mutat Res.

Gehlenborg, N., 2019. UpSetR: A More Scalable Alternative to Venn and Euler Diagrams for Visualizing Intersecting Sets.

Gouliarmou, V., Lostia, A.M., Coecke, S., Bernasconi, C., Bessems, J., Dorne, J. Lou, Ferguson, S., Testai, E., Remy, U.G., Brian Houston, J., Monshouwer, M., Nong, A., Pelkonen, O., Morath, S., Wetmore, B.A., Worth, A., Zanelli, U., Zorzoli, M.C., Whelan, M., 2018. Establishing a systematic framework to characterise in vitro methods for human hepatic metabolic clearance. Toxicology in Vitro 53, 233–244. https://doi.org/10.1016/j.tiv.2018.08.004

Hahsler, M., Buchta, C., Gruen, B., Hornik, K., 2023. arules: Mining Association Rules and Frequent Itemsets.

Hair, J.F., Black, W.C., Babin, B.J., Anderson, R.E., 2019. Multivariate Data Analysis. Cengage.

Harding, C., Viljanto, M., Habershon-Butcher, J., Taylor, P., Scarth, J., 2023. Equine metabolism of the selective androgen receptor modulator YK-11 in urine and plasma following oral administration. Drug Test Anal 15, 388–407. https://doi.org/10.1002/dta.3425

Hashler, M., 2023. arulesViz: Visualizing Association Rules and Frequent Itemsets.

Husson, F., Le, S., Pagès, J., 2017. Exploratory Multivariate Analysis by Example Using R. Chapman and Hall/CRC. https://doi.org/10.1201/b21874

Jacobs, M., 2013. In vitro metabolism and bioavailability tests for endocrine active substances: What is needed next for regulatory purposes? ALTEX 30, 331–351. https://doi.org/10.14573/altex.2013.3.331

Jacobs, M., Janssens, W., Bernauer, U., Brandon, E., Coecke, S., Combes, R., Edwards, P., Freidig, A., Freyberger, A., Kolanczyk, R., Mc Ardle, C., Mekenyan, O., Schmieder, P., Schrader, T., Takeyoshi, M., Burg, B., 2008. The Use of Metabolising Systems for In Vitro Testing of Endocrine Disruptors. Curr Drug Metab 9, 796–826. https://doi.org/10.2174/138920008786049294

James, K.L., Randall, N.P., Haddaway, N.R., 2016. A methodology for systematic mapping in environmental sciences. Environ Evid 5, 7. https://doi.org/10.1186/s13750-016-0059-6

Khalil, H., Tricco, A.C., 2022. Differentiating between mapping reviews and scoping reviews in the evidence synthesis ecosystem. J Clin Epidemiol 149, 175–182. https://doi.org/10.1016/j.jclinepi.2022.05.012

Klimisch, H.-J., Andreae, M., Tillmann, U., 1997. A Systematic Approach for Evaluating the Quality of Experimental Toxicological and Ecotoxicological Data. Regulatory Toxicology and Pharmacology 25, 1–5. https://doi.org/10.1006/rtph.1996.1076

Larsson, J., 2022. eulerr: Area-Proportional Euler and Venn Diagrams with Ellipses.

Lê, S., Josse, J., Husson, F., 2008. FactoMineR : An R Package for Multivariate Analysis. J Stat Softw 25, 253–258. https://doi.org/10.18637/jss.v025.i01

Liu, D., Gao, J., Zhang, C., Ren, X., Liu, Y., Xu, Y., 2011. Identification of carboxylesterases expressed in rat intestine and effects of their hydrolyzing activity in predicting first-pass metabolism of ester prodrugs. Pharmazie 66, 888–893.

Moermond, C.T.A., Kase, R., Korkaric, M., Ågerstrand, M., 2016. CRED: Criteria for reporting and evaluating ecotoxicity data. Environ Toxicol Chem 35, 1297–1309. https://doi.org/10.1002/etc.3259

Moher, D., Shamseer, L., Clarke, M., Ghersi, D., Liberati, A., Petticrew, M., Shekelle, P., Stewart, L.A., 2015. Preferred reporting items for systematic review and meta-analysis protocols (PRISMA-P) 2015 statement. Syst Rev 4, 1. https://doi.org/10.1186/2046-4053-4-1

Morgan, R.L., Thayer, K.A., Santesso, N., Holloway, A.C., Blain, R., Eftim, S.E., Goldstone, A.E., Ross, P., Ansari, M., Akl, E.A., Filippini, T., Hansell, A., Meerpohl, J.J., Mustafa, R.A., Verbeek, J., Vinceti, M., Whaley, P., Schünemann, H.J., 2019. A risk of bias instrument for non-randomized studies of exposures: A users' guide to its application in the context of GRADE. Environ Int 122, 168–184. https://doi.org/10.1016/j.envint.2018.11.004

Munn, Z., Peters, M.D.J., Stern, C., Tufanaru, C., McArthur, A., Aromataris, E., 2018. Systematic review or scoping review? Guidance for authors when choosing between a systematic or scoping review approach. BMC Med Res Methodol 18, 143. https://doi.org/10.1186/s12874-018-0611-x

NTP-OHAT, 2019. Handbook for Conducting a Literature-Based Health Assessment Using OHAT Approach for Systematic Review and Evidence Integration, National Toxicology Program.

OECD, 2018. Revised Guidance Document 150 on Standardised Test Guidelines for Evaluating Chemicals for Endocrine Disruption, OECD Publishing, OECD Series on Testing and Assessment. OECD. https://doi.org/10.1787/9789264304741-en

OECD, 2008. Detailed Review Paper on the State of the Science on Novel In Vitro and In Vivo Screening and Testing Methods and Endpoints for Evaluating Endocrine Disruptors, SERIES ON TESTING AND ASSESMENT, OECD Series on Testing and Assessment. OECD. https://doi.org/10.1787/9789264221352-en

Page, M.J., McKenzie, J.E., Bossuyt, P.M., Boutron, I., Hoffmann, T.C., Mulrow, C.D., Shamseer, L., Tetzlaff, J.M., Akl, E.A., Brennan, S.E., Chou, R., Glanville, J., Grimshaw, J.M., Hróbjartsson, A., Lalu, M.M., Li, T., Loder, E.W., Mayo-Wilson, E., McDonald, S., McGuinness, L.A., Stewart, L.A., Thomas, J., Tricco, A.C., Welch, V.A., Whiting, P., Moher, D., 2021a. The PRISMA 2020 statement: an updated guideline for reporting systematic reviews. BMJ 372, n71. https://doi.org/10.1136/bmj.n71

Page, M.J., Moher, D., Bossuyt, P.M., Boutron, I., Hoffmann, T.C., Mulrow, C.D., Shamseer, L., Tetzlaff, J.M., Akl, E.A., Brennan, S.E., Chou, R., Glanville, J., Grimshaw, J.M., Hróbjartsson, A., Lalu, M.M., Li, T., Loder, E.W., Mayo-Wilson, E., McDonald, S., McGuinness, L.A., Stewart, L.A., Thomas, J., Tricco, A.C., Welch, V.A., Whiting, P., McKenzie, J.E., 2021b. PRISMA 2020 explanation and elaboration: updated guidance and exemplars for reporting systematic reviews. BMJ 372, n160. https://doi.org/10.1136/bmj.n160

Posit team, 2023. RStudio: Integrated Development Environment for R.

Rooney, A.A., Cooper, G.S., Jahnke, G.D., Lam, J., Morgan, R.L., Boyles, A.L., Ratcliffe, J.M., Kraft, A.D., Schünemann, H.J., Schwingl, P., Walker, T.D., Thayer, K.A., Lunn, R.M., 2016. How credible are the study results? Evaluating and applying internal validity tools to literature-based assessments of environmental health hazards. Environ Int 92–93, 617–629. https://doi.org/10.1016/j.envint.2016.01.005

Schneider, K., Schwarz, M., Burkholder, I., Kopp-Schneider, A., Edler, L., Kinsner-Ovaskainen, A., Hartung, T., Hoffmann, S., 2009. "ToxRTool", a new tool to assess the reliability of toxicological data. Toxicol Lett 189, 138–144. https://doi.org/10.1016/j.toxlet.2009.05.013

Shamseer, L., Moher, D., Clarke, M., Ghersi, D., Liberati, A., Petticrew, M., Shekelle, P., Stewart, L.A., 2015. Preferred reporting items for systematic review and meta-analysis protocols (PRISMA-P) 2015: elaboration and explanation. BMJ 349, g7647–g7647. https://doi.org/10.1136/bmj.g7647

Siegers, C.P., Denker, S., Steffen, B., Jelkmann, W., 1987. Biotransformation enzymes in two renal epithelial cell lines (LLC-PK1 and RK-L). Mol Toxicol.

Team, R.C., 2023. R: A language and environment for statistical computing.

Tricco, A.C., Lillie, E., Zarin, W., O'Brien, K.K., Colquhoun, H., Levac, D., Moher, D., Peters, M.D.J., Horsley, T., Weeks, L., Hempel, S., Akl, E.A., Chang, C., McGowan, J., Stewart, L., Hartling, L., Aldcroft, A., Wilson, M.G., Garritty, C., Lewin, S., Godfrey, C.M., Macdonald, M.T., Langlois, E. V., Soares-Weiser, K., Moriarty, J., Clifford, T., Tunçalp, Ö., Straus, S.E., 2018. PRISMA Extension for Scoping Reviews (PRISMA-ScR): Checklist and Explanation. Ann Intern Med 169, 467–473. https://doi.org/10.7326/M18-0850

US EPA, 2018. Application of Systematic Review in Tsca Risk Evaluations.

Whaley, P., Aiassa, E., Beausoleil, C., Beronius, A., Bilotta, G., Boobis, A., de Vries, R., Hanberg, A., Hoffmann, S., Hunt, N., Kwiatkowski, C.F., Lam, J., Lipworth, S., Martin, O., Randall, N., Rhomberg, L., Rooney, A.A., Schünemann, H.J., Wikoff, D., Wolffe, T., Halsall, C., 2020. Recommendations for the conduct of systematic reviews in toxicology and environmental health research (COSTER). Environ Int 143, 105926. https://doi.org/10.1016/j.envint.2020.105926

Wickham, H., 2016. ggplot2: Elegant Graphics for Data Analysis. Springer New York, New York.

Wolffe, T.A.M., Vidler, J., Halsall, C., Hunt, N., Whaley, P., 2020. A Survey of Systematic Evidence Mapping Practice and the Case for Knowledge Graphs in Environmental Health and Toxicology. Toxicological Sciences 175, 35–49. https://doi.org/10.1093/toxsci/kfaa025

Wolffe, T.A.M., Whaley, P., Halsall, C., Rooney, A.A., Walker, V.R., 2019. Systematic evidence maps as a novel tool to support evidence-based decision-making in chemicals policy and risk management. Environ Int 130, 104871. https://doi.org/10.1016/j.envint.2019.05.065

Zhang, Y., Zhang, Q., Ji, G., Xu, H., Zhang, S., Liu, J., Shi, L., 2016. In vitro metabolism of Hydroxylation polybrominated diphenyl ethers in mice liver. Huanjing Kexue Xuebao/Acta Scientiae Circumstantiae. https://doi.org/10.13671/j.hjkxxb.2016.0170

### 5.2 References of the dataset BTS1/endocrine

Allaben, W.T., Louie, S.C., Lazear, E.J., 1979. Synergistic effect of diethylstilbestrol on the mutagenicity of 2-acetylaminofluorene and N-hydroxy-acetylaminofluorene in the Salmonella assay system. Cancer Lett. 7, 109–114. https://doi.org/10.1016/S0304-3835(79)80104-4

Anand, S.S., Serex, T.L., Carpenter, C., Donner, E.M., Hoke, R., Buck, R.C., Loveless, S.E., 2012. Toxicological assessment of tridecafluorohexylethyl methacrylate (6:2 FTMAC). Toxicology 292, 42–52. https://doi.org/10.1016/j.tox.2011.11.016

Beyer, B.K., Juchau, M.R., 1988. Contrasting effects of estradiol-17β and 17α-ethinyl estradiol-17β on cultured whole embryos. J. Steroid Biochem. 29, 629–634. https://doi.org/10.1016/0022-4731(88)90162-8

Borrisser-Pairó, F., Rasmussen, M.K., Ekstrand, B., Zamaratskaia, G., 2015. Gender-related differences in the formation of skatole metabolites by specific CYP450 in porcine hepatic S9 fractions. Animal 9, 635–642. https://doi.org/10.1017/S1751731114002808

Brimer, P.A., Tan, E.-L., Hsie, A.W., 1982. Effect of metabolic activation on the cytotoxicity and mutagenicity of 1,2-dibromoethane in the CHO/HGPRT system. Mutat. Res. Mol. Mech. Mutagen. 95, 377–388. https://doi.org/10.1016/0027-5107(82)90272-X

Broberg, M.N., Knych, H., Bondesson, U., Pettersson, C., Stanley, S., Thevis, M., Hedeland, M., 2021. Investigation of Equine In Vivo and In Vitro Derived Metabolites of the Selective Androgen Receptor Modulator (SARM) ACP-105 for Improved Doping Control. Metabolites 11, 85. https://doi.org/10.3390/metabo11020085

Cabaton, N., Zalko, D., Rathahao, E., Canlet, C., Delous, G., Chagnon, M.-C., Cravedi, J.-P., Perdu, E., 2008. Biotransformation of bisphenol F by human and rat liver subcellular fractions. Toxicol. Vitr. 22, 1697–1704. https://doi.org/10.1016/j.tiv.2008.07.004

Clarke, A., Scarth, J., Teale, P., Pearce, C., Hillyer, L., 2011. The use of in vitro technologies and high-resolution/accurate-mass LC-MS to screen for metabolites of 'designer' steroids in the equine. Drug Test. Anal. 3, 74–87. https://doi.org/10.1002/dta.250

Fahrig, R., 1996. Anti-mutagenic agents are also co-recombinogenic and can be converted into co-mutagens. Mutat. Res. Mol. Mech. Mutagen. 350, 59–67. https://doi.org/10.1016/0027-5107(95)00091-7

Glatt, H., Jung, R., Oesch, F., 1983. Bacterial mutagenicity investigation of epoxides: drugs, drug metabolites, steroids and pesticides. Mutat. Res. - Fundam. Mol. Mech. Mutagen. 111, 99–118. https://doi.org/10.1016/0027-5107(83)90056-8

Hashimoto, S., Ueda, Y., Kurihara, R., Shiraishi, F., 2007. Comparison of the estrogenic activities of seawater extracts from Suruga Bay, Japan, based on chemical analysis or bioassay. Environ. Toxicol. Chem. 26, 279–286. https://doi.org/10.1897/05-689R1.1

Hundal, B.S., Dhillon, V.S., Sidhu, I.S., 1997. Genotoxic potential of estrogens. Mutat. Res. Toxicol. Environ. Mutagen. 389, 173–181. https://doi.org/10.1016/S1383-5718(96)00144-1

HUO, Z.P., FENG, X.C., WANG, Y., TIAN, Y.T., QIU, F., 2021. Sulfite as the substrate of C-sulfonate metabolism of α, β-unsaturated carbonyl containing andrographolide: analysis of sulfite in rats' intestinal tract and the reaction kinetics of andrographolide with sulfite. Chin. J. Nat. Med. 19, 706–712. https://doi.org/10.1016/S1875-5364(21)60094-8

Jeon, B.K., Jang, Y., Lee, E.M., Jung, D.W., Moon, J.H., Lee, H.J., Lee, D.Y., 2021. A systematic approach to metabolic characterisation of thyroid-disrupting chemicals and their in vitro biotransformants based on prediction-assisted metabolomic analysis. J. Chromatogr. A 1649, 462222. https://doi.org/10.1016/j.chroma.2021.462222

Kang, J.S., Choi, J.-S., Kim, W.-K., Lee, Y.-J., Park, J.-W., 2014. Estrogenic potency of bisphenol S, polyethersulfone and their metabolites generated by the rat liver S9 fractions on a MVLN cell using a luciferase reporter gene assay. Reprod. Biol. Endocrinol. 12, 102. https://doi.org/10.1186/1477-7827-12-102

Kojima, M., Fukunaga, K., Sasaki, M., Nakamura, M., Tsuji, M., Nishiyama, T., 2005. Evaluation of estrogenic activities of pesticides using an in vitro reporter gene assay. Int. J. Environ. Health Res. 15, 271–280. https://doi.org/10.1080/09603120500155765

Lakhani, N.J., Sparreboom, A., Xu, X. i. a., Veenstra, T.D., Venitz, J., Dahut, W.L., Figg, W.D., 2007. Characterisation of in vitro and in vivo metabolic pathways of the investigational anticancer agent, 2-methoxyestradiol. J. Pharm. Sci. 96, 1821–1831. https://doi.org/10.1002/jps.20837

Li, M., Yang, Yunjia, Yang, Yi, Yin, J., Zhang, J., Feng, Y., Shao, B., 2013. Biotransformation of Bisphenol AF to Its Major Glucuronide Metabolite Reduces Estrogenic Activity. PLoS One 8, e83170. https://doi.org/10.1371/journal.pone.0083170

Lindblad, W.J., Jackim, E., 1982. Mechanism for the differential induction of mutation by S9 activated benzo[a]pyrene employing either a glucose-6-phosphate-dependent NADPH-regenerating system or an isocitrate-dependent system. Mutat. Res. Mol. Mech. Mutagen. 96, 109–118. https://doi.org/10.1016/0027-5107(82)90021-5

Mollergues, J., Van Vugt-Lussenburg, B., Kirchnawy, C., Bandi, R.A., Van Der Lee, R.B., Marin-Kuan, M., Schilter, B., Fussell, K.C., 2017. Incorporation of a metabolising system in biodetection assays for endocrine active substances. ALTEX 34, 389–398. https://doi.org/10.14573/altex.1611021

Montaño, M., Weiss, J., Hoffmann, L., Gutleb, A.C., Murk, A.J., 2013. Metabolic Activation of Nonpolar Sediment Extracts Results in Enhanced Thyroid Hormone Disrupting Potency. Environ. Sci. Technol. 130716143653008. https://doi.org/10.1021/es4011898

Morrison, R.D., Blobaum, A.L., Byers, F.W., Santomango, T.S., Bridges, T.M., Stec, D., Brewer, K.A., Sanchez-Ponce, R., Corlew, M.M., Rush, R., Felts, A.S., Manka, J., Bates, B.S., Venable, D.F., Rodriguez, A.L., Jones, C.K., Niswender, C.M., Conn, P.J., Lindsley, C.W., Emmitte, K.A., Daniels, J.S., 2012. The role of aldehyde oxidase and xanthine oxidase in the biotransformation of a novel negative allosteric modulator of metabotropic glutamate receptor subtype 5. Drug Metab. Dispos. 40, 1834–1845. https://doi.org/10.1124/dmd.112.046136

Mugford, C.A., Tarloff, J.B., 1997. The contribution of oxidation and deacetylation to acetaminophen nephrotoxicity in female Sprague-Dawley rats. Toxicol. Lett. 93, 15–22. https://doi.org/10.1016/S0378-4274(97)00063-5

Myhr, B.C., Mayo, J.K., 1987. Mutagenicity of rat-liver S9 to L5178Y mouse lymphoma cells. Mutat. Res. Toxicol. 189, 27–37. https://doi.org/10.1016/0165-1218(87)90030-9

Okuda, K., Fukuuchi, T., Takiguchi, M., Yoshihara, S., 2011. Novel Pathway of Metabolic Activation of Bisphenol A-Related Compounds for Estrogenic Activity. Drug Metab. Dispos. 39, 1696–1703. https://doi.org/10.1124/dmd.111.040121

Ousji, O., Ohlund, L., Sleno, L., 2020. Comprehensive In Vitro Metabolism Study of Bisphenol A Using Liquid Chromatography-High Resolution Tandem Mass Spectrometry. Chem. Res. Toxicol. 33, 1468–1477. https://doi.org/10.1021/acs.chemrestox.0c00042

OZAWA, N., WATABE, T., YOSHIMURA, H., KOGA, N., SHUDO, K., 1985. Effect of liver S9 from 3,4,5,3',4'-pentachlorobiphenyl-pretreated rats on the mutagenic activity of the various carcinogens toward Salmonella typhimurium TA 98. J. Pharmacobiodyn. 8, 199–205. https://doi.org/10.1248/bpb1978.8.199

Park, H.S., Oh, J.U.H., Lee, J.H., Lee, Y.J., 2011. Minor effects of the Citrus flavonoids naringin, naringenin and quercetin, on the pharmacokinetics of doxorubicin in rats. Pharmazie 66, 424–429. https://doi.org/10.1691/ph.2011.0857

Parrella, A., Lavorgna, M., Criscuolo, E., Isidori, M., 2013. Mutagenicity, Genotoxicity, and Estrogenic Activity of River Porewaters. Arch. Environ. Contam. Toxicol. 65, 407–420. https://doi.org/10.1007/s00244-013-9928-y

Peng, B., Zhao, H., Keerthisinghe, T.P., Yu, Y., Chen, D., Huang, Y., Fang, M., 2022. Gut microbial metabolite p-cresol alters biotransformation of bisphenol A: Enzyme competition or gene induction? J. Hazard. Mater. 426, 128093. https://doi.org/10.1016/j.jhazmat.2021.128093

Sumida, K., Ooe, N., Nagahori, H., Saito, K., Isobe, N., Kaneko, H., Nakatsuka, I., 2001. An in Vitro Reporter Gene Assay Method Incorporating Metabolic Activation with Human and Rat S9 or Liver Microsomes. Biochem. Biophys. Res. Commun. 280, 85–91. https://doi.org/10.1006/bbrc.2000.4071

Taxvig, C., Olesen, P.T., Nellemann, C., 2011. Use of external metabolising systems when testing for endocrine disruption in the T-screen assay. Toxicol. Appl. Pharmacol. 250, 263–269. https://doi.org/10.1016/j.taap.2010.10.029

UENO, Y., TASHIRO, F., 1981. α-Zearalenol, a Major Hepatic Metabolite in Rats of Zearalenone, an Estrogenic Mycotoxin of Fusarium Species1. J. Biochem. 89, 563–571. https://doi.org/10.1093/oxfordjournals.jbchem.a133232

Vian, L., Bichet, N., Gouy, D., 1993. The in vitro micronucleus test on isolated human lymphocytes. Mutat. Res. Mutagen. Relat. Subj. 291, 93–102. https://doi.org/10.1016/0165-1161(93)90021-Q

Wang, L., Raghavan, N., He, K., Luettgen, J.M., Humphreys, W.G., Knabb, R.M., Pinto, D.J., Zhang, D., 2009. Sulfation of O -Demethyl Apixaban: Enzyme Identification and Species Comparison. Drug Metab. Dispos. 37, 802–808. https://doi.org/10.1124/dmd.108.025593

Wheeler, W.J., Cherry, L.M., Downs, T., Hsu, T.C., 1986. Mitotic inhibition and aneuploidy induction by naturally occurring and synthetic estrogens in Chinese hamster cells in vitro. Mutat. Res. Toxicol. 171, 31–41. https://doi.org/10.1016/0165-1218(86)90006-6

Yoshihara, S. ’i., 2001. Metabolic Activation of Bisphenol A by Rat Liver S9 Fraction. Toxicol. Sci. 62, 221–227. https://doi.org/10.1093/toxsci/62.2.221

Zalko, D., Prouillac, C., Riu, A., Perdu, E., Dolo, L., Jouanin, I., Canlet, C., Debrauwer, L., Cravedi, J.-P., 2006. Biotransformation of the flame retardant tetrabromo-bisphenol A by human and rat sub-cellular liver fractions. Chemosphere 64, 318–327. https://doi.org/10.1016/j.chemosphere.2005.12.053

Zhu, W., Xu, H., Wang, S.W.J., Hu, M., 2010. Breast Cancer Resistance Protein (BCRP) and Sulfotransferases Contribute Significantly to the Disposition of Genistein in Mouse Intestine. AAPS J. 12, 525–536. https://doi.org/10.1208/s12248-010-9209-x

5.3 References of the dataset BTS2/mutagen

Agarwal, D.K., Lawrence, W.H., Nunez, L.J., Autian, J., 1985. Mutagenicity evaluation of phthalic acid esters and metabolites in salmonella typhimurium cultures. J. Toxicol. Environ. Health 16, 61–69. https://doi.org/10.1080/15287398509530719

Amacher, D.E., Turner, G.N., 1980. Promutagen activation by rodent-liver postmitochondrial fractions in the L5178Y/TK cell mutation assay. Mutat. Res. Mutagen. Relat. Subj. 74, 485–501. https://doi.org/10.1016/0165-1161(80)90179-X

Arbillaga, L., Azqueta, A., Ezpeleta, O., Cerain, A.L. d., 2006. Oxidative DNA damage induced by Ochratoxin A in the HK-2 human kidney cell line: evidence of the relationship with cytotoxicity. Mutagenesis 22, 35–42. https://doi.org/10.1093/mutage/gel049

Ashby, J., Tinwell, H., Callander, R.D., Kimber, I., Clay, P., Galloway, S.M., Hill, R.B., Greenwood, S.K., Gaulden, M.E., Ferguson, M.J., Vogel, E., Nivard, M., Parry, J.M., Williamson, J., 1997. Thalidomide: lack of mutagenic activity across phyla and genetic endpoints. Mutat. Res. Mol. Mech. Mutagen. 396, 45–64. https://doi.org/10.1016/S0027-5107(97)00174-7

Babich, H., Borenfreund, E., 1987. Polycyclic aromatic hydrocarbon in vitro cytotoxicity to bluegill BF-2 cells: Mediation by S-9 microsomal fraction and temperature. Toxicol. Lett. 36, 107–116. https://doi.org/10.1016/0378-4274(87)90174-3

Benford, D.J., Reavy, H.J., Hubbard, S.A., 1988. Metabolising systems in cell culture cytotoxicity tests. Xenobiotica 18, 649–656. https://doi.org/10.3109/00498258809041703

Boeira, J.M., Da Silva, J., Erdtmann, B., Henriques, J.A.P., 2001. Genotoxic Effects of the Alkaloids Harman and Harmine Assessed by Comet Assay and Chromosome Aberration Test in Mammalian Cells in vitro. Pharmacol. Toxicol. 89, 287–294. https://doi.org/10.1034/j.1600-0773.2001.d01-162.x

Budroe, J.D., Schol, H.M., Shaddock, J.G., Casciano, D.A., 1988. Inhibition of 7,12-dimethylbenz[ a ]anthracene-induced genotoxicity in Chinese hamster ovary cells by retinol and retinoic acid. Carcinogenesis 9, 1307–1311. https://doi.org/10.1093/carcin/9.7.1307

Cabrera, M., Lavaggi, M.L., Hernández, P., Merlino, A., Gerpe, A., Porcal, W., Boiani, M., Ferreira, A., Monge, A., de Cerain, A.L., González, M., Cerecetto, H., 2009. Cytotoxic, mutagenic and genotoxic effects of new anti-T. cruzi 5-phenylethenylbenzofuroxans. Contribution of phase I metabolites on the mutagenicity induction. Toxicol. Lett. 190, 140–149. https://doi.org/10.1016/j.toxlet.2009.07.006

Chang, L.W., Daniel, F.B., Deangelo, A.B., 1991. DNA strand breaks induced in cultured human and rodent cells by chlorohydroxyfuranones—mutagens isolated from drinking water. Teratog. Carcinog. Mutagen. 11, 103–114. https://doi.org/10.1002/tcm.1770110206

Chen, D.J.-C., Okinaka, R.T., Strniste, G.F., Barnhart, B.J., 1982. Induction of 6-thioguanine-resistant mutations by rat-liver homogenate (S9)-activated promutagens in human embryonic skin fibroblasts. Mutat. Res. Toxicol. 101, 87–98. https://doi.org/10.1016/0165-1218(82)90168-9

Chung, K.-T., Murdock, C.A., Stevens, S.E., Li, Y.-S., Wei, C.-I., Huang, T.-S., Chou, M.W., 1995. Mutagenicity and toxicity studies of p-phenylenediamine and its derivatives. Toxicol. Lett. 81, 23–32. https://doi.org/10.1016/0378-4274(95)03404-8

Chung, K.-T., Murdock, C.A., Zhou, Y., Stevens, S.E., Li, Y.-S., Wei, C.-I., Fernando, S.Y., Chou, M.-W., 1996. Effects of the nitro-group on the mutagenicity and toxicity of some benzamines. Environ. Mol. Mutagen. 27, 67–74. https://doi.org/10.1002/(SICI)1098-2280(1996)27:1<67::AID-EM9>3.0.CO;2-B

de Cássia Ribeiro Gonçalves, R., Rezende Kitagawa, R., Aparecida Varanda, E., Stella Gonçalves Raddi, M., Andrea Leite, C., Regina Pombeiro Sponchiado, S., 2016. Effect of biotransformation by liver S9 enzymes on the mutagenicity and cytotoxicity of melanin extracted from Aspergillus nidulans. Pharm. Biol. 54, 1014–1021. https://doi.org/10.3109/13880209.2015.1091846

Degen, G.H., Lebrun, S., Lektarau, Y., Föllmann, W., 2005. Modulation of ochratoxin A induced DNA-damage in urothelial cell cultures. Mycotoxin Res. 21, 57–60. https://doi.org/10.1007/BF02954819

Demarini, D.M., Brimer, P.A., Hsie, A.W., 1984. Cytotoxicity and mutagenicity of coal oils in the CHO/HGPRT assay. Environ. Mutagen. 6, 517–527. https://doi.org/10.1002/em.2860060405

Diaz, D., Scott, A., Carmichael, P., Shi, W., Costales, C., 2007. Evaluation of an automated in vitro micronucleus assay in CHO-K1 cells. Mutat. Res. Toxicol. Environ. Mutagen. 630, 1–13. https://doi.org/10.1016/j.mrgentox.2007.02.006

Erdinger, L., Schmezer, P., Razdan, R., Kumar, R., Spiegelhalder, B., Preussmann, R., Siddiqi, M., 1993. Caffeine-derived N-nitroso compounds. III: Mutagenicity in S. typhimurium and in vitro induction of DNA single-strand breaks in rat hepatocytes by mononitrosocaffeidine and dinitrosocaffeidine. Mutat. Res. Mutagen. Relat. Subj. 292, 41–49. https://doi.org/10.1016/0165-1161(93)90006-L

Glatt, H., Seidel, A., Bochnitschek, W., Marquardt, H., Marquardt, H., Hodgson, R.M., Grover, P.L., Oesch, F., 1986. Mutagenic and cell-transforming activities of triol-epoxides as compared to other chrysene metabolites. Cancer Res. 46, 4556–65.

Glatt, H., Seidel, A., Ribeiro, O., Kirkby, C., Hirom, P., Oesch, F., 1987. Metabolic activation to a mutagen of 3-hydroxy- trans -7,8-dihydroxy-7,8-dihydrobenzo[a]pyrene, a secondary metabolite of benzo[a]pyrene. Carcinogenesis 8, 1621–1627. https://doi.org/10.1093/carcin/8.11.1621

Goeger, D.., Hsie, A.., Anderson, K.., 1999. Co-mutagenicity of Coumarin (1,2-benzopyrone) with Aflatoxin B1 and Human Liver S9 in Mammalian Cells. Food Chem. Toxicol. 37, 581–589. https://doi.org/10.1016/S0278-6915(99)00046-0

Goeger, D.E., Anderson, K.E., Hsie, A.W., 1998. Coumarin chemoprotection against aflatoxin B1-induced gene mutation in a mammalian cell system: A species difference in mutagen activation and protection with chick embryo and rat liver S9. Environ. Mol. Mutagen. 32, 64–74. https://doi.org/10.1002/(SICI)1098-2280(1998)32:1<64::AID-EM8>3.0.CO;2-B

Haack, T., Erdinger, L., Boche, G., 2001. Mutagenicity in Salmonella typhimurium TA98 and TA100 of nitroso and respective hydroxylamine compounds. Mutat. Res. Toxicol. Environ. Mutagen. 491, 183–193. https://doi.org/10.1016/S1383-5718(01)00140-1

Holme, J.A., Soderlund, E.J., 1985. Species differences in the cytotoxic and genotoxic effects of 2-acetylaminofluorene and its primary metabolites 2-aminofluorene and N-OH-2-acetylaminofluorene. Carcinogenesis 6, 421–425. https://doi.org/10.1093/carcin/6.3.421

Horner, S.A., Fry, J.R., Clothier, R.H., Balls, M., 1985. A comparison of two cytotoxicity assays for the detection of metabolism-mediated toxicity in vitro: a study with cyclophosphamide. Xenobiotica 15, 681–686. https://doi.org/10.3109/00498258509047427

Huang, C.C., McKernan, K., Pantano, J.R., Sirianni, S.R., 1980. An in vitro metabolic activation assay using liver microsomes in diffusion chambers: induction of sister chromatid exchanges and chromosome aberrations by cyclophosphamide or ifosfamide in cultured human and Chinese hamster cells. Carcinogenesis 1, 37–40. https://doi.org/10.1093/carcin/1.1.37

Isabel, R.-R.M., Sandra, G.-A., Rafael, V.-P., Carmen, M.-V., Josefina, C.-E., del Carmen, C.-E.M., Rocío, G.-M., Francisco, A.-H., Elena, C.-S.M., 2012. Evaluation of 8-hydroxy-2′-deoxyguanosine (8-OHdG) adduct levels and DNA strand breaks in human peripheral blood lymphocytes exposed in vitro to polycyclic aromatic hydrocarbons with or without animal metabolic activation. Toxicol. Mech. Methods 22, 170–183. https://doi.org/10.3109/15376516.2011.623330

Kauderer, B., Zamith, H., Paumgartten, F.J.R., Speit, G., Holden, H.E., 1991. Evaluation of the mutagenicity of β-myrcene in mammalian cells in vitro. Environ. Mol. Mutagen. 18, 28–34. https://doi.org/10.1002/em.2850180106

Kim, K.-J., Lee, O.-H., Lee, B.-Y., 2010. Genotoxicity studies on fucoidan from Sporophyll of Undaria pinnatifida. Food Chem. Toxicol. 48, 1101–1104. https://doi.org/10.1016/j.fct.2010.01.032

Kitchin, R.M., Bechtold, W.E., Brooks, A.L., 1988. The structure-function relationships of nitrofluorenes and nitrofluorenones in the Salmonella mutagenicity and CHO sister-chromatid exchange assays. Mutat. Res. Toxicol. 206, 367–377. https://doi.org/10.1016/0165-1218(88)90123-1

Krishna, G., Kropko, M.L., Theiss, J.C., 1989. Use of the cytokinesis-block method for the analysis of munuclei in V79 Chinese hamster lung cells: results with mitomycin C and cyclophosphamide. Mutat. Res. Toxicol. 222, 63–69. https://doi.org/10.1016/0165-1218(89)90036-0

Kugler, U., Bauchinger, M., Schmid, E., Göggelmann, W., 1987. The effectiveness of S9 and microsomal mix on activation of cyclophosphamide to induce genotoxicity in human lymphocytes. Mutat. Res. Toxicol. 187, 151–156. https://doi.org/10.1016/0165-1218(87)90082-6

Lebsanft, J., McMahon, J.B., Steinmann, G.G., Shoemaker, R.H., 1989. A rapid in vitro method for the evaluation of potential antitumor drugs requiring metabolic activation by hepatic S9 enzymes. Biochem. Pharmacol. 38, 4477–4483. https://doi.org/10.1016/0006-2952(89)90659-X

Liewen, M.B., Marth, E.H., 1985. Evaluation of 1,3-pentadiene for mutagenicity by the Salmonella/mammalian microsome assay. Mutat. Res. Toxicol. 157, 49–52. https://doi.org/10.1016/0165-1218(85)90048-5

Lin, M.F., Wu, C.L., Wang, T.C., 1987. Pesticide clastogenicity in Chinese hamster ovary cells. Mutat. Res. Toxicol. 188, 241–250. https://doi.org/10.1016/0165-1218(87)90095-4

Lynch, B., Lau, A., Baldwin, N., Hofman-Hüther, H., Bauter, M.R., Marone, P.A., 2013. Genotoxicity of dried Hoodia parviflora aerial parts. Food Chem. Toxicol. 55, 272–278. https://doi.org/10.1016/j.fct.2013.01.014

Ma, H., An, J., Hsie, A.W., Au, W.W., 1993. Mutagenicity and cytotoxicity of 2-methoxyethanol and its metabolites in Chinese hamster cells (the CHO/HPRT and AS52/GPT assays). Mutat. Res. Toxicol. 298, 219–225. https://doi.org/10.1016/0165-1218(93)90044-E

Machanoff, R., O'Neill, J.P., Hsie, A.W., 1981. Quantitative analysis of cytotoxicity and mutagenicity of benzo[a]pyrene in mammalian cells (CHO/HGPRT system). Chem. Biol. Interact. 34, 1–10. https://doi.org/10.1016/0009-2797(81)90084-3

Maksimova, V., Shalginskikh, N., Vlasova, O., Usalka, O., Beizer, A., Bugaeva, P., Fedorov, D., Lizogub, O., Lesovaya, E., Katz, R., Belitsky, G., Kirsanov, K., Yakubovskaya, M., 2021. HeLa TI cell-based assay as a new approach to screen for chemicals able to reactivate the expression of epigenetically silenced genes. PLoS One 16, e0252504. https://doi.org/10.1371/journal.pone.0252504

Maralhas, A., Monteiro, A., Martins, C., Kranendonk, M., Laires, A., Rueff, J., Rodrigues, A.S., 2006. Genotoxicity and endoreduplication inducing activity of the food flavouring eugenol. Mutagenesis 21, 199–204. https://doi.org/10.1093/mutage/gel017

Martins, C., Cação, R., Cole, K.J., Phillips, D.H., Laires, A., Rueff, J., Rodrigues, A.S., 2012. Estragole: A weak direct-acting food-borne genotoxin and potential carcinogen. Mutat. Res. Toxicol. Environ. Mutagen. 747, 86–92. https://doi.org/10.1016/j.mrgentox.2012.04.009

Maximino, S.C., Dutra, J.A.P., Rodrigues, R.P., Gonçalves, R.C.R., Morais, P.A.B., Ventura, J.A., Schuenck, R.P., Júnior, V.L., Kitagawa, R.R., S. Borges, W., 2020. Synthesis of Eugenol Derivatives and Evaluation of their Antifungal Activity Against Fusarium solani f. sp. piperis. Curr. Pharm. Des. 26, 1532–1542. https://doi.org/10.2174/1381612826666200403120448

Miadokova, E., Vlkova, V., Podstavkova, S., Slaninova, M., Vlek, D., 1998. Unicellular green algaChlamydomonas reinhardtii as an activation system for 2-aminofluorene. Environ. Mol. Mutagen. 31, 383–389. https://doi.org/10.1002/(SICI)1098-2280(1998)31:4<383::AID-EM11>3.0.CO;2-8

Müller, L., Kasper, P., Kaufmann, G., 1992. The clastogenic potential in vitro of pyrrolizidine alkaloids employing hepatocyte metabolism. Mutat. Res. Lett. 282, 169–176. https://doi.org/10.1016/0165-7992(92)90091-U

Nasr, M.L., Goldman, M., Klein, A.K., Dacre, J.C., 1988. SCE induction in Chinese hamster ovary cells (CHO) exposed to G agents. Mutat. Res. Toxicol. 204, 649–654. https://doi.org/10.1016/0165-1218(88)90068-7

O'Donovan, M.R., 1990. Mutation assays of ethyl methanesulphonate, benzidine and benzo[ <italic>a</italic> ]pyrene using Chinese hamster V79 cells. Mutagenesis 5, 9–13. https://doi.org/10.1093/mutage/5.Supplement.9

Oesch-Bartlomowicz, B., Arens, H.J., Richter, B., Hengstler, J.G., Oesch, F., 1997. Control of the mutagenicity of aromatic amines by protein kinases and phosphatases. Arch. Toxicol. 71, 601–611. https://doi.org/10.1007/s002040050433

Otto, M., Hansen, S.H., Dalgaard, L., Dubois, J., Badolo, L., 2008. Development of an in vitro assay for the investigation of metabolism-induced drug hepatotoxicity. Cell Biol. Toxicol. 24, 87–99. https://doi.org/10.1007/s10565-007-9018-x

Patierno, S.R., Lehman, N.L., Henderson, B.E., Landolph, J.R., 1989. Study of the ability of phenacetin, acetaminophen, and aspirin to induce cytotoxicity, mutation, and morphological transformation in C3H/10T1/2 clone 8 mouse embryo cells. Cancer Res. 49, 1038–44.

Perocco, P., Del Ciello, C., Mazzullo, M., Rocchi, P., Ferreri, A.., Paolini, M., Pozzetti, L., Cantelli-Forti, G., 1997. Cytotoxic and cell transforming activities of the fungicide methyl thiophanate on BALB/c 3T3 cells in vitro. Mutat. Res. Toxicol. Environ. Mutagen. 394, 29–35. https://doi.org/10.1016/S1383-5718(97)00120-4

Picada, J.N., da Silva, K.V.C.., Erdtmann, B., Henriques, A.T., Henriques, J.A.., 1997. Genotoxic effects of structurally related β-carboline alkaloids. Mutat. Res. Mol. Mech. Mutagen. 379, 135–149. https://doi.org/10.1016/S0027-5107(97)00116-4

Pirisi, L., Garcea, R., Pascale, R., Ruggiu, M.E., Feo, F., 1987. Control of Glucose-6-Phosphate Dehydrogenase Deficiency on the Formation of Mutagenic and Carcinogenic Metabolites Derived from Benzo(a)pyrene. Toxicol. Pathol. 15, 115–119. https://doi.org/10.1177/019262338701500118

Recio, L., Hsie, A.W., 1987. Modulation of the cytotoxicity and mutagenicity of benzo[a]pyrene and benzo[a]pyrene 7,8-diol by glutathione and glutathione S-transferases in mammalian cells (CHO/HGPRT assay). Mutat. Res. Mol. Mech. Mutagen. 178, 257–269. https://doi.org/10.1016/0027-5107(87)90276-4

Recio, L., Hsie, A.W., 1984. Glucuronide conjugation reduces the cytotoxicity but not the mutagenicity of benzo(a)pyrene in the CHO/HGPRT assay. Teratog. Carcinog. Mutagen. 4, 391–402. https://doi.org/10.1002/tcm.1770040503

Recio, L., Shepard, K.G., Hernandez, L.G., Kedderis, G.L., 2012. Dose-Response Assessment of Naphthalene-Induced Genotoxicity and Glutathione Detoxication in Human TK6 Lymphoblasts. Toxicol. Sci. 126, 405–412. https://doi.org/10.1093/toxsci/kfs012

Reddy, M.V., Storer, R.D., Laws, G.M., Armstrong, M.J., Barnum, J.E., Gara, J.P., McKnight, C.G., Skopek, T.R., Sina, J.F., DeLuca, J.G., Galloway, S.M., 2002. Genotoxicity of naturally occurring indole compounds: correlation between covalent DNA binding and other genotoxicity tests. Environ. Mol. Mutagen. 40, 1–17. https://doi.org/10.1002/em.10088

Ribas, G., Surrallés, J., Carbonell, E., Creus, A., Xamena, N., Marcos, R., 1998. Lack of genotoxicity of the herbicide atrazine in cultured human lymphocytes. Mutat. Res. Toxicol. Environ. Mutagen. 416, 93–99. https://doi.org/10.1016/S1383-5718(98)00081-3

Rogers, C.G., Boyes, B.G., Matula, T.I., Stapley, R., 1992. Evaluation of genotoxicity of tert.-butylhydroquinone in an hepatocyte-mediated assay with V79 Chinese hamster lung cells and in strain D7 of Saccharomyces cerevisiae. Mutat. Res. Toxicol. 280, 17–27. https://doi.org/10.1016/0165-1218(92)90014-Q

Sargent, E.V., Bradley, M.O., 1986. Genotoxic activity of m-nitrobenzaldehyde. Mutat. Res. Lett. 175, 133–137. https://doi.org/10.1016/0165-7992(86)90111-9

Sarrif, A.M., Arce, G.T., Krahn, D.F., O'Neil, R.M., Reynolds, V.L., 1994. Evaluation of carbendazim for gene mutations in the Salmonella/Ames plate-incorporation assay: the role of aminophenazine impurities. Mutat. Res. Toxicol. 321, 43–56. https://doi.org/10.1016/0165-1218(94)90119-8

Sbrana, I., Zaccaro, L., Lascialfari, D., Ceccherini, I., Loprieno, N., 1984. Human lymphocytes assay: Cyclophosphamide metabolic activation by S9 system with low cytotoxicity. Mutat. Res. Mutagen. Relat. Subj. 130, 411–416. https://doi.org/10.1016/0165-1161(84)90013-X

Schmid, E., Göggelmann, W., Bauchinger, M., 1986. Formaldehyde-induced cytotoxic, genotoxic and mutagenic response in human lymphocytes and Salmonella typhimurium. Mutagenesis 1, 427–431. https://doi.org/10.1093/mutage/1.6.427

Sheu, C.-J.W., Lee, J.H.K., Rodriguez, I., Randolph, S.C., 1991. The Use of Uninduced Rat Livfr S-9 to Supplement BALB/3T3 Cells in the In Vitro Transformation Assay. Drug Chem. Toxicol. 14, 113–126. https://doi.org/10.3109/01480549109017871

Šiviková, K., Dianovský, J., 1999. Genotoxic activity of the commercial herbicide containing bifenox in bovine peripheral lymphocytes. Mutat. Res. Toxicol. Environ. Mutagen. 439, 129–135. https://doi.org/10.1016/S1383-5718(98)00184-3

Slameňová, D., Budayová, E., Gábelová, A., Morávková, A., Pániková, L., 1986. Results of genotoxicity testing of mazindol (degonan), lithium carbonicum (contemnol) and dropropizine (ditustat) in Chinese hamster V79 and human EUE cells. Mutat. Res. Toxicol. 169, 171–177. https://doi.org/10.1016/0165-1218(86)90096-0

Slesinski, R.S., Guzzie, P.J., Putman, D.L., Ballantyne, B., 1988. In vitro and in vivo evaluation of the genotoxic potential of 2-ethyl-1,3-hexanediol. Toxicology 53, 179–198. https://doi.org/10.1016/0300-483X(88)90212-0

Slesinski, R.S., Hengler, W.C., Guzzie, P.J., Wagner, K.J., 1983. Mutagenicity evaluation of glutaraldehyde in a battery of in vitro bacterial and mammalian test systems. Food Chem. Toxicol. 21, 621–629. https://doi.org/10.1016/0278-6915(83)90150-3

Sobti, R.C., Krishan, A., Pfaffenberger, C.D., 1982. Cytokinetic and cytogenetic effects of some agricultural chemicals on human lymphoid cells in vitro: organophosphates. Mutat. Res. Toxicol. 102, 89–102. https://doi.org/10.1016/0165-1218(82)90149-5

Suárez, S., Sueiro, R.A., Garrido, J., 2000. Genotoxicity of the coating lacquer on food cans, bisphenol A diglycidyl ether (BADGE), its hydrolysis products and a chlorohydrin of BADGE. Mutat. Res. Toxicol. Environ. Mutagen. 470, 221–228. https://doi.org/10.1016/S1383-5718(00)00109-1

Suter, W., 1987. Mutagenicity of procarbazine for V79 Chinese hamster fibroblasts in the presence of various metabolic activation systems. Mutagenesis 2, 27–32. https://doi.org/10.1093/mutage/2.1.27

Szalay, B., Tátrai, E., Nyírő, G., Vezér, T., Dura, G., 2012. Potential toxic effects of iron oxide nanoparticles in in vivo and in vitro experiments. J. Appl. Toxicol. 32, 446–453. https://doi.org/10.1002/jat.1779

Tafazoli, M., Baeten, A., Geerlings, P., Kirsch-Volders, M., 1998. In vitro mutagenicity and genotoxicity study of a number of short-chain chlorinated hydrocarbons using the micronudeus test and the alkaline single cell gel electrophoresis technique (Comet assay) in human lymphocytes: a structure–activity relationship (Q. Mutagenesis 13, 115–126. https://doi.org/10.1093/mutage/13.2.115

Tafazoli, M., Kirsch-Volders, M., 1996. In vitro mutagenicity and genotoxicity study of 1,2-dichloroethylene, 1,1,2-trichloroethane, 1,3-dichloropropane, 1,2,3-trichloropropane and 1,1,3-trichloropropene, using the micronucleus test and the alkaline single cell gel electrophoresis technique (co. Mutat. Res. Toxicol. 371, 185–202. https://doi.org/10.1016/S0165-1218(96)90107-X

Tan, E.-L., Hsie, A.W., 1981. Effect of calcium phosphate and alumina C γ gels on the mutagenicity and cytotoxicity of dimethylnitrosamine as studied in the CHO/HGPRT system. Mutat. Res. Mol. Mech. Mutagen. 84, 147–156. https://doi.org/10.1016/0027-5107(81)90058-0

Tayama, S., Nakagawa, Y., 1994. Effect of scavengers of active oxygen species on cell damage caused in CHO-K1 cells by phenylhydroquinone, an o-phenylphenol metabolite. Mutat. Res. Lett. 324, 121–131. https://doi.org/10.1016/0165-7992(94)90056-6

Thompson, L.H., Carrano, A.V., Salazar, E., Felton, J.S., Hatch, F.T., 1983. Comparative genotoxic effects of the cooked-food-related mutagens Trp-P-2 and IQ in bacteria and cultured mammalian cells. Mutat. Res. Toxicol. 117, 243–257. https://doi.org/10.1016/0165-1218(83)90125-8

Thust, R., Kneist, S., 1979. Activity of citrinin metabolised by rat and human microsome fractions in clastogenicity and SCE assays on Chinese hamster V79-E cells. Mutat. Res. Toxicol. 67, 321–330. https://doi.org/10.1016/0165-1218(79)90028-4

Thust, R., Schneider, M., Wagner, U., Schreiber, D., 1991. Structure/activity investigations in eight arylalkyltriazenes comparison of chemical stability, mode of decomposition, and SCE induction in Chinese hamster V79-E cells. Cell Biol. Toxicol. 7, 145–165. https://doi.org/10.1007/BF00122828

Umar-Tsafe, N., Mohamed-Said, M.S., Rosli, R., Din, L. Bin, Lai, L.C., 2004. Genotoxicity of goniothalamin in CHO cell line. Mutat. Res. Toxicol. Environ. Mutagen. 562, 91–102. https://doi.org/10.1016/j.mrgentox.2004.05.011

Wells, D.A., Thomas, H.F., Digenis, G.A., 1988. Mutagenicity and cytotoxicity of n-methyl-2-pyrrolidinone and 4-(methylamino)butanoic acid in theSalmonella/microsome assay. J. Appl. Toxicol. 8, 135–139. https://doi.org/10.1002/jat.2550080211

Wening, J.V., Marquardt, H., Katzer, A., Jungbluth, K.H., Marquardt, H., 1995. Cytotoxicity and mutagenicity of Kevlar®: an in vitro evaluation. Biomaterials 16, 337–340. https://doi.org/10.1016/0142-9612(95)93262-C

Wetmore, B., 1999. Evidence for site-specific bioactivation of alachlor in the olfactory mucosa of the Long-Evans rat. Toxicol. Sci. 49, 202–212. https://doi.org/10.1093/toxsci/49.2.202

White, A.D., Hesketh, L.C., 1980. A method utilising human lymphocytes with in vitro metabolic activation for assessing chemical mutagenicity by sister-chromatid exchange analysis. Mutat. Res. Mol. Mech. Mutagen. 69, 283–291. https://doi.org/10.1016/0027-5107(80)90093-7

Yu, R.C.-T.C.-T., 1999. Genetic toxicity of cocaine. Carcinogenesis 20, 1193–1199. https://doi.org/10.1093/carcin/20.7.1193

Zetouni, N.C., Siraki, A.G., Weinfeld, M., Pereira, A.D.S., Martin, J.W., 2017. Screening of genotoxicity and mutagenicity in extractable organics from oil sands process-affected water. Environ. Toxicol. Chem. 36, 1397–1404. https://doi.org/10.1002/etc.3670

Zhang, Z., Fu, J., Yao, B., Zhang, X., Zhao, P., Zhou, Z., 2011. In vitro genotoxicity of danthron and its potential mechanism. Mutat. Res. Toxicol. Environ. Mutagen. 722, 39–43. https://doi.org/10.1016/j.mrgentox.2011.02.006

Zhu, S., Cunningham, M.L., Gray, T.E., Nettesheim, P., 1991. Cytotoxicity, genotoxicity and transforming activity of 4-(methylnitrosamino)-1-(3-pyridyl)-1-butanone (NNK) in rat tracheal epithelial cells. Mutat. Res. Toxicol. 261, 249–259. https://doi.org/10.1016/0165-1218(91)90040-S

Zhuge, J., 2003. Heterologous expression of human cytochrome P450 2E1 in HepG2 cell line. World J. Gastroenterol. 9, 2732. https://doi.org/10.3748/wjg.v9.i12.2732

Zwanenburg, T.S.B., 1988. Comparative analysis of the clastogenicity and cytotoxicity of airborne particulate matter generated during the fire at Schweizerhalle on November 1, 1986. Mutat. Res. Toxicol. 206, 395–409. https://doi.org/10.1016/0165-1218(88)90126-7

### 5.4 References of the dataset BTS3/historical

Abdallah, M.A.-E., Nguyen, K.-H., Moehring, T., Harrad, S., 2019. First insight into human extrahepatic metabolism of flame retardants: Biotransformation of EH-TBB and Firemaster-550 components by human skin subcellular fractions. Chemosphere 227, 1–8. https://doi.org/10.1016/j.chemosphere.2019.04.017

Abdallah, M.A.-E., Uchea, C., Chipman, J.K., Harrad, S., 2014. Enantioselective Biotransformation of Hexabromocyclododecane by in Vitro Rat and Trout Hepatic Sub-Cellular Fractions. Environ. Sci. Technol. 48, 2732–2740. https://doi.org/10.1021/es404644s

Adehin, A., Tan, K.S., Lu, Z., Cheng, Q., Tan, W., 2019. In vitro metabolic stability and biotransformation of isosteviol in human and rat liver fractions. Drug Metab. Pharmacokinet. 34, 194–200. https://doi.org/10.1016/j.dmpk.2019.02.005

Ames, B.N., Durston, W.E., Yamasaki, E., Lee, F.D., 1973. Carcinogens are Mutagens: A Simple Test System Combining Liver Homogenates for Activation and Bacteria for Detection. Proc. Natl. Acad. Sci. 70, 2281–2285. https://doi.org/10.1073/pnas.70.8.2281

Anderson, D., Phillips, B.J., 1985. Nitrofurazone—Genotoxicity studies in mammalian cells in vitro and in vivo. Food Chem. Toxicol. 23, 1091–1098. https://doi.org/10.1016/0278-6915(85)90057-2

Arukwe, A., Carteny, C.C., Eggen, T., Möder, M., 2018. Novel aspects of uptake patterns, metabolite formation and toxicological responses in Salmon exposed to the organophosphate esters—Tris(2-butoxyethyl)- and tris(2-chloroethyl) phosphate. Aquat. Toxicol. 196, 146–153. https://doi.org/10.1016/j.aquatox.2018.01.014

Ashrap, P., Zheng, G., Wan, Y., Li, T., Hu, W., Li, W., Zhang, H., Zhang, Z., Hu, J., 2017. Supporting Information for Discovery of a Widespread Metabolic Pathway within and among Phenolic Xenobiotics Pahriya. Proc. Natl. Acad. Sci. U. S. A. 114, 6062–6067. https://doi.org/10.1073/pnas.1700558114

Au, W.W., Johnston, D.A., Collie-Bruyere, C., Hsu, T.C., 1980. Short-term cytogenetic assays of nine cancer chemotherapeutic drugs with metabolic activation. Environ. Mutagen. 2, 455–464. https://doi.org/10.1002/em.2860020404

Azadniya, E., Mollergues, J., Stroheker, T., Billerbeck, K., Morlock, G.E., 2020. New incorporation of the S9 metabolising system into methods for detecting acetylcholinesterase inhibition. Anal. Chim. Acta 1129, 76–84. https://doi.org/10.1016/j.aca.2020.06.033

Balabanič, D., Filipič, M., Krivograd Klemenčič, A., Žegura, B., 2017. Raw and biologically treated paper mill wastewater effluents and the recipient surface waters: Cytotoxic and genotoxic activity and the presence of endocrine disrupting compounds. Sci. Total Environ. 574, 78–89. https://doi.org/10.1016/j.scitotenv.2016.09.030

Bernacki, D.T., Bryce, S.M., Bemis, J.C., Kirkland, D., Dertinger, S.D., 2016. γH2AX and p53 responses in TK6 cells discriminate promutagens and nongenotoxicants in the presence of rat liver S9. Environ. Mol. Mutagen. 57, 546–558. https://doi.org/10.1002/em.22028

Bimboes Detlev, Greim Helmut, 1976. Human lymphocytes as target cells in a metabolising test system in vitro for detecting potential mutagens. Mutat. Res. Mol. Mech. Mutagen. 35, 155–159. https://doi.org/10.1016/0027-5107(76)90177-9

Boon, J.P., Sleiderink, H.M., Helle, M.S., Dekker, M., Van Schanke, A., Roex, E., Hillebrand, M.T.J., Klamer, H.J.C., Govers, B., Pastor, D., Morse, D., Wester, P.G., De Boer, J., 1998. The use of a microsomal in vitro assay to study phase I biotransformation of chlorobornanes (toxaphene®) in marine mammals and birds: Possible consequences of biotransformation for bioaccumulation and genotoxicity. Comp. Biochem. Physiol. - C Pharmacol. Toxicol. Endocrinol. 121, 385–403. https://doi.org/10.1016/S0742-8413(98)10058-0

Borenfreund, E., Puerner, J.A., 1987. Short-term quantitative in vitro cytotoxicity assay involving an S-9 activating system☆. Cancer Lett. 34, 243–248. https://doi.org/10.1016/0304-3835(87)90173-X

Brendt, J., Crawford, S.E., Velki, M., Xiao, H., Thalmann, B., Hollert, H., Schiwy, A., 2021a. Is a liver comparable to a liver? A comparison of different rat-derived S9-fractions with a biotechnological animal-free alternative in the Ames fluctuation assay. Sci. Total Environ. 759, 143522. https://doi.org/10.1016/j.scitotenv.2020.143522

Brendt, J., Lackmann, C., Heger, S., Velki, M., Crawford, S.E., Xiao, H., Thalmann, B., Schiwy, A., Hollert, H., 2021b. Using a high-throughput method in the micronucleus assay to compare animal-free with rat-derived S9. Sci. Total Environ. 751, 142269. https://doi.org/10.1016/j.scitotenv.2020.142269

Brusick, D., Myhr, B., Galloway, S., Rundell, J., Jagannath, D.R., Tarka, S., 1986. Genotoxicity of theobromine in a series of short-term assays. Mutat. Res. Toxicol. 169, 105–114. https://doi.org/10.1016/0165-1218(86)90089-3

Burkina, V., Sakalli, S., Giang, P.T., Grabicová, K., Staňová, A.V., Zamaratskaia, G., Zlabek, V., 2020. In Vitro Metabolic Transformation of Pharmaceuticals by Hepatic S9 Fractions from Common Carp (Cyprinus carpio). Molecules 25, 2690. https://doi.org/10.3390/molecules25112690

Butt, C.M., Muir, D.C.G., Mabury, S.A., 2010. Biotransformation of the 8:2 fluorotelomer acrylate in rainbow trout. 2. In vitro incubations with liver and stomach S9 fractions. Environ. Toxicol. Chem. 29, 2736–2741. https://doi.org/10.1002/etc.348

Cabaton, N., Dumont, C., Severin, I., Perdu, E., Zalko, D., Cherkaoui-Malki, M., Chagnon, M.-C., 2009. Genotoxic and endocrine activities of bis(hydroxyphenyl)methane (bisphenol F) and its derivatives in the HepG2 cell line. Toxicology 255, 15–24. https://doi.org/10.1016/j.tox.2008.09.024

Charles, G.D., Bartels, M.J., Gennings, C., Zacharewski, T.R., Freshour, N.L., Bhaskar Gollapudi, B., Carney, E.W., 2000. Incorporation of S-9 activation into an ER-α transactivation assay☆. Reprod. Toxicol. 14, 207–216. https://doi.org/10.1016/S0890-6238(00)00070-8

Chen, M.-H., Zhang, S.-H., Jia, S.-M., Wang, L.-J., Ma, W.-L., 2022. In vitro biotransformation of tris(1,3-dichloro-2-propyl) phosphate and triphenyl phosphate by mouse liver microsomes: Kinetics and key CYP isoforms. Chemosphere 288, 132504. https://doi.org/10.1016/j.chemosphere.2021.132504

Chen, M., Guo, T., He, K., Zhu, L., Jin, H., Wang, Q., Liu, M., Yang, L., 2019. Biotransformation and bioconcentration of 6:2 and 8:2 polyfluoroalkyl phosphate diesters in common carp (Cyprinus carpio): Underestimated ecological risks. Sci. Total Environ. 656, 201–208. https://doi.org/10.1016/j.scitotenv.2018.11.297

Chen, M., Qiang, L., Pan, X., Fang, S., Han, Y., Zhu, L., 2015. In Vivo and in Vitro Isomer-Specific Biotransformation of Perfluorooctane Sulfonamide in Common Carp (Cyprinus carpio). Environ. Sci. Technol. 49, 13817–13824. https://doi.org/10.1021/acs.est.5b00488

Choi, J.M., Oh, S.J., Lee, J.-Y., Jeon, J.S., Ryu, C.S., Kim, Y.-M., Lee, K., Kim, S.K., 2015. Prediction of Drug-Induced Liver Injury in HepG2 Cells Cultured with Human Liver Microsomes. Chem. Res. Toxicol. 28, 872–885. https://doi.org/10.1021/tx500504n

Choi, K., Joo, H., Rose, R.L., Hodgso, E., 2006. Metabolism of chlorpyrifos and chlorpyrifos oxon by human hepatocytes. J. Biochem. Mol. Toxicol. 20, 279–291. https://doi.org/10.1002/jbt.20145

Coldham, N.G., Horton, R., Byford, M.F., Sauer, M.J., 2002. A binary screening assay for pro-oestrogens in food: metabolic activation using hepatic microsomes and detection with oestrogen sensitive recombinant yeast cells. Food Addit. Contam. 19, 1138–1147. https://doi.org/10.1080/0265203021000014789

Cox, J.A., Fellows, M.D., Hashizume, T., White, P.A., 2016. The utility of metabolic activation mixtures containing human hepatic post-mitochondrial supernatant (S9) for in vitro genetic toxicity assessment. Mutagenesis 31, 117–130. https://doi.org/10.1093/mutage/gev082

de Rijke, E., Essers, M.L., Rijk, J.C.W., Thevis, M., Bovee, T.F.H., van Ginkel, L.A., Sterk, S.S., 2013. Selective androgen receptor modulators: in vitro and in vivo metabolism and analysis. Food Addit. Contam. Part A 30, 1517–1526. https://doi.org/10.1080/19440049.2013.810346

Deisenroth, C., DeGroot, D.E., Zurlinden, T., Eicher, A., McCord, J., Lee, M.Y., Carmichael, P., Thomas, R.S., 2020. The alginate immobilisation of metabolic enzymes platform retrofits an estrogen receptor transactivation assay with metabolic competence. Toxicol. Sci. 178, 281–301. https://doi.org/10.1093/toxsci/kfaa147

Dubreil, E., Sczubelek, L., Burkina, V., Zlabek, V., Sakalli, S., Zamaratskaia, G., Hurtaud-Pessel, D., Verdon, E., 2020. In vitro investigations of the metabolism of Victoria pure blue BO dye to identify main metabolites for food control in fish. Chemosphere 238, 124538. https://doi.org/10.1016/j.chemosphere.2019.124538

Ellenton, J.A., Douglas, G.R., Nestmann, E.R., 1981. MUTAGENIC EVALUATION OF 1,1,2,3-TETRACHLORO-2- PROPENE, A CONTAMINANT IN PULP MILL EFFLUENTS, USING A BATTERY OF IN VITRO MAMMALIAN AND MICROBIAL TESTS. Can. J. Genet. Cytol. 23, 17–25. https://doi.org/10.1139/g81-003

Fang, M., Webster, T.F., Ferguson, P.L., Stapleton, H.M., 2015. Characterising the Peroxisome Proliferator-Activated Receptor (PPAR γ ) Ligand Binding Potential of Several Major Flame Retardants, Their Metabolites, and Chemical Mixtures in House Dust. Environ. Health Perspect. 123, 166–172. https://doi.org/10.1289/ehp.1408522

Fic, A., Žegura, B., Sollner Dolenc, M., Filipič, M., Peterlin Mašič, L., 2013. Mutagenicity and DNA Damage of Bisphenol a and its Structural Analogues in Hepg2 Cells. Arch. Ind. Hyg. Toxicol. 64, 189–200. https://doi.org/10.2478/10004-1254-64-2013-2319

Galloway, S.M., Bloom, A.D., Resnick, M., Margolin, B.H., Nakamura, F., Archer, P., Zeiger, E., 1985. Development of a standard protocol for in vitro cytogenetic testing with Chinese hamster ovary cells: Comparison of results for 22 compounds in two laboratories. Environ. Mutagen. 7, 1–51. https://doi.org/10.1002/em.2860070102

Géniès, C., Jacques-Jamin, C., Duplan, H., Rothe, H., Ellison, C., Cubberley, R., Schepky, A., Lange, D., Klaric, M., Hewitt, N.J., Grégoire, S., Arbey, E., Fabre, A., Eilstein, J., 2020. Comparison of the metabolism of 10 cosmetics-relevant chemicals in EpiSkinTM S9 subcellular fractions and in vitro human skin explants. J. Appl. Toxicol. 40, 313–326. https://doi.org/10.1002/jat.3905

Gomez, C.F., Constantine, L., Huggett, D.B., 2010. The influence of gill and liver metabolism on the predicted bioconcentration of three pharmaceuticals in fish. Chemosphere 81, 1189–1195. https://doi.org/10.1016/j.chemosphere.2010.09.043

Gonzalez, R.., Tarloff, J.., 2001. Evaluation of hepatic subcellular fractions for Alamar blue and MTT reductase activity. Toxicol. Vitr. 15, 257–259. https://doi.org/10.1016/S0887-2333(01)00014-5

Gorman, G.S., Coward, L., Kerstner-Wood, C., Freeman, L., Hebert, C.D., Kapetanovic, I.M., 2009. In-vitro and in-vivo metabolic studies of the candidate chemopreventative pentamethylchromanol using liquid chromatography/tandem mass spectrometry. J. Pharm. Pharmacol. 61, 1309–1318. https://doi.org/10.1211/jpp/61.10.0006

Guesmi, A., Sleno, L., 2020. In vitro metabolism of triclosan studied by liquid chromatography–high-resolution tandem mass spectrometry. Anal. Bioanal. Chem. 412, 335–342. https://doi.org/10.1007/s00216-019-02239-6

Gupta, R.S., Singh, B., 1982. Mutagenic responses of five independent geentic loci in CHO cells to a variety of mutagens. Mutat. Res. Mol. Mech. Mutagen. 94, 449–466. https://doi.org/10.1016/0027-5107(82)90307-4

Han, X., O'Connor, J.C., Donner, E.M., Nabb, D.L., Mingoia, R.T., Snajdr, S.I., Clarke, J.J., Kaplan, A.M., 2009. Non-coplanar 2,2′,3,3′,4,4′,5,5′,6,6′-decachlorobiphenyl (PCB 209) did not induce cytochrome P450 enzyme activities in primary cultured rat hepatocytes, was not genotoxic, and did not exhibit endocrine-modulating activities. Toxicology 255, 177–186. https://doi.org/10.1016/j.tox.2008.10.013

Hashimoto, Y., Moriguchi, Y., Oshima, H., Kawaguchi, M., Miyazaki, K., Nakamura, M., 2001. Measurement of estrogenic activity of chemicals for the development of new dental polymers. Toxicol. Vitr. 15, 421–425. https://doi.org/10.1016/S0887-2333(01)00046-7

Heflich, R.H., Casciano, D.A., Zhuo, Z., Djurić, Z., Fullerton, N.F., Beland, F.A., 1988. Metabolism of 2-acetylaminofluorene in the chinese hamster ovary cell mutation assay. Environ. Mol. Mutagen. 11, 167–181. https://doi.org/10.1002/em.2850110203

Hölzel, B.N., Pfannkuche, K., Allner, B., Allner, H.T., Hescheler, J., Derichsweiler, D., Hollert, H., Schiwy, A., Brendt, J., Schaffeld, M., Froschauer, A., Stahlschmidt-Allner, P., 2020. Following the adverse outcome pathway from micronucleus to cancer using H2B-eGFP transgenic healthy stem cells. Arch. Toxicol. 94, 3265–3280. https://doi.org/10.1007/s00204-020-02821-3

Jacobsen, N.W., Brooks, B.W., Halling-Sørensen, B., 2012. Suggesting a testing strategy for possible endocrine effects of drug metabolites. Regul. Toxicol. Pharmacol. 62, 441–448. https://doi.org/10.1016/j.yrtph.2012.02.003

Jaeg, J.P., Perdu, E., Dolo, L., Debrauwer, L., Cravedi, J.-P., Zalko, D., 2004. Characterisation of New Bisphenol A Metabolites Produced by CD1 Mice Liver Microsomes and S9 Fractions. J. Agric. Food Chem. 52, 4935–4942. https://doi.org/10.1021/jf049762u

Janer, G., LeBlanc, G.A., Porte, C., 2005. A comparative study on androgen metabolism in three invertebrate species. Gen. Comp. Endocrinol. 143, 211–221. https://doi.org/10.1016/j.ygcen.2005.03.016

Jeon, J., Hollender, J., 2019. In vitro biotransformation of pharmaceuticals and pesticides by trout liver S9 in the presence and absence of carbamazepine. Ecotoxicol. Environ. Saf. 183, 109513. https://doi.org/10.1016/j.ecoenv.2019.109513

Johanning, K., Hancock, G., Escher, B., Adekola, A., Bernhard, M.J., Cowan-Ellsberry, C., Domoradzki, J., Dyer, S., Eickhoff, C., Embry, M., Erhardt, S., Fitzsimmons, P., Halder, M., Hill, J., Holden, D., Johnson, R., Rutishauser, S., Segner, H., Schultz, I., Nichols, J., 2012. Assessment of Metabolic Stability Using the Rainbow Trout ( Oncorhynchus mykiss ) Liver S9 Fraction. Curr. Protoc. Toxicol. 53, 1–28. https://doi.org/10.1002/0471140856.tx1410s53

Joo, H., Choi, K., Hodgson, E., 2010. Human metabolism of atrazine. Pestic. Biochem. Physiol. 98, 73–79. https://doi.org/10.1016/j.pestbp.2010.05.002

Kitamura, S., Ohmegi, M., Sanoh, S., Sugihara, K., Yoshihara, S., Fujimoto, N., Ohta, S., 2003a. Estrogenic activity of styrene oligomers after metabolic activation by rat liver microsomes. Environ. Health Perspect. 111, 329–334. https://doi.org/10.1289/ehp.5723

Kitamura, S., Sanoh, S., Kohta, R., Suzuki, T., Sugihara, K., Fujimoto, N., Ohta, S., 2003b. Metabolic Activation of Proestrogenic Diphenyl and Related Compounds by Rat Liver Microsomes. J. Heal. Sci. 49, 298–310. https://doi.org/10.1248/jhs.49.298

Kropf, C., Begnaud, F., Gimeno, S., Berthaud, F., Debonneville, C., Segner, H., 2020. In Vitro Biotransformation Assays Using Liver S9 Fractions and Hepatocytes from Rainbow Trout ( Oncorhynchus mykiss ): Overcoming Challenges with Difficult to Test Fragrance Chemicals. Environ. Toxicol. Chem. 39, 2396–2408. https://doi.org/10.1002/etc.4872

Ladd, M.A., Fitzsimmons, P.N., Nichols, J.W., 2016. Optimisation of a UDP-glucuronosyltransferase assay for trout liver S9 fractions: activity enhancement by alamethicin, a pore-forming peptide. Xenobiotica 46, 1066–1075. https://doi.org/10.3109/00498254.2016.1149634

Lai, Y., Cai, Z., 2012. In vitro metabolism of hydroxylated polybrominated diphenyl ethers and their inhibitory effects on 17β-estradiol metabolism in rat liver microsomes. Environ. Sci. Pollut. Res. 19, 3219–3227. https://doi.org/10.1007/s11356-012-0828-x

Legler, J., Dennekamp, M., Vethaak, A.D., Brouwer, A., Koeman, J.H., van der Burg, B., Murk, A.J., 2002. Detection of estrogenic activity in sediment-associated compounds using in vitro reporter gene assays. Sci. Total Environ. 293, 69–83. https://doi.org/10.1016/S0048-9697(01)01146-9

Li, J., Chen, M., Wang, Z., Ma, M., Peng, X., 2011. Analysis of environmental endocrine disrupting activities in wastewater treatment plant effluents using recombinant yeast assays incorporated with exogenous metabolic activation system. Biomed. Environ. Sci. 24, 132–9. https://doi.org/10.3967/0895-3988.2011.02.007

Li, S., Zhao, J., Huang, R., Santillo, M.F., Houck, K.A., Xia, M., 2019. Use of high-throughput enzyme-based assay with xenobiotic metabolic capability to evaluate the inhibition of acetylcholinesterase activity by organophosphorous pesticides. Toxicol. Vitr. 56, 93–100. https://doi.org/10.1016/j.tiv.2019.01.002

Li, S., Zhao, J., Huang, R., Travers, J., Klumpp-Thomas, C., Yu, W., MacKerell, A.D., Sakamuru, S., Ooka, M., Xue, F., Sipes, N.S., Hsieh, J.-H., Ryan, K., Simeonov, A., Santillo, M.F., Xia, M., 2021. Profiling the Tox21 Chemical Collection for Acetylcholinesterase Inhibition. Environ. Health Perspect. 129, EHP6993. https://doi.org/10.1289/EHP6993

Lopardo, L., Adams, D., Cummins, A., Kasprzyk-Hordern, B., 2018. Verifying community-wide exposure to endocrine disruptors in personal care products – In quest for metabolic biomarkers of exposure via in vitro studies and wastewater-based epidemiology. Water Res. 143, 117–126. https://doi.org/10.1016/j.watres.2018.06.028

Lopardo, L., Cummins, A., Rydevik, A., Kasprzyk-Hordern, B., 2017. New Analytical Framework for Verification of Biomarkers of Exposure to Chemicals Combining Human Biomonitoring and Water Fingerprinting. Anal. Chem. 89, 7232–7239. https://doi.org/10.1021/acs.analchem.7b01527

Madle, E., Tiedemann, G., Madle, S., Ott, A., Kaufmann, G., 1986. Comparison of S9 mix and hepatocytes as external metabolising systems in Mammalian cell cultures: Cytogenetic effects of 7,12-dimethylbenzanthracene and aflatoxin B1. Environ. Mutagen. 8, 423–437. https://doi.org/10.1002/em.2860080311

Miller, G.E., Brabec, M.J., Kulkarni, A.P., 1986. Mutagen activation of 1,2-dibromo-3-chloropropane by cytosolic glutathione s-transferases and microsomal enzymes. J. Toxicol. Environ. Health 19, 503–518. https://doi.org/10.1080/15287398609530948

Montaño, M., Cocco, E., Guignard, C., Marsh, G., Hoffmann, L., Bergman, Å., Gutleb, A.C., Murk, A.J., 2012. New Approaches to Assess the Transthyretin Binding Capacity of Bioactivated Thyroid Hormone Disruptors. Toxicol. Sci. 130, 94–105. https://doi.org/10.1093/toxsci/kfs228

Morohoshi, K., Yamamoto, H., Kamata, R., Shiraishi, F., Koda, T., Morita, M., 2005. Estrogenic activity of 37 components of commercial sunscreen lotions evaluated by in vitro assays. Toxicol. Vitr. 19, 457–469. https://doi.org/10.1016/j.tiv.2005.01.004

Murk, A., Morse, D., Boon, J., Brouwer, A., 1994. In vitro metabolism of 3,3′,4,4′-tetrachlorobiphenyl in relation to ethoxyresorufin-O-deethylase activity in liver microsomes of some wildlife species and rat. Eur. J. Pharmacol. Environ. Toxicol. Pharmacol. 270, 253–261. https://doi.org/10.1016/0926-6917(94)90069-8

Natarajan, A.T., Tates, A.D., van Buul, P.P.W., Meijers, M., de Vogel, N., 1976. Ctrogenetic effects of mutagens/carcinogens after activation in a microsomal system in vitro I. Induction of chromosome aberrations and sister chromatid exchanges by diethylnitrosamine (DEN) and dimethylnitrosamine (DMN) in CHO cells in the presence of ra. Mutat. Res. Mol. Mech. Mutagen. 37, 83–90. https://doi.org/10.1016/0027-5107(76)90057-9

Neft, R.E., Schol, H.M., Fu, P.P., Casciano, D.A., 1990. The induction of aneuploidy by 3-nitrobenzo[a]pyrene in Chinese hamster ovary cells. Mutagenesis 5, 221–228. https://doi.org/10.1093/mutage/5.3.221

Nguyen, K.-H., Abou-Elwafa Abdallah, M., Moehring, T., Harrad, S., 2017. Biotransformation of the Flame Retardant 1,2-Dibromo-4-(1,2-dibromoethyl)cyclohexane (TBECH) in Vitro by Human Liver Microsomes. Environ. Sci. Technol. 51, 10511–10518. https://doi.org/10.1021/acs.est.7b02834

Nichols, J.W., Ladd, M.A., Fitzsimmons, P.N., 2018. Measurement of Kinetic Parameters for Biotransformation of Polycyclic Aromatic Hydrocarbons by Trout Liver S9 Fractions: Implications for Bioaccumulation Assessment. Appl. Vitr. Toxicol. 4, 365–378. https://doi.org/10.1089/aivt.2017.0005

Obringer, C., Wu, S., Troutman, J., Karb, M., Lester, C., 2021. Effect of chain length and branching on the in vitro metabolism of a series of parabens in human liver S9, human skin S9, and human plasma. Regul. Toxicol. Pharmacol. 122, 104918. https://doi.org/10.1016/j.yrtph.2021.104918

Ousji, O., Ohlund, L., Sleno, L., 2020. Comprehensive In Vitro Metabolism Study of Bisphenol A Using Liquid Chromatography-High Resolution Tandem Mass Spectrometry. Chem. Res. Toxicol. 33, 1468–1477. https://doi.org/10.1021/acs.chemrestox.0c00042

Pelkonen, O., 2009. Comparison of metabolic stability and metabolite identification of 55 ECVAM/ICCVAM validation compounds between human and rat liver homogenates and microsomes - a preliminary analysis. ALTEX 26, 214–222. https://doi.org/10.14573/altex.2009.3.214

Peng, B., Liu, M., Han, Y., Wanjaya, E.R., Fang, M., 2019. Competitive Biotransformation among Phenolic Xenobiotic Mixtures: Underestimated Risks for Toxicity Assessment. Environ. Sci. Technol. 53, 12081–12090. https://doi.org/10.1021/acs.est.9b04968

Phillips, A.L., Herkert, N.J., Ulrich, J.C., Hartman, J.H., Ruis, M.T., Cooper, E.M., Ferguson, P.L., Stapleton, H.M., 2020. In Vitro Metabolism of Isopropylated and tert-Butylated Triarylphosphate

Esters Using Human Liver Subcellular Fractions. Chem. Res. Toxicol. 33, 1428–1441. https://doi.org/10.1021/acs.chemrestox.0c00002

Pratt, R.M., Willis, W.D., 1985. In vitro screening assay for teratogens using growth inhibition of human embryonic cells. Proc. Natl. Acad. Sci. 82, 5791–5794. https://doi.org/10.1073/pnas.82.17.5791

Recio, L., Hsie, A.W., 1984. Glucuronide conjugation reduces the cytotoxicity but not the mutagenicity of benzo(a)pyrene in the CHO/HGPRT assay. Teratog. Carcinog. Mutagen. 4, 391–402. https://doi.org/10.1002/tcm.1770040503

Richardson, S., Bai, A., A. Kulkarni, A., F. Moghaddam, M., 2016. Efficiency in Drug Discovery: Liver S9 Fraction Assay As a Screen for Metabolic Stability. Drug Metab. Lett. 10, 83–90. https://doi.org/10.2174/1872312810666160223121836

Rijk, J.C.W., Bovee, T.F.H., Groot, M.J., Peijnenburg, A.A.C.M., Nielen, M.W.F., 2008. Evidence of the indirect hormonal activity of prohormones using liver S9 metabolic bioactivation and an androgen bioassay. Anal. Bioanal. Chem. 392, 417–425. https://doi.org/10.1007/s00216-008-2275-6

Ritter, C.L., Bennett, K.K., Fullerton, N.F., Beland, F.A., Malejka-Giganti, D., 1996. Effect of ovariectomy on the in vitro and in vivo activation of carcinogenic N -2-fluorenylhydroxamic acids by rat mammary gland and liver. Carcinogenesis 17, 2411–2418. https://doi.org/10.1093/carcin/17.11.2411

Schmidt, J., Kotnik, P., Trontelj, J., Knez, Ž., Mašič, L.P., 2013. Bioactivation of bisphenol A and its analogs (BPF, BPAF, BPZ and DMBPA) in human liver microsomes. Toxicol. Vitr. 27, 1267–1276. https://doi.org/10.1016/j.tiv.2013.02.016

Shao, Y., Schiwy, A., Glauch, L., Henneberger, L., König, M., Mühlenbrink, M., Xiao, H., Thalmann, B., Schlichting, R., Hollert, H., Escher, B.I., 2020. Optimisation of a pre-metabolisation procedure using rat liver S9 and cell-extracted S9 in the Ames fluctuation test. Sci. Total Environ. 749, 141468. https://doi.org/10.1016/j.scitotenv.2020.141468

Shen, M., Cheng, J., Wu, R., Zhang, S., Mao, L., Gao, S., 2012. Metabolism of polybrominated diphenyl ethers and tetrabromobisphenol A by fish liver subcellular fractions in vitro. Aquat. Toxicol. 114–115, 73–79. https://doi.org/10.1016/j.aquatox.2012.02.010

Slesinski, R.S., Hengler, W.C., Guzzie, P.J., Wagner, K.J., 1983. Mutagenicity evaluation of glutaraldehyde in a battery of in vitro bacterial and mammalian test systems. Food Chem. Toxicol. 21, 621–629. https://doi.org/10.1016/0278-6915(83)90150-3

Takatori, S., Kitagawa, Y., Oda, H., Miwa, G., Nishikawa, J., Nishihara, T., Nakazawa, H., Hori, S., 2003. Estrogenicity of Metabolites of Benzophenone Derivatives Examined by a Yeast Two-Hybrid Assay. J. Heal. Sci. 49, 91–98. https://doi.org/10.1248/jhs.49.91

Takehisa, S., Kanaya, N., Rieger, R., 1988. Promutagen activation by Vicia faba: An assay based on the induction of sister-chromatid exchanges in Chinese hamster ovary cells. Mutat. Res. Mol. Mech. Mutagen. 197, 195–205. https://doi.org/10.1016/0027-5107(88)90093-0

Terasaki, M., Kosaka, K., Kunikane, S., Makino, M., Shiraishi, F., 2011. Assessment of thyroid hormone activity of halogenated bisphenol A using a yeast two-hybrid assay. Chemosphere 84, 1527–1530. https://doi.org/10.1016/j.chemosphere.2011.04.045

Thibaut, R., Schnell, S., Porte, C., 2009. Assessment of metabolic capabilities of PLHC-1 and RTL-W1 fish liver cell lines. Cell Biol. Toxicol. 25, 611–622. https://doi.org/10.1007/s10565-008-9116-4

van Lipzig, M.M.H., Vermeulen, N.P.E., Gusinu, R., Legler, J., Frank, H., Seidel, A., Meerman, J.H.N., 2005. Formation of estrogenic metabolites of benzo[a]pyrene and chrysene by cytochrome P450 activity and their combined and supra-maximal estrogenic activity. Environ. Toxicol. Pharmacol. 19, 41–55. https://doi.org/10.1016/j.etap.2004.03.010

van Vugt-Lussenburg, B.M.A., van der Lee, R.B., Man, H.-Y., Middelhof, I., Brouwer, A., Besselink, H., van der Burg, B., 2018. Incorporation of metabolic enzymes to improve predictivity of reporter gene assay results for estrogenic and anti-androgenic activity. Reprod. Toxicol. 75, 40–48. https://doi.org/10.1016/j.reprotox.2017.11.005

Vervliet, P., Den Plas, J. Van, De Nys, S., Duca, R.C., Boonen, I., Elskens, M., Van Landuyt, K.L., Covaci, A., 2019. Investigating the in vitro metabolism of the dental resin monomers BisGMA, BisPMA, TCD-DI-HEA and UDMA using human liver microsomes and quadrupole time of flight mass spectrometry. Toxicology 420, 1–10. https://doi.org/10.1016/j.tox.2019.03.007

Vignati, L., Turlizzi, E., Monaci, S., Grossi, P., Kanter, R. De, Monshouwer, M., 2005. An in vitro approach to detect metabolite toxicity due to CYP3A4-dependent bioactivation of xenobiotics. Toxicology 216, 154–167. https://doi.org/10.1016/j.tox.2005.08.003

Whitehead, F.W., San, R.H.C., Stich, H.F., 1983. An intestinal cell-mediated chromosome aberration test for the detection of genotoxic agents. Mutat. Res. Mol. Mech. Mutagen. 111, 209–217. https://doi.org/10.1016/0027-5107(83)90064-7

Winckler, K., Obe, G., Madle, S., Nau, H., 1984. Mutagenic activities of cyclophosphamide (NSC-26271) and its main metabolites in Salmonella typhimurium, human peripheral lymphocytes and Chinese hamster ovary cells. Mutat. Res. Mol. Mech. Mutagen. 129, 47–55. https://doi.org/10.1016/0027-5107(84)90122-2

Winuthayanon, W., Suksen, K., Boonchird, C., Chuncharunee, A., Ponglikitmongkol, M., Suksamrarn, A., Piyachaturawat, P., 2009. Estrogenic activity of diarylheptanoids from Curcuma comosa Roxb. requires metabolic activation. J. Agric. Food Chem. 57, 840–845. https://doi.org/10.1021/jf802702c

Wu, W.-N., McKown, L.A., Rybczynski, P.J., Demarest, K., 2010. Hepatic biotransformation of the new calcium-mimetic agent, RWJ-68025, in the rat and in man — API-MS/MS identification of metabolites. J. Pharm. Pharmacol. 55, 631–637. https://doi.org/10.1211/002235703765344531

Yahagi, T., Degawa, M., Seino, Y., Matsushima, T., Nagao, M., Sugimura, T., Hashimoto, Y., 1975. Mutagenicity of carcinogenic azo dyes and their derivatives. Cancer Lett. 1, 91–96. https://doi.org/10.1016/S0304-3835(75)95563-9

Yoshihara, S. ’i., 2004. Potent Estrogenic Metabolites of Bisphenol A and Bisphenol B Formed by Rat Liver S9 Fraction: Their Structures and Estrogenic Potency. Toxicol. Sci. 78, 50–59. https://doi.org/10.1093/toxsci/kfh047

Zhuang, S., Lv, X., Pan, L., Lu, L., Ge, Z., Wang, Jiaying, Wang, Jingpeng, Liu, J., Liu, W., Zhang, C., 2017. Benzotriazole UV 328 and UV-P showed distinct antiandrogenic activity upon human CYP3A4-mediated biotransformation. Environ. Pollut. 220, 616–624. https://doi.org/10.1016/j.envpol.2016.10.011

Zwart, N., Nio, S.L., Houtman, C.J., de Boer, J., Kool, J., Hamers, T., Lamoree, M.H., 2018. High-Throughput Effect-Directed Analysis Using Downscaled in Vitro Reporter Gene Assays To Identify Endocrine Disruptors in Surface Water. Environ. Sci. Technol. 52, 4367–4377. https://doi.org/10.1021/acs.est.7b06604

1756 6. Appendix

1757 6.1 DEERS protocol

| Data Extraction, Evaluation, and Reliability Schema (DEERS) | | |
|---|---|---|
| **Criteria/(Sub-)Domain** | **Score** | **Definition - Comment** |
| Documentation of observations with importance to BTS study relevance and identification (not part of the reliability assessment) | | |
| Authors | | |
| Title | | |
| Year | | |
| Journal | | |
| Bibliographic reference | | |
| Assigned ID number | | |
| Define test system (organism). | | |
| Which endpoints were investigated (e.g., mode of action)? | | |
| What methods were applied for endpoint recording? | | |
| What type of external BTS was applied? | | |
| Reliability assessment of BTS method reporting | | |
| If assessable, a score is given for every category. In the specific cases that data items are not applicable, e.g. "strains" for human BTS, the score is defined as "na" and will get an automatic full score. Scores are dichotomous (1 – reported, 0 – not reported). | | |
| **Criteria Group I: BTS characterisation** | | |
| 1) Is the origin, producer, or vendor of the BTS defined? (if yes, define) | | |
| 2) Is the species of BTS origin defined? | | |

| |
|---|
| 3) Is the strain of the test animal defined? |
| 4) Is the sex of individuals defined? (individuals used for BTS pooling) |
| 5) Is the pooling of original liver tissue for BTS production defined? (number of individuals) |
| 6) Are details on husbandry given? (diet, weight, age, etc.) |
| 7) Was the BTS induced in the test animal before extraction? (if yes, define compound or mixture) |
| **Criteria Group II: BTS reaction components** |
| 8) Is the BTS protein concentration defined? (if yes, define in mg/mL) |
| 9) Is the BTS reaction buffer system defined? (if yes, define type and concentration in mM) |
| 10) Are the overall final concentrations listed of all relevant components partaking in the BTS reaction? (if a dilution factor is applied, define) |
| 11) Are primary BTS cofactors mentioned? (if yes, define and give concentration in mM) |
| 12) Are secondary BTS cofactors mentioned? (if yes, define and give concentration in mM) |
| 13) Is there a solvent (as an exposure vehicle) involved in the BTS reaction? (if yes, define) |
| **Criteria Group III: BTS experimental setup** |
| 14) Is a specific BTS incubation period named during which the reaction takes place? (if yes, define in min or h) |

| | | |
|---|---|---|
| 15) Is the BTS incubation reaction temperature named? (define in °C) | | |
| 16) Have BTS-specific controls been employed? (define) | | |
| Other observations of study relevance (not part of the reliability assessment) | | |
| What is the exposure regime? (define chemical and range, if feasible) | | |
| Name post BTS procedures | | |
| **Criteria/(Sub-)Domain** | **Score** | **Definition - Comment** |

6.2 R-Code

```
################################################################################
################################################################################
### MCAs ###
################################################################################
simpmca <- read.csv('simpmca2.csv')

# yr5, group, total points are qualitative supplementary variables

require(tidyverse) #
require(FactoMineR) #
require(factoextra) #
require(dplyr) #
require(scales) #
require(egg) #
require(scales) #
################################################################################
################################################################################
# MCA - prep data
missing_counts <- colSums(is.na(simpmca))
missing_counts # no NAs
simpmca$total_points <- as.character(simpmca$total_points)

# example fix capital letters (done for all)
simpy <- simpmca
simpy <- simpy %>%
  mutate(strain = ifelse(strain == "other", "Other", strain))

################################################################################
```

```r
mca1 <- MCA(simpy[2:19],quali.sup=16:18, level.ventil = 0.01, graph = F) # exclude ID
# check that correct columns are selected (yr5, group, total_points as quali.sup)
summary(mca1)

# calculate the squared singular values
squared_singular_values <- mca1$svd$vs^2
# calculate the percentage of total variance explained by each dimension
variance_explained <- (squared_singular_values / sum(squared_singular_values)) * 100

# plot theme (example - margin changed depending on labels)
theme <- theme(axis.title.x = element_text(hjust = 0.5, size = 15, margin = margin(0.5,0,0,0,
'cm')),  axis.title.y = element_text(hjust = 0.5, size = 15, margin = margin(0,0.5,0,0,
'cm')),  legend.title = element_text(hjust = 0, size = 15, margin = margin(0,0,0.25,0, 'cm')),
legend.text = element_text(hjust = 0, vjust = 0.5, size = 15, margin = margin(0,0,0,0.25,
'cm')),   panel.border = element_rect(colour = 'black', fill = 0, linewidth = 0.5),
axis.text.y= element_text(hjust = 0.5, size = 10, margin = margin(0.1,0.1,0.1,0.1, 'cm')),
axis.text.x= element_text(hjust = 0.5, vjust = 0.9, angle = 0, size = 10, margin =
margin(0.1,0.1,0.1,0.1, 'cm')),  legend.key.size = unit(1.25, 'cm'), axis.ticks.length =
unit(0.25, 'cm'),  legend.spacing.y = unit(0.5, 'cm'), legend.margin = margin(0.5,0,0.5,0.5,
'cm'), legend.box.spacing = unit(0.5, 'cm'), panel.background = element_rect(fill = 'white',
colour = 'black', linetype='solid', linewidth = 0.5), legend.background =
element_rect(fill='white',colour = 'white'),  legend.key = element_rect(fill = 'white', colour
= 'white'), plot.margin = margin(1,0.5,0,0.5, 'cm'))

################################################################################
# PLOT (EXAMPLE - STRAIN)
################################################################################
### STRAIN
mca1_obs_df = data.frame(mca1$ind$coord, group = simpy$strain)

# test plot
fviz_mca_biplot(mca1, geom=c('point'), pointsize = 1.5, label ='none',
                ellipse.alpha = 0.5,ellipse.border.remove = FALSE,
                invisible ='var', habillage= 'strain',ellipse.level=0.05,
                addEllipses=F, alpha = 1)
table(mca1_obs_df$group)

mca1_obs_df$group <- factor(mca1_obs_df$group , levels = c('Fi', 'SD', 'Wi', 'Other', 'nd',
'na'))

mca1_obs_df <- na.omit(mca1_obs_df)

require(dplyr)
centroids <- mca1_obs_df %>%
  group_by(group) %>%
  summarize(Centroid_Dim.1 = mean(Dim.1), Centroid_Dim.2 = mean(Dim.2))

plot.1 <- ggplot(data= mca1_obs_df,
       aes(x = Dim.1, y = Dim.2, colour = group)) +
  geom_hline(yintercept = 0, colour = "black", linetype = 'dashed', linewidth = 0.25) +
geom_vline(xintercept = 0, colour = "black", linetype = 'dashed', linewidth = 0.25) +
```

```
geom_abline(intercept = 0, slope = 1, linetype = 'solid', linewidth= 1.5, color =
'#4D4D4DFF')+  stat_ellipse(aes(fill = group, colour = group), linewidth = 0.75, geom =
'polygon', alpha = 0, level = 0.95, show.legend = F, data = mca1_obs_df)+
geom_point(aes(colour=group), size = 1.5, alpha = 0.8)+   scale_colour_manual(values =
c('royalblue4', 'palegreen4', 'goldenrod2', '#4D4D4DFF', '#BABABAFF', '#689FB0FF'))+  labs(x =
paste("Dim.1 (", round(variance_explained[1],1), "%)"),  y = paste("Dim.2 (",
round(variance_explained[2],1), "%)"), fill = '', tag = '', caption = '', title = '') +
geom_point(data = centroids, aes(x = Centroid_Dim.1, y = Centroid_Dim.2), shape = 2, stroke =
2, size = 8) + theme +  guides(color = guide_legend(override.aes = list(size = 8), 'Strain'))+
scale_x_continuous(labels = label_number(accuracy = 0.5)) +  scale_y_continuous(labels =
label_number(accuracy = 0.5))

plot.1

########################################################################

### scores ###

########################################################################

require(dplyr) # mutate

require(tidyr) # pivot_longer

require(ggplot2) # plotting

require(tidytext) # reorder_within

require(cowplot) # print plots

########################################################################

## absolute scores all domains

scores.sorted <- read.csv('scores.sorted.csv')

colnames(scores.sorted)[1] <- 'no'

# pivot longer for all domains in same column

scores.long <- pivot_longer(scores.sorted[c(1:2,19:22)], -c(no, id, total_points), names_to =
'domain', values_to = 'score')

# order domain

scores.long$domain <- factor(scores.long$domain, levels = c('total_bts_experimental_setup',
'total_bts_reaction', 'total_bts_characterisation'))

scores.a <- scores.long %>%

  mutate(id = reorder_within(id, no, id)) %>%

  ggplot(aes(x = id, y = score, fill = id)) +

  ggtitle(label = 'All primary domains')+

  theme(plot.title = element_text(hjust = 0.5, face = 'bold', size = 10, margin =
margin(0,0,0.7,0, 'cm')), axis.title.x = element_text(hjust = 0.5, face = 'bold', size = 10,
margin = margin(0.3,0,0,0, 'cm')), axis.title.y = element_text(hjust = 0.5, face = 'bold',
size = 10, margin = margin(0,0.3,0,0, 'cm')), axis.text.x = element_text(hjust = 0.5, face =
'bold', size = 10), legend.position = 'none', axis.text.y = element_text(hjust = 0.5, face =
'bold', size = 3.5), axis.ticks.length.y = unit(0, 'cm'), axis.line = element_line(colour =
'black', linewidth = 0.1, linetype = 'solid'), panel.background = element_rect(fill =
'white'), plot.margin = margin(1,0.5,0,0.5, 'cm'))+  labs(x = 'Study ID', y = 'Absolute
score', fill = 'Domain', caption = '', tag = '') +   coord_flip() +  scale_x_reordered() +
geom_bar(stat = 'identity', aes(x = id, y = score, fill = domain), alpha = 0.7, width = 0.9,
position = position_stack(reverse = FALSE)) +   geom_hline(yintercept = c(12.8, 16), colour =
c('red', 'darkgreen'), alpha = 0.7, linewidth = 0.7, linetype = 'dashed')+
scale_fill_manual(values = c('#000066', '#FFCC00', '#CC0000'), labels = c('BTS experimental
setup','BTS reaction', 'BTS characterisation'))

scores.a

## absolute scores per domain

# barplot scores - one plot per domain

scores <- read.csv('lungu.scores_20230305.csv')

colnames(scores)[1] <- 'id'
```

```
# pivot longer for all domains in same column
tot.scores <- pivot_longer(scores[c(1,18:21)], -c(id, total_points), names_to = 'domain', values_to = 'score')

tot.scores$domain <- factor(tot.scores$domain, levels = c('total_bts_characterisation',
                                                             'total_bts_reaction',
                                                             'total_bts_experimental_setup'))

dom <- as_labeller(c(`total_bts_characterisation` = 'BTS\ncharacterisation', `total_bts_reaction` = 'BTS\nreaction', `total_bts_experimental_setup` = 'BTS\nexperimental setup'))

scores.b <- tot.scores %>%
  group_by(domain) %>%
  ungroup %>%
  mutate(domain = as.factor(domain), id = reorder_within(id, -score, domain)) %>%
  ggplot(aes(id, score, fill = domain)) +  theme(axis.title.x = element_text(hjust = 0.5, face = 'bold', size = 10, margin = margin(0.5,0,0,0, 'cm')),  axis.title.y = element_text(hjust = 0.5, face = 'bold', size = 10, margin = margin(0,0.3,0,0, 'cm')),  axis.text.x = element_text(hjust = 0.5, face = 'bold', size = 10), legend.position = 'none', axis.text.y = element_text(hjust = 0.5, face = 'bold', size = 1.5), axis.ticks.length.y = unit(0, 'cm'), panel.background = element_rect(fill = 'white'), plot.margin = margin(1,0.5,0,0.5, 'cm'), strip.background = element_rect(color = 'white', fill = 'white', linewidth  = 1.5, linetype = 'solid'), strip.text.x = element_text(hjust = 0.5, face = 'bold', size = 10, margin = margin(0,0,0.7,0, 'cm')), axis.line = element_line(colour = 'black', linewidth = 0.1, linetype = 'solid'))+ labs(x = 'Study ID', y = 'Absolute score', fill = 'Domain', caption = '', tag = '') + geom_col(show.legend = FALSE, alpha = 0.7, width = 0.9) + facet_wrap(~domain, scales = 'free_y', labeller = dom) + coord_flip() +  scale_x_reordered() + scale_fill_manual(values = c('#CC0000','#FFCC00', '#000066'))
scores.b

## relative scores per domain
# rel scores already ordered:
rel.scores <- read.csv('rel.score.csv') # as of 20230307 - needs update?
colnames(rel.scores)[1] <- 'no'
# pivot longer for all domains in same column
rel.scores.long <- pivot_longer(rel.scores[1:4], -no, names_to = 'domain', values_to = 'rel.score')

# order domains for plot
rel.scores.long$domain <- factor(rel.scores.long$domain, levels = c('tot_bts_char_percentage','tot_bts_rea_percentage','tot_bts_ex_percentage'))
# step/line plot relative score per domain
scores.c <- ggplot(rel.scores.long, mapping = aes(x = no, y = rel.score/100, colour = domain)) +  geom_point(size=0, alpha=0.7)+   geom_hline(yintercept = c(0.8, 1), colour = c('red', 'darkgreen'), alpha = 0.7, linewidth = 0.7, linetype = 'dashed')+  geom_step(linewidth = 1.2, alpha = 0.8) +  theme(plot.title = element_text(hjust = 0.5, face = 'bold', size = 10, margin = margin(0,0,0.7,0, 'cm')), axis.title.x = element_text(hjust = 0.5, face = 'bold', size = 10, margin = margin(0.5,0.5,0.8,0, 'cm')), axis.title.y = element_text(hjust = 0.5, face = 'bold', size = 10, margin = margin(0,0.3,0,0, 'cm')), axis.text.x = element_text(hjust = 0.5, face = 'bold', size = 10), legend.position = 'bottom', legend.justification = 'left', axis.text.y = element_text(hjust = 0.5, face = 'bold', size = 10), axis.ticks.length = unit(.25, 'cm'), panel.background = element_rect(fill = 'white'), legend.background = element_rect(fill = 'white',colour = 'white'), legend.text = element_text(hjust = 0, face = 'bold', size = 12, margin = margin(0,0,0.5,0, 'cm')), legend.key = element_rect(fill = 'white', colour = 'white'), plot.margin = margin(1,0.5,0.75,0.5, 'cm'), axis.line = element_line(colour = 'black', linewidth = 0.1, linetype = 'solid'))+ scale_y_continuous(labels = scales::percent)+ scale_colour_manual(values = c('#CC0000','#FFCC00','#000066'), labels = c('BTS characterisation','BTS reaction','BTS experimental setup'))+ labs(x = 'Number of publications', y = 'Relative score', colour = '',title = 'Relative score per domain', caption
```

```
= '', tag = '')+ guides(colour = guide_legend(override.aes = list(size=8,linetype=0), nrow=3,
label.vjust=-2))
scores.c

################################################################################
#################################################################################
### euler, upset, histograms ###
#################################################################################
require(UpSetR)
require(cowplot)
require(eulerr)
require(tidyr)
require(ggplot2)
################################################################################
# PLOTS (EXAMPLE - FIELD/TOPIC)
################################################################################
### FIELD/TOPIC
df <- read.csv('field.csv')
df.hist <- read.csv('field.hist.csv')
#############################################################################
# UPSET
plot.aa <- upset(df, nsets= 13, order.by = c('freq'), point.size = 3, line.size = 1,
                 mainbar.y.label = 'Publications per intersection', mb.ratio = c(0.65, 0.35),
                 sets.x.label = 'Publ. per field', set_size.show = T,
                 set_size.scale_max = 175, text.scale = 2, decreasing = c(T))

plot.a <- plot_grid(NULL, plot.aa$Main_bar, plot.aa$Sizes, plot.aa$Matrix,
                    nrow=2, align = 'hv', rel_heights = c(2,1.1,2,1), rel_widths = c(1.1, 2,
1.1, 2))
plot.a # 717 x 950 (1200x900)
#############################################################################
# EULER
venn <- euler(df[6:12])
venn
#'BioMed','Tox', 'Env', 'Nut', 'BioSci', 'AnChem', 'Vet'

plot.b <- plot(venn, quantities = list(labels =
c('9','88','40','11','4','3','2','11','1','','','2','','16',

'8','11','4','','1','3','','1','','1','','1','','2','2','1',

'','1','','','','','','','','','','','','','1',

'4','','','','','','','','','','','','','','','','',

'1','','','','','','','','','','','','','','','','','','','','','',
```

```
'','','','','','','','','','','','','','','','','','','','',''), font = 1, fontsize = 16),
list(font = 1, fontsize=17),
               edges = list(lty = 1, lwd = 2), col =
c('#3969ACFF','#11A579FF','#F2B701FF','#D9565CFF','#7F3C8DFF',
                                                       '#FF9933FF','#088BBEFF'),
               fills = c('#3969ACFF','#11A579FF','#F2B701FF','#D9565CFF','#7F3C8DFF',
                         '#FF9933FF','#088BBEFF'), alpha=0.5,
               legend = list(labels = c(' BioMed  ',' Tox  ', ' Env  ', ' Nut  ', ' BioSci  ',
                                        ' AnChem  ', ' Vet  '),
                             fontsize = 18, side = 'bottom', nrow = 1, ncol = 7, alpha = 0.8))
plot.b # 900 x 900

############################################################################
# HISTOGRAM
long_df <- pivot_longer(df.hist[c(2,4,6:12)], -c(id, yr5), names_to = 'variable', values_to =
'pub')
long_df$variable <- factor(long_df$variable, levels = c('BioMed','Tox', 'Env', 'Nut',
'BioSci', 'AnChem', 'Vet'))
plot.c <-
  ggplot(long_df)+
  theme(axis.title.x = element_text(hjust = 0.5, size = 18, margin = margin(1,0,0,0, 'cm')),
        axis.title.y = element_text(hjust = 0.5, size = 18, margin = margin(0,1,0,0, 'cm')),
        legend.text = element_text(hjust = 0, vjust = 0.5, size = 18, margin = margin(0,0,0,1,
'cm')),
        panel.border = element_rect(colour = 'black', fill = 0, linewidth = 0.5),
        axis.text.y= element_text(hjust = 0.5, size = 16, margin = margin(0.1,0.1,0.1,0.1,
'cm')),
        axis.text.x= element_text(hjust = 1, vjust = 0.9, angle = 45, size = 16, margin =
margin(0.1,0.1,0.1,0.1, 'cm')),
        legend.key.size = unit(1, 'cm'), axis.ticks.length = unit(0.5, 'cm'),
        legend.spacing.y = unit(1.0, 'cm'), legend.margin = margin(0.5,0.5,0.5,0.5, 'cm'),
        legend.box.spacing = unit(1.0, 'cm'),
        panel.background = element_rect(fill = 'white', colour = 'black', linetype='solid',
linewidth = 0.5),
        legend.background = element_rect(fill='white',colour = 'white'),
        legend.key = element_rect(fill = 'white', colour = 'white'), plot.margin =
margin(0.5,0.5,0.5,0.5, 'cm'))+
  labs(x = 'Year', y = 'Number of publications', fill = '', tag = '', caption = '', title =
'',
       subtitle = '')+
  scale_fill_manual(values = c('#3969ACFF','#11A579FF','#F2B701FF','#D9565CFF','#7F3C8DFF',
                               '#FF9933FF','#088BBEFF'))+
  geom_bar(stat = 'identity', aes(x = yr5, y = pub, fill = variable), width = 1,
           position = position_stack(reverse = FALSE))
plot.c # 1024 x 717

###############################################################################
################################################################################
```

```
### networks ###
################################################################################
simpmca <- read.csv('simpmca3.csv')
colnames(simpmca)[19] <- 'Dataset'

simpmca$total_points <- as.character(simpmca$total_points)
simpmca$X <- NULL

# example fix capital letters etc. (done for all)
require(dplyr)
simpy <- simpmca
simpy <- simpy %>%
  mutate(Strain = ifelse(Strain == "other", "Other", Strain))

allfactors <- c(colnames(simpy[c(2:18, 20)]))
require(dplyr)
simp.factor <- simpy %>%
  mutate_at(vars(allfactors), as.factor)

factors <- sapply(simp.factor, is.factor) # identify all categorical variables.
xFactor <- simp.factor[ , factors]
###############################################################################
require(arules)
require('arulesViz')

x <- apriori(xFactor, support = 0.1, confidence = 0.8, maxlen = 500)
color_palette <- colorRampPalette(c('#c8e6c9','#a5d6a7','#66bb6a','#43a047','#388e3c',
                                    "#2e7d32"))(n = 100)

plot(x, method = "graph", engine = "htmlwidget", itemCol = '#4D4D4DFF',
     nodeCol = rev(color_palette),
     max = 500, degree_highlight = 1) %>%
  visNodes(font=list(color="white", face="Arial"))

library(ggplot2)
x@quality$Lift <- x@quality$lift
x@quality$Support <- x@quality$support
#x@itemInfo$Labels <- x$labels
plot(x, method = "graph",engine = "ggplot2",
     control = list(edges = ggraph::geom_edge_link(
         end_cap = ggraph::circle(4, "mm"),
         start_cap = ggraph::circle(4, "mm"),
         # color = "black",
         arrow = arrow(length = unit(1, "mm"), angle = 20, type = "closed"),
```

```
          alpha = .3, max = 50 #, nodeCol = color_palette),
        #nodes = ggraph::geom_node_point(aes(color="labels", size = 10)),
        nodes = ggraph::geom_node_point(aes_string(size = "Support", color = "Lift")),
        nodetext = ggraph::geom_node_label(aes_string(label = "label"), alpha = .9,
                                           repel = T)), limit = 100) +
  scale_color_gradient2(low = '#c8e6c9', mid = '#66bb6a', high = '#2e7d32', midpoint = 4) +
  #scale_color_gradient2(colorRampPalette(c('lightgrey', '#3969ACFF','#000066'))(100)) +
  scale_size(range = c(3, 10))

# pdf 8 x 12
```

### 6.3 Abbreviations

A list of abbreviations, as appearing in the main article, SM, and supplementary information material.

| | |
|---|---|
| 3-MC | 3-methylcholanthrene |
| AChE | Acetylcholinesterase |
| ACN | Acetonitrile |
| alc | Alcoholic solvents (EtOH, MeOH) |
| AnChem | analytical chemistry |
| AOP | Adverse outcome pathway |
| AR | androgen receptor |
| Aro | Aroclor and other PCBs |
| BioMed | biomedicine (other than tox.) |
| BioSci | bioscience (non-medicine) |
| BNF | Beta-naphthoflavone |
| BTS | externally added biotransformation system |
| BTS1-3 | investigated literature databases |
| CA | chemical analysis |
| cult | culture medium |
| CYP | Cytochromes P450 |
| CYP450 (x) | Cytochromes P450, X families not defined here |
| Cyto | Cytotoxicity |
| DEERS | Data Extraction, Evaluation, and Reliability Schema |
| dh | dehydrogenese |
| DMSO | Dimethyl sulfoxide |
| DTT | Dithiothreitol |
| EC JRC EURL ECVAM | European Commission, Joint Research Centre. EU Reference Laboratory - European Centre for the Validation of Alternative Methods |
| EDC | endocrine disruption |
| EDCs | endocrine disruptive chemicals |
| Env | environmental sciences (env. Tox. + env. Chem.) |
| ER | estrogene receptor |
| EtOH | ethanol |
| ex | external |
| F | female |

| | |
|---|---|
| Fi | Fisher |
| G6P | Glucose 6-phosphate |
| G6P-dh | Glucose 6-phosphate dehydrogenase |
| GSH | Glutathione |
| GST | Glutathione S-transferase |
| H2O | water based: saline buffers, media, H20 itself |
| HPLC | High-performance liquid chromatography |
| HTS | high throughput screening |
| IATA | integrated approaches to testing and assessment |
| in | internal |
| inab | inactivated BTS |
| iso | isocitrate |
| isocitrate-dh | isocitrate dehydrogenase |
| KE | Key event |
| LE | Long-Evans |
| M | male |
| MCA | multiple correspondence analysis |
| MCA | Methylcholanthrene |
| MeOH | methanol |
| Meta | metabolites identification & characterisation |
| MIE | Molecular initiating events |
| MutGen | Mutagenicity & Genotoxcicty |
| na | not applicable/assessable, neutral scoring |
| NAD | Nicotinamide adenine dinucleotide, oxidised, NAD+ |
| NADH | Nicotinamide adenine dinucleotide, reduced |
| NADP | Nicotinamide adenine dinucleotide phosphate, oxidised, NADP+ |
| NADPH | Nicotinamide adenine dinucleotide phosphate, reduced |
| NADPx | Nicotinamide adenine dinucleotide phosphate, various forms NADPH, NADP+ |
| NADPx | Nicotinamide adenine dinucleotide phosphate, x stands for both types used, oxidised and reduced |
| NAM | new approach method |
| nc | not clear, negative scoring |
| nd | not defined, negative scoring |
| NeuDev | neuronal & developmental toxicity |
| Nut | nutrition |
| oBioSci | Other biosciences |
| oMammal | Other mammals |
| org | organic solvents (ACN, acetone, etc.) |
| PAPS | 3′-Phosphoadenosine-5′-phosphosulfate |
| PB | Phenobarbital |
| PBS | Phosphate-buffered saline |
| PCA | Principle component analysis |
| PCB | Polychlorinated biphenyl |
| PCB-PAH | Polychlorinated biphenyls and polycyclic aromatic hydrocarbons |
| PCDD | Polychlorinated dibenzodioxin |

| | |
|---|---|
| PCDF | Polychlorinated dibenzofuran |
| ph1 | phase 1 metabolism |
| ph2 | phase 2 metabolism |
| PICO/PECO(TS) | population, intervention, exposure, comparator, outcomes, target conditions, study design |
| PPAR (x) | peroxisome proliferator-activated receptor, x types not defined here |
| PR | progesterone receptor |
| S9 | supernatant 9000, supernatant from liver homogenate fractionation by centrifuging at 9000g |
| ScR | scoping review |
| SD | Sprague-Dawley, BALB/c |
| SEM | systematic evidence map |
| SM | supplementary manuscript |
| SR | systematic review |
| SULT | Sulfurtransferases |
| TH | thyroid hormone |
| Tox | toxicology |
| TR | thyroid receptor |
| Tris | Tris-HCl |
| TTR | Transthyretin |
| UDPGA | uridine 5'-diphospho-glucuronic acid |
| UGT | Uridine 5'-diphospho-glucuronosyltransferase |
| Vet | veterinary sciences |
| w/o | without |
| Wi | Wistar |
| wob | w/o BTS |
| woc | w/o cofactors |
| XenMet | xenobiotic metabolism |
| xPO4 | phosphate buffer, X stands for various cations used |
