## Supplementary material for "On the utilisation and characterisation of external biotransformation systems in *in vitro* toxicology: a critical review of the scientific literature with guidance recommendations": BTS reporting checklist

| BTS reporting checklist | |
|---|---|
| Reporting category | Check/details |
| **I. BTS characterisation** | |
| I. Determine BTS origin (internally produced or externally purchased). | |
| I. A) Define the type of BTS, e.g., microsomal or S9. | |
| I. B) Define the species of BTS origin. | |
| I. C) Define the strain of the above species, if applicable. | |
| I. D) Provide details on BTS pooling procedures (number of individuals and sex). | |
| I. E) Define the biotransformation-inducing chemical agents. State if no chemical induction was performed. | |
| **II. BTS reaction components** | |
| II. Clearly define all reaction components and their respective concentrations within the “final BTS reaction mixture”. | |
| II. A) Define the BTS buffer system in which the final BTS reaction takes place. | |
| II. B) Specify the utilised solvent(s) for chemical exposure, especially the solvent concentrations within the “final BTS reaction mixture”. | |
| II. C) Provide the S9 or microsomal fraction protein concentration in mg/mL, not percentages. | |
| II. D) Name the utilised cofactors and define their molar concentrations. | |
| II. E) Define, report, or measure BTS enzymatic activity. | |
| **III. BTS experimental setup** | |
| III. A) Define BTS incubation time and temperature. | |
| III. B) Define BTS-related controls. | |
